# Age-dependent genetic variants associated with longitudinal changes in brain structure across the lifespan

**DOI:** 10.1101/2020.04.24.031138

**Authors:** Rachel M. Brouwer, Marieke Klein, Katrina L. Grasby, Hugo G. Schnack, Neda Jahanshad, Jalmar Teeuw, Sophia I. Thomopoulos, Emma Sprooten, Carol E. Franz, Nitin Gogtay, William S. Kremen, Matthew S. Panizzon, Loes M. Olde Loohuis, Christopher D. Whelan, Moji Aghajani, Clara Alloza, Dag Alnæs, Eric Artiges, Rosa Ayesa-Arriola, Gareth J. Barker, Mark E. Bastin, Elisabet Blok, Erlend Bøen, Isabella A. Breukelaar, Joanna K. Bright, Elizabeth E. L. Buimer, Robin Bülow, Dara M. Cannon, Simone Ciufolini, Nicolas A. Crossley, Christienne G. Damatac, Paola Dazzan, Casper L. de Mol, Sonja M. C. de Zwarte, Sylvane Desrivières, Covadonga M. Díaz-Caneja, Nhat Trung Doan, Katharina Dohm, Juliane H. Fröhner, Janik Goltermann, Antoine Grigis, Dominik Grotegerd, Laura K. M. Han, Mathew A. Harris, Catharina A. Hartman, Sarah J. Heany, Walter Heindel, Dirk J. Heslenfeld, Sarah Hohmann, Bernd Ittermann, Philip R. Jansen, Joost Janssen, Tianye Jia, Jiyang Jiang, Christiane Jockwitz, Temmuz Karali, Daniel Keeser, Martijn G. J. C. Koevoets, Rhoshel K. Lenroot, Berend Malchow, René C. W. Mandl, Vicente Medel, Susanne Meinert, Catherine A. Morgan, Thomas W. Mühleisen, Leila Nabulsi, Nils Opel, Víctor Ortiz-García de la Foz, Bronwyn J. Overs, Marie-Laure Paillère Martinot, Erin B. Quinlan, Ronny Redlich, Tiago Reis Marques, Jonathan Repple, Gloria Roberts, Gennady V. Roshchupkin, Nikita Setiaman, Elena Shumskaya, Frederike Stein, Gustavo Sudre, Shun Takahashi, Anbupalam Thalamuthu, Diana Tordesillas-Gutiérrez, Aad van der Lugt, Neeltje E. M. van Haren, Joanna M. Wardlaw, Wei Wen, Henk-Jan Westeneng, Katharina Wittfeld, Alyssa H. Zhu, Andre Zugman, Nicola J. Armstrong, Gaia Bonfiglio, Janita Bralten, Shareefa Dalvie, Gail Davies, Marta Di Forti, Linda Ding, Gary Donohoe, Andreas J. Forstner, Javier Gonzalez-Peñas, Joao P. O. F. T. Guimaraes, Georg Homuth, Jouke-Jan Hottenga, Maria J. Knol, John B. J. Kwok, Stephanie Le Hellard, Karen A. Mather, Yuri Milaneschi, Derek W. Morris, Markus M. Nöthen, Sergi Papiol, Marcella Rietschel, Marcos L. Santoro, Vidar M. Steen, Jason L. Stein, Fabian Streit, Rick M. Tankard, Alexander Teumer, Dennis van ‘t Ent, Dennis van der Meer, Kristel R. van Eijk, Evangelos Vassos, Javier Vázquez-Bourgon, Stephanie H. Witt, Alzheimer’s Disease Neuroimaging Initiative, the IMAGEN consortium, Hieab H. H. Adams, Ingrid Agartz, David Ames, Katrin Amunts, Ole A. Andreassen, Celso Arango, Tobias Banaschewski, Bernhard T. Baune, Sintia I. Belangero, Arun L. W. Bokde, Dorret I. Boomsma, Rodrigo A. Bressan, Henry Brodaty, Jan K. Buitelaar, Wiepke Cahn, Svenja Caspers, Sven Cichon, Benedicto Crespo Facorro, Simon R. Cox, Udo Dannlowski, Torbjørn Elvsåshagen, Thomas Espeseth, Peter G. Falkai, Simon E. Fisher, Herta Flor, Janice M. Fullerton, Hugh Garavan, Penny A. Gowland, Hans J. Grabe, Tim Hahn, Andreas Heinz, Manon Hillegers, Jacqueline Hoare, Pieter J. Hoekstra, Mohammad A. Ikram, Andrea P. Jackowski, Andreas Jansen, Erik G. Jönsson, Rene S. Kahn, Tilo Kircher, Mayuresh S. Korgaonkar, Axel Krug, Herve Lemaitre, Ulrik F. Malt, Jean-Luc Martinot, Colm McDonald, Philip B. Mitchell, Ryan L. Muetzel, Robin M. Murray, Frauke Nees, Igor Nenadic, Jaap Oosterlaan, Roel A. Ophoff, Pedro M. Pan, Brenda W. J. H. Penninx, Luise Poustka, Perminder S. Sachdev, Giovanni A. Salum, Peter R. Schofield, Gunter Schumann, Philip Shaw, Kang Sim, Michael N. Smolka, Dan J. Stein, Julian Trollor, Leonard H. van den Berg, Jan H. Veldink, Henrik Walter, Lars T. Westlye, Robert Whelan, Tonya White, Margaret J. Wright, Sarah E. Medland, Barbara Franke, Paul M. Thompson, Hilleke E. Hulshoff Pol

**Affiliations:** Department of Psychiatry, University Medical Center Utrecht Brain Center, Utrecht University, Utrecht, The Netherlands; Department of Complex Trait Genetics, Center for Neurogenomics and Cognitive Research, Amsterdam Neuroscience, VU Amsterdam, Amsterdam, The Netherlands; Department of Psychiatry, University of California San Diego, La Jolla, CA, USA; Department of Human Genetics, Radboud University Medical Center, Nijmegen, The Netherlands; Donders Institute for Brain, Cognition and Behaviour, Radboud University, Nijmegen, The Netherlands; Psychiatric Genetics, QIMR Berghofer Medical Research Institute, Brisbane, QLD, Australia; Imaging Genetics Center, Mark and Mary Stevens Neuroimaging and Informatics Institute, Keck School of Medicine, University of Southern California, Marina del Rey, CA, USA; Department of Cognitive Neuroscience, Donders Institute for Brain, Cognition and Behaviour, Radboud University Medical Center, Nijmegen, The Netherlands; Department of Psychiatry and Center for Behavior Genetics of Aging, University of California San Diego, La Jolla, CA, USA; American Psychiatric Association, Washington, DC, USA; VA San Diego Center of Excellence for Stress and Mental Health, San Diego, CA, USA; Center for Neurobehavioral Genetics, University of California, Los Angeles, Los Angeles, CA, USA; Biogen Research and Development, Cambridge, MA, USA; Department of Psychiatry, Amsterdam Public Health and Amsterdam Neuroscience, Amsterdam UMC, Vrije Universiteit & GGZinGeest, Amsterdam, The Netherlands; Department of Child and Adolescent Psychiatry, Institute of Psychiatry and Mental Health, Hospital General Universitario Gregorio Marañón, IiSGM, CIBERSAM, School of Medicine, Universidad Complutense, Madrid, Spain; NORMENT Centre, University of Oslo, Oslo, Norway; Division of Mental Health and Addiction, Oslo University Hospital, Oslo, Norway; INSERM U A10 1000 Trajectoires Développementales en Psychiatrie, Université Paris-Saclay, Ecole Normale Supérieure Paris-Saclay, CNRS, Centre Borelli, Gif-sur-Yvette, France; Valdecilla Biomedical Research Institute (IDIVAL), Marqués de Valdecilla University Hospital (HUMV), School of Medicine, University of Cantabria, Santander, Spain; Department of Neuroimaging, King’s College London, London, UK; Lothian Birth Cohorts group, Department of Psychology, University of Edinburgh, Edinburgh, UK; Centre for Clinical Brain Sciences, University of Edinburgh, Edinburgh, UK; Edinburgh Imaging, University of Edinburgh, Edinburgh, UK; Department of Child and Adolescent Psychiatry/Psychology, Sophia Children’s Hospital, Erasmus University Medical Centre Rotterdam, Rotterdam, The Netherlands; Psychosomatic and CL Psychiatry, Oslo University Hospital, Oslo, Norway; Brain Dynamics Centre, Westmead Institute for Medical Research, The University of Sydney, Westmead, NSW, Australia; Institute of Diagnostic Radiology and Neuroradiology, University Medicine Greifswald, Greifswald, Germany; Centre for Neuroimaging & Cognitive Genomics (NICOG), Clinical Neuroimaging Laboratory, NCBES Galway Neuroscience Centre, College of Medicine Nursing and Health Sciences, National University of Ireland Galway, Galway, Ireland; Department of Psychosis Studies, Institute of Psychiatry, Psychology and Neuroscience, King’s College London, London, UK; Department of Psychiatry, School of Medicine, Pontificia Universidad Católica de Chile, Santiago, Chile; Department of Psychological Medicine, Institute of Psychiatry, Psychology and Neuroscience, King’s College London, London, UK; Department of Neurology, Erasmus University Medical Centre Rotterdam, Rotterdam, The Netherlands; Social, Genetic and Developmental Psychiatry Centre, Institute of Psychiatry, Psychology & Neuroscience, King’s College London, London, UK; Institute for Translational Psychiatry, University of Münster, Münster, Germany; Section of Systems Neuroscience Department of Psychiatry and Psychotherapy Technische Universität Dresden, Dresden, Germany; Université Paris-Saclay, CEA, Neurospin, Gif-sur-Yvette, France; University of Groningen, University Medical Center Groningen, Department of Psychiatry, Interdisciplinary Center Psychopathology and Emotion regulation (ICPE), Groningen, The Netherlands; Department of Psychiatry and Mental Health, University of Cape Town, Cape Town, South Africa; Institute of Clinical Radiology, University and University Hospital Münster, Münster, Germany; Departments of Experimental and Clinical Psychology, Amsterdam, The Netherlands; Department of Child and Adolescent Psychiatry and Psychotherapy, Central Institute of Mental Health, Medical Faculty Mannheim/Heidelberg University, Mannheim, Germany; Physikalisch-Technische Bundesanstalt (PTB), Berlin, Germany; Department of Clinical Genetics, VUmc, Amsterdam UMC, Amsterdam, The Netherlands; Centre for Population Neuroscience and Precision Medicine (PONS), Institute of Science and Technology for Brain-Inspired Intelligence and MoE Key Laboratory of Computational Neuroscience and Brain-Inspired Intelligence, Fudan University, Shanghai, China; Centre for Population Neuroscience and Precision Medicine (PONS), Institute of Psychiatry, Psychology and Neuroscience, SGDP Centre, King’s College London, London, UK; Centre for Healthy Brain Ageing (CHeBA), School of Psychiatry, University of New South Wales, Sydney, NSW, Australia; Institute of Neuroscience and Medicine (INM-1), Research Centre Jülich, Jülich, Germany; Department of Psychiatry, Psychotherapy and Psychosomatics, RWTH Aachen University, Medical Faculty, Aachen, Germany, Aachen, Germany; Department of Psychiatry and Psychotherapy, University Hospital LMU, Munich, Germany; Department of Radiology, University Hospital LMU, Munich, Germany; Munich Center for Neurosciences (MCN) – Brain & Mind, Planegg-Martinsried, Germany; School of Psychiatry, University of New South Wales, Sydney, NSW, Australia; School of Psychiatry and Behavioral Sciences, School of Medicine, University of New Mexico, Albuquerque, NM, USA; Neuroscience Research Australia, Sydney, NSW, Australia; Department of Psychiatry and Psychotherapy, University Medical Center Göttingen, Göttingen, Germany; School of Psychology and Centre for Brain Research, The University of Auckland, Auckland, New Zealand; Brain Research New Zealand, New Zealand; Cécile and Oskar Vogt Institute for Brain Research, Medical Faculty, University Hospital Düsseldorf, Heinrich Heine University Düsseldorf, Düsseldorf, Germany; Department of Biomedicine, University of Basel, Basel, Switzerland; Neuroimaging Unit, Technological Facilities, Valdecilla Biomedical Research Institute IDIVAL, Santander, Spain; INSERM U1299 Trajectoires Développementales en Psychiatrie, Centre Borelli UMR9010, Ecole Normale Supérieure Paris-Saclay, CNRS, Centre Borelli, Gif-sur-Yvette, France; Department of Mental Health, Institute of Translational Psychiatry, The University of Münster, Münster, Germany; Psychiatric Imaging Group, MRC London Institute of Medical Sciences (LMS), Imperial College London, London, UK; Department of Epidemiology, Erasmus University Medical Centre Rotterdam, Rotterdam, The Netherlands; Department of Radiology & Nuclear Medicine, Erasmus MC, University Medical Center Rotterdam, Rotterdam, The Netherlands; Department of Psychiatry and Psychotherapy, Philipps-University Marburg, Marburg, Germany; Social and Behavioral Research Branch, National Human Genome Research Institute, Bethesda, MD, USA; Department of Neuropsychiatry, Wakayama Medical University, Wakayama, Japan; CIBERSAM, Biomedical Research Network on Mental Health Area, Santander, Spain; Centre for Clinical Brain Sciences and UK Dementia Research Institute Centre, University of Edinburgh, Edinburgh, UK; Department of Neurology, University Medical Center Utrecht Brain Center, Utrecht University, Utrecht, The Netherlands; German Center for Neurodegenerative Diseases (DZNE), Site Rostock/Greifswald, Greifswald, Germany; Department of Psychiatry and Psychotherapy, University Medicine Greifswald, Greifswald, Germany; Laboratory of Integrative Neuroscience (LiNC), Department of Psychiatry, Universidade Federal de São Paulo (UNIFESP), São Paulo, SP, Brazil; National Institute of Developmental Psychiatry for Children and Adolescents (INPD), CNPq, São Paulo, SP, Brazil; Mathematics and Statistics, Curtin University, Perth, WA, Australia; Department of Psychology, University of Edinburgh, Edinburgh, UK; Centre for Neuroimaging & Cognitive Genomics (NICOG), School of Psychology and Discipline of Biochemistry, National University of Ireland Galway, Galway, Ireland; Centre for Human Genetics, Philipps-University Marburg, Marburg, Germany; Institute of Human Genetics, University of Bonn, School of Medicine & University Hospital Bonn, Bonn, Germany; Interfaculty Institute for Genetics and Functional Genomics, University Medicine Greifswald, Greifswald, Germany; Department of Biological Psychology, Vrije Universiteit Amsterdam, Amsterdam, The Netherlands; Faculty of Medicine and Health, The University of Sydney, Sydney, NSW, Australia; School of Medical Sciences, University of New South Wales, Sydney, NSW, Australia; NORMENT Centre of Excellence, Department of Clinical Science, University of Bergen, Bergen, Norway; Dr. Einar Martens Research Group for Biological Psychiatry, Department of Medical Genetics, Haukeland University Hospital, Bergen, Norway; Institute of Psychiatric Phenomics and Genomics (IPPG), University Hospital LMU, Munich, Germany; Department of Genetic Epidemiology, Central Institute of Mental Health, Medical Faculty Mannheim, Heidelberg University, Mannheim, Germany; Department of Morphology and Genetics, Universidade Federal de São Paulo (UNIFESP), São Paulo, SP, Brazil; Department of Genetics & UNC Neuroscience Center, University of North Carolina at Chapel Hill, Chapel Hill, NC, USA; Institute for Community Medicine, University Medicine Greifswald, Greifswald, Germany; School of Mental Health and Neuroscience, Faculty of Health, Medicine and Life Sciences, Maastricht University, Maastricht, The Netherlands; NIHR Maudsley Biomedical Research Centre, South London and Maudsley NHS Trust, London, UK; Department of Psychiatry, Marqués de Valdecilla University Hospital, Valdecilla Biomedical Research Institute (IDIVAL), Santander, Spain; Department of Clinical Genetics, Erasmus University Medical Centre Rotterdam, Rotterdam, The Netherlands; Centre for Psychiatry Research, Department of Clinical Neuroscience, Karolinska Institutet, & Stockholm Health Care Services, Stockholm County Council, Stockholm, Sweden; Department of Psychiatric Research, Diakonhjemmet Hospital, Oslo, Norway; Academic Unit for Psychiatry of Old Age, University of Melbourne, Parkville, VIC, Australia; National Ageing Research Institute, Parkville, VIC, Australia; Department of Psychiatry, The University of Melbourne, Melbourne, VIC, Australia; Florey Institute of Neuroscience and Mental Health, The University of Melbourne, Melbourne, VIC, Australia; Discipline of Psychiatry and Trinity College Institute of Neuroscience, Trinity College Dublin, Dublin, Ireland; Netherlands Twin Register, Department of Biological Psychology, Vrije Universiteit, Amsterdam, The Netherlands; Instituto Ame Sua Mente, São Paulo, SP, Brazil; Karakter Child and Adolescent Psychiatry University Centre, Nijmegen, The Netherlands; Altrecht Science, Altrecht Mental Health Institute, Utrecht, The Netherlands; Institute for Anatomy I, Medical Faculty, Heinrich Heine University Düsseldorf, Düsseldorf, Germany; Institute of Medical Genetics and Pathology, University Hospital Basel, University of Basel, Basel, Switzerland; Department of Psychiatry, Virgen del Rocio University Hospital, School of Medicine, University of Seville, IBIS, Seville,Spain; Scottish Imaging Network, A Platform for Scientific Excellence (SINAPSE) Collaboration, Edinburgh, UK; NORMENT Centre, Oslo University Hospital, Oslo, Norway; Department of Neurology, Oslo University Hospital, Oslo, Norway; Institute of Clinical Medicine, University of Oslo, Oslo, Norway; Department of Psychology, University of Oslo, Oslo, Norway; Bjørknes College, Oslo, Norway; Language and Genetics Department, Max Planck Institute for Psycholinguistics, Nijmegen, The Netherlands; Department of Cognitive and Clinical Neuroscience, Central Institute of Mental Health, Medical Faculty Mannheim, Heidelberg University, Mannheim, Germany; Department of Psychiatry, University of Vermont, Burlington, VT, USA; Sir Peter Mansfield Imaging Centre, School of Physics and Astronomy, University of Nottingham, Nottingham, UK; Charité Universitätsmedizin Berlin, Berlin, Germany; Department of Child and Adolescent Psychiatry, Groningen, The Netherlands; Core-Facility Brainimaging, Faculty of Medicine, University of Marburg, Marburg, Germany; Department of Psychiatry, Icahn School of Medicine at Mount Sinai, New York, NY, USA; VISN 2 Mental Illness Research, Education & Clinical Center (MIRECC) James J. Peters Department of Veterans Affairs Medical Center, Bronx, NY, USA; Department of Psychiatry and Psychotherapy, University of Bonn, Bonn, Germany; Groupe d’Imagerie Neurofonctionnelle, Institut des Maladies Neurodégénératives, CNRS UMR 5293, Université de Bordeaux, Centre Broca Nouvelle-Aquitaine, Bordeaux, France; Unit for Psychosomatic Medicine and C-L Psychiatry, University of Oslo, Oslo, Norway; Black Dog Institute, Sydney, NSW, Australia; Institute of Medical Psychology and Medical Sociology, University Medical Center Schleswig-Holstein, Kiel University, Kiel, Germany; Emma Children’s Hospital, Amsterdam UMC, University of Amsterdam, Emma Neuroscience Group, department of Pediatrics, Amsterdam Reproduction & Development, Amsterdam, The Netherlands, Amsterdam, The Netherlands; Department of Psychiatry, Erasmus Medical Center, Erasmus University, Rotterdam, The Netherlands; Department of Child and Adolescent Psychiatry, University Medical Center Goettingen, Germany, Göttingen, Germany; Neuropsychiatric Institute, The Prince of Wales Hospital, Sydney, NSW, Australia; Department of Psychiatry and Legal Medicine, Universidade Federal do Rio Grande do Sul, Porto Alegre, RS, Brazil; Section on Negative Affect and Social Processes, Hospital de Clínicas de Porto Alegre, Porto Alegre, RS, Brazil; Fudan-KCL PONS Centre, Institute for Science and Technology for Brain-inspired Intelligence (ISTBI), Fudan University, Shanghai, China; PONS Research Group, Department of Psychiatry and Psychotherapy, Charite, Humboldt University, Berlin, Germany; National Institute of Mental Health, National Institutes of Health, Bethesda, MD, USA; West Region, Institute of Mental Health, Singapore; Yong Loo Lin School of Medicine, National University of Singapore, Singapore; Department of Psychiatry and Neuroimaging Center, Technische Universität Dresden, Dresden, Germany; SAMRC Unit on Risk & Resilience in Mental Disorders, Department of Psychiatry & Neuroscience Institute, University of Cape Town, Cape Town, South Africa; Department of Developmental Disability Neuropsychiatry, School of Psychiatry, UNSW Sydney, Sydney, NSW, Australia; Charité Universitätsmedizin Berlin, corporate member of Freie Universität Berlin, Humboldt-Universität zu Berlin, and Berlin Institute ofr Health, Berlin, Germany; Trinity College Institute of Neuroscience, Trinity College Dublin, Dublin, Ireland; Queensland Brain Institute, The University of Queensland, Brisbane, QLD, Australia; Centre for Advanced Imaging, The University of Queensland, Brisbane, QLD, Australia; QIMR Berghofer Medical Research Institute, Brisbane, QLD, Australia; Department of Psychiatry, Radboud University Medical Center, Nijmegen, The Netherlands

## Abstract

Human brain structure changes throughout our lives. Altered brain growth or rates of decline are implicated in a vast range of psychiatric, developmental, and neurodegenerative diseases. Here, we identified common genetic variants that affect rates of brain growth or atrophy, in the first genome-wide association meta-analysis of changes in brain morphology across the lifespan. Longitudinal MRI data from 15,640 individuals were used to compute rates of change for 15 brain structures. The most robustly identified genes *GPR139, DACH1* and *APOE* are associated with metabolic processes. We demonstrate global genetic overlap with depression, schizophrenia, cognitive functioning, insomnia, height, body mass index and smoking. Gene-set findings implicate both early brain development and neurodegenerative processes in the rates of brain changes. Identifying variants involved in structural brain changes may help to determine biological pathways underlying optimal and dysfunctional brain development and ageing.

## Main

Under the influence of genes and a varying environment, human brain structure changes throughout the lifespan. Even in adulthood, when the brain seems relatively stable, individuals differ in the profile and rate of brain changes^1^. Longitudinal studies are crucial to identify genetic and environmental factors that influence the rate of these brain changes throughout development^2^ and ageing^3^. Inter-individual differences in brain development are associated with general cognitive function^4,5^, and risk for psychiatric disorders^6,7^ and neurological diseases^8,9^. Genetic factors involved in brain development and ageing overlap with those for cognition^10^ and risk for neuropsychiatric disorders^11^. A recent cross-sectional study showed a genetic component to advanced brain age in several brain disorders^12^. Yet, we still lack information on which genetic variants influence individual brain changes throughout life, since this requires longitudinal data. Discovering genetic factors for brain changes may reveal key biological pathways that drive normal development and ageing, and may contribute to identifying disease risk and resilience: a crucial goal given the urgent need for new treatments for aberrant brain development and ageing worldwide.

As part of the Enhancing Imaging Genetics through Meta-Analysis (ENIGMA) consortium^13^ the ENIGMA Plasticity Working Group recently quantified the overall genetic contribution to longitudinal brain changes by combining evidence from multiple twin cohorts across the world^14^. Most global and subcortical brain measures showed genetic influences on change over time, with a higher genetic contribution in the elderly (heritability 16 – 42%). Genetic factors that influence longitudinal changes were partially independent of those that influence baseline volumes of brain structures, suggesting that there might be genetic variants that specifically affect the rate of development or ageing. Even so, the genes involved in these processes are still not known. So far, only a single, small-scale genome-wide association study (GWAS) was performed for brain change^15^. Here, we set out to find genetic variants that may influence rates of brain changes over time, using genome-wide analysis in individuals scanned with magnetic resonance imaging (MRI) on more than one occasion. We also aimed to identify age-dependent effects of genomic variation on longitudinal brain changes in mostly healthy, but also neurological and psychiatric, populations.

In our GWAS meta-analysis, we sought genetic loci associated with annual change rates in 8 global and 7 subcortical morphological brain measures in a coordinated two-phased analysis using data from 40 longitudinal cohorts. Global and subcortical brain measures were extracted, and annual change rates were analysed using additive genetic association analyses to estimate effects of genetic variants on rates of change within each cohort. As brain change is not constant over age^1^ and gene expression also changes during development and ageing^16^, we determined whether the estimated genetic variants were age-dependent, i.e., differentially affected rates of brain changes at different stages of life using genome-wide meta-regression models with linear or quadratic age effects (Methods). We employed a rolling cumulative meta-analysis and -regression approach^17^. In phase 1, we analysed the cohorts from European descent (N=9,623). We sought replication by adding three additional cohorts that made their data available after the first analysis: one developmental and two in ageing populations (N =5,477; all European descent; total N=15,100 in phase 2). In all follow-up analyses, results from this second phase were used. Finally, we added cohorts from non-European ancestry (total N=15,640).

## Longitudinal trajectories

Change in global brain measures showed different trajectories of change with age (Figure 1 and Extended Data Movie 1), characterized by either monotonic increases (lateral ventricles), monotonic decreases (cortex volume, cerebellar grey matter volume, cortical thickness, surface area, total brain volume), or increases followed by stabilization and subsequently decreases (cerebral and cerebellar white matter, thalamus, caudate, putamen, nucleus accumbens, pallidum, hippocampus and amygdala volumes). Each brain structure showed a characteristic trajectory of change. Rates of change within individuals generally showed low correlations in both childhood and older age in our data (Extended Data Fig. 2), with the exception of change rates of cortical thickness with cortex volume. Therefore, we chose to investigate all brain structures separately, maximizing sensitivity of the GWAS for identification of region-specific associations of genetic variants. Using the correlation structure, we estimated the effective number of independent variables through matrix spectral decomposition on the rates of change yielding 14 independent traits for multiple testing corrections (Methods).

**Figure 1:**
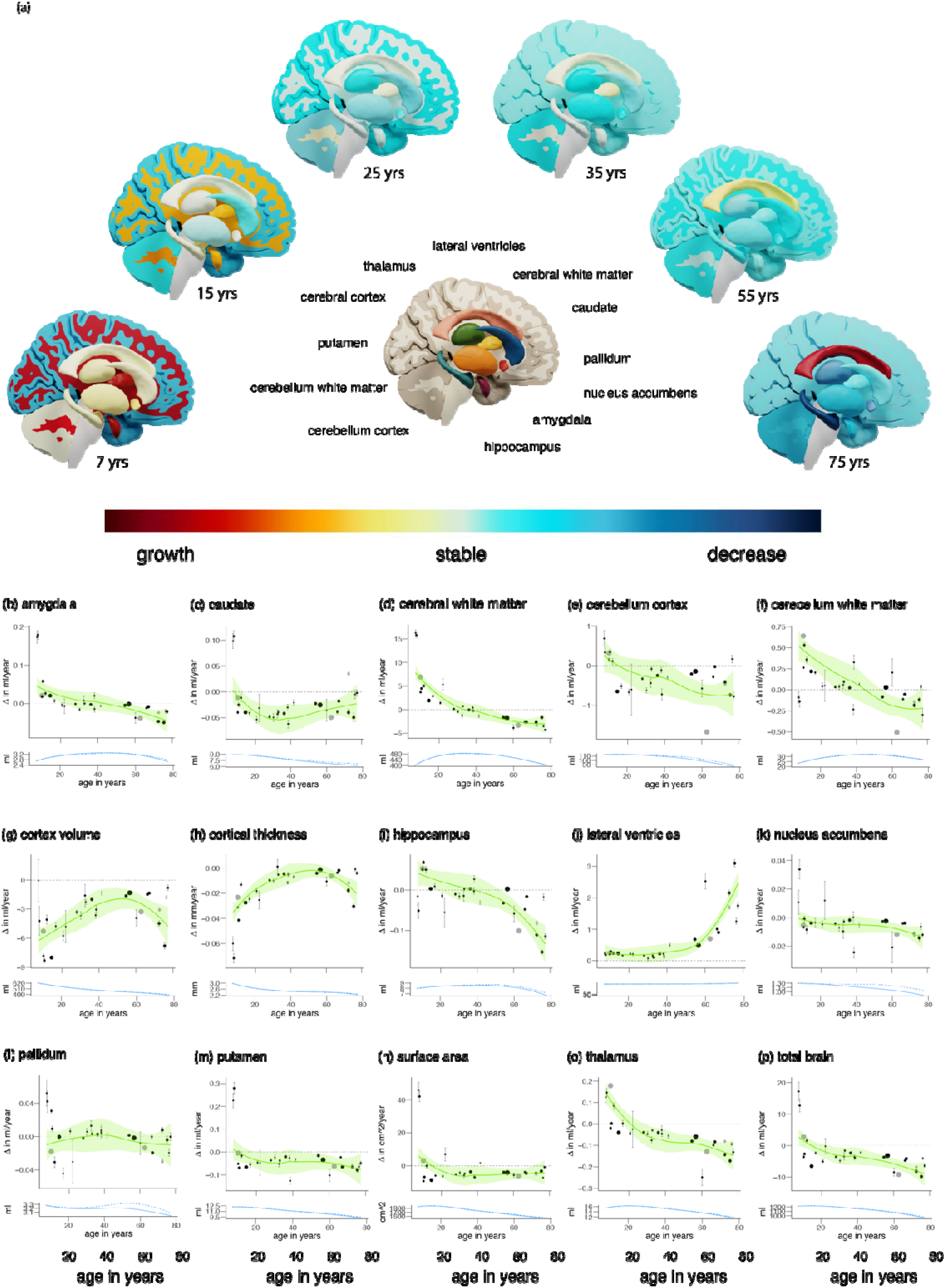
Phenotypic brain changes throughout the lifespan. Visualization of growth and decline of brain structures throughout the lifespan. The subcortical structures are shown in exploded view (a). Annual rates of change Δ per cohort for each structure (b-p). The estimated trajectories with confidence intervals (*in green*) are displayed in the top row. The size of the points represents the relative size of the cohorts. Standard errors are displayed in grey. Means and standard deviations are based on raw data – no covariates were included. Cohorts that were added in phase2 are displayed in grey. Only cohorts that satisfy N>75 and mean interval > 0.5 years are shown. The estimated trajectories of the volumes themselves are displayed in the bottom row, for all subjects (*solid line*) and for subjects not part of diagnostic groups (*dashed line*).

## Age-independent associations

Two loci showed genome-wide significant effects on the rate of brain change in phase 1, one of which was also genome-wide significant in phase 2 (Figure 2; Supplementary Table 4). This lead SNP, rs72772740 on chromosome 16, is an intronic variant located in the *GPR139* gene and was associated with change in lateral ventricle volume (Figure 3). Functional annotation identified numerous significant expression quantitative trait loci (eQTL) associations (FDR < 0.05) in different datasets and highlighted genes by either eQTL mapping (*GPRC5B, IQCK, KNOP1, C16orf62*) or chromatin interaction mapping (*ACSM1, ACSM5, UMOD, GP2). GPR139* is the G-protein-coupling receptor gene 139, which encodes a member of the rhodopsin family of G-protein coupled receptors. The gene is almost exclusively expressed in the central nervous system, with highest expression from 12 to 26 weeks post-conception, and has been suggested as a therapeutic target for metabolic syndromes and motor diseases^18^. *GPR139* may play a role in foetal brain development^19^ and mice lacking *GPR139* exhibited schizophrenia-like symptomatology^20^. Additionally, functional cell-assays confirmed the inhibitory influence of *GPR139* on dopamine receptor 2 (D2R) signalling^20^. The second lead SNP, rs449998, an intronic variant on chromosome 21 located in the Down Syndrome Cell Adhesion Molecule (*DSCAM*) gene, was associated with change in nucleus accumbens volume but was not significant in phase 2. Three SNPs (intergenic SNP rs10990953 on chr 2, associated with rate of change in lateral ventricles; intergenic and located in long intergenic non-protein coding RNA SNP rs1425034 on chr 2, associated with rate of change in pallidum; and rs1425034, intron of *CDH8* on chr 16, associated with rate of change in total brain volume) were significant in the phase 2 analysis only (Supplementary Table 5; Extended Data Figs. 3,4 provide Manhattan plots, QQ plots, locus plots and circos plots). The *CDH8* association with total brain volume change is particularly interesting, since it has been associated earlier with learning disability and autism, with macrocephaly as a risk factor^21^. *CDH8* is a protein coding gene and encodes a type II classical cadherin from the cadherin superfamily, integral membrane proteins that mediate calcium-dependent cell-cell adhesion. Genome-wide significant SNPs in phase 1 or phase 2 did not show heterogeneity (I2 < 10.2; p(I2) > 0.31; Supplementary Tables 4,5, Extended Data Fig 5 for forest plots).

**Figure 2:**
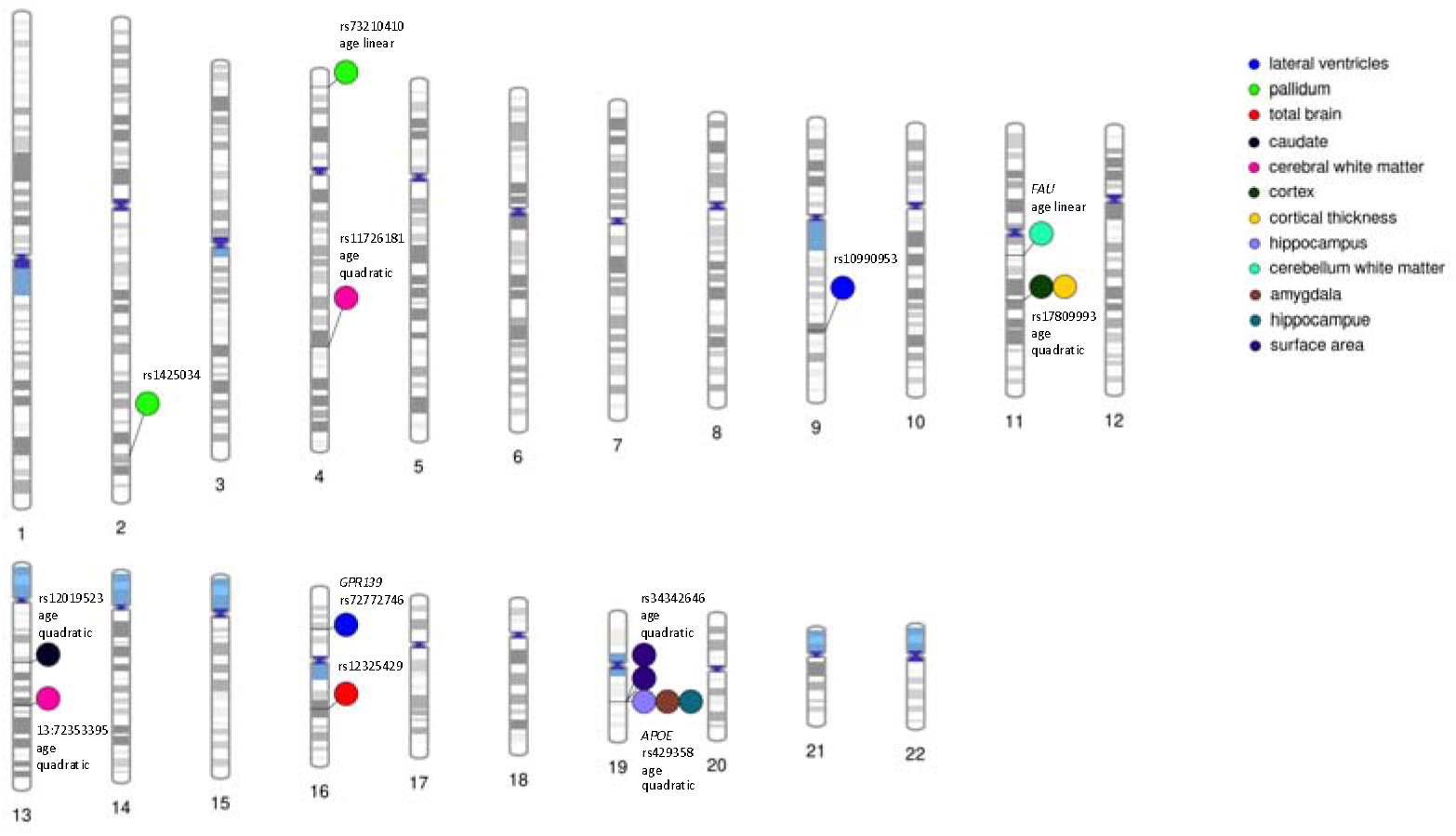
Genetic effects on rates of brain changes throughout the lifespan. Genome-wide significant SNPs and genes with effects on brain changes at their respective loci across the human genome. This plot was created using PhenoGram (http://visualization.ritchielab.org).

**Figure 3:**
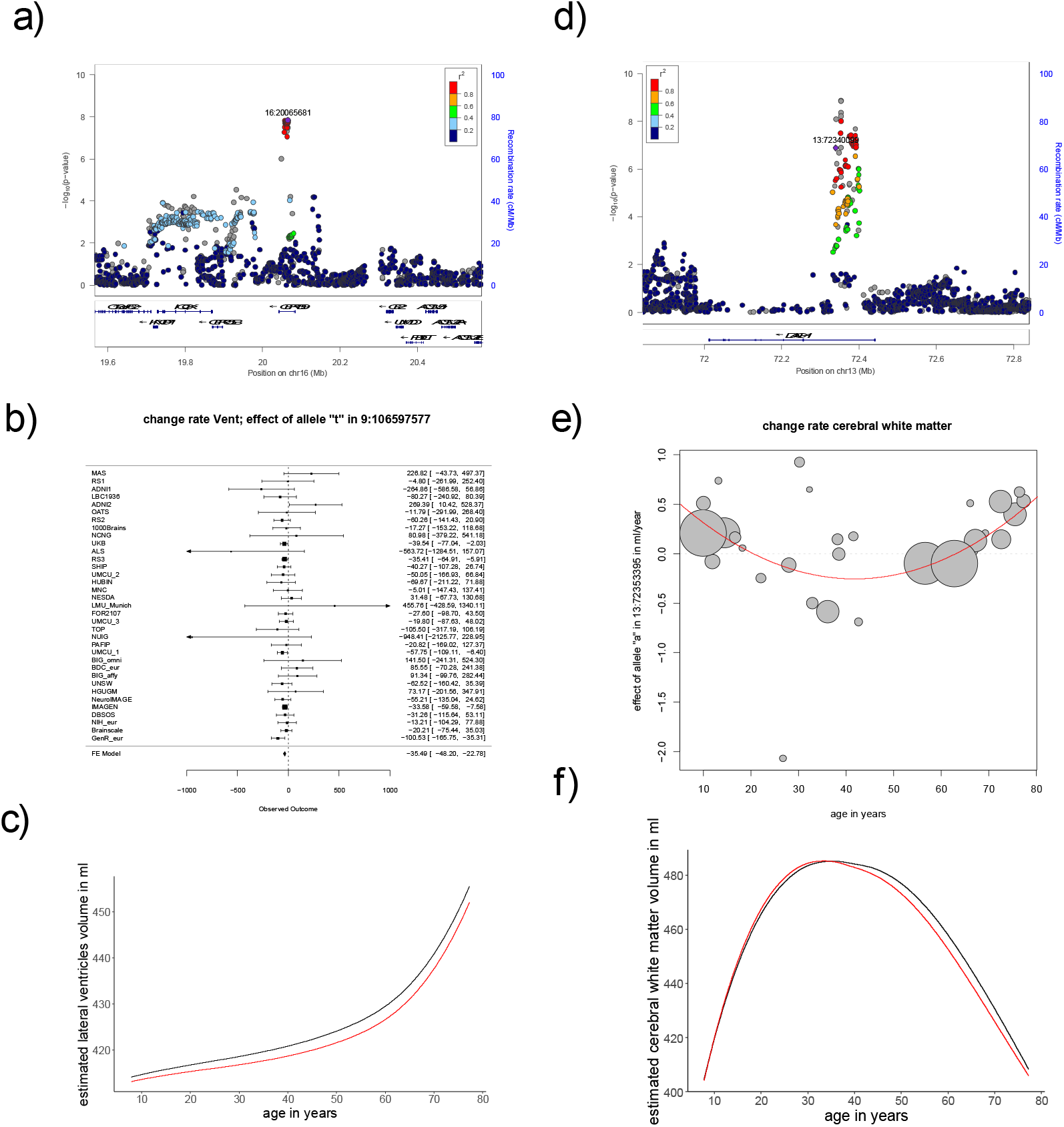
Summary of findings for two top-SNPs. Shown here is a summary of findings for a top-SNP of an age independent effect (rs72772746; intron to *GPR139*; associated with rate of change of lateral ventricle volume; left column) and a top-SNP of an age dependent effect (13:72353395; intron to *DACH1*; associated with rate of change in cerebral white matter volume; right column). Displayed are the locus plots (a) and (d), forest plot (b) and plot of meta-regression (e) and inferred lifespan trajectories for carriers (in red) and non-carriers of the effect allele (in black) (c) and (f). Note that 13:72353395 was not in the reference dataset containing LD structure; the displayed LD structure is based on 13:7234009, R2 = 0.87 with the top-SNP.

## Age-dependent associations

The association of three additional loci with rate of change was variable across the lifespan in phase 1 (Figure 2; Supplementary Tables 6,8), two of which remained significant in the phase 2 analysis. White matter cerebrum volume change was affected by rs573983368 (intronic variant) in the Dachshund Family Transcription Factor 1 (*DACH1*) gene, and rs6864758 (intergenic and located in long intergenic non-protein coding RNA LINC02227) on chromosome 5 had an age-dependent effect on the change in surface area (Figure 3; Supplementary Tables 6-9). White matter cerebellum volume change was affected by the intronic rs10674957 in the Thyrotropin Releasing Hormone Degrading Enzyme (*TRHDE*) gene, but this third locus was not significant in phase 2. The *DACH1* locus shows significant chromatin interaction, which can play an important role in gene expression regulation. *DACH1* encodes a chromatin-associated protein that associates with DNA-binding transcription factors to regulate gene expression and cell fate determination during development. *DACH1* is highly expressed in the proliferating neural progenitor cells of the developing cortical ventricular and subventricular regions, and in the striatum^22^. We found the effect of *DACH1* to have a quadratic age-dependence, with the variant being associated with faster growth in childhood and earlier but slower decline with ageing (Figure 3). Seven additional loci showed a significant dependency on age in phase 2 (Supplementary Tables 7,9; Extended Data Figs. 3,4 provide Manhattan plots, QQ plots, locus plots and circos plots). One of these, rs429358, a missense variant of the Alzheimer’s disease (AD)-related^23^ Apolipoprotein E gene (*APOE*) gene, was associated with change rate in hippocampus, showing faster decay of the hippocampus for carriers of the AD risk variant. *APOE* plays a role in maintenance of cellular cholesterol homeostasis by delivering cholesterol to neurons on apoE-containing lipoprotein particles. Cholesterol is important for synapse and dendrite formation, and cholesterol depletion has been shown to cause synaptic and dendritic degeneration^24^. Other findings include rs12019523, an intronic variant in the *CAB39L* gene associated with rate of change of the caudate; rs34342646, an intronic variant in the *NECTIN2* gene associated with rate of change in surface area and rs73210410, an intronic variant in the *SORCS2* gene associated with rate of change in the pallidum. To visualize the age-dependent effects, we plotted the meta-regression results for the significant loci (Methods, Extended Data Fig. 5). Genome-wide significant SNPs in phase 1 or phase 2 did not show significant residual heterogeneity (p > 0.23; except for the age-dependent effect of rs429358 on hippocampus change rate (p=0.02)). A summary of the genome-wide significant results and the top-10 loci for each phenotype and age model are presented in Supplementary Tables 4-9.

## Gene-based analyses

Gene-based associations with all phenotypes were estimated using MAGMA (Methods). We found six genome-wide significant genes influencing structural rates of change in the phase 1, four of which were also significant in phase 2 (Supplementary Table 10,11); among these, *DACH1* and *GPR139*, which were implicated through SNP-based GWAS, also reached genome-wide significance in this gene-based GWAS. In addition, we found *APOE* to be associated with change rates for both hippocampus and amygdala. The phase 2 analysis showed two new findings: an association of the *FAU* gene with rate of change in cerebellum white matter, and again *APOE*, associated with rate of change in surface area. Of note, the *APOE* findings were based on GWAS and subsequent gene analysis, and we did not investigate the classical *APOE* status, since that is determined by a combination of two SNPs. However, we observed that the effect of *APOE* on change rate of hippocampus and amygdala was fully driven by rs429358, with the risk variant for AD causing faster increases in childhood for amygdala and faster decay for both amygdala and hippocampus later in life. To visualize the age-dependent effects, we plotted the meta-regression results for the top SNP in each of the significant genes (Extended Data Fig. 5). Supplementary Tables 10,11 display the top-10 genes for each phenotype and each age model. Supplementary Table 12 details putative biological functions of associated genes and genes harbouring genome-wide significant associated loci.

## Gene-set analyses

To test whether genetic findings for brain structure change converged onto functional gene sets and pathways, we conducted gene-set analyses using MAGMA (Methods). Competitive testing was used and 10 and 12 genome-wide significant gene sets were found for phase 1 and phase 2, respectively (Supplementary Tables 13, 14 for top-10 gene sets and genes included). Two main themes emerge as biological functions of the gene sets converge onto involvement of genes in early brain development on the one hand and neurodegeneration on the other.

One gene set was significant in both the phase 1 and phase 2 analyses: GO_neural_nucleus_development is associated with genes involved in neural nucleus development and related to rates of change in cerebellar white matter volume. Two other gene sets, significant in phase 1 (GO_substantia_nigra_development associated with cerebellum white matter rate of change) and phase 2 (GO_midbrain_development associated with quadratic age-dependent surface area rates of change) were closely related to neural nucleus development in gene ontology terms.

The most significant gene set was GO_response_to_phorbol_13_acetate_12_myristate, (p-value=1.42e-08) in phase 2, related to surface area change. Phorbol 13-acetate 12-myristate is a phorbol ester and an activator of protein kinase C (PKC)^25^. Two other gene sets, significant in phase 2 (GO_tau_protein_binding and GO_tau_protein_kinase_activity, both associated with caudate change), imply genes involved in interacting with tau protein. Tau is a microtubule-associated protein, implicated in Alzheimer’s disease, Down Syndrome and amyotrophic lateral sclerosis (ALS).

## Overlap with cross-sectional findings

SNP-based heritability estimates (*h^2^*) of the rates of change based on linkage disequilibrium score regression (LDSC; Methods) were small overall (Supplementary Table 15). For all phenotypes, the h^2^ z-score was below 4, so we tested for genetic overlap with cross-sectional brain data and other phenotypes by applying approaches other than LDSC: to investigate whether cross-sectional GWAS for brain structure and our GWAS on rates of change identify the same or different genetic variants, we investigated overlap between rate of change and earlier published data on cross-sectional brain structure of the same structure, where available (Methods). Extended Data Fig. 6 displays the number of overlapping genes tested against the expected number of overlapping genes that would occur by chance, in the first 1-1,000 ranked genes. Supplementary Table S11 lists the top-10 gene findings for each of the 15 change rate phenotypes and compares these with the gene ranks from cross-sectional data. In the top-10 ranked genes, *APOE* for hippocampus occurred in the top-10 for both crosssectional data^26^ and age-dependent effects on rate of change (p=0.006). No overlap was seen for the other measured phenotypes. Extending this search to the top 200 (~1% of genes), we found overlapping genes above chance level for cortical thickness of quadratic age-dependent genes and cross-sectional findings (p = 8.39e-05). In the top 1,000 ranked genes (~5% of genes), further overlapping genes did emerge (Extended Data Fig. 6). Overlapping genes at such a high aggregate level imply that largely different genetic backgrounds underlie changes in brain structure and brain structure per se.

To test for global genomic overlap between our findings and GWAS of cross-sectional volumes we applied independent SNP-Effect Concordance Analyses (iSECA) (Methods) and tested for pleiotropy. We found no significant pleiotropy between longitudinal and cross-sectional results, confirming a largely different genetic background for changes in brain structure and brain structure per se (Figure 4).

**Figure 4:**
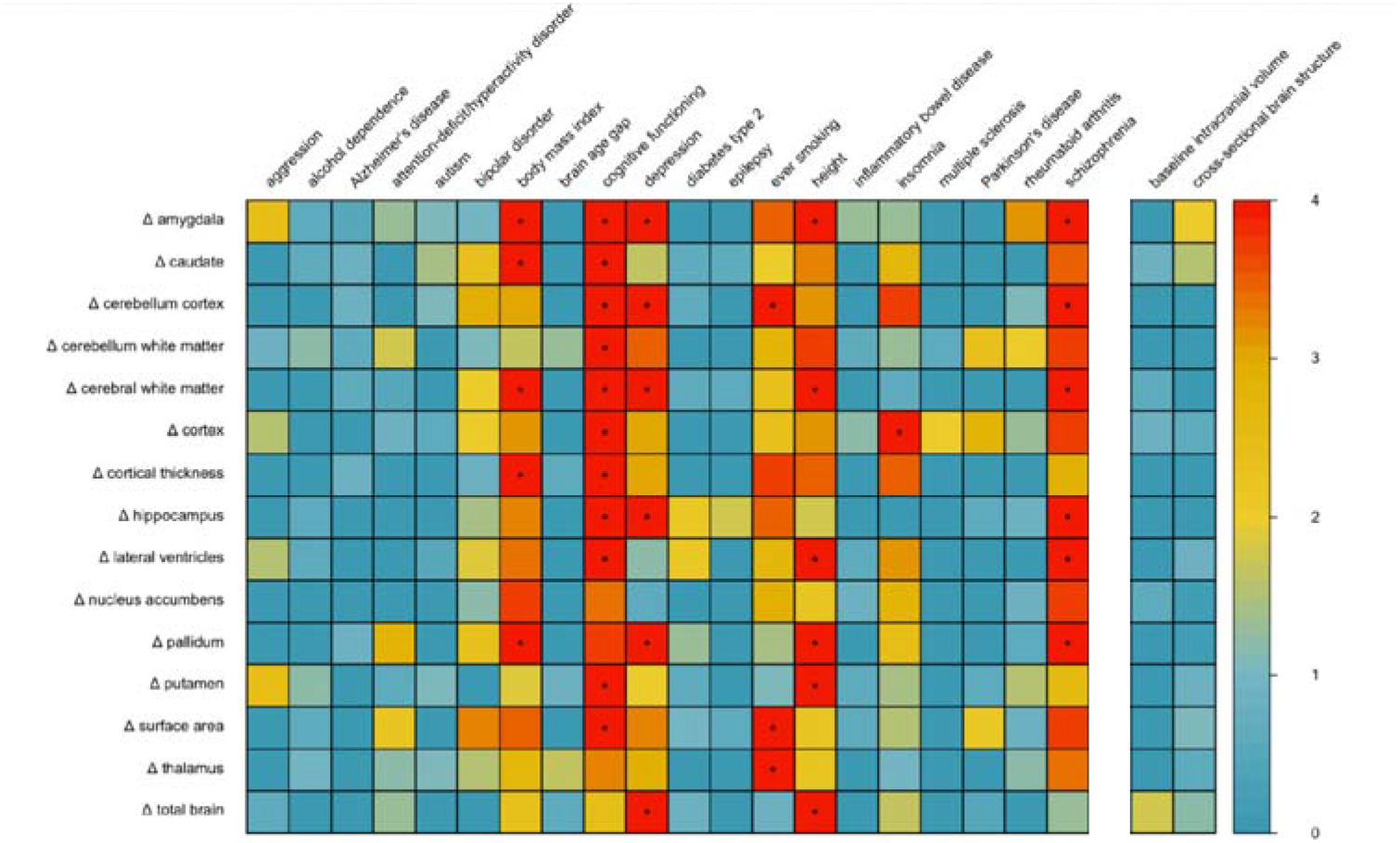
Genetic overlap with other phenotypes. *P*-values for pleiotropy between change rates of structural brain measures (rows, indicated by Δ for change rate) and neuropsychiatric, disease-related and psychological traits (columns left of colour legend). *P*-values for pleiotropy between change rates of structural brain measures and head size (intracranial volume) and the cross-sectional brain measure are displayed on the right (columns right of colour legend). Significant overlap (*p* < 1.6e-04) is marked with *.

## Overlap with other traits

We applied iSECA for overlap between our age-independent summary statistics for structural brain changes and several neuropsychiatric, neurological, physical, ageing and disease-related phenotypes and psychological traits. We found significant genomic overlap (*p* < 1.6e-04) with genetic variants associated with depression^27^, schizophrenia^28^, cognitive functioning^29^, height^30^, insomnia^31^, body mass index (BMI)^30^ and ever smoking^32^. Despite significant pleiotropy between rates of change and these traits, we did not find evidence for concordance or discordance of effects (Figure 4, External Data Fig. 7). For comparison, we computed the genomic overlap between cross-sectional volumes and these phenotypes using the same method. In general, cross-sectional volumes showed overlap for the same traits and several others. Of note, there was also little overlap between the summary statistics for the longitudinal brain measures and summary statistics for the corresponding volumes, based on cross-sectional data. This implies that despite the fact that both cross-sectional brain volume and rates of changes are associated with traits such as schizophrenia or cognitive functioning, these associations are likely not driven by the same genomic locations. Additionally, there was little overlap in the genetic loci associated with the longitudinal brain measures and intracranial volume at baseline, indicating that overall head size did not drive our findings (Figure 4).

## Gene expression in the brain across the lifespan

We determined mRNA expression for genome-wide significant genes and genes associated with genome-wide significant SNPs (Supplementary Tables S5,7) in 54 tissue types and in both the developing and adult human brain (Methods). For the prioritized genes, a gene expression heatmap was created, based on GTEx v8 RNAseq data^33^. This revealed considerable expression levels across several brain tissues for the following genes: *APOE, CAB39L, FAU, NECTIN2* (alias *PVRL2*) and *SORCS2*, the latter showing higher relative expression in brain tissue compared to all other tissue types (Extended Data Fig. 8A). Expression heatmaps based on BrainSpan data^34^ revealed that *DACH1* shows highest relative expression during early prenatal stages (8-9 post conception weeks), compared to postnatal stages. A second cluster of genes demonstrated stable high relative expression levels throughout development and across the lifespan (*APOE, CAB39L, FAU, NECTIN2* (alias *PVRL2*)). One additional gene, *CDH8*, showed lower relative expression in the early prenatal stages and higher expression later in life (Extended Data Fig. 8B).

## Phenome-wide associations

For the prioritized SNPs and genes (Supplementary Tables 5,7), exploratory pheWAS (i.e., “phenome-wide”) analysis was performed to systematically analyse many phenotypes for association with the genotype and individual genes (Supplementary Table 16). PheWAS was performed using publicly available data from the GWASAtlas^32^ (https://atlas.ctglab.nl). Gene associations of *DACH1, GPR139* and *SORCS2* showed pleiotropic effects mainly in the metabolic domain, e.g., with estimated glomerular filtration rate and BMI (Supplementary Table 16, Extended Data Fig. 9). *SORCS2* and *CDH8* also showed significant associations with psychiatric and cognitive traits. Both *APOE* and *NECTIN2* showed strongest associations with Alzheimer’s disease, cholesterol and lipids (Supplementary Table 16, Extended Data Fig. 9).

## Sensitivity analyses

We repeated the SNP and gene analyses in various subgroups: 1) by adding four cohorts of non-European or mixed ancestry (N=540; total N=15,640); 2) by omitting cohorts that did not meet a minimum sample size criterion (N>75) or a minimum scanning interval (> 0.5 years) leaving N=14,601; 3) by excluding diagnostic groups in each cohort leaving N=13,034, and 4) by including a covariate adjusting for disease status (Supplementary Tables 17,18). In SNP-based and gene-based analyses, effect sizes of SNPs were very similar in all subgroups, suggesting that our results are also applicable for individuals of non-European ancestry - with the caveat that the non-European subgroup was rather small - and not driven by the smaller cohorts. Findings were also similar in the healthy subgroup and when correcting for disease status, with one notable exception: the *APOE* finding for hippocampus and amygdala rate of change showing increasing influence of the top SNP with age, was no longer present when correcting for disease (see Supplementary Tables 1 for diagnoses). This suggests that these *APOE* findings for hippocampus and amygdala were in part driven by the presence of patients, which could be either explained through disease related genes that also influence rates of change, or brain changes as a consequence of the disease.

Given that our main analyses included patients and iSECA analyses showed several associations with disease, we repeated iSECA analyses excluding diagnostic groups in each cohort. These analyses implicate the same traits, associated with largely the same rates of change of brain measures. (Extended Data Fig. 7).

## Discussion

Here, we present the first GWAS investigating influences of common genetic variants on brain-structural changes in over 15,000 subjects covering the lifespan. The longitudinal design of our study combined with the large age range assessed provides a flexible framework to detect age-independent and age-dependent effects of genetic variants on rates of structural brain changes. We identified genetic variants for structural brain changes between 4 and 99 years of age. Some of these were independent of age, showing effects of stable strength and direction throughout life, suggesting that these genetic variants are equally crucial for early brain development as for brain ageing. In addition, we identified age-dependent genetic variants, suggesting that some genetic variants are predominantly associated with brain development while others are mainly associated with brain ageing.

Amongst our top findings is the *APOE* gene, a major risk factor for AD^23^, and specifically a missense variant in that gene, which influences amygdala and hippocampus rates of change with varying and differential effects across the lifespan, with probably most pronounced effects in those affected with brain disorders. However, while most of the additional genetic loci identified here have not previously been associated with any brain-plasticity-related phenotypes, several others were also linked to brain disorders, including psychiatric (e.g., *GPR139* and *CDH8*) and neurodegenerative disorders (e.g., *NECTIN2*). Notably, *DACH1* and *NECTIN2* show increased expression during early development, while other genes’ brain expression patterns are most pronounced during adulthood (e.g., *APOE* and *CDH8*), suggesting that these genes may exert specific effects during different developmental periods.

Gene-set analysis also implies a role for both developmental and neurodegenerative processes. We found a gene-set involved in neural nucleus development to influence rates of change in cerebellar white matter, and other closely related gene ontology terms: neural nucleus development in the substantia nigra and midbrain, associated with rates of change of cerebral white matter and surface area. These implicate the biological process of progression of a neural nucleus from its initial condition or formation to its mature state. This would also suggest that genes involved in early developmental mechanisms of subcortical nuclei are related with cortical changes later in life. In addition, we found several gene-sets interacting with tau-protein associated with rate of change in caudate volume, and a gene-set associated with rate of change in surface area, implicating phorbol 13-acetate 12-myristate, an activator of protein kinase C (PKC)^25^. PKC is a family of enzymes whose members transduce a large variety of cellular signals and plays a key role in controlling the balance between cell survival and cell death. Its loss of function is generally associated with cancer, whereas its enhanced activity is associated with neurodegeneration. PKC both directly phosphorylates tau and indirectly causes the dephosphorylation of tau, and has been suggested to play a key role in the pathology of Alzheimer’s disease^35^. Together these results suggest involvement of genes in ageing and neurodegeneration.

At the global, genome-wide level, we found significant genomic overlap with genetic variants associated with depression, schizophrenia, cognitive functioning, insomnia, height, body mass index (BMI) and ever-smoking. Several of these traits, such as schizophrenia, smoking, cognitive functioning, and body mass index, have been associated with longitudinal brain-structural changes^5,36–38^. The global overlap coincides with findings at the individual gene level: several of the identified genetic variants and genes were linked to metabolic processes (*APOE, DACH1, GPR139, NECTIN2*), cognitive functioning (*CDH8*), psychiatric traits (*GPR139, SORCS2, CDH8*) and Alzheimer’s disease (*NECTIN2* and *APOE*) as apparent from the pheWAS results. Despite the pleiotropic effects, concordance of effects was generally null. This is not surprising, as rates of change measures for brain structures are not constant and often switch sign over the course of the lifespan^1,39^, while the GWAS for other traits assume stability of both the phenotype and the genetic influences on the phenotype over time. As such, concordance and discordance of effects are not to be expected.

The advantage of longitudinal analyses is that each individual acts as their own control, allowing us to separate the genetic effects on cross-sectional volumes from those on the rates of change^14^. Indeed, we found little overlap between the two: top genes identified in the GWAS on cross-sectional brain structure^26,40–42^ generally did not overlap with the top genes for the corresponding rates of change. Longitudinal analyses have for long been shown to provide different information than cross-sectional approaches. On a phenotypic level, comparisons between cross-sectional and longitudinal ageing patterns of the hippocampus show different results^43^. On a genetic level, a study including a within-sample SNP times age interaction in the ADNI cohort, which is included in this study, showed larger power to detect genetic associations in a longitudinal design compared to a cross-sectional analysis^44^. Of note, that study also identified rs429358 in *APOE* associated with longitudinal hippocampal and amygdala volume change in older age. We now show this variant to exert a lifespan effect through our meta-regression approach, with the risk variant for AD causing faster increases in childhood for amygdala and faster decay for both amygdala and hippocampus later in life.

Given the dynamics of brain structural changes during the lifespan, we investigated both age-independent and age-dependent genetic effects. The age-independent effects can be interpreted as neurodevelopmental influences that also impact brain structure at older ages^45,46^, whereas the age-dependent effects can be interpreted as possible changing effects of genes or gene expression during life^16^. The genome-wide meta-regression approach employed here may enable future GWAS for other phenotypes that change over the human lifespan.

We chose to analyse longitudinal changes for 15 separate brain structures, because we observed generally low correlations between these phenotypic changes. This approach allowed us to find brain structure specific associations. However, there are several longitudinal studies describing phenotypic correlations between structural changes ^39,47,48^, and combining several phenotypes could be an alternative approach to identify genetic variants exerting a global effect. Of note, cohort and age are intertwined in meta-regression analysis and in that sense, we cannot be sure that differences between cohorts are exclusively attributed to age. Mega-analysis would circumvent this problem, but was not feasible in practice. Moreover, we imposed the same stringent criteria of genome-wide significance for the age-independent meta-analysis and age-dependent meta-regression, which renders chance findings equally unlikely in either type of analysis. In addition, residual heterogeneity for the top findings was generally small. That said, our sample size is still relatively modest for GWAS purposes, and replication in larger samples and inclusion of other ancestries is needed, once more longitudinal data becomes available.

How exactly variation in these genes impacts brain changes in health and disease cannot be answered based on genome-wide association studies. To this end, our findings may direct future studies into brain development and ageing, and prevention and treatment of brain disorders. For example, biological pathways that guide neural nucleus development in the foetal subcortical brain may be particularly relevant to the cerebral white matter growth and cortical thinning that takes place during childhood and adolescence. Neurodegenerative disorders might be better understood when we identify genetic variants that influence brain atrophy over time, compared with identification of static genetic differences. In conclusion, our study shows that our genetic architecture is associated with the dynamics of human brain structure throughout life.

## Supporting information

Supplementary Note & Suppelementary Figure legends

## Methods

### Ethical approval

All participants gave written informed consent and all participating sites obtained approval from local research ethics committees/institutional review boards. Ethics approval for meta-analyses within the ENIGMA consortium was granted by the QIMR Berghofer Medical Research Institute Human Research Ethics Committee in Australia (approval: *P2204*).

### Inclusion criteria

Cohorts that had longitudinal magnetic resonance imaging (MRI) data of the brain and genotyped data extracted from blood or saliva available were invited to participate, irrespective of disease status and age. Patients were not excluded as aberrant brain trajectories are often observed and we hypothesize that genetic risk for disease may be associated with genetic influences on rates of change. We included cohorts that had a preferred sample size of at least 75 subjects and a follow up duration (for repeated MRI scans) of at least six months. After quality control of individual subject’s imaging and genotyping data, not all the cohorts could meet these criteria. In total, we included 15,640 subjects aged 4 to 99 (49% female, 14% patients). Please see Extended Data Fig. 1 and Supplementary Table 1 for further description of the cohorts.

### Longitudinal imaging

Eight global brain measures (total brain including cerebellum and excluding brainstem, surface area measured at the grey-white matter boundary, average cortical thickness, total lateral ventricle volume, and cortical and cerebellar grey and white matter volume) and seven subcortical structures (thalamus, caudate, putamen, pallidum, hippocampus, amygdala and nucleus accumbens) were extracted from the FreeSurfer processing pipeline^49–51^; see Supplementary Table 2 for details per cohort). We chose these measures based on the fact that they show generally high test-retest reliability for cross-sectional measures^52–54^, thereby selecting those measures that would have sufficient signal to noise in change measures. Image processing and quality control were performed at the level of the cohorts, following harmonized protocols (http://enigma.ini.usc.edu/protocols/imaging-protocols/) which included visual inspection of the segmentation. Annual rates of change were computed in each individual for each phenotype by subtracting baseline brain measures from follow up measures and dividing by the number of years of follow-up duration. We chose not to correct for overall head size in the main analysis: while it is common practice to correct for intracranial volume when investigating cross-sectional brain volumes^55^, the associations between intracranial volume and brain changes over time are small (Extended Data Fig. 2) and GWAS findings are very similar with and without correction (Supplementary Note). Distributions of baseline and follow-up measures - as well as annual rates of changes - were visually inspected and change rates were centrally compared for consistency.

Longitudinal trajectories of brain structure rates of change were estimated by applying locally, cohort-size weighted, estimated scatterplot smoothing with a Gaussian kernel, local polynomials of degree 2 and a span of 1 (LOWESS^56^) implemented in R^57^. Integrating these trajectories and then fitting these to the baseline values of the phenotypes in the cohorts provides trajectories throughout the lifespan. Trajectories were estimated in the full dataset including patients and by excluding diagnostic groups in each cohort separately.

### Genome-wide association analysis

At each participating site, genotypes were imputed using the 1000 Genomes project dataset^58^ through the Michigan imputation server^59^ (https://imputationserver.sph.umich.edu/) or the Sanger imputation server^60^ (Supplementary Table 3). Subsequently, each site ran the same multidimensional scaling (MDS) analysis protocol, computing MDS components from the combination of their cohort’s data with the HapMap3 population^61^. This ensured that all sites corrected for ancestry in a consistent manner. See http://enigma.ini.usc.edu/protocols/genetics-protocols/ for the imputation and MDS analysis protocol. Within each cohort genome-wide association was conducted using an additive model, modelling change rate as a function of the genetic variant plus covariates age, sex, age*sex, age^2^, age^2^*sex and ancestry (the first four MDS components). While it is possible that rates of brain structural changes are different in males and females, we did not have the power to perform analyses separating the sexes. Dummy variables were added where appropriate, e.g., when multiple scanners were used. We re-ran these analyses adding a covariate for disease status if the cohorts contained patients and controls. Most sites used our harmonized GWAS protocol, which used *raremetalworker*^62^ for analysis (Supplementary Table 3). Regardless of the study design, a kinship matrix was incorporated in these analyses, accounting for relatedness in family studies, or possible unknown kinship in the other studies. Given the small sample sizes of the individual cohorts, a stringent cohort level quality control was enforced, to exclude variants with a minor allele frequency (MAF) < 0.05 or variants with imputation R^2^ / info score < 0.75. Across cohorts and phenotypes, GWAS summary plots (Manhattan plots and QQ plots) were visually inspected at the central site. If a given cohort / trait showed deviation from expectations, sites were asked to re-analyse their data, which usually involved removal of outliers in the phenotypic data. QQ plots per cohort, per phenotype can be found in Extended Data Figure 12.

### Meta-analysis and Meta-regression

In the phase 1 cohorts of European ancestry (N=9,604) we aggregated the cohort-level data for each phenotype, using standard-error weighted meta-analysis or meta-regression. We employed a cumulative meta-analysis and meta-regression approach for replication, in phase 2 (N=15,100). We tested three models. Under the assumption that effect sizes of single nucleotide polymorphisms (SNPs) were consistent across the lifespan (i.e., a standard meta-analytic approach), where the subscript C denotes a cohort and ∑ an error term:

1) Effect_SNP_C_ ~ b_0_ + ∑_C_, under the null hypothesis that b_0_ = 0.

Given that brain changes throughout life are dependent on age, the effects of a genetic variant on brain change is likely to depend on age too. Within cohorts, such an age by SNP effect analysis would not have been feasible since longitudinal cohorts that span the age-range between 4-99 years do not exist. Given the widespread mean age among the cohorts included (Extended Data Fig. 1 and Supplementary Table 1), it was possible to calculate the age-dependent effects across the life span by comparing effects of loci between cohorts, through meta-regression. Meta-regression is a sophisticated tool for addressing heterogeneity between cohorts in meta-analyses when the source of heterogeneity is known (in this case, age)^63^. We estimated the following model under the assumption that the effects of SNPs may vary in size or direction across the lifespan:

2) Effect_SNP_C_ ~ b_0_ + b_1_*age_C_ + ∑_C_ under the null hypothesis that b_1_=0 (1 degree of freedom), and
3) Effect_SNP_C_ ~ b_0_ + b_1_*age_C_ + b_2_*age_C_^2^ + ∑_C_ under the null hypothesis that (b_1_=b_2_=0, 2 degrees of freedom).

SNP data were aligned using METAL^64^ for all three analyses. The age-independent effect of SNPs (model 1) was computed in METAL. For the age-dependent analyses (model 2 for linear age effects and model 3 for quadratic age effects) the aligned data were imported into R^52^ and fixed effects meta-regression was performed using the R-package metafor^65^ (version 2.0-0). Results were filtered on SNPs that were present for at least 50% of the cohorts and in at least 50% of the subjects.

### Functional mapping

Functional mapping was performed using the FUMA platform designed for prioritization, annotation and interpretation of GWAS results^66^. As the first step, independent significant SNPs in the individual GWAS meta-analysis summary statistics were identified based on their *p*-value (p < 5 × 10^−8^) and independence of each other (r2 < 0.6 in the 1000G phase 3 reference) within a 1Mb window. Thereafter, lead SNPs were identified from independent significant SNPs, which are independent of each other (r2 < 0.1). We used FUMA to annotate lead SNPs in genomic risk loci based on the following functional consequences on genes: eQTL data (GTEx v6 and v7^67^), blood eQTL browser^68^, BIOS QTL browser^69^, BRAINEAC^70^, MuTHER^71^, xQTLServer^72^, the CommonMind Consortium^73^ and 3D chromatin interactions from HI-C experiments of 21 tissues/cell types^74^. Next for eQTL mapping and chromatin interaction mapping, genes were mapped using positional mapping, which is based on a maximum distance between SNPs (default 10kb) and genes. Chromatin interaction mapping was performed with significant chromatin interactions (defined as FDR < 1 × 10^−6^). The two ends of significant chromatin interactions were defined as follows: region 1 – a region overlapping with one of the candidate SNPs, and region 2 – another end of the significant interaction, used to map to genes based on overlap with a promoter region (250bp upstream and 50bp downstream of the transcription start site).

### Visualization of SNP effects

We visualized the effects of our top SNPs on the lifespan trajectory, assuming no effects of the other SNPs, for easier interpretation of the direction of effect. Similar to the estimation of the lifespan trajectory, we estimated a smoothed version *f(x)* of the phenotypic change rate using LOWESS (see above) and integrated the rate of change. We added the unknown volume *C* at the start of our age range by fitting the integrated curve to the baseline data. Suppose *h(x)* is the unknown rate of change for non-carriers. The additional change rate *g(x)* for carriers was estimated through the meta-analysis or meta-regression. The full dataset contained a fraction *p* of the carriers of the tested allele. Assuming *p + q = 1, f(x) = p*(h(x) + g(x)) + q*h(x) = h(x) + p*g(x)*. We created a rate of change curve for non-carriers as *f(x)-p*g(x)* and a rate of change curve of carriers as *f(x)+q*g(x)*. The offset *C* is potentially different in carriers and non-carriers, so we estimated this difference by taking the effect of the cross-sectional GWAS data (see below) in this SNP, or a proxy SNP in high linkage disequilibrium (LD).

### Gene-based and gene-set analyses

Gene-based associations with 15 phenotypes were estimated using MAGMA^75^ (version 1.09a) using the summary statistics from age-independent and age-dependent GWAS metaanalyses of rate of change of global brain measures. Gene names and locations were based on NCBI 37.3 locations as provided by MAGMA. Association was tested using the SNP-wise mean model, in which the sum of −log(SNP *p*-value) for SNPs located within the transcribed region was used as the test statistic. LD correction was based on estimates from the 1000 Genomes Project Phase 3 European ancestry samples^58^. To describe the direction of the age effect for significant genes in the age-dependent analyses, we subsequently identified the SNPs that were used in the gene-based *p*-value and plotted the age-dependent effect of the top SNP that contributed to the gene-based *p*-value.

The generated gene-based *p*-values were used to analyse sets of genes in order to test for association of genes belonging to specific biological pathways or processes. MAGMA applies a competitive test to analyse if the genes of a gene set are more strongly associated with the trait than other genes, while correcting for a series of confounding effects such as gene length and size of the gene set. For gene sets we used 9,975 sets with 10 −1,000 genes from the Gene Ontology sets^76^ curated from MsigDB 7.0^77^.

### Multiple testing corrections

We investigated annual rates of change for 15 brain phenotypes, but these are correlated to some extent (Extended Data Fig. 2). We therefore estimated the effective number of independent variables based on matrix spectral decomposition^78^ for the largest adolescent cohort (IMAGEN; N=1,068) and for the largest elderly cohort from the phase 1 sample (ADNI2; N=626). The most conservative estimate of the number of independent traits was 13.93. Despite the fact that models 2 and 3 are nested and therefore not independent, we also corrected for performing three analyses per trait. The study-wide significant threshold for the genome was therefore set at *p* < 1.2e-09 (5e-08/13.93*3). For gene-based significance, we applied a genome-wide significance level of 0.05/17541= 2.85e-06, and a study wide significance of 2.85e-06/(13.93*3), i.e. *p* < 6.82e-08. For gene-set significance, we applied a genome-wide significance level of 0.05/9,975 = 5.01e-06 and a study-wide significance level of 5.01e-06/(13.93*3), i.e. *p* < 1.20e-07.

### SNP heritability

SNP heritabilities, *h^2^_SNP_*, were estimated by using linkage disequilibrium (LD) score regression^79^ (LDSR) for the European-ancestry brain change GWASs to ensure matching of population LD structure. For LDSR, we used precomputed LD scores based on the European-ancestry samples of the 1000 Genomes Project^58^ restricted to HapMap3 SNPs^61^. The summary statistics with standard LDSC filtering were regressed onto these scores. SNP heritabilities were estimated based on the slope of the LD score regression, with heritabilities on the observed scale calculated. To ensure sufficient power for the genetic correlations, r_g_ was calculated if the Z-score of the *h^2^_SNP_* for the corresponding GWAS was 4 or higher^79^.

### Comparison with cross-sectional results

For the genome-wide significant genes and genes associated with genome-wide significant SNPs, we compared our findings with cross-sectional GWAS summary statistics when available. To this end, datasets^26,40–42^ were requested and downloaded from http://enigma.ini.usc.edu/research/download-enigma-gwas-results/ and http://big.stats.ox.ac.uk/download_page. Gene-based association analyses for cross-sectional brain GWAS summary statistics were performed using MAGMA (as described above). Additionally, we compared the overlap in the first 1,000 ranked genes to the expected number of overlapping genes based on chance. False discovery rate correction^80^ was applied to determine over- or under-representation of genes from our longitudinal GWAS to the cross-sectional previously published GWAS^26,40–42^.

### Genetic overlap with cross-sectional results and other traits

To investigate genetic overlap with other traits across the genome we applied an adapted version of iSECA^81^ (independent SNP effect concordance analysis) which examines pleiotropy and concordance of the direction of effects between two phenotypes by comparing expected and observed overlap in sets of SNPs from both phenotypes that are thresholded at different levels. From the results at each threshold, heatmap plots were generated containing binomial tests for pleiotropy and Fisher’s exact tests for concordance. An empirical *p*-value for overall pleiotropy and concordance was then generated through permutation testing. Our implementation of iSECA also included a *p*-value for overall discordance, as we expect some phenotypes to negatively influence brain-structural change rates. *P*-values were computed using a two-step approach: we first ran 1,000 permutations. If the *p*-value for pleiotropy was below 0.05/15 we reran the analyses with 10,000 permutations to obtain a more precise *p*-value. Summary statistics of change rates were first filtered on SNPs for which > 95% of the subjects contributed data to remove the sample size dependency of *p*-values and subsequently clumped (*p*=1, kb=1000) to ensure independence of input SNPs.

We investigated the genetic overlap between brain-structural changes and risk for 20 neuropsychiatric, neurological and somatic disorders, and physical and psychological traits. Summary statistics were downloaded or requested for aggression^82^, alcohol dependence^83^, Alzheimer’s disease^84^, attention-deficit/hyperactivity disorder^85^, autism^86^, bipolar disorder^87^, body mass index^30^, brain age gap^12^, cognitive functioning^29^, depression^27^, diabetes type 2^88^, ever smoking^32^, focal epilepsy^89^, height^30^, inflammatory bowel disease^90^, insomnia^31^, multiple sclerosis^91^, Parkinson’s disease^92^, rheumatoid arthritis^93^ and schizophrenia^28^. These phenotypes were chosen because of known associations with brain structure or function, and availability of summary statistics based on large GWA-studies. For comparison, we computed the genetic overlap between cross-sectional brain structure and these phenotypes, using the same method.

Apart from these, we also 1) included intracranial volume^94^ to investigate the effect of overall head size and 2) tested the overlap between each structure’s longitudinal change measure against its cross-sectional brain structure. Pleiotropy, concordance or discordance was considered significant when the *p*-value was smaller than 0.05/15*22 (#change rates * #phenotypes tested) = 1.6e-04.

### Brain gene expression

GENE2FUNC, a core process of FUMA^66^ (Functional Mapping and Annotation of Genomewide Association Studies), was employed to analyse gene expression patterns. For this, a set of 8 genes was used as input, including all genome-wide significant genes and genes harbouring genome-wide significant SNPs (compare Supplementary Tables 4-7). Gene expression heatmap was constructed employing GTEx v8^33^; 54 tissue types) and BrainSpan RNA-seq data across 29 different ages or 11 different developmental stages^32^. The average of normalized expression per label (zero means across samples) was displayed on the corresponding heatmaps. Expression values are TPM (Transcripts Per Million) for GTEx v8 and RPKM (Read per Kilobase Million) in the case of the BrainSpan data set.

### Phenome-wide association studies

To identify phenotypes associated with the candidate SNPs and genes (defined as genome-wide significant SNPs and the genome-wide significant genes and genes associated with genome-wide significant SNPs), a phenome-wide association study (pheWAS) was done for each SNP and/or gene. PheWAS was performed using public data provided by GWASAtlas^32^(https://atlas.ctglab.nl). To correct for multiple testing, the total number of GWASs (4,756) was considered (including GWASs in which the searched SNP or gene was not tested) and the number of tested SNPs and genes (n=14), resulting in a Bonferroni corrected *p*-value threshold of 1.05e-05/14, i.e., *p* < 7.51e-07.

### Sensitivity analyses

The phase 2 analyses include available data from all cohorts with European ancestry (N=15,100). The four cohorts of non-European and mixed ancestry together consist of 540 subjects, who are predominantly children and adolescents (Supplementary Table 3). The number of subjects, heterogeneity in ancestry and the age-distribution do not allow for separate meta-analysis or meta-regression. We therefore added the cohorts of non-European ancestry to the original datasets and reran analyses (N=15,640). In a second analysis, we excluded the 9 cohorts that had N < 75 or mean scanning interval < 0.5 years (Supplementary Table 2), leaving N=14,601 subjects. The main analyses include data from all subjects combined, without correction for disease. This approach was chosen because many neurological and neuropsychiatric diseases are characterized by aberrant brain changes over time, and genes involved in the disease may also be involved in these brain changes. To check whether our results were confounded by disease, we repeated the main analyses excluding diagnostic groups of each cohort (N=13,0349) and by correcting for disease status.

## Data and code availability

This work is a meta-analysis. Upon publication, the meta-analytic results will be made available from the ENIGMA consortium webpage (http://enigma.ini.usc.edu/research/download-enigma-gwas-results). Cohort level data can be shared upon request, after permission of cohort principle investigators. Individual level data can be shared with interested investigators, subject to local and national ethics regulations and legal requirements that respect the informed consent forms and national laws of the country of origin of the persons scanned. Code for the meta-regression is available through Github https://github.com/RMBrouwer/GWAS_meta_regression.

## Funding

The ENIGMA-Plasticity working group is part of the ENIGMA World Aging Center, funded by NIA grant 1R56AG058854-01. The ENIGMA Consortium core funding is supported by NIH Consortium grant U54 EB020403, supported by a cross-NIH alliance that funds Big Data to Knowledge Centers of Excellence.

### 1000BRAINS

1000BRAINS is a population-based cohort based on the Heinz-Nixdorf Recall Study and is supported in part by the German National Cohort. We thank the Heinz Nixdorf Foundation (Germany) for their generous support in terms of the Heinz Nixdorf Study. The authors are supported by the Initiative and Networking Fund of the Helmholtz Association (Svenja Caspers) and the European Union’s Horizon 2020 Research and Innovation Programme under Grant Agreements 785907 (Human Brain Project SGA2; Svenja Caspers, Sven Cichon, and Katrin Amunts). This work was further supported by the German Federal Ministry of Education and Research (BMBF) through the Integrated Network IntegraMent (Integrated Understanding of Causes and Mechanisms in Mental Disorders) under the auspices of the e:Med Program (grant 01ZX1314A; Sven Cichon), and by the Swiss National Science Foundation (SNSF, grant 156791; Sven Cichon).

### ABCD

Data used in the preparation of this article were obtained from the Adolescent Brain Cognitive DevelopmentSM (ABCD) Study (https://abcdstudy.org), held in the NIMH Data Archive (NDA). This is a multisite, longitudinal study designed to recruit more than 10,000 children age 9-10 and follow them over 10 years into early adulthood. The ABCD Study® is supported by the National Institutes of Health and additional federal partners under award numbers U01DA041048, U01DA050989, U01DA051016, U01DA041022, U01DA051018, U01DA051037, U01DA050987, U01DA041174, U01DA041106, U01DA041117, U01DA041028, U01DA041134, U01DA050988, U01DA051039, U01DA041156, U01DA041025, U01DA041120, U01DA051038, U01DA041148, U01DA041093, U01DA041089, U24DA041123, U24DA041147. A full list of supporters is available at https://abcdstudy.org/federal-partners.html. A listing of participating sites and a complete listing of the study investigators can be found at https://abcdstudy.org/consortium_members/. ABCD consortium investigators designed and implemented the study and/or provided data but did not necessarily participate in the analysis or writing of this report. This manuscript reflects the views of the authors and may not reflect the opinions or views of the NIH or ABCD consortium investigators. The ABCD data repository grows and changes over time. The ABCD data used in this report came from Data Release 3.0 (http://dx.doi.org/10.15154/1519007).

### ADNI

Data collection and sharing for this project was funded by the Alzheimer’s Disease Neuroimaging Initiative (ADNI) (National Institutes of Health Grant U01 AG024904) and DOD ADNI (Department of Defense award number W81XWH-12-2-0012). ADNI is funded by the National Institute on Aging, the National Institute of Biomedical Imaging and Bioengineering, and through generous contributions from the following: AbbVie, Alzheimer’s Association; Alzheimer’s Drug Discovery Foundation; Araclon Biotech; BioClinica, Inc.; Biogen; Bristol-Myers Squibb Company; CereSpir, Inc.; Cogstate; Eisai Inc.; Elan Pharmaceuticals, Inc.; Eli Lilly and Company; EuroImmun; F. Hoffmann-La Roche Ltd and its affiliated company Genentech, Inc.; Fujirebio; GE Healthcare; IXICO Ltd.;Janssen Alzheimer Immunotherapy Research & Development, LLC.; Johnson & Johnson Pharmaceutical Research & Development LLC.; Lumosity; Lundbeck; Merck & Co., Inc.; Meso Scale Diagnostics, LLC.; NeuroRx Research; Neurotrack Technologies; Novartis Pharmaceuticals Corporation; Pfizer Inc.; Piramal Imaging; Servier; Takeda Pharmaceutical Company; and Transition Therapeutics. The Canadian Institutes of Health Research is providing funds to support ADNI clinical sites in Canada. Private sector contributions are facilitated by the Foundation for the National Institutes of Health (www.fnih.org). The grantee organization is the Northern California Institute for Research and Education, and the study is coordinated by the Alzheimer’s Therapeutic Research Institute at the University of Southern California. ADNI data are disseminated by the Laboratory for Neuro Imaging at the University of Southern California.

### ALS Utrecht

The authors acknowledge grants supporting their work from the European Union’s Horizon 2020 Research and Innovation Programme (H2020/2014–2020) under grant agreements 667302 (CoCA), 728018 (Eat2beNICE), 785907 (HBP SGA2), and 772376 (EScORIAL) and the Netherlands ALS Foundation.

### BDC

Brain Dynamics Centre (BDC), Sydney - cohort is funded by a National Health & Medical Research Council of Australia Project Grant (APP1008080).

### BHRCS

The Brazilian High Risk Cohort Study (BHRCS) was supported by the National Institute of Developmental Psychiatry for Children and Adolescent (INPD) Grant: Fapesp 2014/50917-0 CNPq 465550/2014-2.

### BIG

This study used the BIG database, which was established in Nijmegen in 2007. This resource is now part of Cognomics, a joint initiative by researchers of the Donders Centre for Cognitive Neuroimaging, the Human Genetics and Cognitive Neuroscience departments of the Radboud university medical center, and the Max Planck Institute for Psycholinguistics. The Cognomics Initiative is supported by the participating departments and centres and by external grants, including grants from the Biobanking and Biomolecular Resources Research Infrastructure (Netherlands) (BBMRI-NL) and the Hersenstichting Nederland. In particular, the authors would also like to acknowledge grants supporting their work from the Netherlands Organization for Scientific Research (NWO), i.e. the NWO Brain & Cognition Excellence Program (grant 433-09-229) and the Vici Innovation Program (grant 016–130–669 to BF). Additional support is received from the European Community’s Seventh Framework Programme (FP7/2007 – 2013) under grant agreements n° 602805 (Aggressotype), n° 603016 (MATRICS), n° 602450 (IMAGEMEND), and n° 278948 (TACTICS), and from the European Community’s Horizon 2020 Programme (H2020/2014 – 2020) under grant agreements n° 643051 (MiND) and n° 667302 (CoCA).

### BrainSCALE

The BrainSCALE study is a collaborative project between Netherlands Twin Register (NTR) at the Vrije Universiteit (VU) Amsterdam and University Medical Center Utrecht (UMCU). The BrainSCALE study was funded by Nederlandse Organisatie voor Wetenschappelijk Onderzoek (NWO 51.02.061 to H.H., NWO 51.02.062 to D.B., NWO-NIHC Programs of excellence 433-09-220 to H.H., NWO-MagW 480-04-004 to D.B., and NWO/SPI 56-464-14192 to D.B.); FP7 Ideas: European Research Council (ERC-230374 to D.B.), Universiteit Utrecht (High Potential Grant to H.H.), Netherlands Twin Registry Repository (NWO-Groot 480-15-001/674 to DB) and Neuroscience Campus Amsterdam (NCA). Biomolecular Resources Research Infrastructure (BBMRI–NL, 184.021.007 and 184.033.111) Developmental trajectories of psychopathology (NIMH 1RC2 MH089995); and the Avera Institute for Human Genetics, Sioux Falls, South Dakota (USA).

### Capetown

The CTAAC study was supported by grant No R01-HD074051.

### DBSOS

The DBSOS study is partially funded by the Brain and behavior Foundation (NARSAD) by an Independent Investigator grant; No 20244. The generation R Study is made possible by financial support from the Erasmus Medical center, Rotterdam and the Netherlands organization for health research and development (ZonMW). The neuroimaging infrastructure is supported by ZonMW TOP (No: 912110210), The NWO Physical Sciences Division, and SURFsara supercomputing center (Cartesius Compute Cluster).

### FOR2107

This work is part of the German multicenter consortium “Neurobiology of Affective Disorders. A translational perspective on brain structure and function”, funded by the German Research Foundation (Deutsche Forschungsgemeinschaft DFG; Forschungsgruppe/Research Unit FOR2107). Grant agreements included the following: FOR2107 DA1151/5-1 and DA1151/5-2 to UD; SFB-TRR58, Projects C09 and Z02 to UD; the Interdisciplinary Center for Clinical Research (IZKF) of the medical faculty of Münster (grant Dan3/012/17 to UD); KR 3822/7-1 and KR 3822/7-2 to AK; KI 588/14-1, KI 588/14-2; NO 246/10-1 and NO 246/10-2 to MMN. AJ was in particular involved as PI in WP6, multi-method data analytics (JA 1890/7-1, JA 1890/7-2). FOR2107 study was also supported by the German Federal Ministry of Education and Research (BMBF), through ERA-NET NEURON, “SynSchiz - Linking synaptic dysfunction to disease mechanisms in schizophrenia - a multilevel investigation” (01EW1810 to MR) and the German Research Foundation (DFG grant FOR2107; RI908/11-2 to MR).

### Generation R

Netherlands Organization for Health Research and Development (ZonMw) TOP project number 91211021. Sophia Children’s Hospital Foundation (Stichting Vrienden van het Sophia) project number S18-68. The Generation R sample further reports the following support: Super computing resources for imaging processing were supported by the NWO Physical Sciences Division (Exacte Wetenschappen) and SURFsara (Cartesius compute cluster, https://www.surf.nl); neuroimaging data analysis was supported in part by Sophia Foundation Project S18-20 and Erasmus University Fellowship awarded to RLM.

### HGUGM

This work was supported by: Spanish Ministry of Science and Innovation, Instituto de Salud Carlos III (SAM16PE07CP1, PI16/02012, PI19/024), co-financed by ERDF Funds from the European Commission, “A way of making Europe”, CIBERSAM; Madrid Regional Government (B2017/BMD-3740 AGES-CM-2), European Union Structural Funds; European Union Seventh Framework Program under grant agreements, FP7-HEALTH-2013-2.2.1-2-603196 (Project PSYSCAN) and European Union H2020 Program under the Innovative Medicines Initiative 2 Joint Undertaking (grant agreement No 115916, Project PRISM, and grant agreement No 777394, Project AIMS-2-TRIALS), Fundación Familia Alonso, Fundación Alicia Koplowitz and Fundación Mutua Madrileña.

### HUBIN

The HUBIN study was funded by: Swedish Research Council (2003-5485, 2006-2992, 2006-986, 2008-2167, K2012-61X-15078-09-3, 521-2011-4622, 521-2014-3487, 2017-00949); regional agreement on medical training and clinical research between Stockholm County Council and the Karolinska Institutet; Knut and Alice Wallenberg Foundation.

### IMAGEN

This work received support from the following sources: the European Union-funded FP6 Integrated Project IMAGEN (Reinforcement-related behaviour in normal brain function and psychopathology) (LSHM-CT-2007-037286), the Horizon 2020 funded ERC Advanced Grant ‘STRATIFY’ (Brain network based stratification of reinforcement-related disorders) (695313), ERANID (Understanding the Interplay between Cultural, Biological and Subjective Factors in Drug Use Pathways) (PR-ST-0416-10004), BRIDGET (JPND: BRain Imaging, cognition Dementia and next generation GEnomics) (MR/N027558/1), Human Brain Project (HBP SGA 2, 785907), the FP7 project MATRICS (603016), the Medical Research Council Grant ‘c-VEDA’ (Consortium on Vulnerability to Externalizing Disorders and Addictions) (MR/N000390/1), the National Institute for Health Research (NIHR) Biomedical Research Centre at South London and Maudsley NHS Foundation Trust and King’s College London, the Bundesministeriumfür Bildung und Forschung (BMBF grants 01GS08152; 01EV0711; Forschungsnetz AERIAL 01EE1406A, 01EE1406B), the Deutsche Forschungsgemeinschaft (DFG grants SM 80/7-2, SFB 940, TRR 265, NE 1383/14-1), the Medical Research Foundation and Medical Research Council (grants MR/R00465X/1 and MR/S020306/1), the National Institutes of Health (NIH) funded ENIGMA (grants 5U54EB020403-05 and 1R56AG058854-01). Further support was provided by grants from: - the ANR (ANR-12-SAMA-0004, AAPG2019 – GeBra), the Eranet Neuron (AF12-NEUR0008-01 – WM2NA; and ANR-18-NEUR00002-01 – ADORe), the Fondation de France (00081242), the Fondation pour la Recherche Médicale (DPA20140629802), the Mission Interministérielle de Lutte-contre-les-Drogues-et-les-Conduites-Addictives (MILDECA), the Assistance-Publique-Hôpitaux-de-Paris and INSERM (interface grant), Paris Sud University IDEX 2012, the Fondation de l’Avenir (grant AP-RM-17-013), the Fédération pour la Recherche sur le Cerveau; the National Institutes of Health, Science Foundation Ireland (16/ERCD/3797), U.S.A. (Axon, Testosterone and Mental Health during Adolescence; RO1 MH085772-01A1), and by NIH Consortium grant U54 EB020403, supported by a cross-NIH alliance that funds Big Data to Knowledge Centres of Excellence.

### LBC1936

We thank the Lothian Birth Cohort 1936 members who took part in this study, and Lothian Birth Cohort 1936 research team members and radiographers who collected, entered and checked data used in this paper. Magnetic Resonance Image acquisition and analyses were conducted at the Brain Research Imaging Centre, Neuroimaging Sciences, University of Edinburgh (www.bric.ed.ac.uk) which is part of SINAPSE (Scottish Imaging Network—A Platform for Scientific Excellence) collaboration (www.sinapse.ac.uk) funded by the Scottish Funding Council and the Chief Scientist Office. The LBC1936 and this research are supported by Age UK (Disconnected Mind project), the UK Medical Research Council [MRC; G0701120, G1001245, MR/M013111/1, MR/R024065/1], and the University of Edinburgh.

### NCNG

The NCNG sample collection was supported by grants from the Bergen Research Foundation and the University of Bergen, the Dr Einar Martens Fund, the K.G. Jebsen Foundation, the Research Council of Norway, to SLH, VMS and TE.

### NESDA

The infrastructure for the NESDA study (www.nesda.nl) is funded through the Geestkracht program of the Netherlands Organisation for Health Research and Development (ZonMw, grant No 10-000-1002) and financial contributions by participating universities and mental health care organizations (VU University Medical Center, GGZ inGeest, Leiden University Medical Center, Leiden University, GGZ Rivierduinen, University Medical Center Groningen, University of Groningen, Lentis, GGZ Friesland, GGZ Drenthe, Rob Giel Onderzoekscentrum).

### NeuroIMAGE

The NeuroIMAGE study was supported by NIH Grant R01MH62873 (to Stephen V. Faraone), NWO Large Investment Grant 1750102007010 (to Jan Buitelaar), ZonMW grant 60-60600-97-193, NWO grants 056-13-015 and 433-09-242, and matching grants from Radboud University Nijmegen Medical Center, University Medical Center Groningen and Accare, and Vrije Universiteit Amsterdam. The research leading to these results also received support from the European Community’s Seventh Framework Programme (FP7/2007-2013) under grant agreement No 278948 (TACTICS), 602805 (Aggressotype), 603016 (MATRICS) and 602450 (Imagemend), and the Innovation Medicine Initiative grants 115300 (EU-AIMS) and 777394 (AIMS-2-TRIALS).

### NUIG

We would like to thank the radiologists at the University Hospital Galway and the participants who generously gave their time to make this study possible. The NUIG sample was supported and funded by the National University of Ireland Galway (NUIG) Millennium Fund and the Health Research Board (HRA_POR/2011/100).

### OATS

We gratefully acknowledge and thank the OATS participants, their supporters and the Research Team. The Older Australian Twin Study (OATS) is supported by the Australian NHMRC/Australian Research Council Strategic Award (Grant 401162) and the NHMRC Project grant 1405325. This study was facilitated through Twins Research Australia, a national resource in part supported by a Centre for Research Excellence from the NHMRC. DNA was extracted by Genetic Repositories Australia (NHMRC Grant 401184). Genomewide genotyping at the Diamantina Institute, University of Queensland, was partly funded by a CSIRO Flagship Collaboration Fund Grant.

### PAFIP

PAFIP was supported by the Instituto de Salud Carlos III (PI14/00639, PI14/00918 and PI17/01056) and Fundación Instituto de Investigación Marqués de Valdecilla (NCT0235832 and NCT02534363). No pharmaceutical company has financially supported the study.

### Rotterdam study

The GWAS datasets are supported by the Netherlands Organization of Scientific Research NWO Investments (nr. 175.010.2005.011, 911-03-012), the Genetic Laboratory of the Department of Internal Medicine, Erasmus MC, the Research Institute for Diseases in the Elderly (014-93-015; RIDE2), the Netherlands Genomics Initiative (NGI)/Netherlands Organization for Scientific Research (NWO) Netherlands Consortium for Healthy Aging (NCHA), project no. 050-060-810. We thank Pascal Arp, Mila Jhamai, Marijn Verkerk, Lizbeth Herrera and Marjolein Peters, MSc, and Carolina Medina-Gomez, MSc, for their help in creating the GWAS database, and Karol Estrada, PhD, Yurii Aulchenko, PhD, and Carolina Medina-Gomez, MSc, for the creation and analysis of imputed data. The Rotterdam Study is funded by Erasmus Medical Center and Erasmus University, Rotterdam, Netherlands Organization for the Health Research and Development (ZonMw), the Research Institute for Diseases in the Elderly (RIDE), the Ministry of Education, Culture and Science, the Ministry for Health, Welfare and Sports, the European Commission (DG XII), and the Municipality of Rotterdam. The authors are grateful to the study participants, the staff from the Rotterdam Study and the participating general practitioners and pharmacists.

### SHIP

The SHIP study is part of the Community Medicine Research net of the University of Greifswald, Germany, which is funded by the Federal Ministry of Education and Research (grants no. 01ZZ9603, 01ZZ0103, and 01ZZ0403), the Ministry of Cultural Affairs and the Social Ministry of the Federal State of Mecklenburg-West Pomerania. MRI scans in SHIP and SHIP-TREND have been supported by a joint grant from Siemens Healthineers, Erlangen, Germany and the Federal State of Mecklenburg-West Pomerania.

### Sydney MAS

We gratefully acknowledge and thank the Sydney MAS participants, their supporters and the Research Team. The Sydney Memory and Ageing Study (MAS) is supported by a National Health & Medical Research Council of Australia Program Grant (Grants 350833, 568969, 109308) and a Capacity Building Grant (Grant 568940). DNA samples were extracted by Genetic Repositories Australia, an Enabling Facility, which is supported by a National Health & Medical Research Council of Australia Grant, 401184.

### UK Biobank

This research has been conducted using the UK Biobank Resource under Application Number ‘11559’.

### UMCU

The UMCU cohort contains a.o. UTWINS and GROUP. UTWINS was funded by the Netherlands Organization for Health Research and Development ZonMw (908.02.123 and 917.46.370 to H.H.), and by the European Union Marie-Curie Research Training Network (MRTN-CT-2006-035987). The GROUP study is partially funded through the Geestkracht programme of the Dutch Health Research Council (Zon-Mw, grant No 10-000-1001), and matching funds from participating pharmaceutical companies (Lundbeck, AstraZeneca, Eli Lilly, Janssen Cilag) and universities and mental health care organizations (Amsterdam: Academic Psychiatric Centre of the Academic Medical Center and the mental health institutions: GGZ Ingeest, Arkin, Dijk en Duin, GGZ Rivierduinen, Erasmus Medical Centre, GGZ Noord Holland Noord. Groningen: University Medical Center Groningen and the mental health institutions: Lentis, GGZ Friesland, GGZ Drenthe, Dimence, Mediant, GGNet Warnsveld, Yulius Dordrecht and Parnassia psycho-medical center The Hague. Maastricht: Maastricht University Medical Centre and the mental health institutions: GGzE, GGZ Breburg, GGZ Oost-Brabant, Vincent van Gogh voor Geestelijke Gezondheid, Mondriaan, Virenze riagg, Zuyderland GGZ, MET ggz, Universitair Centrum Sint-Jozef Kortenberg, CAPRI University of Antwerp, PC Ziekeren Sint-Truiden, PZ Sancta Maria Sint-Truiden, GGZ Overpelt, OPZ Rekem. Utrecht: University Medical Center Utrecht and the mental health institutions Altrecht, GGZ Centraal and Delta.).

### UNSW

The UNSW study was supported by the Australian National Medical and Health Research Council (NHMRC) Program Grant 1037196, Project Grant 1066177, and the Lansdowne Foundation. We gratefully acknowledge the Janette Mary O’Neil Research Fellowship to JMF.

### Personal funding

ALWB received funding from the National Children’s Foundation Tallaght, Ireland. CD-C was supported by Instituto de Salud Carlos III, Juan Rodés Grant (JR19/00024). CEF was supported by R01 AG050595; R01 AG022381; P01 AG055367; R01R56 AG037985. DAl was supported by South-Eastern Norway Regional Health Authority (2019107). DJS is supported by the SAMRC. DvdM was supported by Research Council of Norway grant No 276082. EGJ was supported by Swedish Research Council (2003-5485, 2006-2992, 2006-986, 2008-2167, K2012-61X-15078-09-3, 521-2011-4622, 521-2014-3487, 2017-00949); regional agreement on medical training and clinical research between Stockholm County Council and the Karolinska Institutet; Knut and Alice Wallenberg Foundation; HUBIN project. ESP is supported by Hypatia Tenure Track Grant (Radboudumc); NARSAD Young Investigator Grant (Brain and Behavior Research Foundation ID:25034); Christine Mohrmann Fellowship. EV was supported by National Institute for Health Research (NIHR) Biomedical Research Centre at South London and Maudsley NHS Foundation Trust and King’s College London. FN was supported by German Research Foundation NE 1383/14-1. HB was supported by NHMRC Australia. GAS was supported by Conselho Nacional de Desenvolvimento Científico e Tecnológico (CNPq, Brazil; grant No 573974/2008-0), the Coordenação de Aperfeiçoamento de Pessoal de Nível Superior (CAPES, Brazil), the Fundação de Amparo à Pesquisa do Estado de São Paulo (FAPESP, Brazil; grant No 2008/57896-8) and the Fundação de Amparo à Pesquisa do Estado do Rio Grande do Sul (FAPERGS, Brazil). HJG has received research funding from the EU “Joint Programme Neurodegenerative Disorders” (JPND). HHHA was supported by the Netherlands Organization for Health Research and Development (ZonMW, grant No 916.19.151). IAB was supported by University of Sydney Post-graduate Award. IN was supported by DFG Ne2254/1-2. JBJK was supported by NHMRC Dementia Research Team Grant APP1095127. JH was supported by R21MH107327-01. JLS was supported by grant Nos R01MH118349, R01MH120125. JW was supported by the UK Dementia Research Institute which receives its funding from DRI Ltd, funded by the UK Medical Research Council, Alzheimer’s Society and Alzheimer’s Research UK (JW), the Row Fogo Charitable Trust through the Row Fogo Centre for Research into Ageing and the Brain (Ref No: AD.ROW4.35. BRO-D.FID3668413) and the Fondation Leducq Transatlantic Network of Excellence for the Study of Perivascular Spaces in Small Vessel Disease, ref no. 16 CVD 05. KLG was supported by grant No APP1173025. KS was supported by research grants from the National Healthcare Group, Singapore (SIG/05004; SIG/05028; SIG /1103), and the Singapore Bioimaging Consortium (RP C009/2006). LD was supported by R01AG059874 and R01MH117601. JHF was supported by SFB 940/2 and the German Ministry of Education and Research (BMBF Grants 01EV0711 & 01EE1406B). LHvdB was supported by the Netherlands ALS Foundation. OAA was supported by Research Council of Norway (223273), KG Jebsen Stiftelsen, H2020 CoMorMent (847776). LMOL was supported by K99MH116115. LP received funding from the German Research Foundation (DFG), the Ministry of Science and Education (BMBF) and EU. LTW is funded by the European Research Council under the European Union’s Horizon 2020 research and innovation program (ERC Starting Grant 802998), the Research Council of Norway (249795), the South-East Norway Regional Health Authority (2019101), and the Department of Psychology, University of Oslo. MK was supported by funding from the Dutch National Science Agenda NeurolabNL project (grant No 400-17-602). MLPM was supported by the French funding agency ANR (ANR-12-SAMA-0004), the Assistance-Publique-Hôpitaux-de-Paris and INSERM (interface grant), Paris-Descartes-University (collaborative-project-2010), Paris-Sud-University (IDEX-2012). MLS was supported by FAPESP: 2016/13737-0 and 2016/04983-7. MMN was supported by the German Research Foundation (DFG grant FOR2107; NO246/10-2). MNS was supported by the Deutsche Forschungsgemeinschaft (DFG grants TRR 265; SFB 940; SM 80/7-2) and the German Ministry of Education and Research (BMBF grants 01EV0711; 01EE1406B). MR was supported by DFG FOR2107 RI 908/11-1 & RI 908/11-2, BMBF Neuron Eranet Synschiz 01EW1810. MSK was supported by the National Health and Medical Research Council, Australia Project Grant (GNT1008080) and Career Development Fellowship (GNT1090148). MSP was supported by NIA R01AG02238.NJ and LD were supported by R01AG059874 and R01MH117601. PMT and SIT were supported by NIH U54 EB020403, R56AG058854 to the ENIGMA World Aging Center, R01MH116147 and P41EB015922. PRS was supported by National Health and Medical Research Council, Australia grant Nos 1037196, 1063960, 1176716. RA-A is funded by a Miguel Servet contract from the Carlos III Health Institute (CP18/00003), carried out on Fundación Instituto de Investigación Marqués de Valdecilla. PGF received funding from the German Research Foundation, the European Union and the Federeal Ministry of Science. RAB was supported by the European Research Council. SIB was supported by FAPESP 2016/04983-7; FAPESP 2011/50740-5; INCT (CNPq/FAPESP) 2014/50917-0. SEF was supported by the Max Planck Society. SEM was supported by NHMRC APP1103623; APP1172917; APP1158127. SHW was supported by DFG FOR2107 Wi3439/3-2, BMBF Neuron ERANET Synschiz 01EW1810. SLH was supported by the University of Bergen, Trond Mohn Research Foundation, Helse Vest. SRC was supported by a Sir Henry Dale Fellowship jointly funded by the Wellcome Trust and the Royal Society (Grant Number 221890/Z/20/Z). TEl was funded by the Research Council of Norway, the South-Eastern Norway Regional Health Authority, Oslo University Hospital and a research grant from Mrs. Throne-Holst. TH was supported by grants from the Interdisciplinary Center for Clinical Research (IZKF) of the medical faculty of Münster (grant MzH 3/020/20) and the German Research Foundation (DFG grants HA7070/2-2, HA7070/3, HA7070/4). TJ was supported by National Natural Science Foundation of China (81801773, 81930095, 91630314), the Shanghai Pujiang Project (18PJ1400900), the Key Project of Shanghai Science and Technology Innovation Plan (16JC1420402), the Shanghai Municipal Science and Technology Major Project (No.2018SHZDZX01) and ZHANGJIANG LAB. TRM was supported by Medical Research Council (UK). TW was supported by Netherlands Organization for Health Research and Development (ZonMw) TOP project No 91211021; Sophia Children’s Hospital Foundation (Stichting Vrienden van het Sophia) project No S18-68.VM was supported by CONICYT fellowships 21180871. UFM was supported by the Throne-Holst foundation. VMS was supported by Research Council of Norway (grant No 223273 NORMENT). WSK was supported by NIA grants R01 AG050595, R01 AG022381, R01AG060470, R01 AG054002, and NIAAA grant R01 AA026881.

## Author contributions

Conceptualization: BF, CEF, HEH, MSP, NJ, PMT, RMB, SEM, WSK. Central analysis and coordination: BF, CDW, ES, HEH, HGS, MK, KLG, NJ, PMT, RMB, SEM, SIT. Core writing team: BF, HEH, KLG, MK, NJ, PMT, RMB, SEM. Visualization: JT, MK, RMB. Cohort principle investigators: AH, AJ, AK, ALWB, APJ, BCF, BTB, BWJHP, CAr, CMcD, DAm, DIB, EGJ, FN, GAS, GSc, HB, HEHP, HF, HG, HHHA, HJG, HL, HW, IA, IN, J-LM, JH, JHV, JKBu, JMF, JO, JTr, KA, KS, LHvdB, LP, LTW, MAI, MHJ, MJW, MNS, MSK, OAA, PAG, PBM, PGF, PJH, PMP, PRS, PS, PSS, RAB, RAO, RLM, RMM, RSK, RW, SCa, SCic, SEF, SIB, SRC, TB, TEl, TEs, TH, TKi, TW, UD, UFM, WC. Imaging data collection: AG, ALWB, APJ, AZ, AvdL, BI, BM, CAM, CAh, CD-C, CJ, CMcD, DAm, DG, DIB, DJH, DMC, DT-G, EA, EBQ, EBø, EGJ, ESh, FN, FSte, GJB, GR, GSu, H-JW, HF, HHHA, IA, IAB, J-LM, JG, JH, JHF, JJa, JMF, JMW, JR, JTr, KA, KD, KS, LHvdB, LTW, MAI, MEB, MGJCK, MJW, MLPM, MNS, MSK, NEMvH, NO, NS, OAA, PAG, PD, PMP, PRJ, PS, RA-A, RB, RKL, RR, SDe, SH, SM, TRM, TW, TWM, UD, VOG, WH, WW. Genetic data collection: AJF, ALWB, BM, BTB, BWJHP, CAr, CD-C, CMcD, DIB, DWM, EA, EBQ, EGJ, FN, FSte, FStr, GH, GSu, HF, HHHA, IAB, J-LM, JBJK, JG-P, JH, JJH, JMF, JR, JTr, JV-B, KAM, KS, LHvdB, MDF, MJW, MLPM, MLS, MMN, MNS, MR, MSK, NS, OAA, PD, PMP, PRS, PS, RAO, RR, SCic, SDe, SEF, SHW, SIB, SLH, SM, TRM, TWM, UD, VMS. Imaging data analysis: AG, AHZ, APJ, ATh, AZ, BJO, BM, CAl, CGD, CJ, CLdM, DAl, DG, DK, DMC, DT-G, EBl, EELB, ESh, ESp, FN, GB, GSu, GVR, H-JW, HJG, IA, IAB, JJi, JKBr, JMW, KS, KW, LKMH, LN, LTW, MA, MAH, MGJCK, MSK, NAC, NEMvH, NJ, NS, NT, RB, RCWM, RMB, RR, SCiu, SIT, SJH, SMCdZ, SRC, ST, TJ, TKa, TW, UD, VM, WH, WW. Genetic data analysis: AJF, ATe, ATh, BM, BTB, CGD, CLdM, DvE, DvdM, EBl, ESp, EV, FStr, GB, GDa, GDo, GSu, GVR, JB, JBJK, JG-P, JLS, JMF, JPOFTG, JTe, KR, KS, LD, LMOL, MAI, MJK, MLS, MR, NJ, NJA, PRJ, RMB, RMT, SDa, SEM, SHW, SIB, SLH, SMCdZ, SP, TJ, YM.

## Additional information

Supplementary Data is available for this paper.

Correspondence and requests for materials should be addressed to Rachel M Brouwer

**Supplementary Table 1.** Cohort characteristics.

**Supplementary Table 2.** Description of imaging per study cohort.

**Supplementary Table 3.** Description of genetics per study cohort.

**Supplementary Table 4.** Summary of genome-wide significant SNPs and top-10 loci for main effect of genetic variants on brain morphology change rates in phase 1 + results for same SNPs in phase 2.

**Supplementary Table 5.** Summary of genome-wide significant SNPs and top-10 loci for main effect of genetic variants on brain morphology change rates in phase 2.

**Supplementary Table 6.** Summary of genome-wide significant SNPs and top-10 loci for linear age effects of genetic variants on brain morphology change rates in phase 1 + results for same SNPs in phase 2.

**Supplementary Table 7.** Summary of genome-wide significant SNPs and top-10 loci for linear age effects of genetic variants on brain morphology change rates in phase 2.

**Supplementary Table 8**. Summary of genome-wide significant SNPs and top-10 loci for quadratic age effects of genetic variants on brain morphology change rates in phase 1 + results for same SNPs in phase 2.

**Supplementary Table 9.** Summary of genome-wide significant SNPs and top-10 loci for quadratic age effects of genetic variants on brain morphology change rates in phase 2.

**Supplementary Table 10.** Summary of genome-wide significant genes and top-10 genes for brain morphology change rates in phase 1 + results for same genes in phase 2.

**Supplementary Table 11.** Summary of genome-wide significant genes, top-10 genes for brain morphology change rates in phase 2 sample, and look-up results for top 10 genes in cross-sectional data.

**Supplementary Table 12.** Biological functions for top SNPs and genes.

**Supplementary Table 13.** Summary of genome-wide significant effects and top-10 gene-sets for brain morphology change rates in phase 1 + results for same gene sets in phase 2.

**Supplementary Table 14.** Summary of genome-wide significant effects and top-10 gene-sets for brain morphology change rates in phase 2.

**Supplementary Table 15.** SNP-based heritabilities as estimated using LDSC.

**Supplementary Table 16.** Phenome-wide association results for genome-wide significant loci and genes.

**Supplementary Table 17.** Loci for age-(in)dependent effect on longitudinal brain changes in subgroups.

**Supplementary Table 18.** Genes for age-(in)dependent effect on longitudinal brain changes in subgroups.

**Extended Data Movie.** Rates of change for brain structure throughout the lifespan.

**Extended Data Figure 1: Demographics and analysis**

Overview of demographics (left). Per cohort, an age distribution is displayed, based on mean and standard deviation of the age at baseline. Cohorts of European ancestry are displayed in green, non-European cohorts are displayed in yellow. On the right, the total number of included subjects is displayed and a pie-chart of the distribution of diagnostic groups (pink) and subjects not belonging to diagnostic groups - often healthy subjects (aqua). Overview of analysis pipeline (right).

**Extended Data Figure 2: Correlations between change rates**

Pearson correlations between rates of change and between baseline intracranial volume and rates of change in the largest adolescent cohort (top) and the largest cohort in older age (bottom) in phase 1. The size of the correlations is displayed by color and size of the circles.

**Extended Data Figure 3: Phenotype and GWAS overview**

Top: Change rates per cohort and estimated trajectories of the change rate with confidence intervals (in green) are displayed above. The size of the points represents the relative size of the cohorts. Cohorts that were added in phase2 are displayed in grey. Only cohorts that satisfy N>75 are shown. The estimated trajectories of the volumes themselves are displayed below, for all subjects (solid line) and healthy subjects only (dashed line). Bottom rows contain Manhattan plots and QQ-plots for age-independent, age linear and age quadratic GWASses for rate of change. 3A: Amygdala; 3B: Caudate; 3C: Cerebellum grey matter; 3D: Cerebellum white matter; 3E: Cerebral white matter; 3F: Cortex volume; 3G: Cortical thickness; 3H: Hippocampus; 3I: Lateral ventricles; 3J: Nucleus accumbens; 3K: Pallidum; 3L: Putamen; 3M: Surface area; 3N: Thalamus; 3O: Total brain.

**Extended Data Figure 4: Locusplots, eQTL and chromatin interaction mapping for genome-wide significant loci.**

S4: Locusplots, eQTL and chromatin interaction mapping for genome-wide significant loci. A) rs1425034; change rate pallidum; independent of age; B) rs73210410; change rate pallidum; linear age dependency; C) rs11726181; change rate cerebral white matter; quadratic age dependency; D) 5:157751672; change rate surface area; linear age dependency*; E) rs10990953; change rate ventricles; independent of age; F) rs17809993; change rate cortex volume quadratic age dependency; G) rs17809993; change rate cortical thickness; quadratic age dependency; H) rs12019523; change rate caudate; quadratic age dependency; I) 13:72353395; change rate cerebral white matter volume; quadratic age dependency; note that this SNP was not in the reference dataset containing LD structure; displayed LD structure is based on 13:7234009, R2 = 0.87 with the top-SNP J) rs72772746; change rate lateral ventricles; independent of age; K) rs12325429; change rate total brain volume; independent of age; L) rs34342646; change rate surface area; quadratic age dependency; M) rs429358; change rate hippocampus; quadratic age dependency. Locus plots were created with locus zoom^95^. Circos plots were created with FUMA^66^. *not present in 1000G reference file, no circos plot available.

**Extended Data Figure 5: Top SNP results for genome-wide significant SNPs and genes**

Age-(in)dependent effect of the significant SNPs/top-SNPs in significant genes. The top figure displays a forest-plot for age-independent effects, or the estimated effect of the tested allele on the change rate in each cohort against age otherwise. In the latter case, the red line displays the estimated age-effect with 95% confidence interval from the meta analysis/meta-regression. The bottom figure shows a visualization of the effect of the tested allele on the phenotype itself. The red line represents the lifespan trajectory for the carriers of the effect allele, the black line represents the lifespan trajectory of the non-carriers. A) rs1425034; change rate pallidum; independent of age; B) rs73210410; change rate pallidum; linear age dependency; C) rs11726181; change rate cerebral white matter; quadratic age dependency; D) rs10990953; change rate ventricles; independent of age; E) *FAU* – top SNP rs769440 change rate cerebellum white matter; linear age dependency; F) rs17809993; change rate cortex volume quadratic age dependency; G) rs17809993; change rate cortical thickness; quadratic age dependency; H) rs12019523; change rate caudate; quadratic age dependency; I) *DACH1* and 13:72353395; change rate cerebral white matter volume; quadratic age dependency; J) *GPR139* and rs72772746; change rate lateral ventricles; independent of age; K) rs12325429; change rate total brain volume; independent of age; L) *APOE* – top SNP rs769449; change rate surface area; quadratic age dependency; M) rs34342646; change rate surface area, quadratic age dependency; N) *APOE* and rs429358; change rate hippocampus; quadratic age dependency O) *APOE* – top SNP rs429358; change rate amygdala; linear* age dependency.

*\*APOE* also showed a significant quadratic age dependency for change rate of amygdala, the most parsimonious model is shown.

**Extended Data Figure 6: Lookup longitudinal versus cross-sectional GWAS**

Expected versus actual overlap for the first top-1000 ranked genes. Results from age-independent analysis(red), linear age -dependent analysis (green) and quadratic age-dependent analysis (blue) are shown in one figure. Top-N ranks are marked for nominally (dots) or FDR-corrected (within the top-1000 genes for this phenotype; triangles) significance for over- or underrepresentation of genes associated with brain structural rates of change amongst the top-N ranked genes for cross-sectional brain measures. For lateral ventricles and cerebellum grey and white matter, summary statistics for the cross-sectional phenotype were only available for left and right lateral and inferior lateral ventricle, and left and right cerebellum grey and white matter, separately. Therefore, for those measures we show curves for overlap with the separate cross-sectional phenotypes.

**Extended Data Figure 7: Overlap with other phenotypes**

iSECA results for overlap between GWAS summary statistics of structural brain change with GWAS summary statistics of other phenotypes testing for pleiotropy (A), concordance and discordance of effects (B) and pleiotropy in the subgroup excluding subjects with a diagnosis (C). For comparison, we also present the same analysis for cross-sectional volumes, again showing pleiotropy results (D), concordance and discordance (E). Colors display the significance level on a 10-log scale. Associations that are significant based are marked with *. For a fair comparison, the cross-sectional analyses (D-E) used the same significance threshold as the change analyses (A-C); even though the latter contained more brain structures.

**Extended Data Figure 8: Gene expression for prioritized genes**

Heatmaps display normalized expression value (zero mean normalization of log2 transformed expression) for prioritized genes, for GTEx v8 RNAseq data (A) and BrainSpan data (B). Darker red means higher relative expression of that gene in each label, compared to a darker blue color in the same label. Note that *PVRL2* is an alias for *NECTIN2*.

**Extended Data Figure 9: PheWas results for study-wide significant genes**

*APOE* (A), *CAB39L* (B), *CDH8* (C), *DACH1* (D), *FAU* (E), *GPR139* (F), *NECTIN2* (G) and *SORCS2* (H). PheWAS plots show the significance of a gene on a range of traits based on MAGMA gene-based tests (Bonferroni corrected P-value threshold: 7.51e-07), as obtained from GWASAtlas^32^ (https://atlas.ctglab.nl). Redundant traits were removed for visualization and trait names were shortened. Full list of significant gene-based associations is in Supplementary Table 16.

**Extended Data Figure 10: Associations between absolute and relative change rates.**

Scatter plots showing the SNP effects for hippocampus change rate (absolute, x-axes) and hippocampus change rate divided by intracranial volume (relative; y-axes) for the three cohorts added in phase 2. SNPs were clumped at r2 < 0.1 for visualization purposes.

**Extended Data Figure 11: Power to detect top SNPs under multiplicative scanner effects.**

For each of the genome-wide significant SNPs in phase2, we simulated multiplicative scanner effects (independently drawn from N(1,s) per cohort; repeated 1000 times) and applied these to the original effect sizes and standard errors per cohort, after which we recalculated the meta-analysis or meta-regression. The x-axis shows the variation of the simulated effects, the y-axis shows the percentage of cases where the top-findings were still significant. Colors represent the different SNPs. The black squares are the average power over all SNPs tested.

**Extended Data Figure 12: QQ plots separately for each participating cohort**

QQ plots of summary statistics uploaded per phenotype and per participating cohort, showing expected (x-axis; under the null hypothesis of no genetic signal) versus observed (y-axis) minus log10-transformed p-values.

