## Supplementary Note & Suppelementary Figure legends for "Age-dependent genetic variants associated with longitudinal changes in brain structure across the lifespan"

**Supplementary Note and Extended Data Figures**

*Correction for intracranial volume*

It is common practice in studies of (cross-sectional) brain volumes to correct for intracranial volume or total brain volume: either to be sure that the differences that are found are region specific and not caused by a difference in overall brain size, or to be able to distinguish developmental effects (maximal brain size, approximated by the intracranial volume) from effects that occur later in life. When considering longitudinal changes over time, the choice of whether one has to correct translates into the question whether one wants to investigate absolute change (in e.g. ml/year, no correction) versus relative change (in %/year, correcting for baseline volume). However, neither total brain volume nor intracranial volume^96^ are stable across the lifespan, which complicates interpretation when these are used as correction factors^97^, especially when one combines data from across the lifespan. Therefore, we chose not to include overall brain volume in our main GWAS.

As a post-hoc analysis, we assessed the influence of adding intracranial volume to our analysis in the replication cohorts. To this end, we calculated relative annual change by subtracting baseline volume from follow-up volume, and divided by follow-up duration in years times intracranial volume at baseline. We excluded the phenotypes mean cortical thickness and total surface area from these analyses, since these are global measures themselves and a one- or two- dimensional brain measure should not be corrected by a three-dimensional volume. The association between relative and absolute annual change was very high in all three cohorts (all correlations > 0.97), which can be explained by intracranial volume having a much lower standard deviation compared to the mean than annual change rates. This implies that applying a correction for ICV to annual change rates is approximately a scaling factor which should not influence the GWAS findings. As expected, the effect estimates and p-values from the GWAS for relative and absolute annual change were also highly correlated (> 0.96; Extended Data Figure S10).

*Potential scanner effects*

Combining imaging data from multiple sources could be subject to biases driven by scanner manufacturer of scanning sequence, which could be a problem if data included in the study is not well distributed across different acquisitions with respect to the contrast of interest. Different approaches for pooling of imaging data have been proposed, including pre-acquisition harmonization of protocols, or post-acquisition methods (e.g. COMBAT^98^). The latter requires access to individual data which was not feasible for our study. However, through our design of meta-analysis of change rates, several potential scanner effects are already accounted for. First, we would like to mention that all results are based on the effect of a genetic variant *within a cohort* and the mean of a change rate in a cohort is therefore not influencing our genetic findings. That is, a potential within-cohort offset between baseline and follow-up measurement does influence the average change rate, but not the GWAS output, as the latter is essentially based on the ranking between subjects.

We further argue that potential additive and multiplicative scanner effects (as are assumed in COMBAT) will not bias our meta-analytic findings: additive scanner effects (an offset) is already accounted by computation of the annual rate of change, since there we subtract the follow-up from the baseline volume. Any multiplicative factors would not influence the Z-scores (or p-values) within a cohort as those are present in both the SNP-effect and the associated standard error. This implies that a potential scanner effect will only affect the weights in our meta-analyses, and this will not change the null distribution. Therefore, if anything, potential scanner effects will introduce noise, but should not introduce bias.

For our top-findings, we ran a simulation analyses incorporating potential multiplicative scanner effects which indeed show that our main findings are robust to relatively large perturbations: we assumed a multiplicative effect on the beta-SNP estimates for each cohort/scanner, by simulating a multiplicative factor from a standard normal distribution with mean 1 and a standard deviation ranging from 0.01 to 0.05. This multiplication factor should be interpreted as a deviation from the true volume, i.e. a factor of 0.95 suggests that brain volume is underestimated in all subjects (more so for larger brains than for smaller brains in absolute terms). Hence, also the change rate and associated SNP-beta obtained from GWAS is underestimated. The upper bound of 0.05 corresponds to the extreme case where relatively large perturbations occur regularly (in 5% of cases, the error will be larger than 10% of the volume). After simulating the multiplication factor for each cohort, we multiplied mean and standard error by this factor and repeated the meta-analysis or meta-regression and repeated for 1000 times. This analysis was done for all our genome-wide significant findings. Extended Data Figure 11 shows the percentage of cases where the top-findings were still significant, as a function of the variation of simulated multiplicative scanner effects. Power to detect our findings was generally high, indicating that our findings are robust under such modest perturbations.
