## Supplementary Figures 1-12 for "Age-dependent genetic variants associated with longitudinal changes in brain structure across the lifespan"

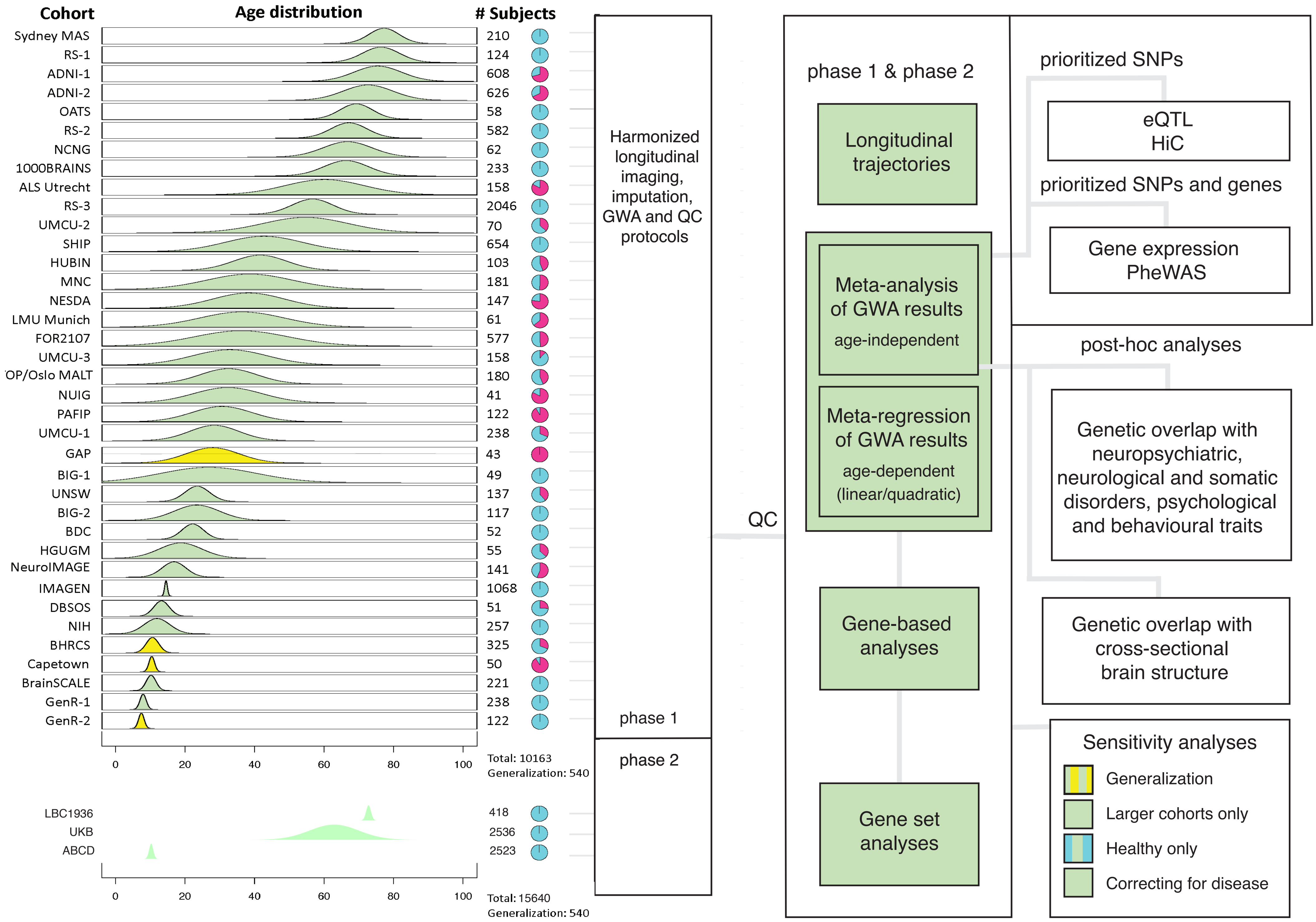
Cohort
Age distribution
### Subjects
Sydney MAS 210
RS-1 124
ADNI-1 608
ADNI-2 626
OATS 58
RS-2 582
NCNG 62
1000BRAINS 233
ALS Utrecht 158
RS-3 2046
UMCU-2 70
SHIP 654
HUBIN 103
MNC 181
NESDA 147
LMU Munich 61
FOR2107 577
UMCU-3 158
OP/Oslo MALT 180
NUIG 41
PAFIP 122
UMCU-1 238
GAP 43
BIG-1 49
UNSW 137
BIG-2 117
BDC 52
HGUGM 55
NeuroIMAGE 141
IMAGEN 1068
DBSOS 51
NIH 257
BHRCS 325
Capetown 50
BrainSCALE 221
GenR-1 238
GenR-2 122
0 20 40 60 80 100
Total: 10163
Generalization: 540
LBC1936 418
UKB 2536
ABCD 2523
0 20 40 60 80 100
Total: 15640
Generalization: 540
Harmonized longitudinal imaging, imputation, GWA and QC protocols
phase 1
phase 2
QC
phase 1 & phase 2
Longitudinal trajectories
Meta-analysis of GWA results
age-independent
Meta-regression of GWA results
age-dependent (linear/quadratic)
Gene-based analyses
Gene set analyses
prioritized SNPs
eQTL
HiC
prioritized SNPs and genes
Gene expression
PheWAS
post-hoc analyses
Genetic overlap with neuropsychiatric, neurological and somatic disorders, psychological and behavioural traits
Genetic overlap with cross-sectional brain structure
Sensitivity analyses
Generalization
Larger cohorts only
Healthy only
Correcting for disease

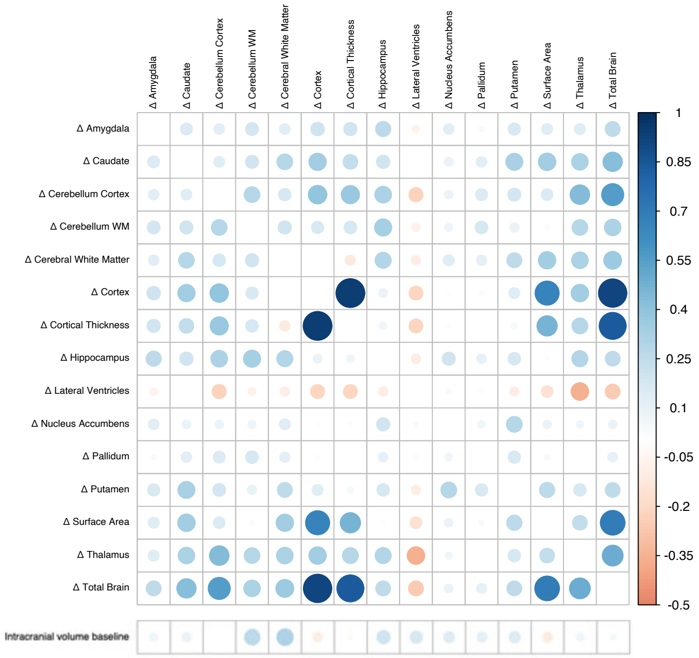


late adolescence
(IMAGEN)

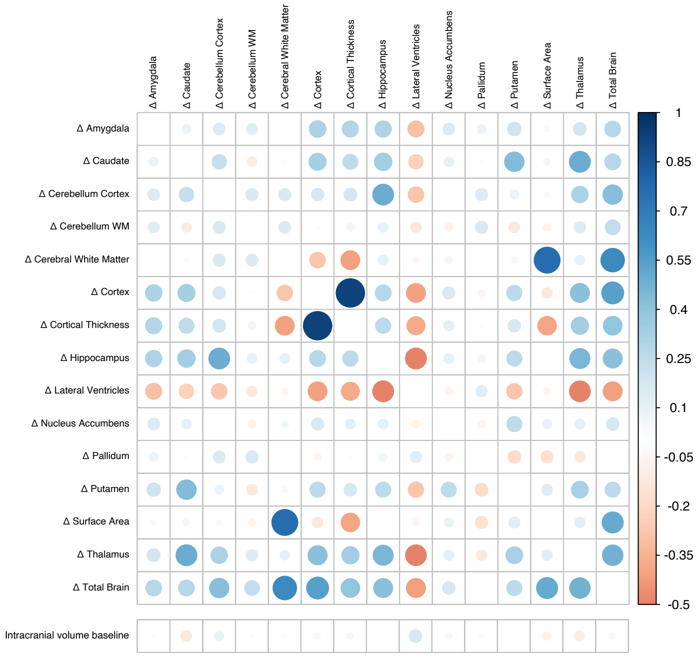


older age
(ADNI2)

### 3A: change rate amygdala volume

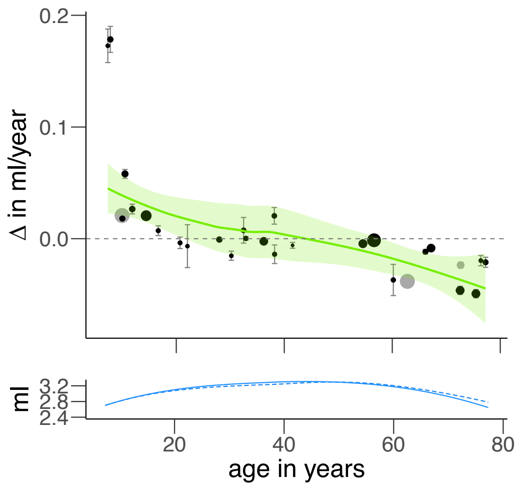


age
independent

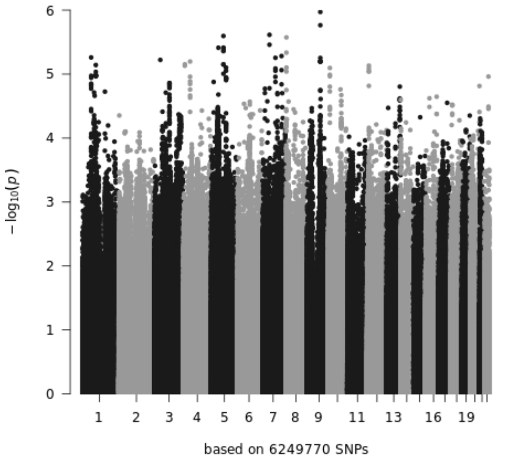


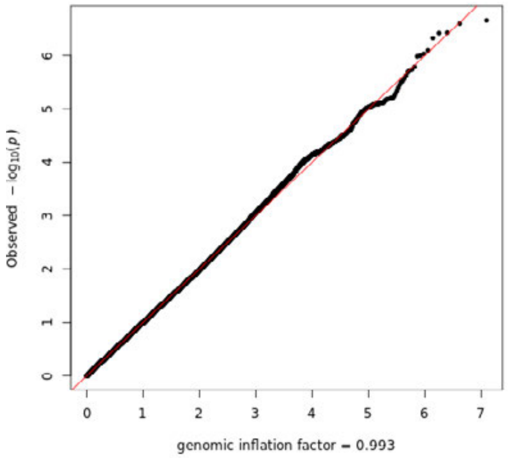


linear
age-dependent

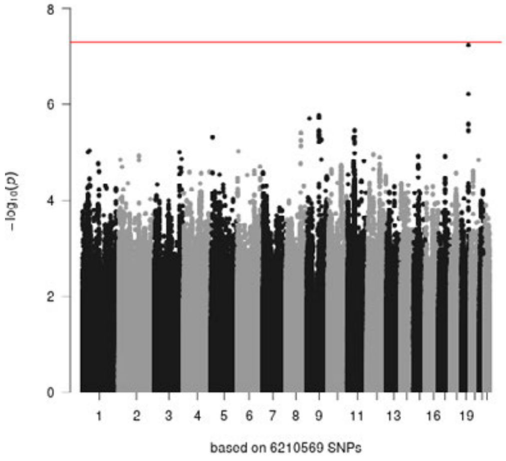


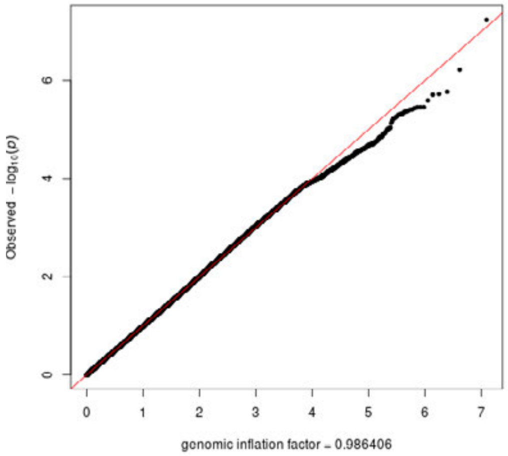


quadratic
age-dependent

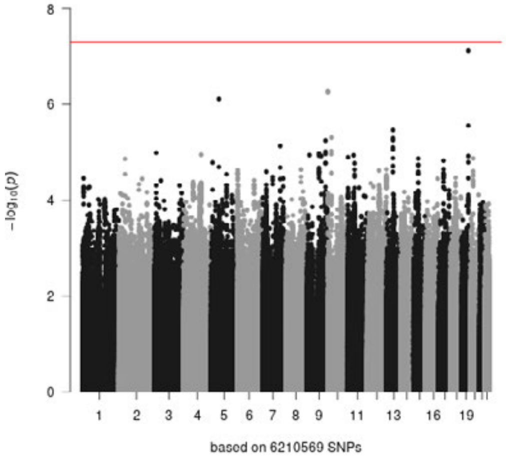


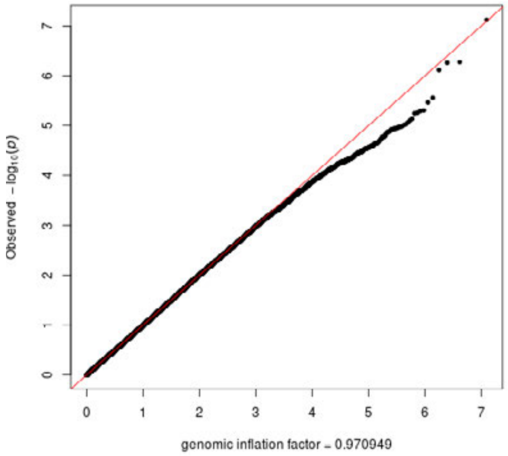

### 3B: change rate caudate volume

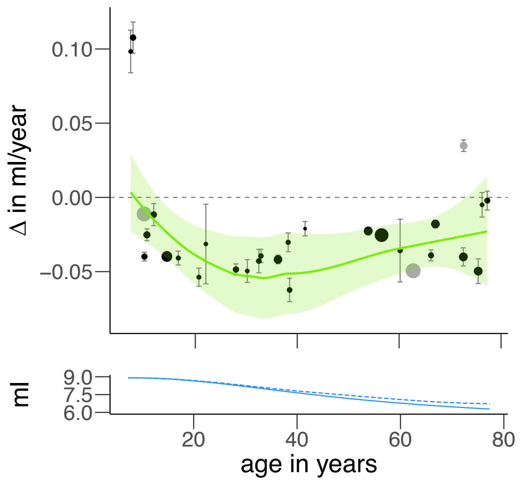


age
independent

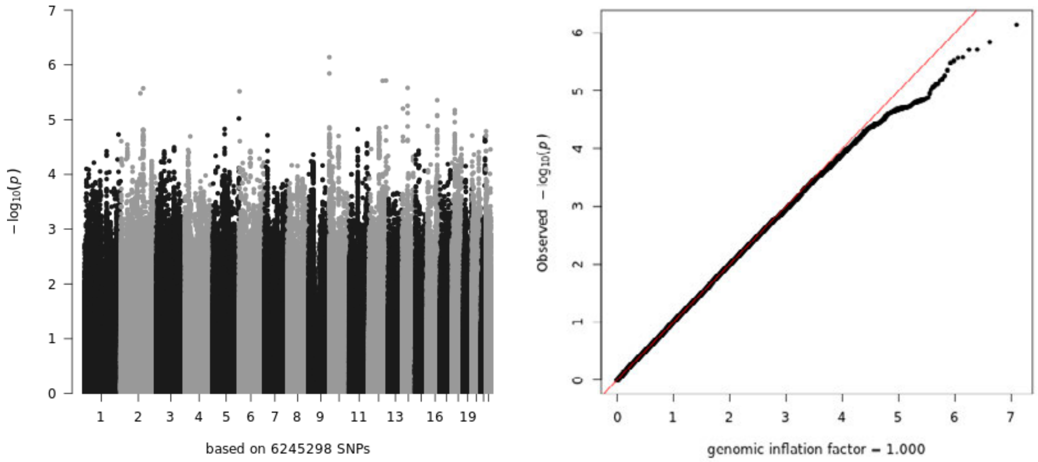


linear
age-dependent

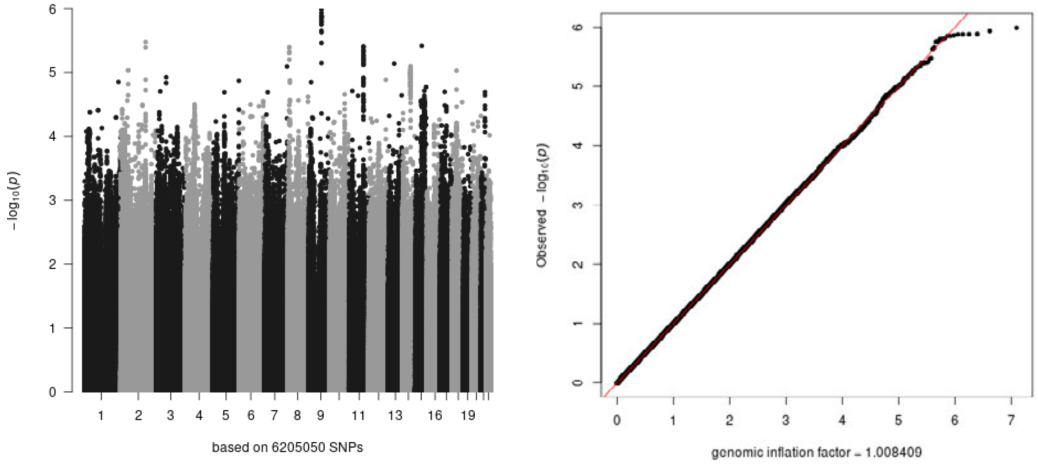


quadratic
age-dependent

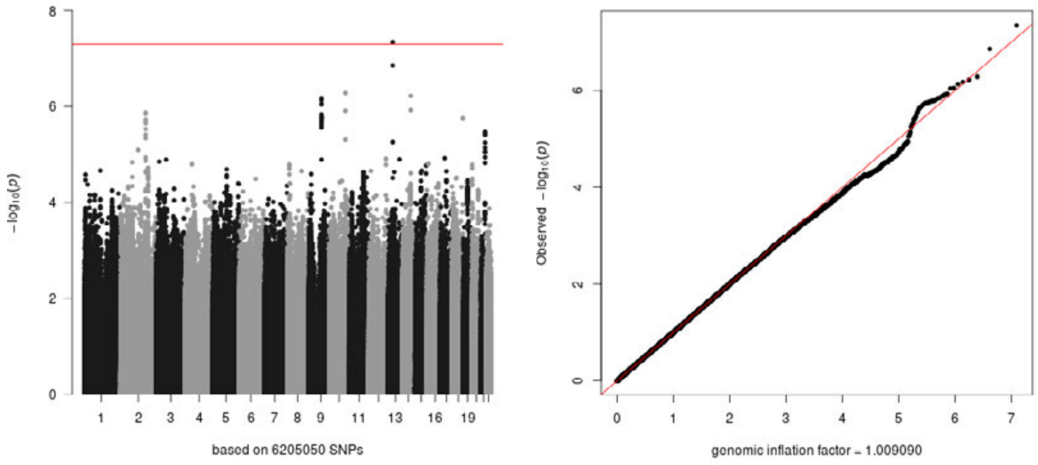

### 3C: change rate cerebellum white matter volume

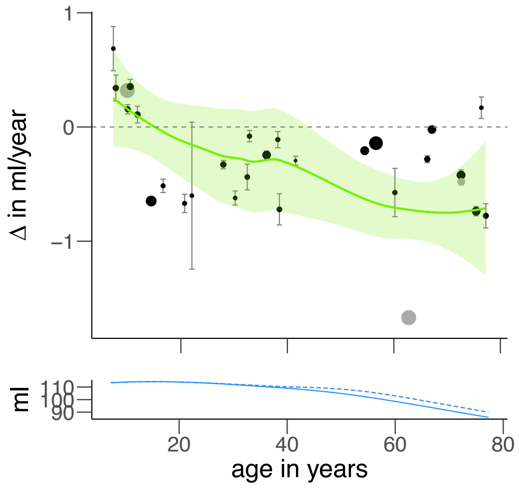


age independent

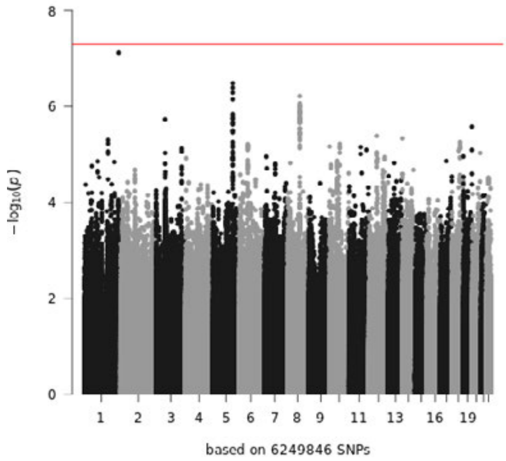


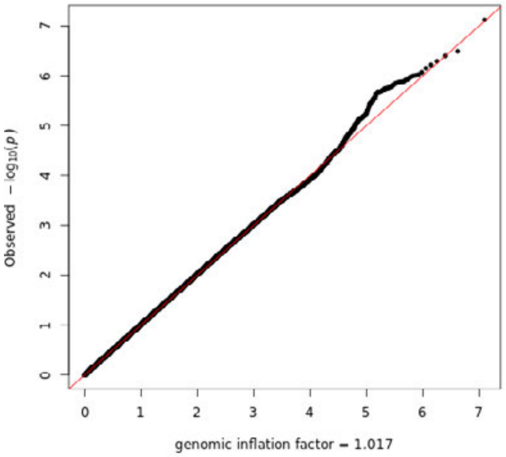


linear age-dependent

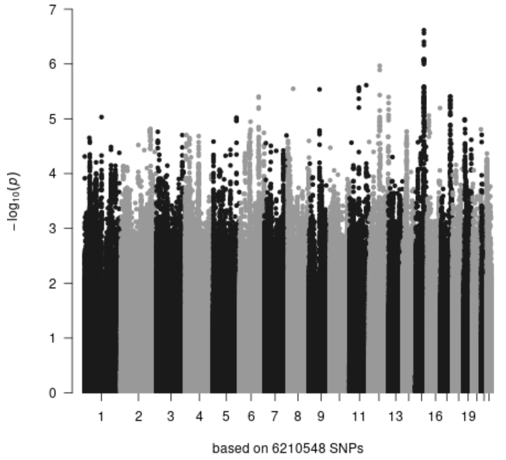


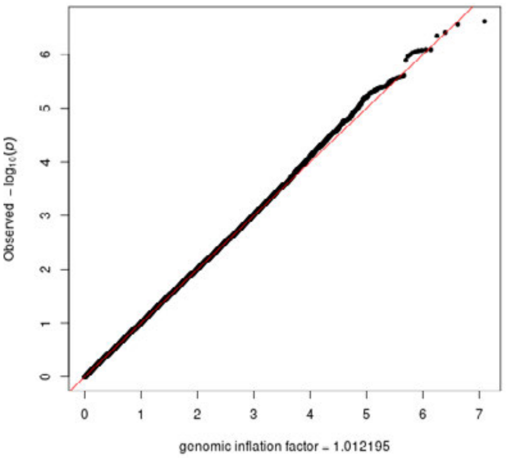


quadratic age-dependent

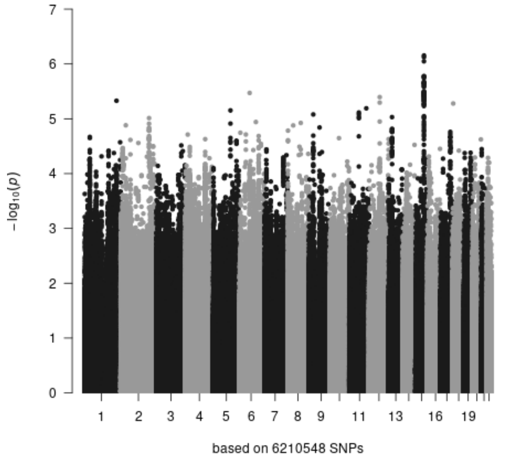


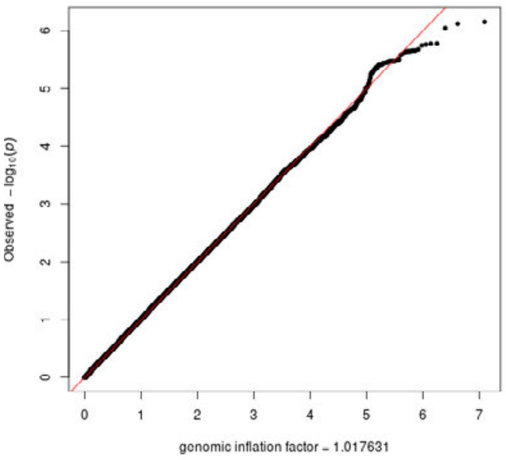

### 3D: change rate cerebellum gray matter volume

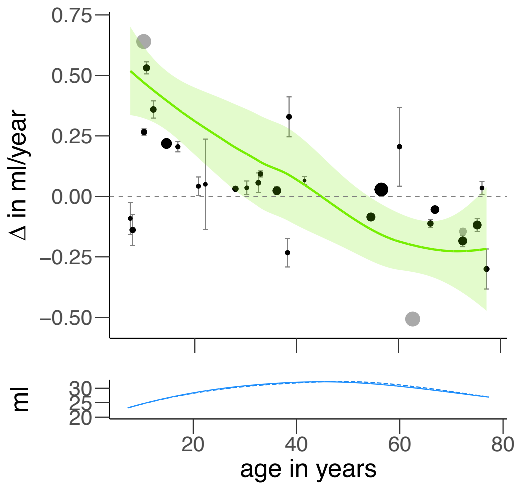


age independent

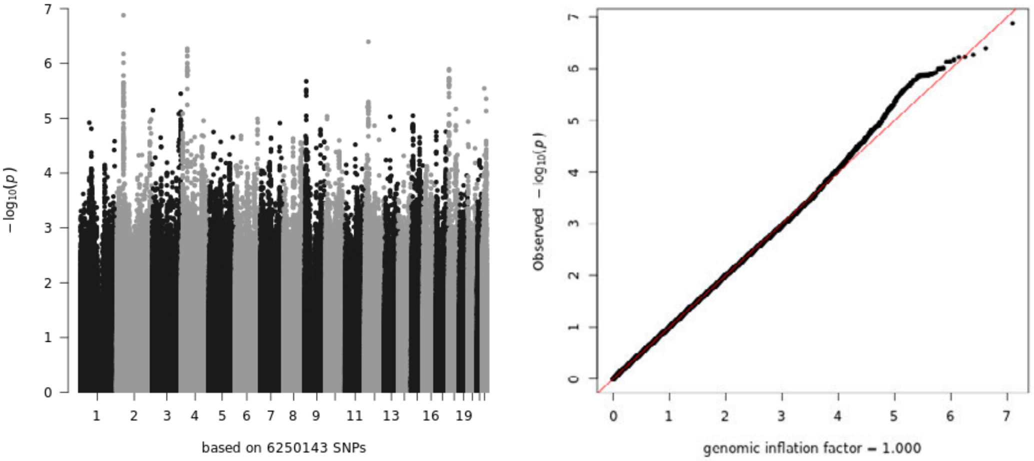


linear age-dependent

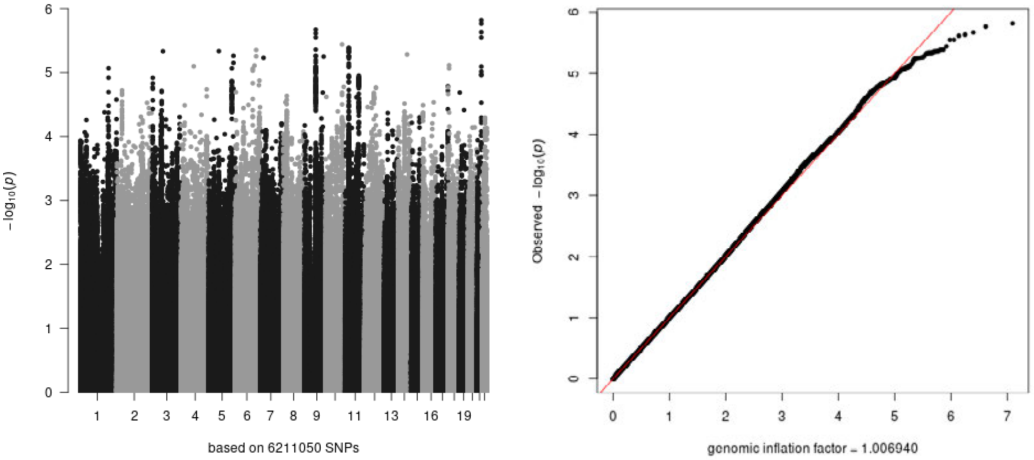


quadratic age-dependent

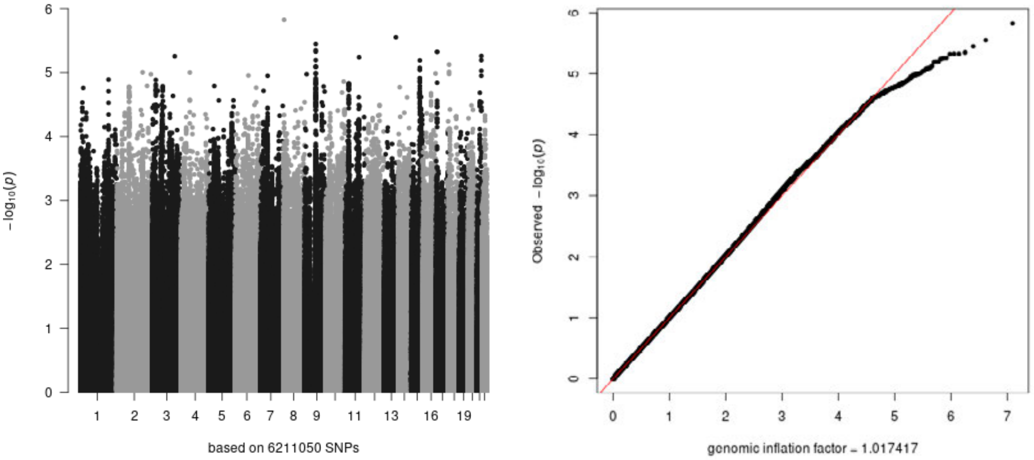

### 3E: change rate cerebral white matter volume

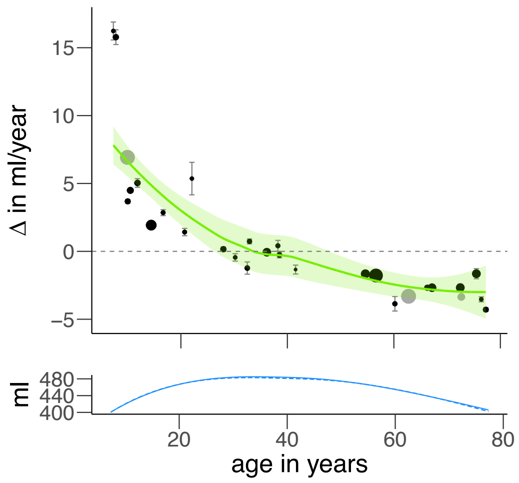


age independent

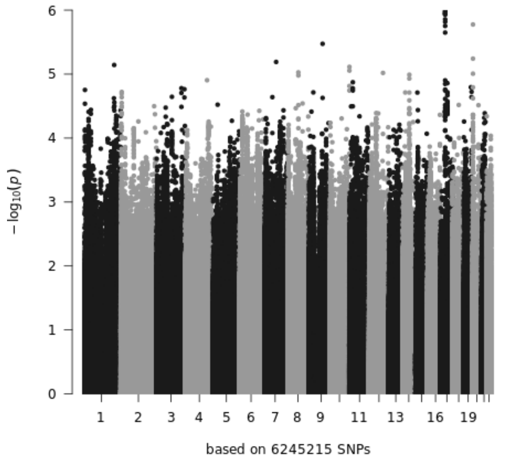


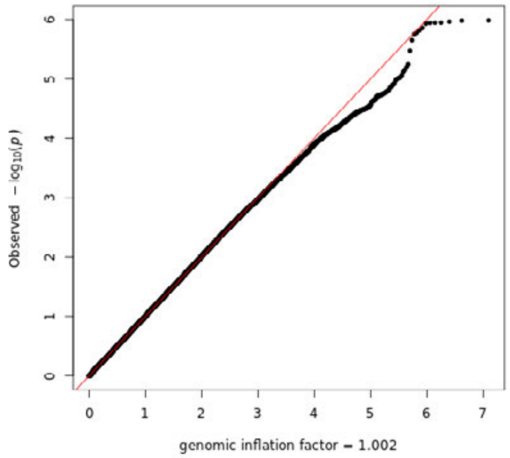


linear age-dependent

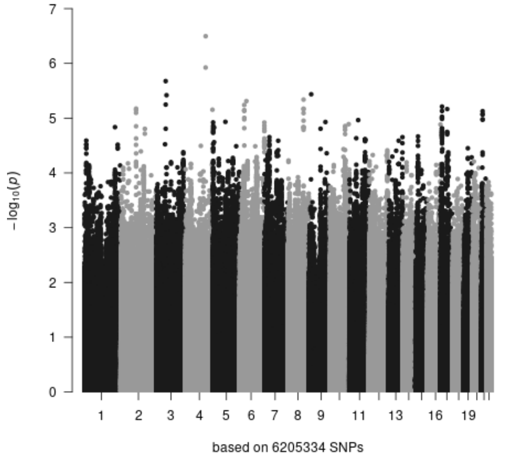


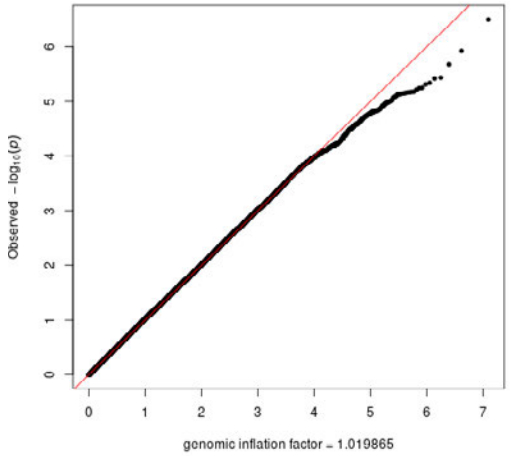


quadratic age-dependent

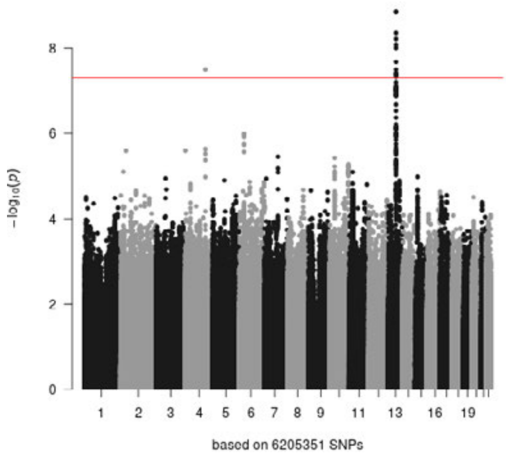


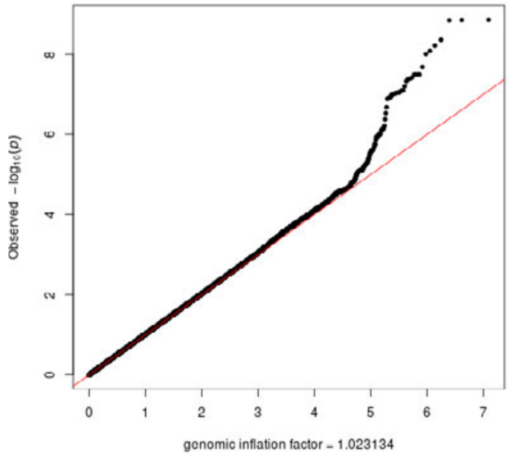

### 3F: change rate cortex volume

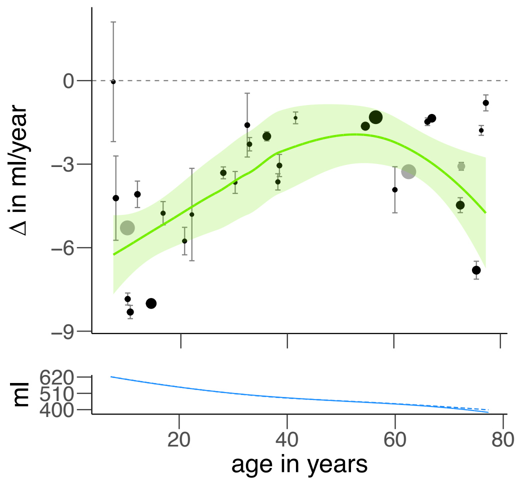


age
independent

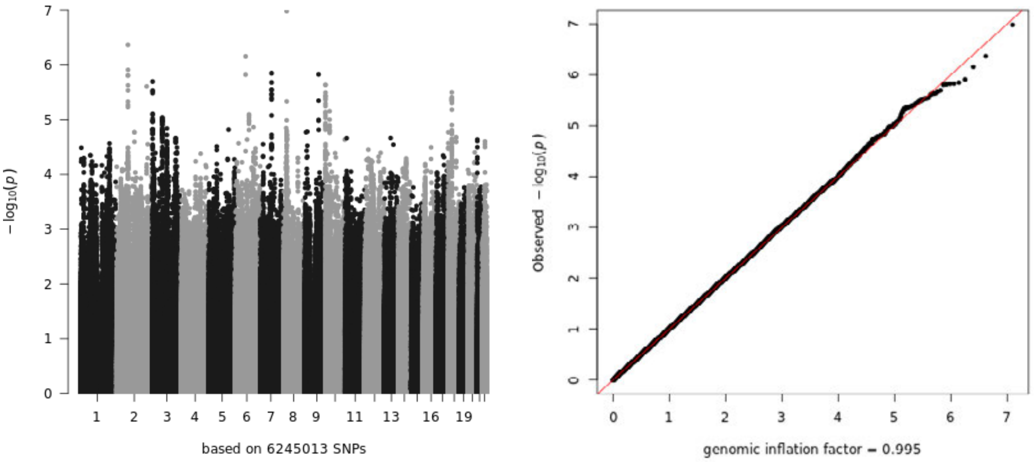


linear
age-dependent

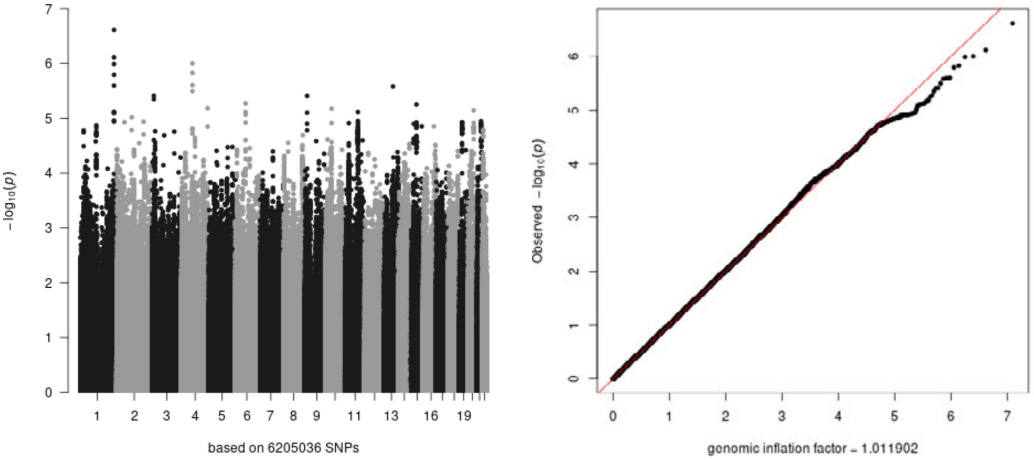


quadratic
age-dependent

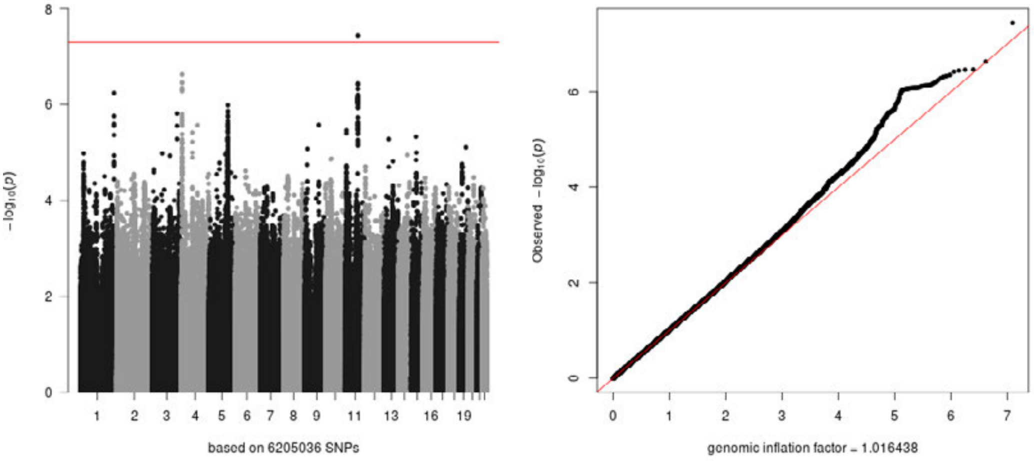

### 3G: change rate cortical thickness

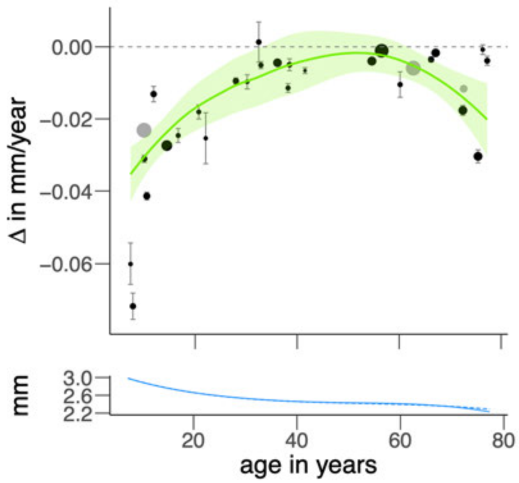


age
independent

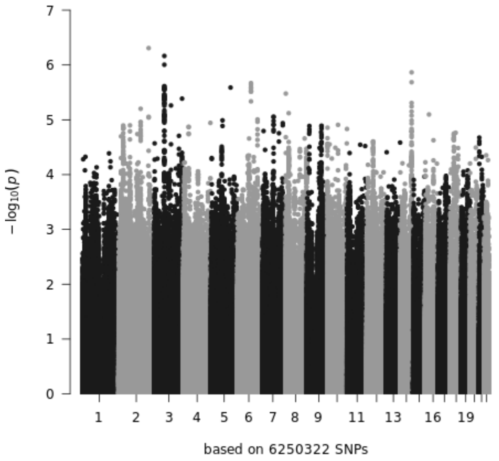


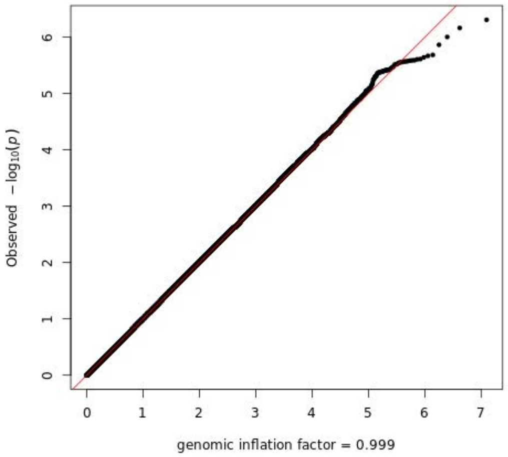


linear
age-dependent

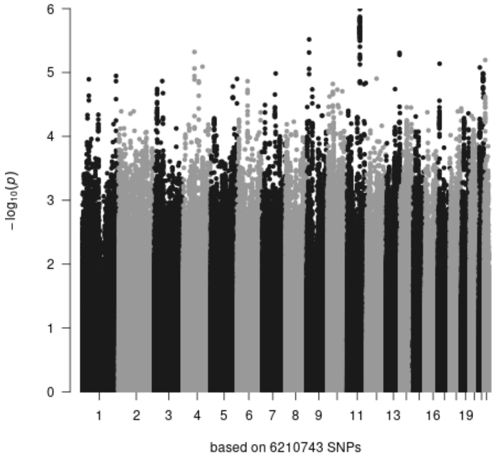


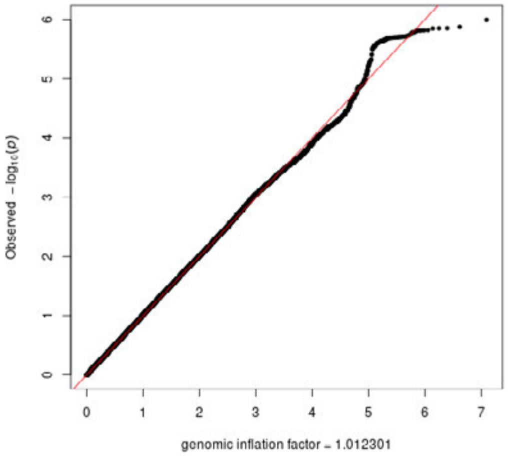


quadratic
age-dependent

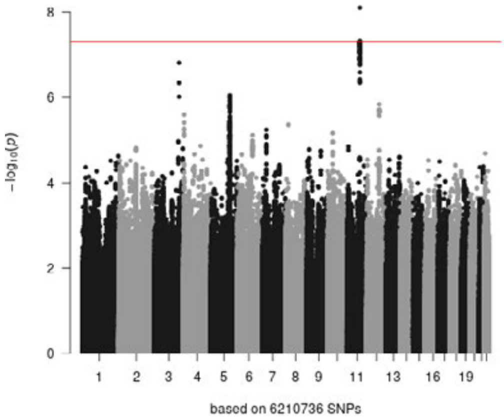


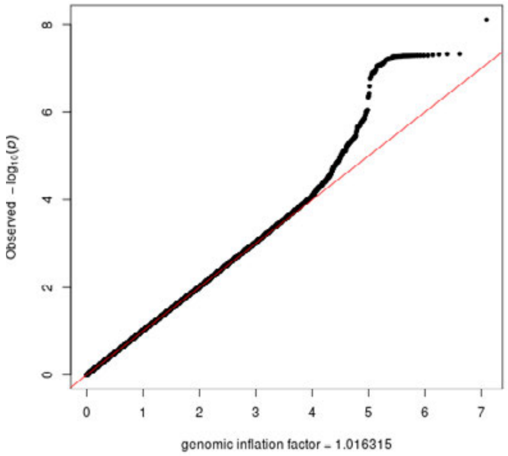

### 3H: change rate hippocampus

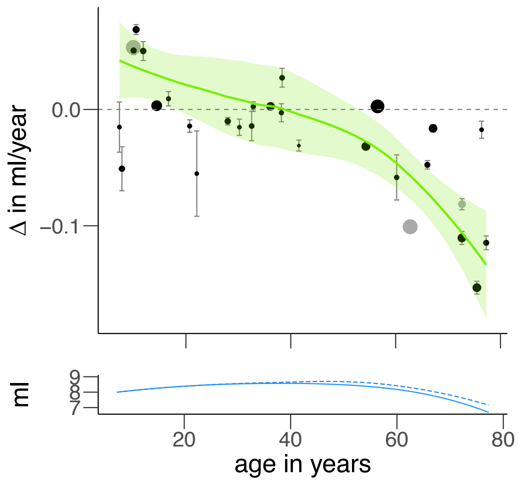


age
independent

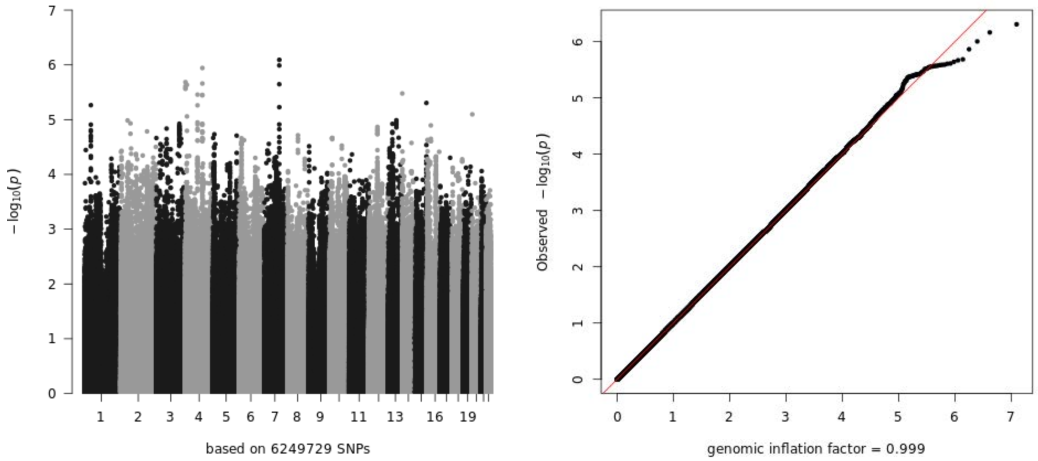


linear
age-dependent

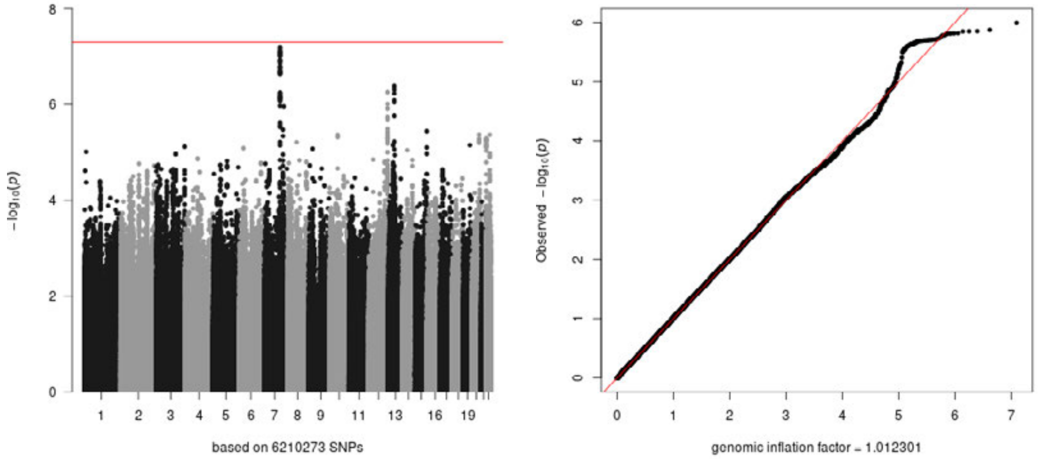


quadratic
age-dependent

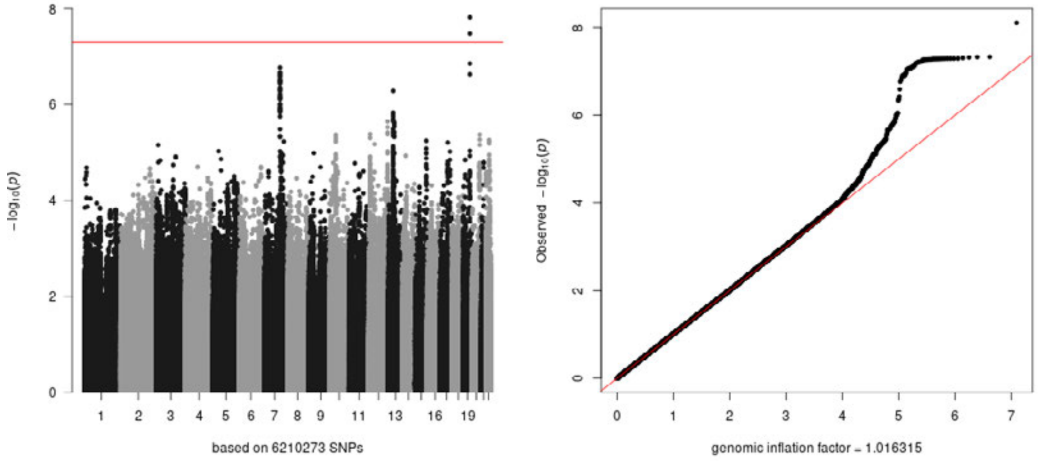

### 3I: change rate lateral ventricle volume

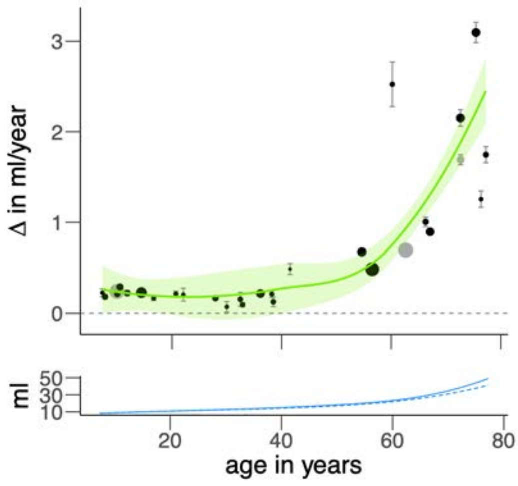


age
independent

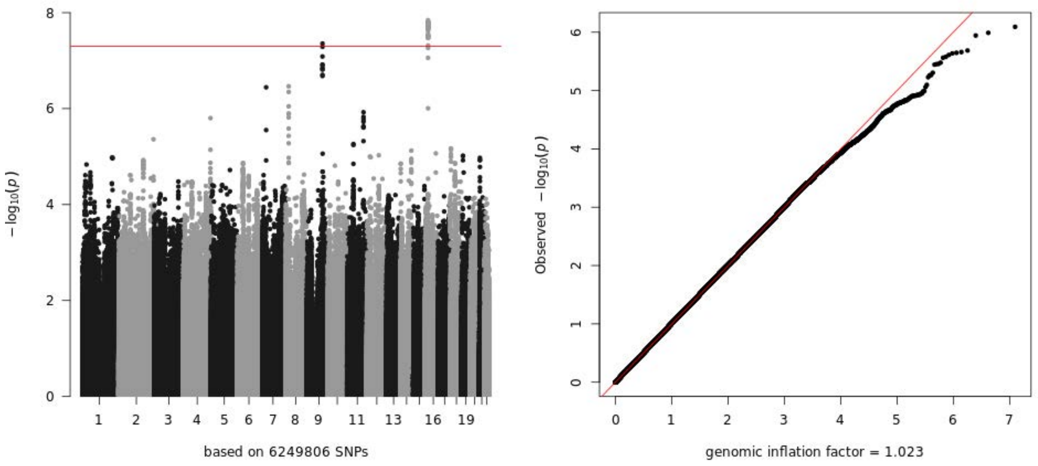


linear
age-dependent

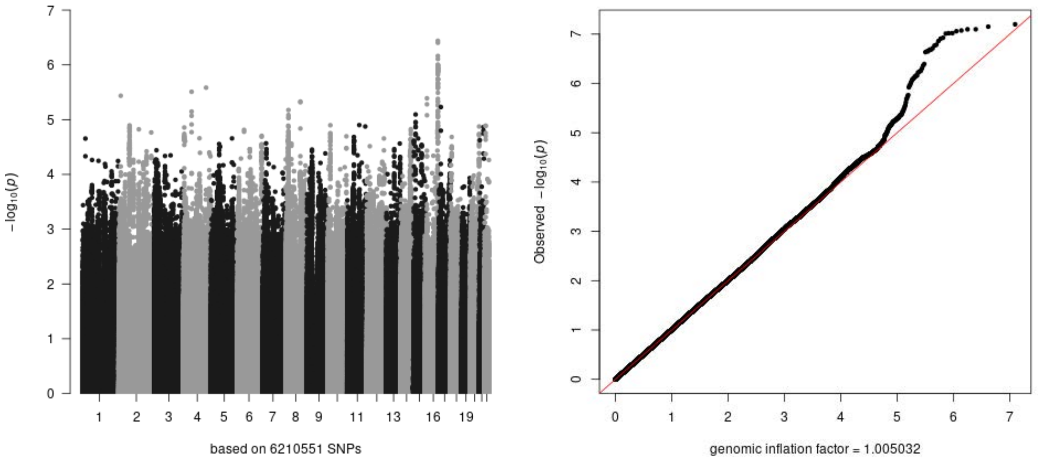


quadratic
age-dependent

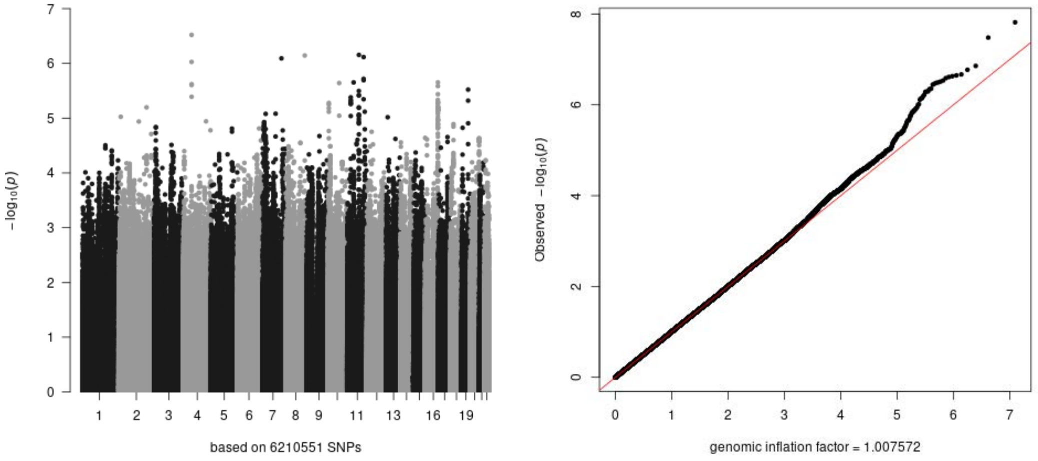

### 3J: change rate nucleus accumbens volume

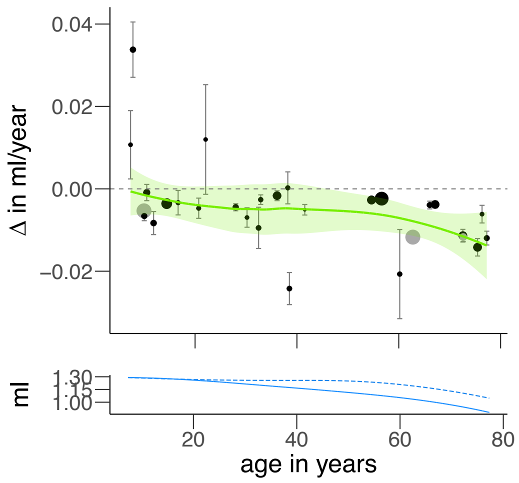


age
independent

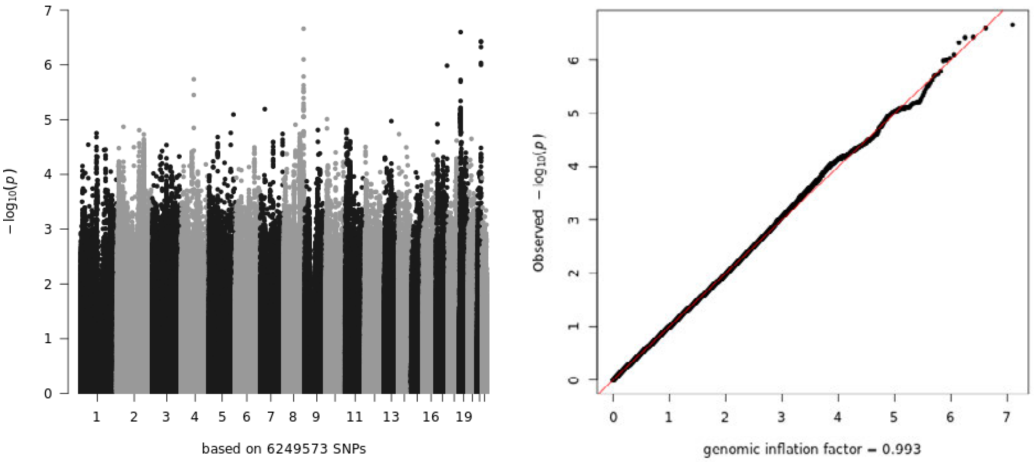


linear
age-dependent

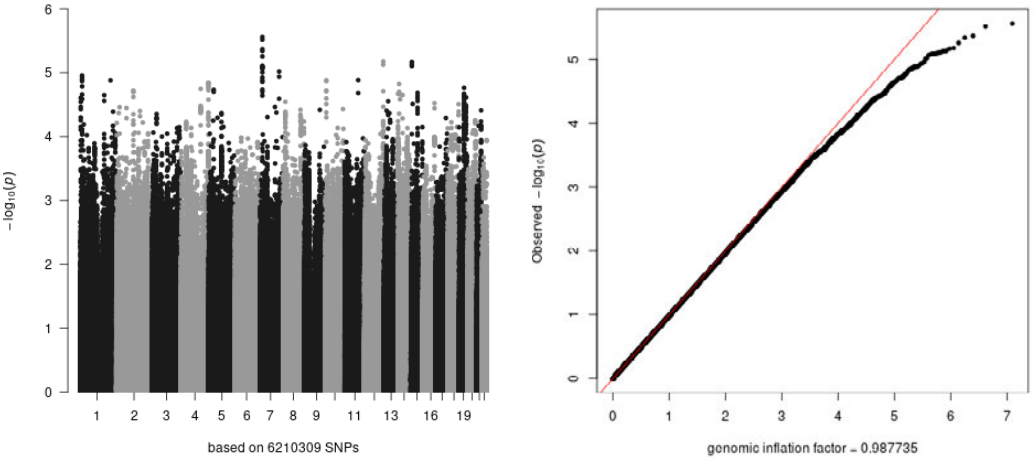


quadratic
age-dependent

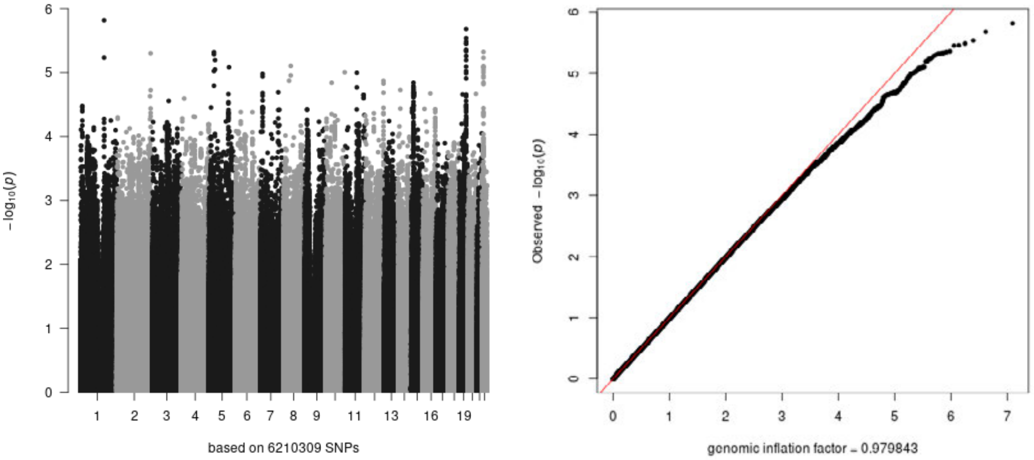

### 3K: change rate pallidum volume

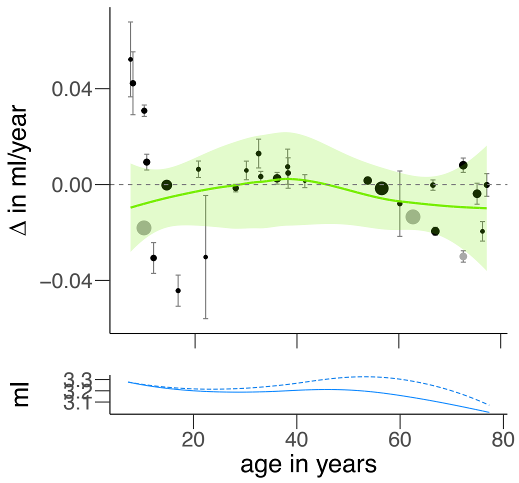


age
independent

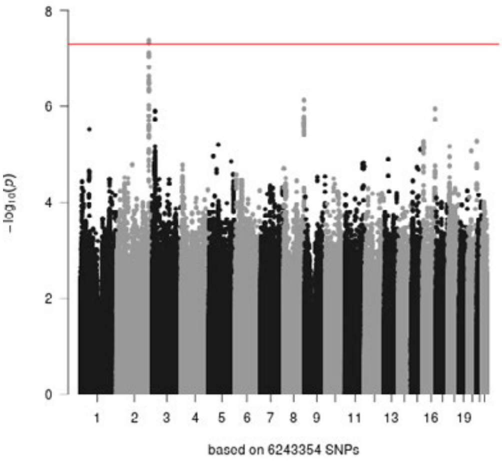


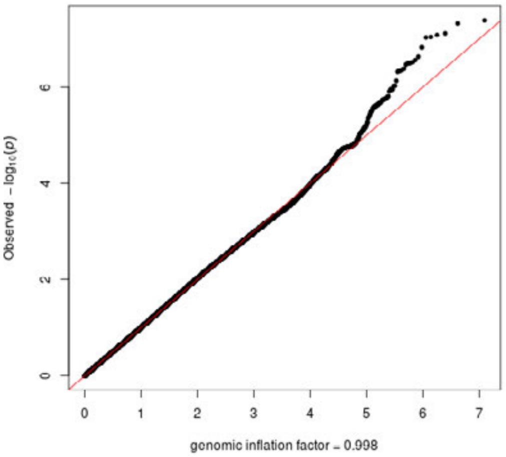


linear
age-dependent

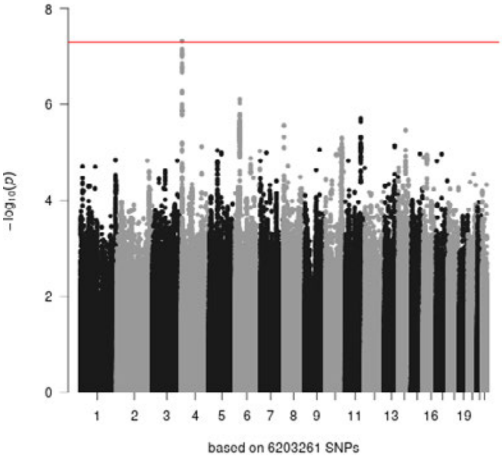


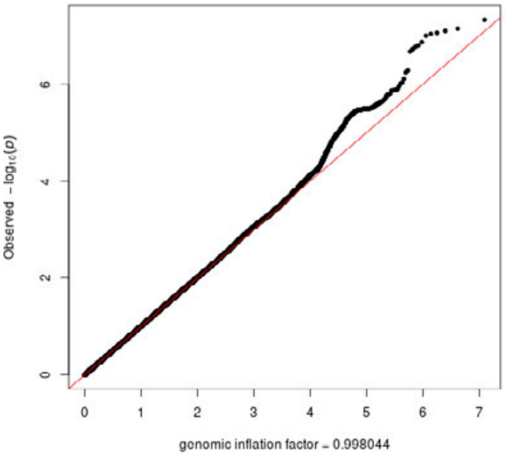


quadratic
age-dependent

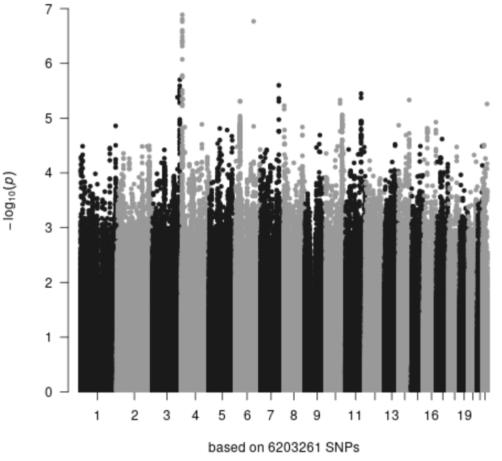


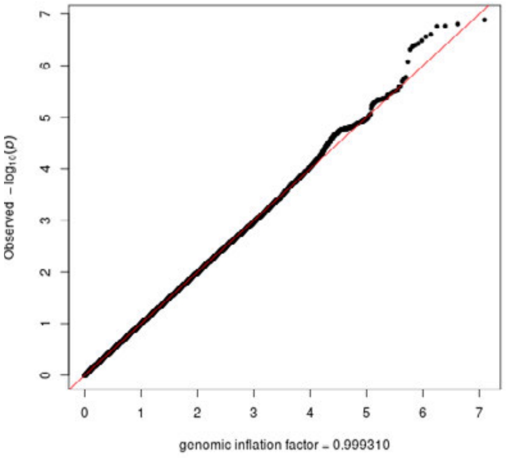

### 3L: change rate putamen volume

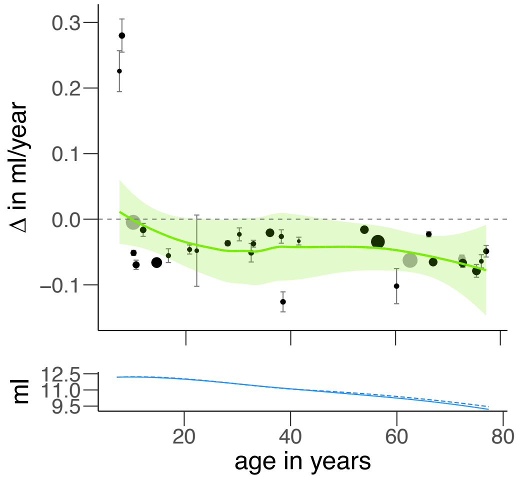


age
independent

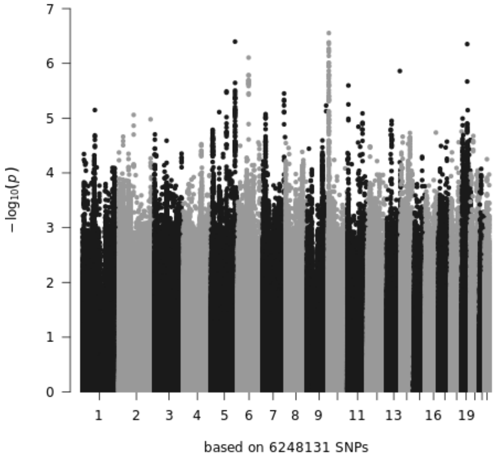


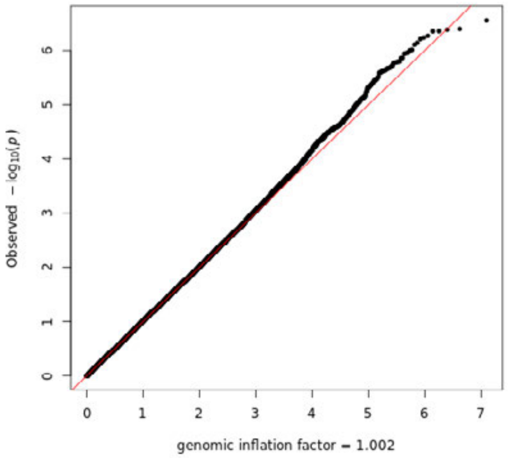


linear
age-dependent

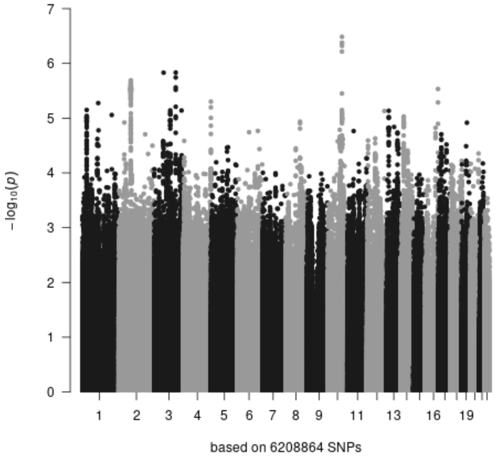


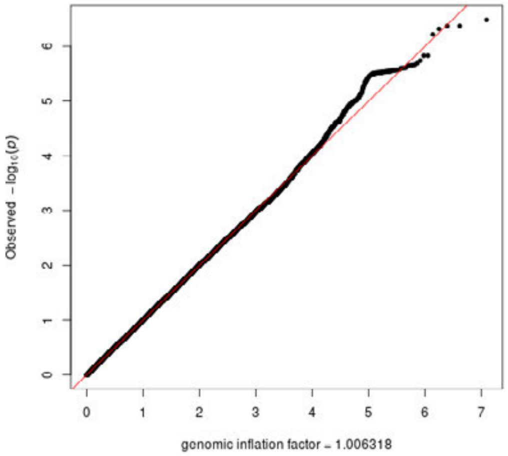


quadratic
age-dependent

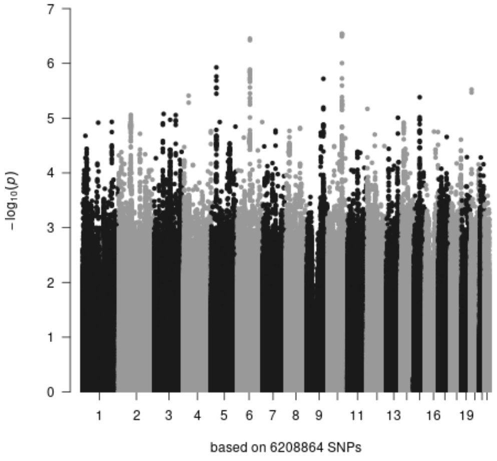


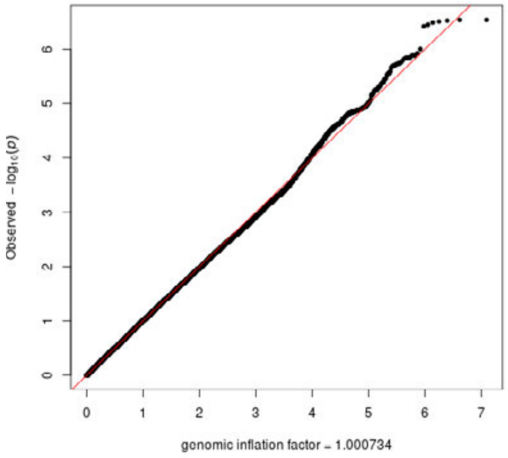

### 3M: change rate surface area

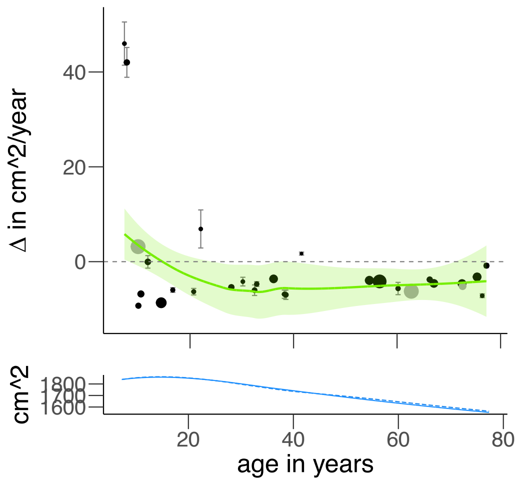


age
independent

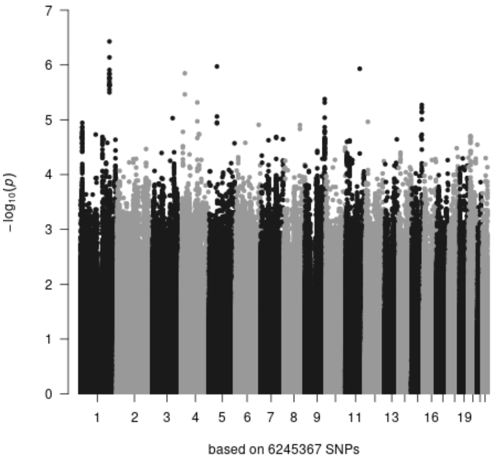


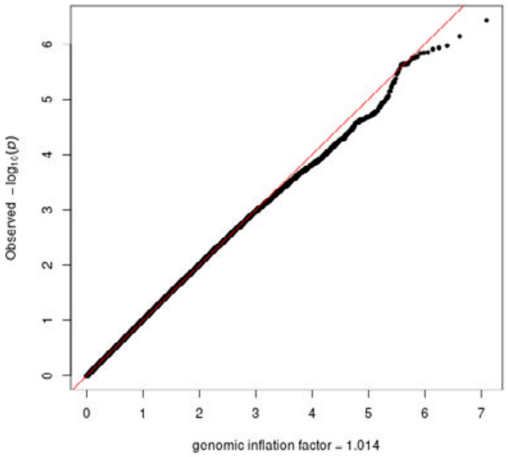


linear
age-dependent

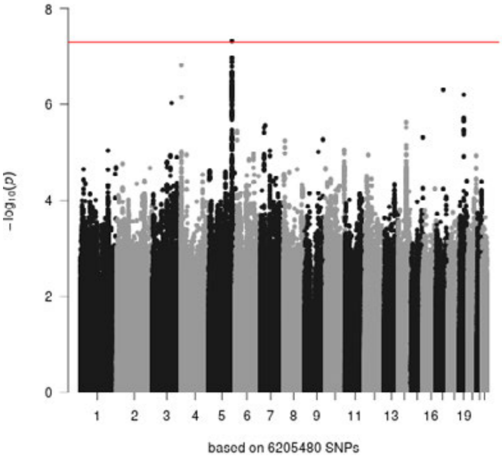


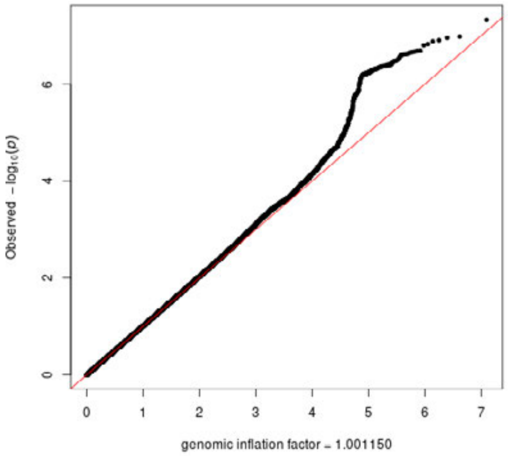


quadratic
age-dependent

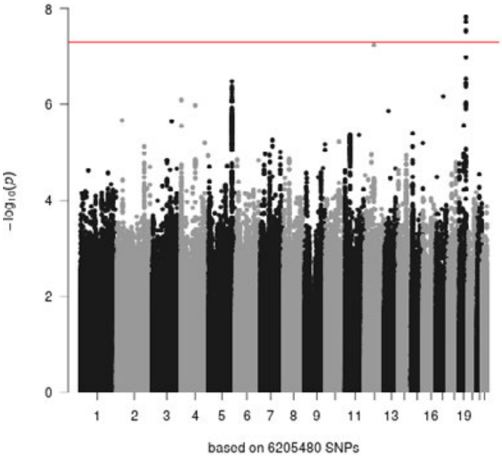


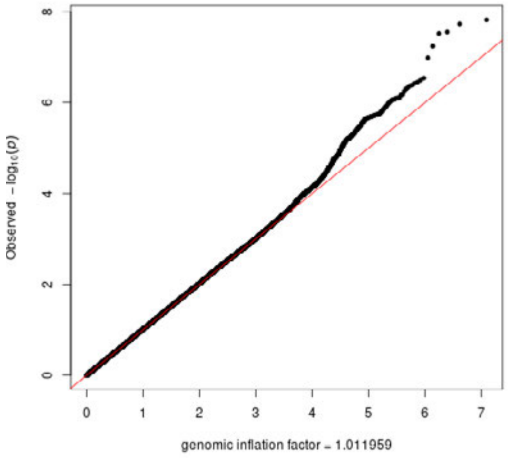

### 3N: change rate thalamus volume

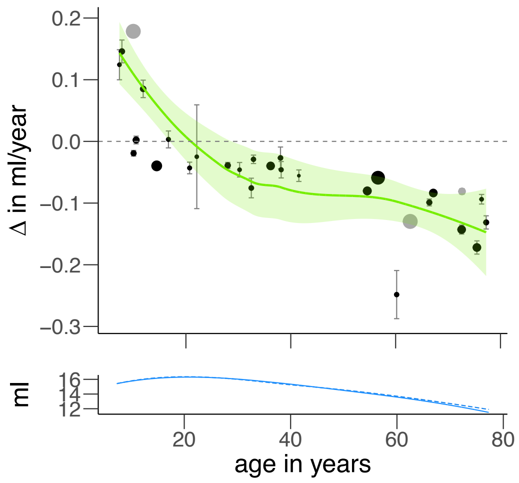


age independent

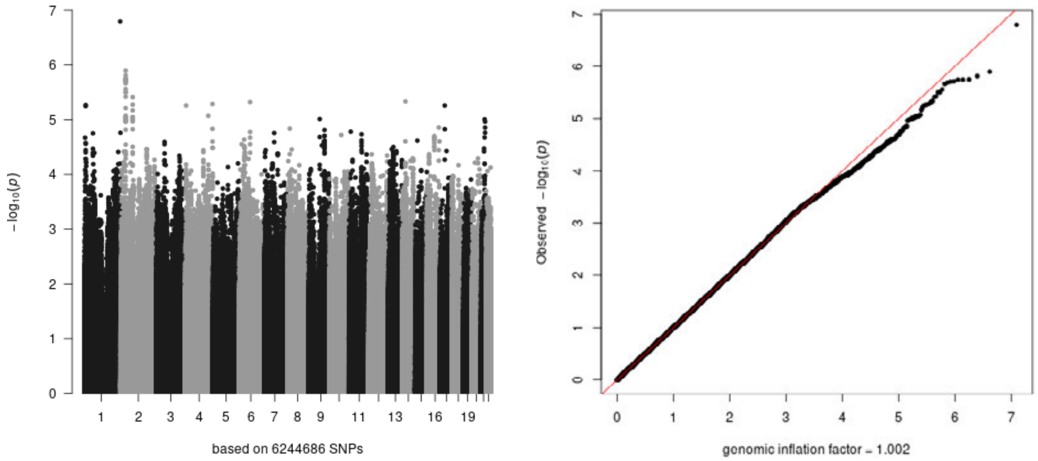


linear age-dependent

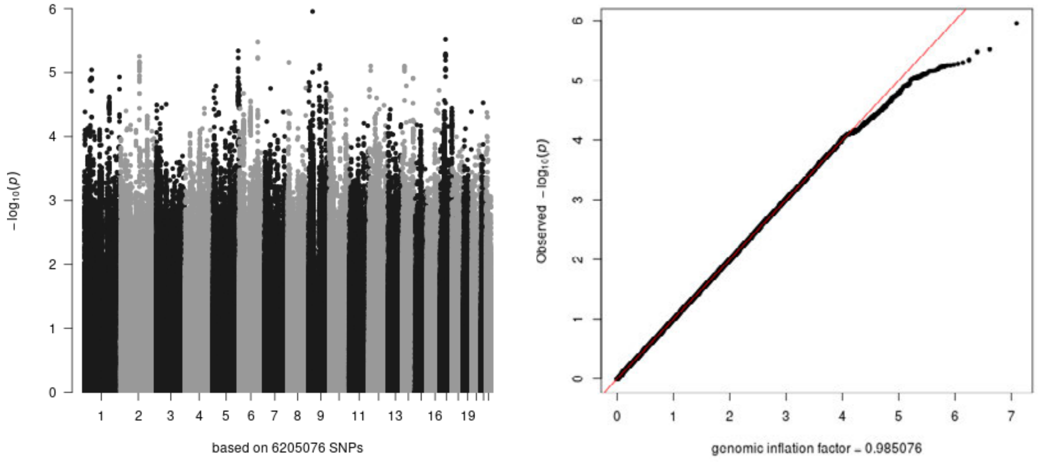


quadratic age-dependent

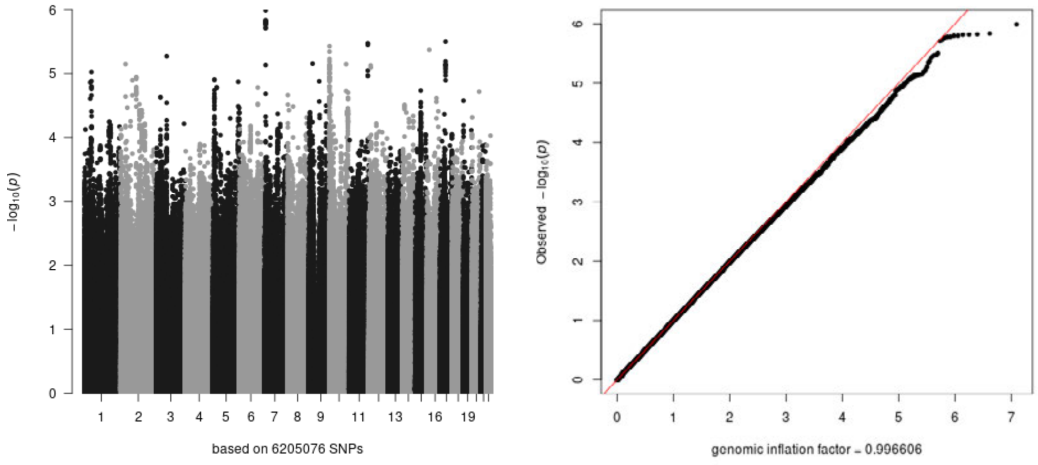

### 3O: change rate total brain volume

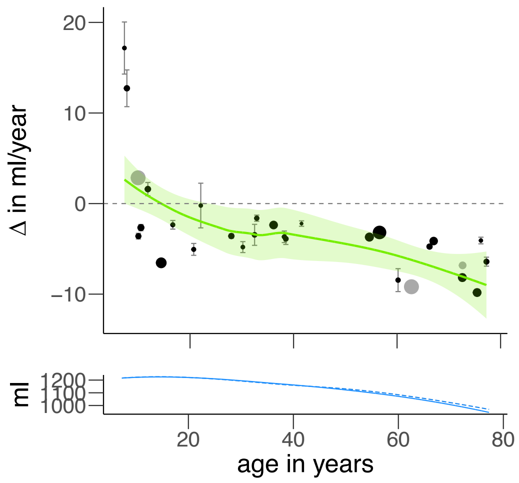


age
independent

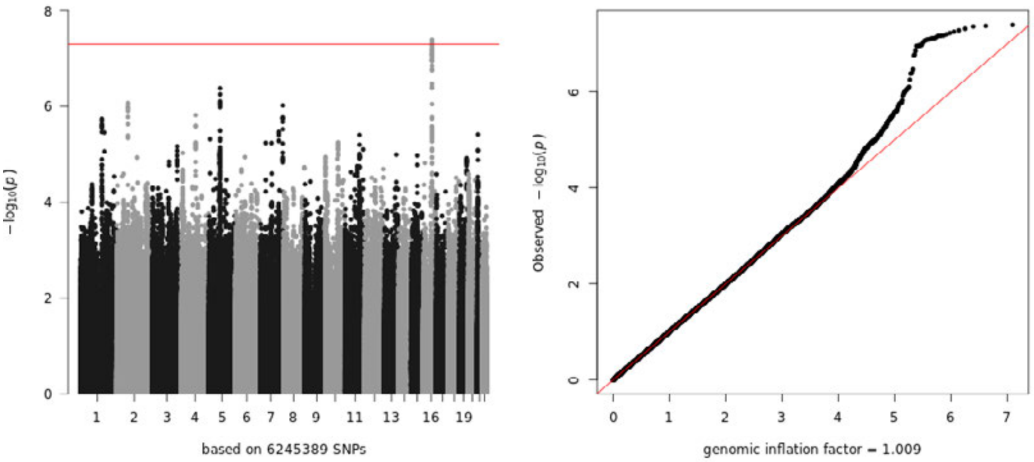


linear
age-dependent

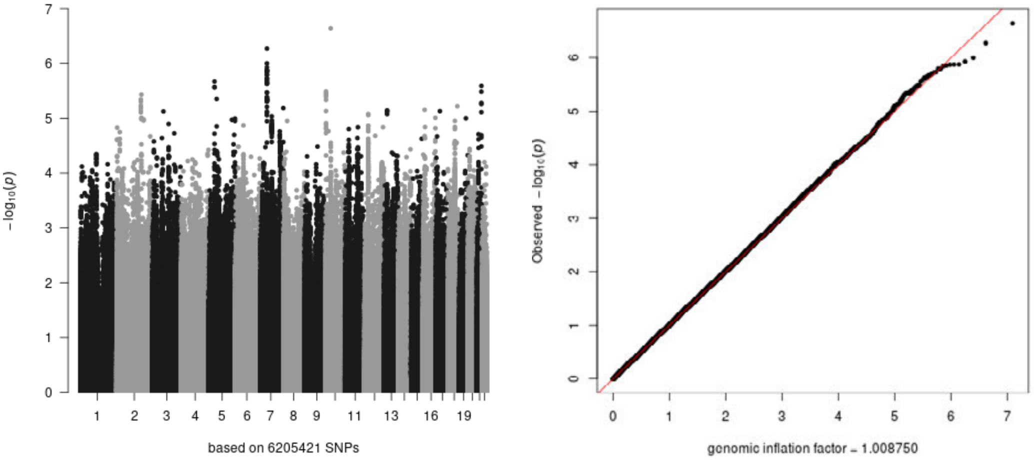


quadratic
age-dependent

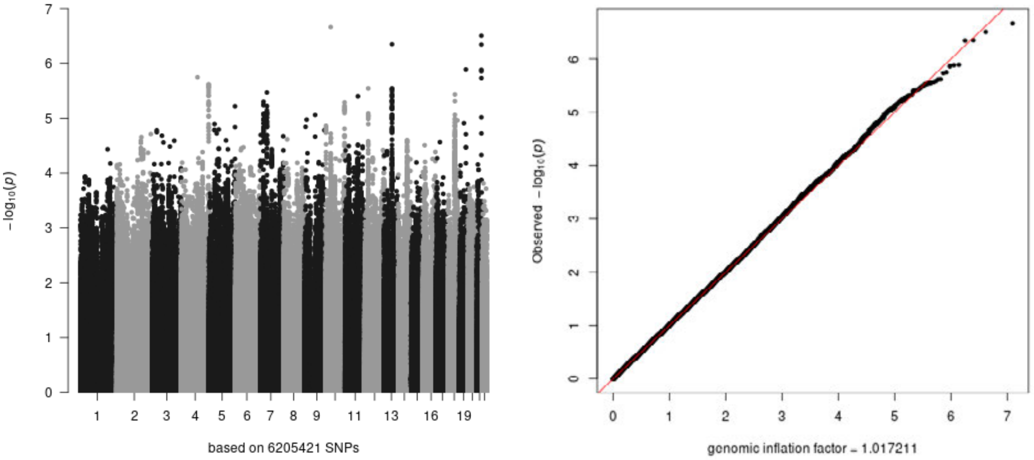

4A

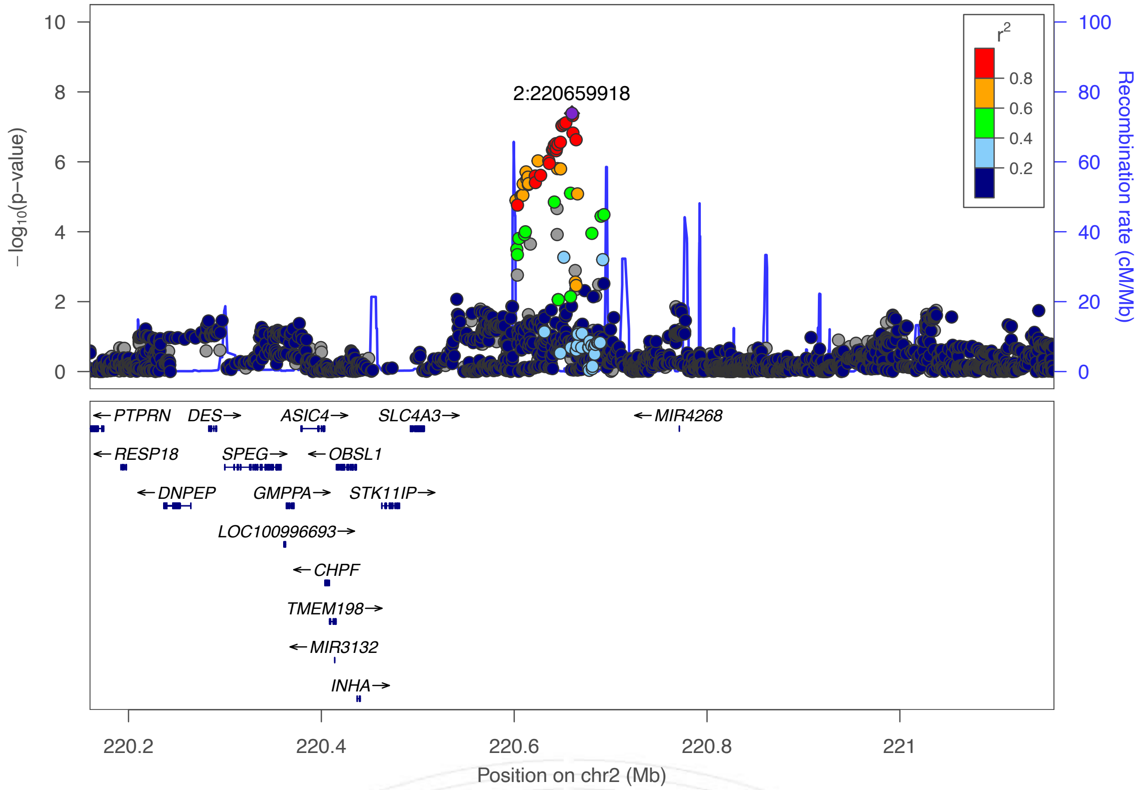
2:220659918
−log10(p−value)
Recombination rate (cM/Mb)
r2
0.8
0.6
0.4
0.2
PTPRN
DES
ASIC4
SLC4A3
MIR4268
RESP18
SPEG
OBSL1
DNPEP
GMPPA
STK11IP
LOC100996693
CHPF
TMEM198
MIR3132
INHA
220.2
220.4
220.6
220.8
221
Position on chr2 (Mb)

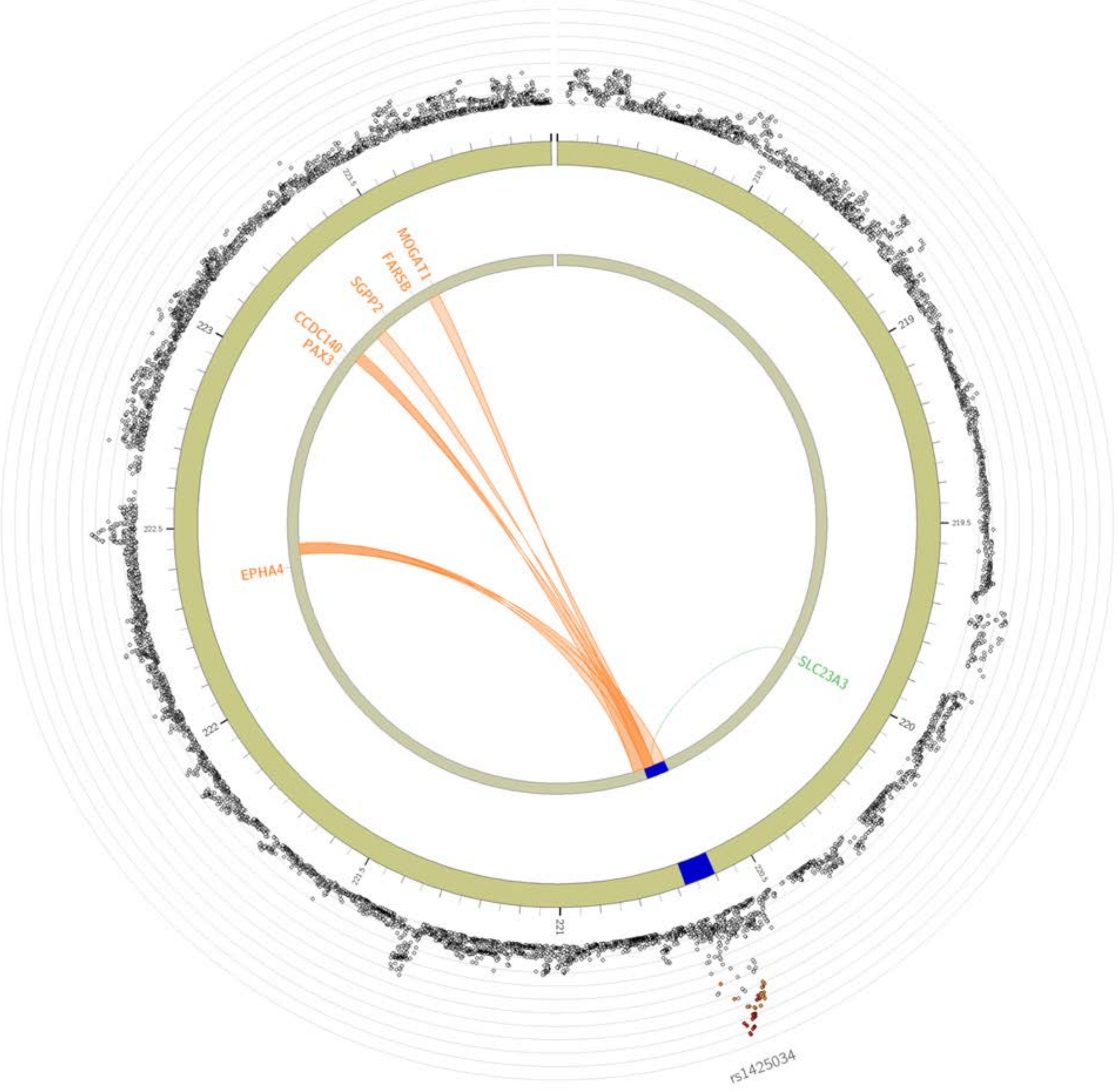
MOGAT1
FARSB
SGPP2
CCDC140
PAX3
EPHA4
SLC23A3
rs1425034

4B

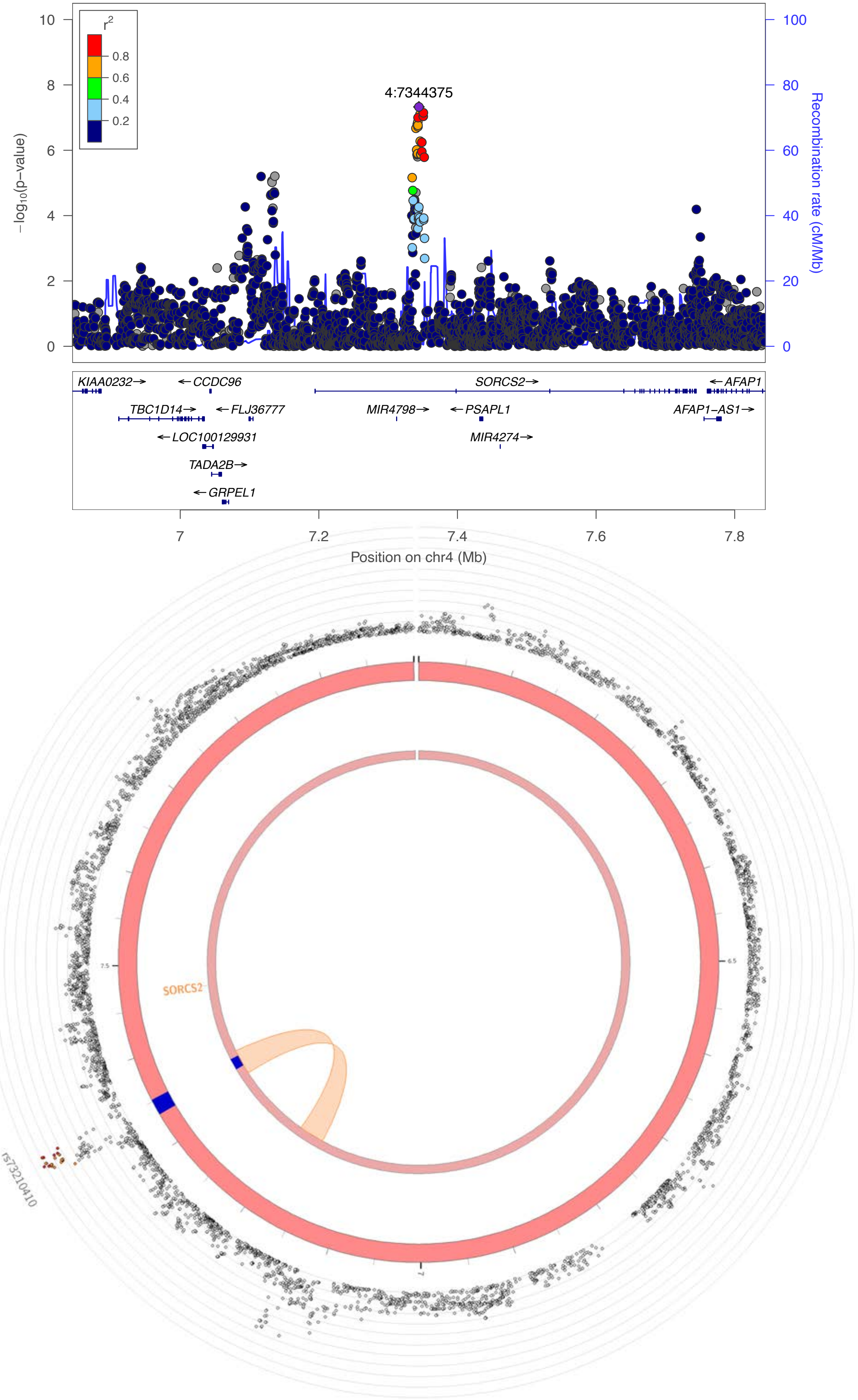

4:7344375
r2
0.8
0.6
0.4
0.2
-log10(p-value)
Recombination rate (cM/Mb)
10
8
6
4
2
0
100
80
60
40
20
0
KIAA0232→
←CCDC96
SORCS2→
←AFAP1
TBC1D14→
←FLJ36777
MIR4798→
←PSAPL1
AFAP1-AS1→
←LOC100129931
MIR4274→
TADA2B→
←GRPEL1
7
7.2
7.4
7.6
7.8
Position on chr4 (Mb)
SORCS2
rs73210410

4C

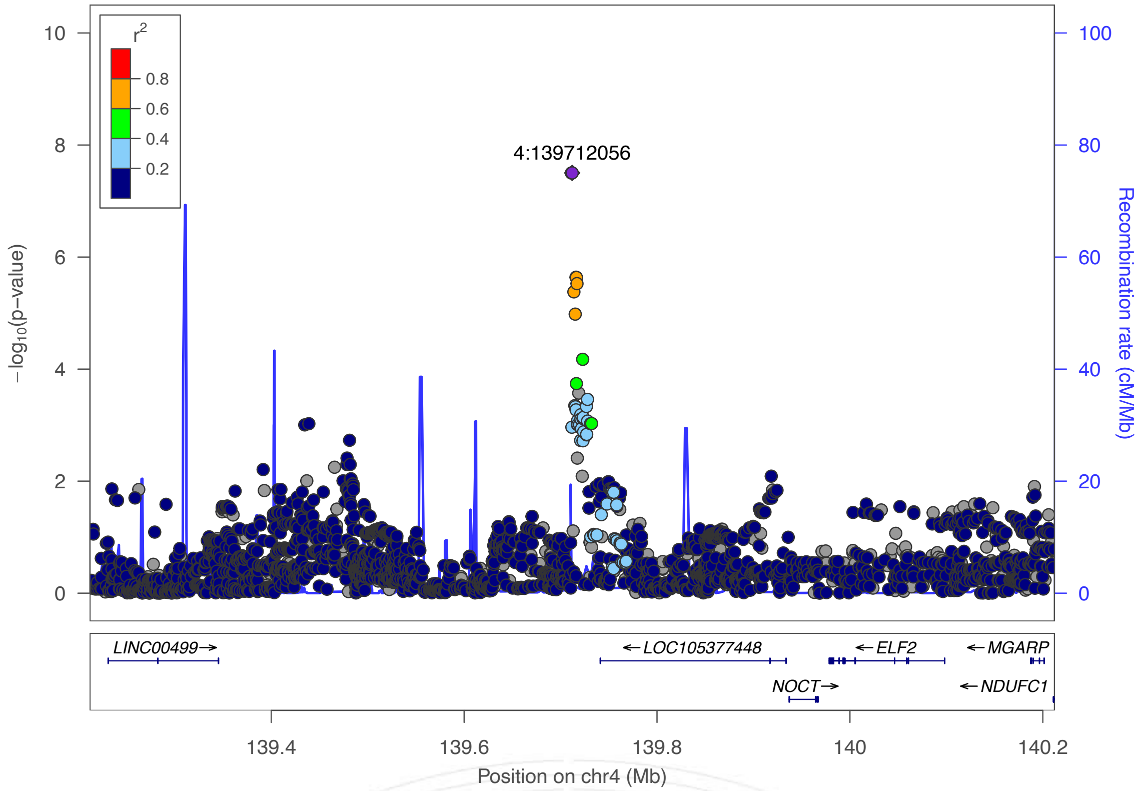

r2
0.8
0.6
0.4
0.2
4:139712056
−log10(p−value)
Recombination rate (cM/Mb)
LINC00499→
←LOC105377448
←ELF2
←MGARP
NOCT→
←NDUFC1
139.4
139.6
139.8
140
140.2
Position on chr4 (Mb)


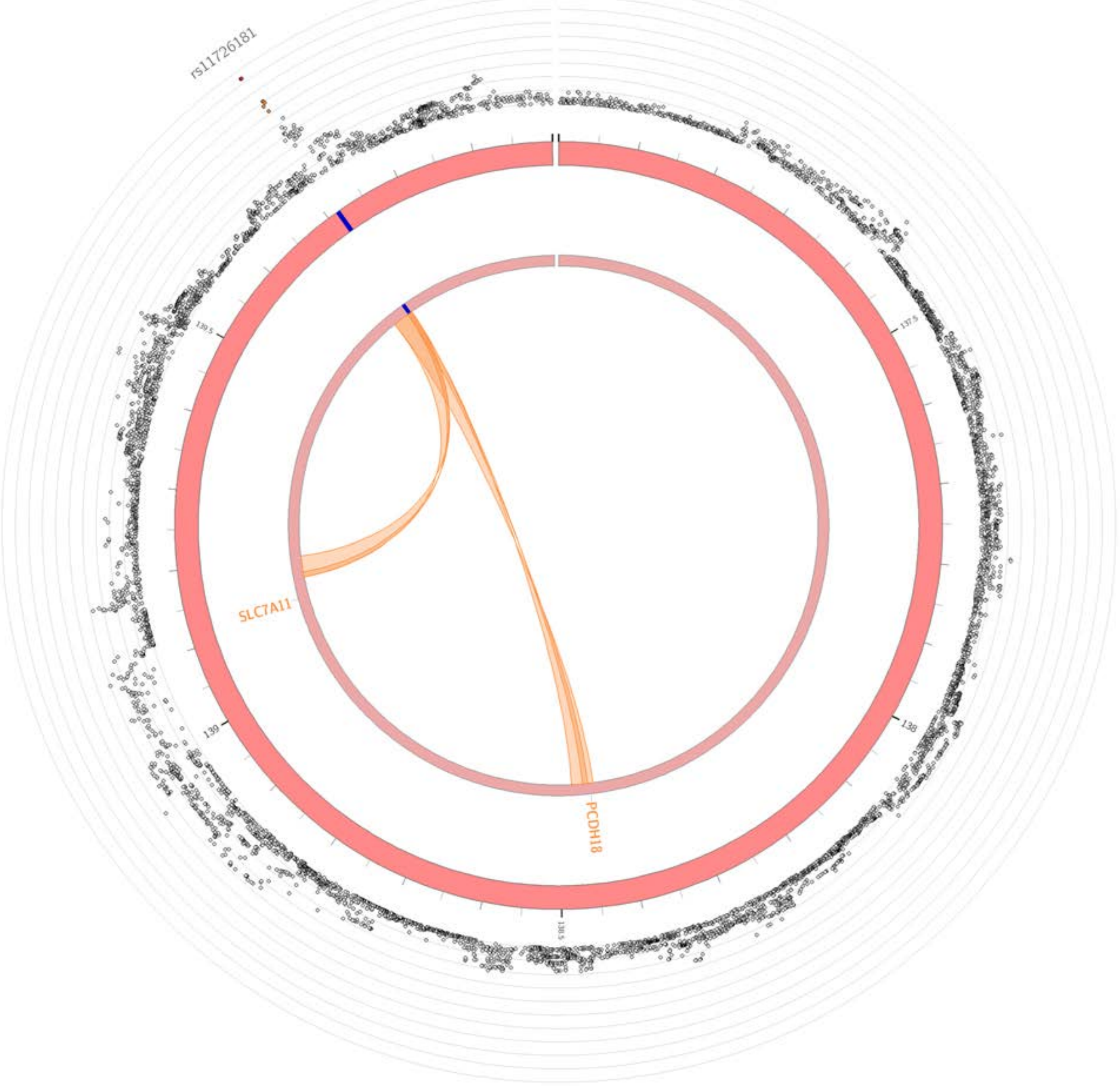

rs11726181
SLC7A11
PCDH18

4D

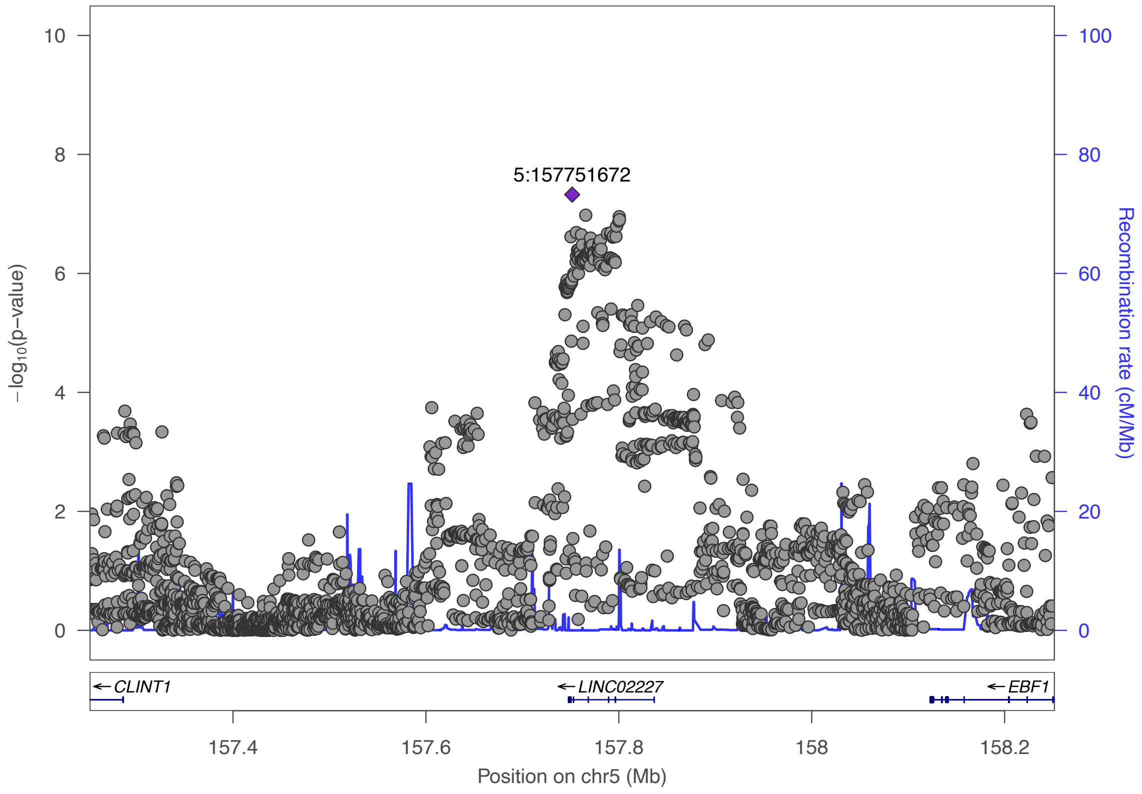

5:157751672
−log10(p−value)
Recombination rate (cM/Mb)
0
2
4
6
8
10
0
20
40
60
80
100
CLINT1
LINC02227
EBF1
157.4
157.6
157.8
158
158.2
Position on chr5 (Mb)

4E

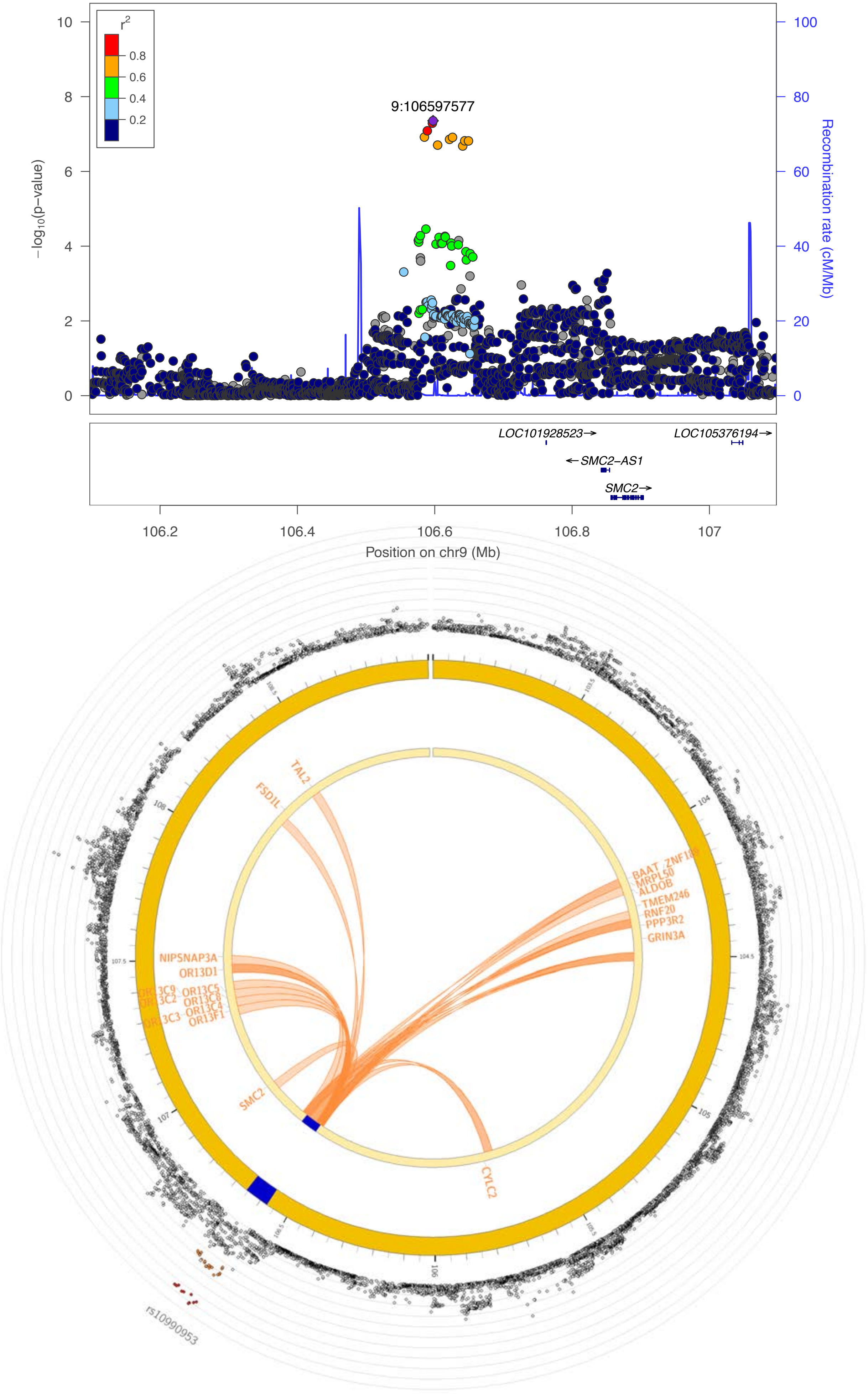

r2
0.8
0.6
0.4
0.2
9:106597577
−log10(p−value)
Recombination rate (cM/Mb)
LOC101928523→
LOC105376194→
←SMC2−AS1
SMC2→
106.2
106.4
106.6
106.8
107
Position on chr9 (Mb)
TAL2
FSD1L
BAAT
ZNF189
MRPL50
ALDOB
TMEM246
RNF20
PPP3R2
GRIN3A
NIPSNAP3A
OR13D1
OR13C9
OR13C2
OR13C5
OR13C8
OR13C4
OR13C3
OR13F1
SMC2
CYLC2
rs10990953

4F

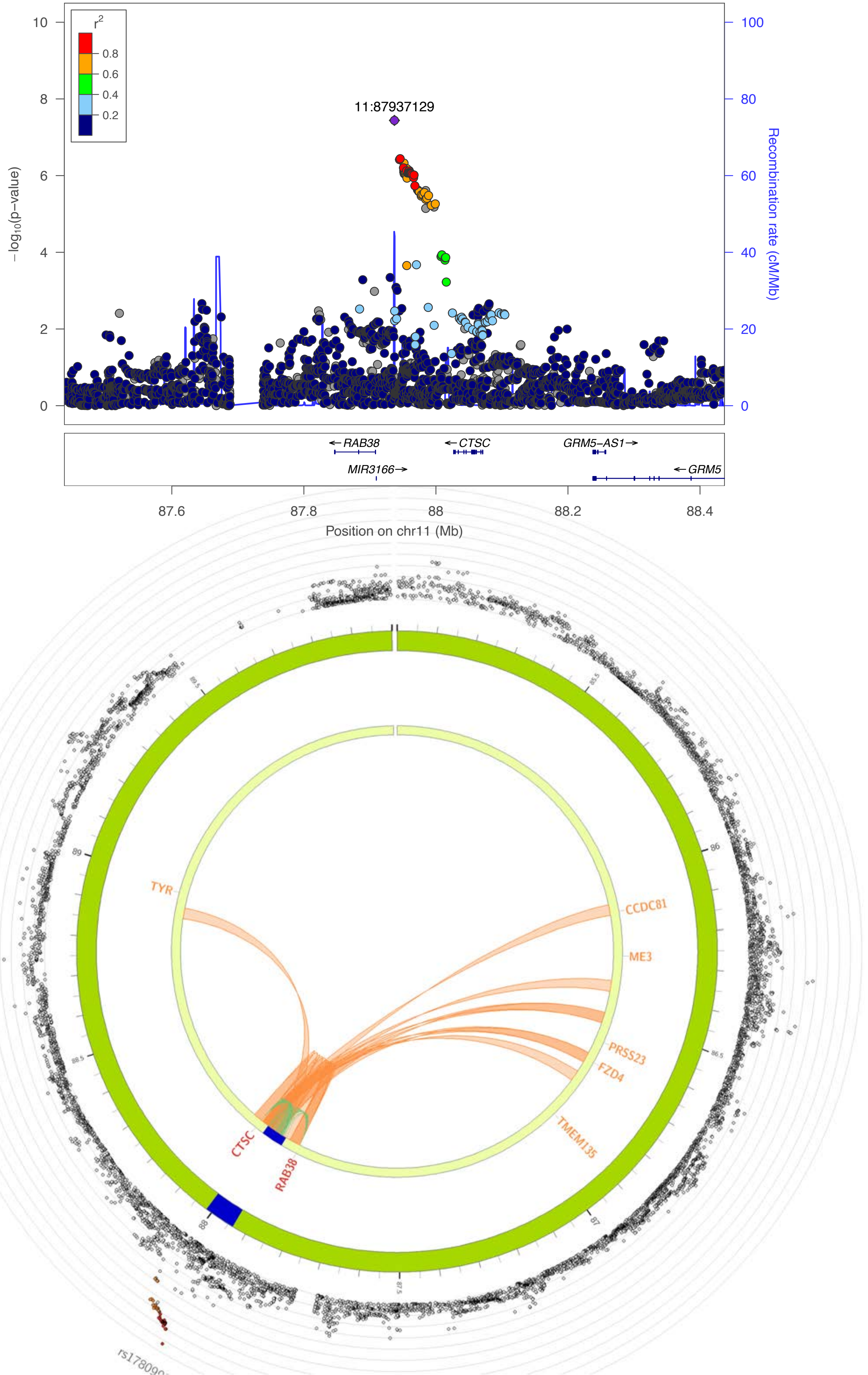

r2
0.8
0.6
0.4
0.2
11:87937129
−log10(p−value)
Recombination rate (cM/Mb)
10
8
6
4
2
0
100
80
60
40
20
0
RAB38
CTSC
GRM5−AS1
MIR3166
GRM5
87.6
87.8
88
88.2
88.4
Position on chr11 (Mb)
TYR
CCDC81
ME3
PRSS23
FZD4
TMEM135
CTSC
RAB38
85.5
86
86.5
87
87.5
88
88.5
89
89.5

4G

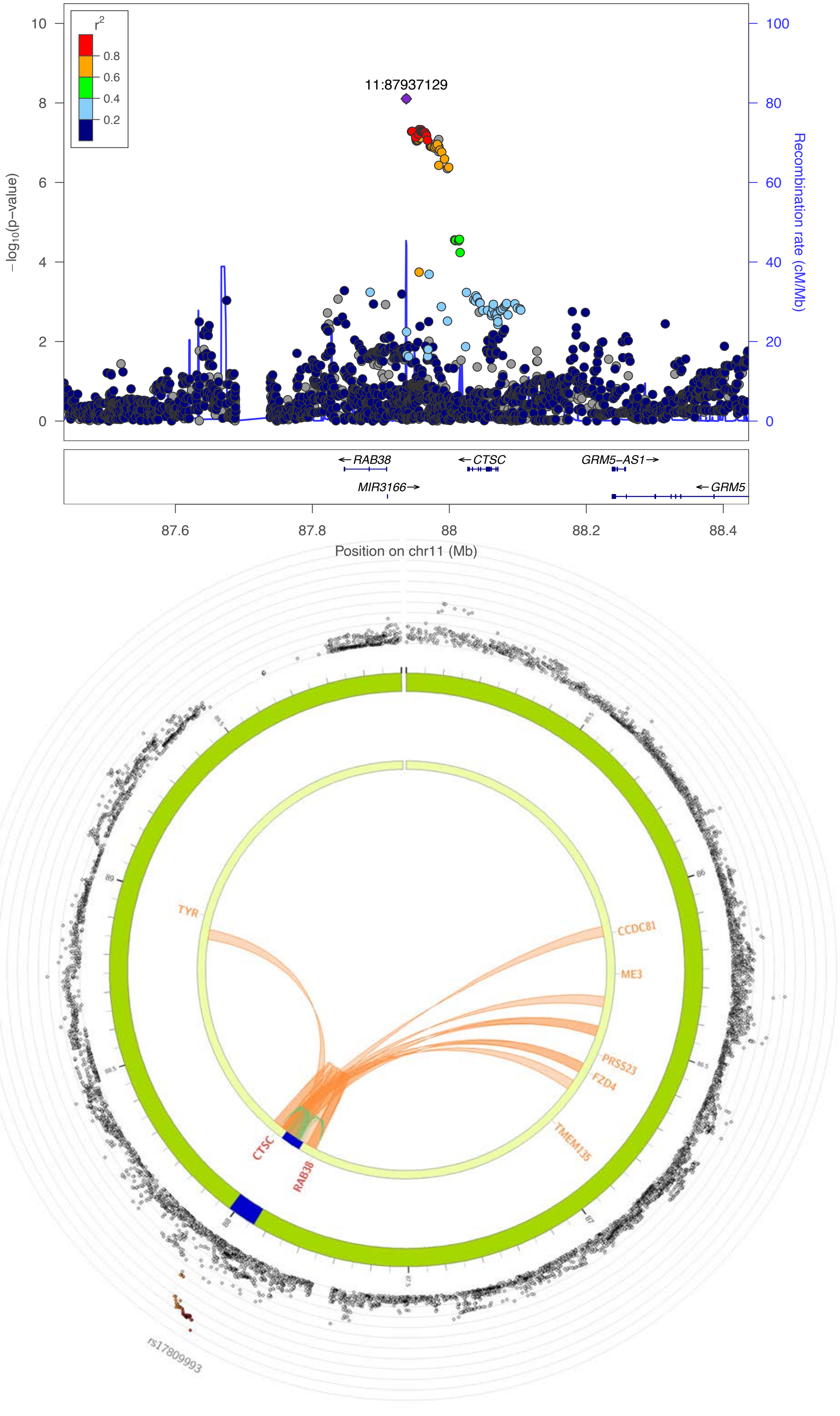
r2
0.8
0.6
0.4
0.2
11:87937129
−log10(p−value)
Recombination rate (cM/Mb)
RAB38
CTSC
GRM5−AS1
MIR3166
GRM5
Position on chr11 (Mb)
TYR
CCDC81
ME3
PRSS23
FZD4
TMEM135
CTSC
RAB38
rs17809993

4H

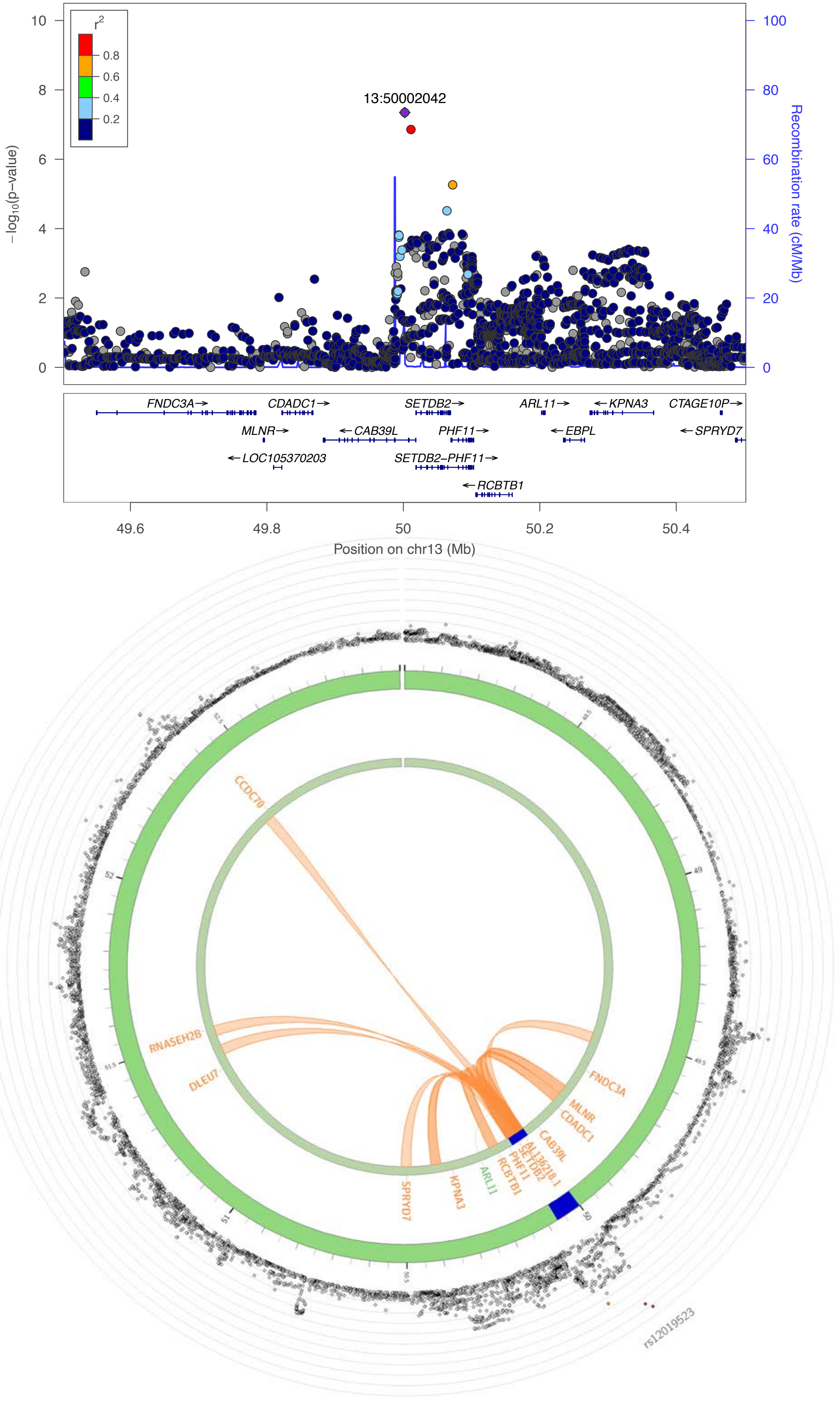
r2
0.8
0.6
0.4
0.2
13:50002042
−log10(p−value)
Recombination rate (cM/Mb)
FNDC3A→
CDADC1→
SETDB2→
ARL11→
←KPNA3
CTAGE10P→
MLNR→
←CAB39L
PHF11→
←EBPL
←SPRYD7
←LOC105370203
SETDB2−PHF11→
←RCBTB1
49.6
49.8
50
50.2
50.4
Position on chr13 (Mb)
CCDC70
RNASEH2B
DLEU7
FNDC3A
MLNR
CDADC1
CAB39L
AL136218.1
SETDB2
PHF11
RCBTB1
ARL11
KPNA3
SPRYD7
rs12019523

4I

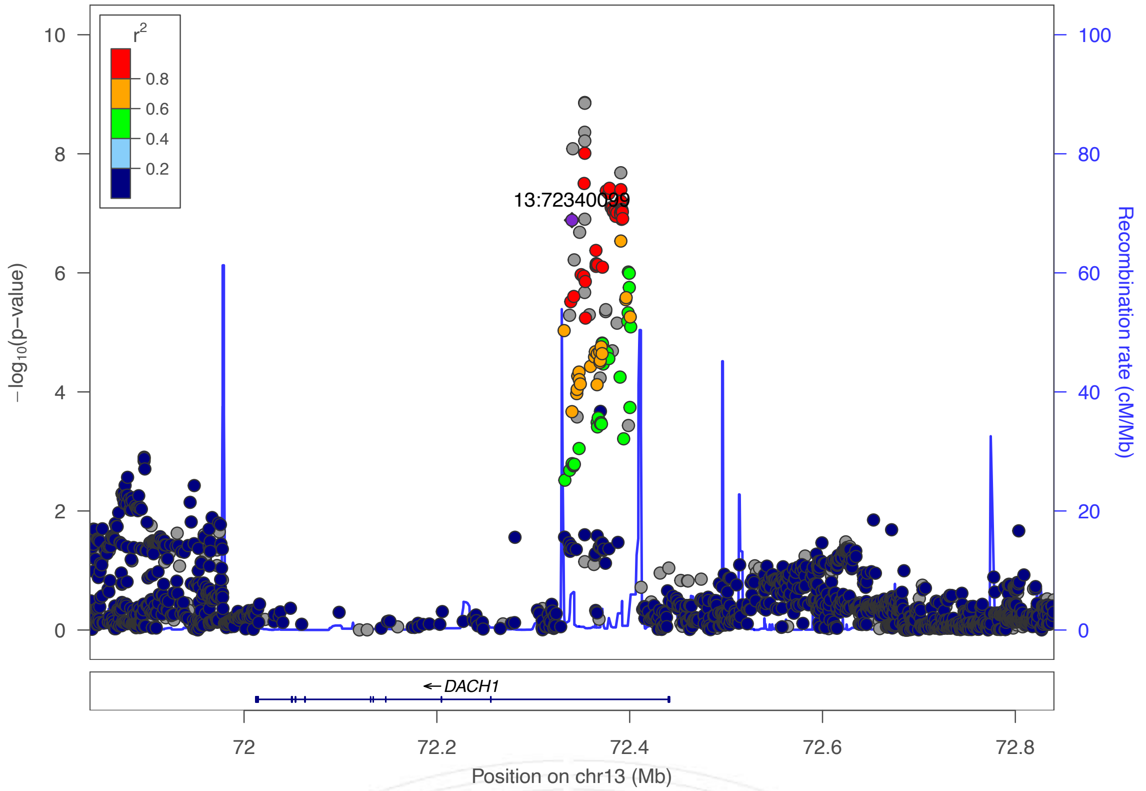

r2
0.8
0.6
0.4
0.2
13:72340099
−log10(p−value)
Recombination rate (cM/Mb)
DACH1
72
72.2
72.4
72.6
72.8
Position on chr13 (Mb)


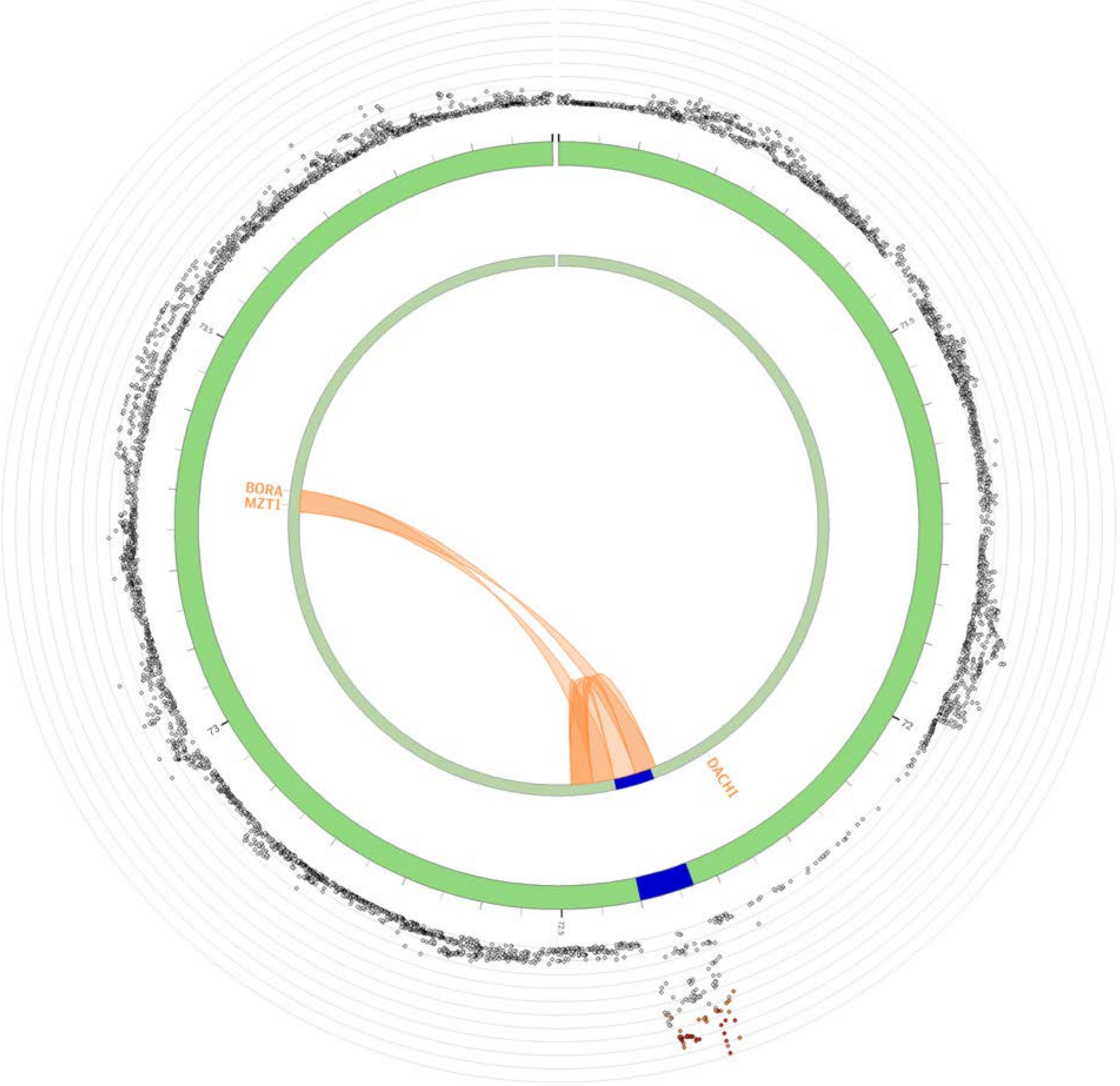

BORA
MZT1
DACH1

4J

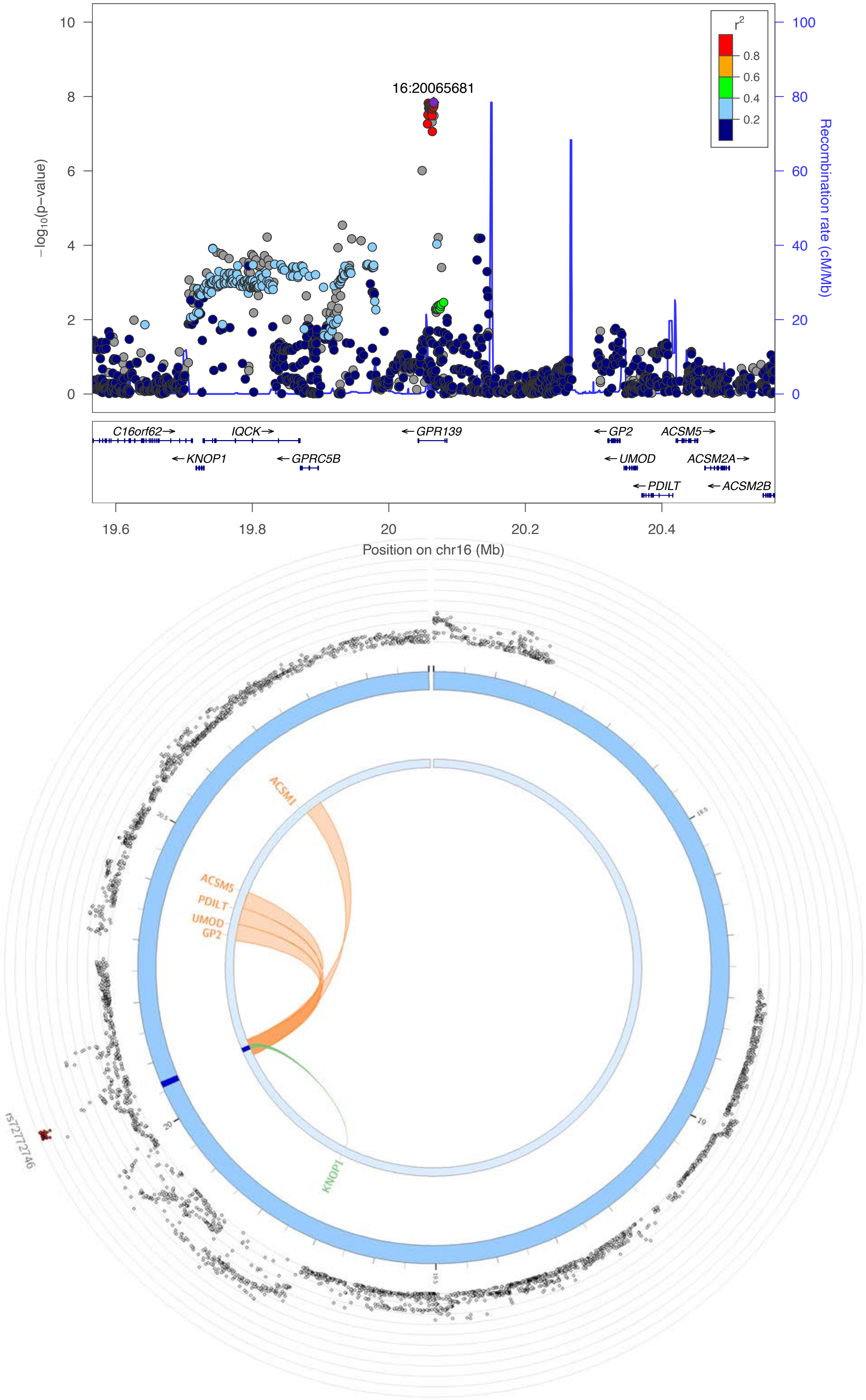

16:20065681
r2
0.8
0.6
0.4
0.2
-log10(p-value)
Recombination rate (cM/Mb)
0
2
4
6
8
10
20
40
60
80
100
C16orf62
IQCK
GPR139
GP2
ACSM5
KNOP1
GPRC5B
UMOD
ACSM2A
PDILT
ACSM2B
19.6
19.8
20
20.2
20.4
Position on chr16 (Mb)
ACSM1
ACSM5
PDILT
UMOD
GP2
KNOP1
rs72772746

4K

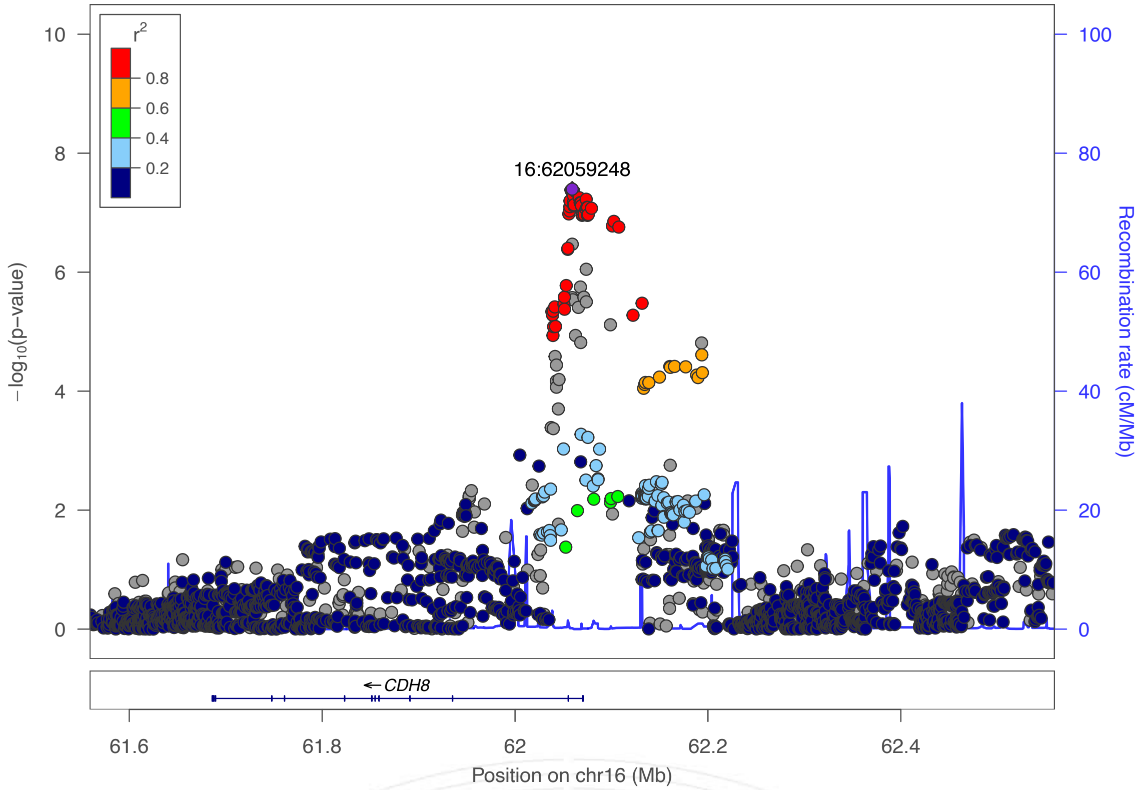

r2
0.8
0.6
0.4
0.2
10
8
6
4
2
0
−log10(p−value)
100
80
60
40
20
0
Recombination rate (cM/Mb)
16:62059248
← CDH8
61.6
61.8
62
62.2
62.4
Position on chr16 (Mb)


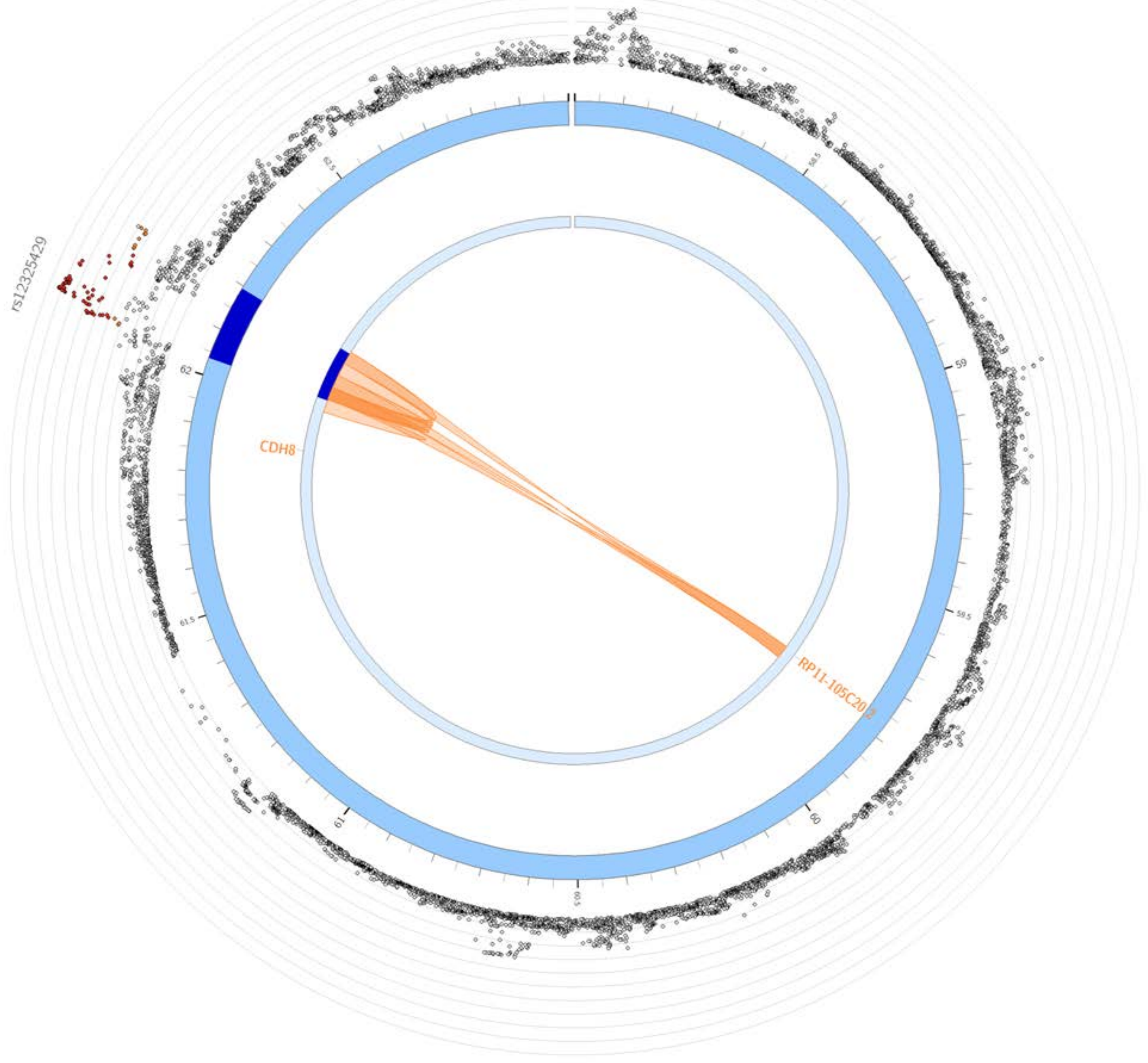

rs12325429
CDH8
RP11-105C20.2

4L

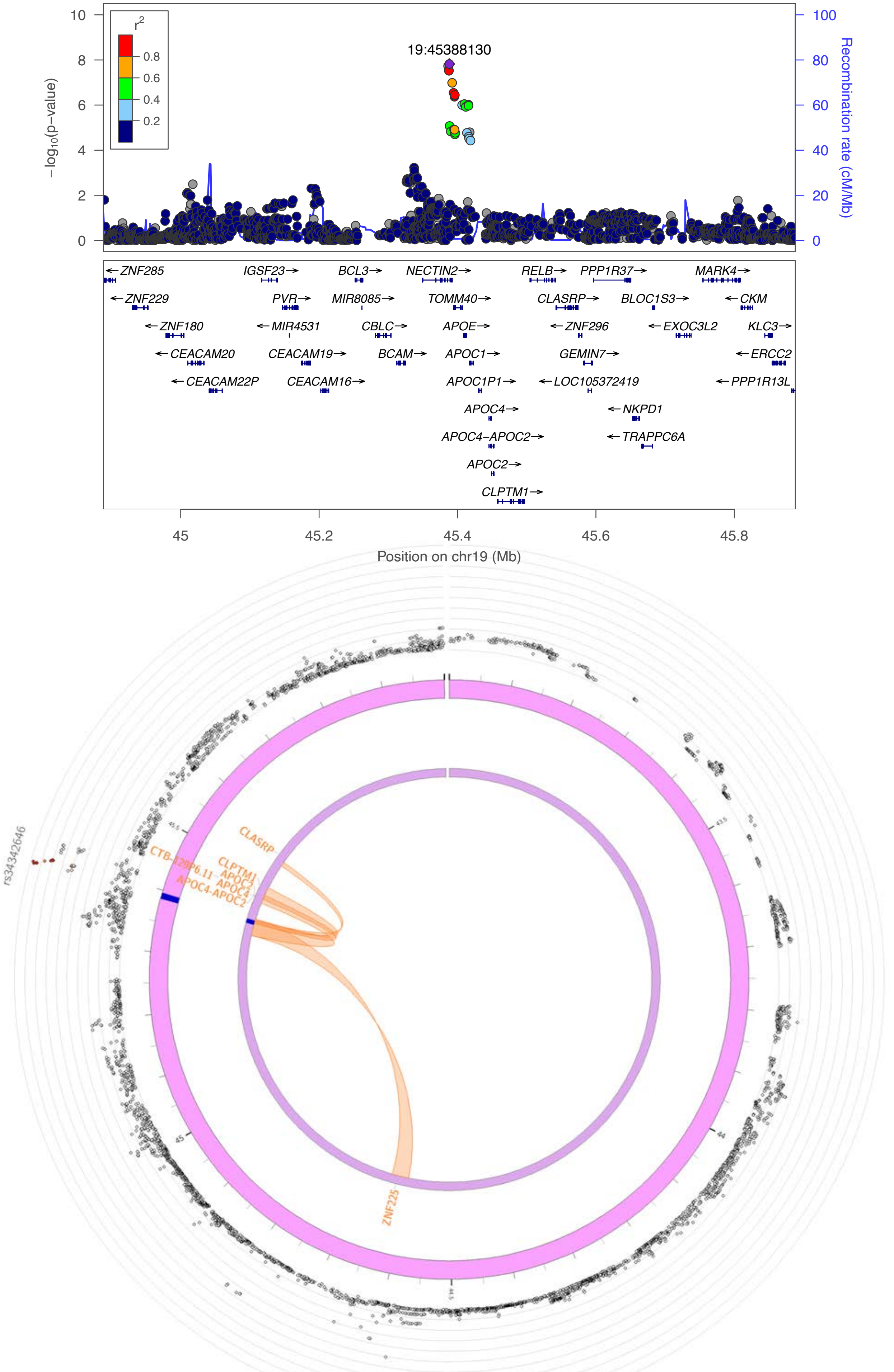

19:45388130
r2
0.8
0.6
0.4
0.2
-log10(p-value)
Recombination rate (cM/Mb)
ZNF285
ZNF229
ZNF180
CEACAM20
CEACAM22P
IGSF23
PVR
MIR4531
CEACAM19
CEACAM16
BCL3
MIR8085
CBLC
BCAM
NECTIN2
TOMM40
APOE
APOC1
APOC1P1
APOC4
APOC4-APOC2
APOC2
CLPTM1
RELB
CLASRP
ZNF296
GEMIN7
LOC105372419
PPP1R37
BLOC1S3
EXOC3L2
NKPD1
TRAPPC6A
MARK4
CKM
KLC3
ERCC2
PPP1R13L
Position on chr19 (Mb)
rs3434646
CLASRP
CLPTM1
APOC2
APOC4
APOC4-APOC2
CTB-129P6.11
ZNF225

4M

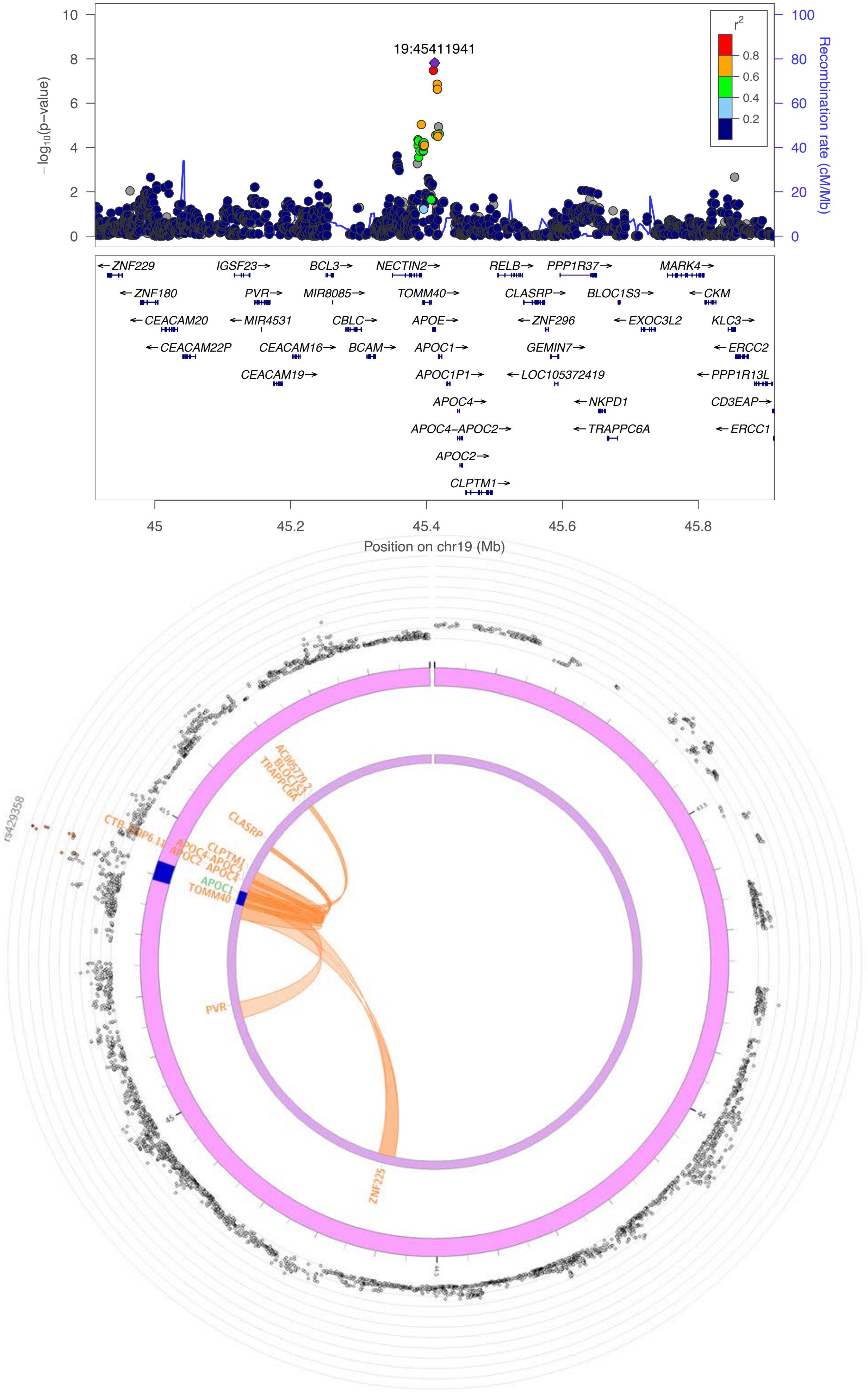
19:45411941
−log10(p−value)
Recombination rate (cM/Mb)
r2
0.8
0.6
0.4
0.2
ZNF229
IGSF23
BCL3
NECTIN2
RELB
PPP1R37
MARK4
ZNF180
PVR
MIR8085
TOMM40
CLASRP
BLOC1S3
CKM
CEACAM20
MIR4531
CBLC
APOE
ZNF296
EXOC3L2
KLC3
CEACAM22P
CEACAM16
BCAM
APOC1
GEMIN7
ERCC2
CEACAM19
APOC1P1
LOC105372419
PPP1R13L
APOC4
NKPD1
CD3EAP
APOC4−APOC2
TRAPPC6A
ERCC1
APOC2
CLPTM1
Position on chr19 (Mb)
rs429358
CLASRP
PVR
ZNF225
APOC1
TOMM40

5A

**change rate pal; effect of allele "a" in 2:220659918**

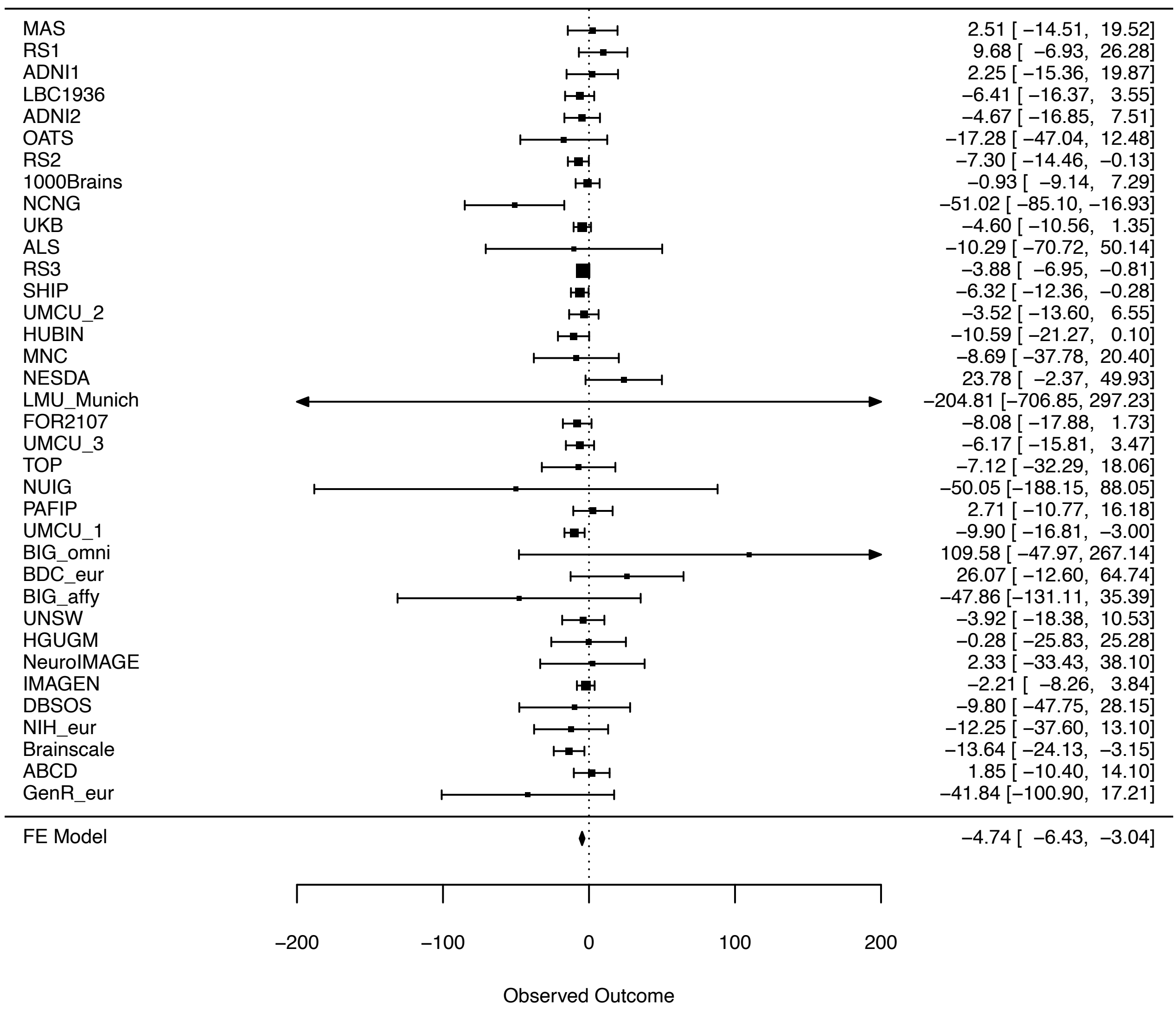


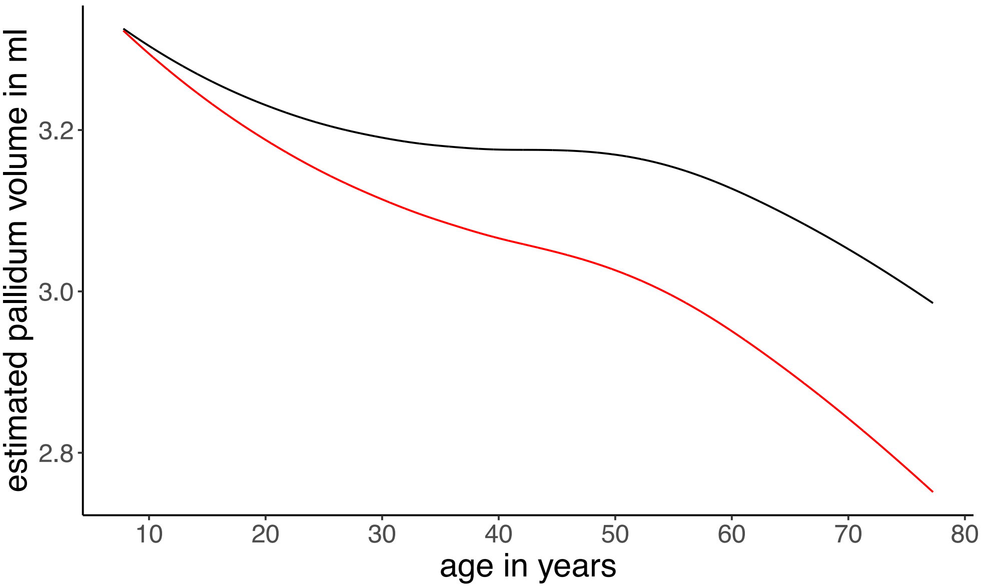

5B

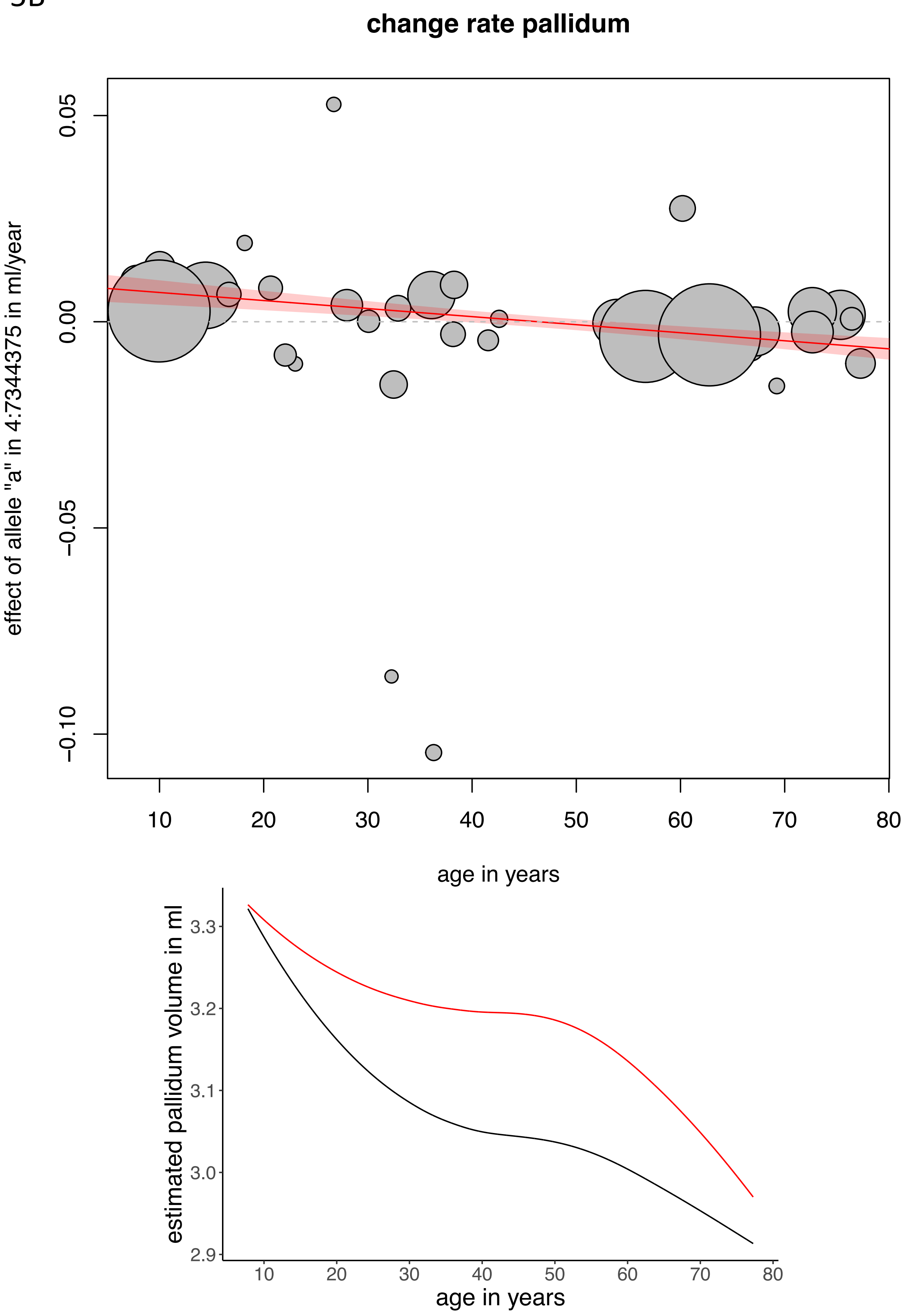
change rate pallidum
effect of allele "a" in 4:7344375 in ml/year
0.05
0.00
−0.05
−0.10
10
20
30
40
50
60
70
80
age in years
estimated pallidum volume in ml
3.3
3.2
3.1
3.0
2.9
10
20
30
40
50
60
70
80
age in years

5C

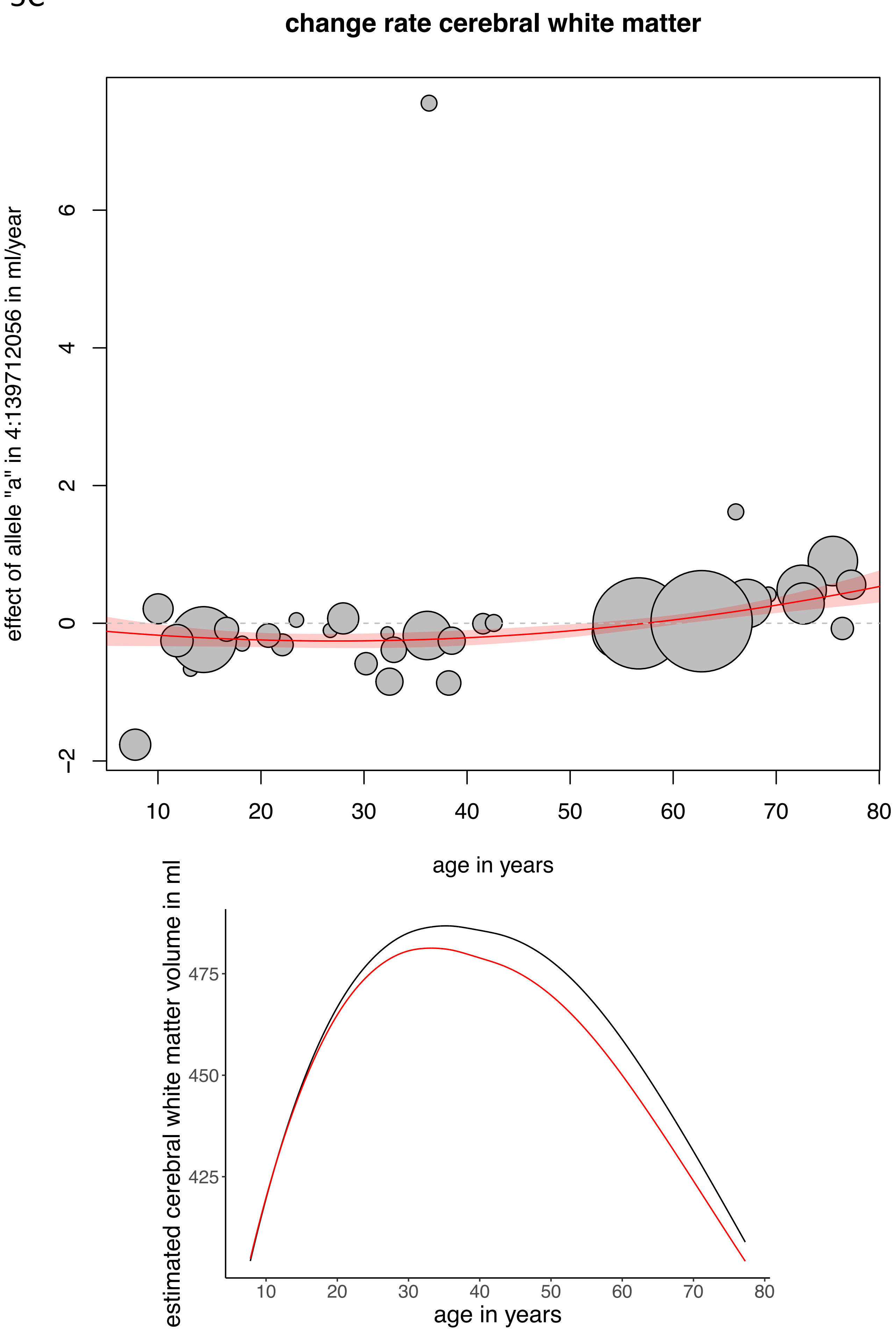

change rate cerebral white matter
effect of allele "a" in 4:139712056 in ml/year
−2
0
2
4
6
10
20
30
40
50
60
70
80
age in years
estimated cerebral white matter volume in ml
425
450
475
10
20
30
40
50
60
70
80
age in years

5D

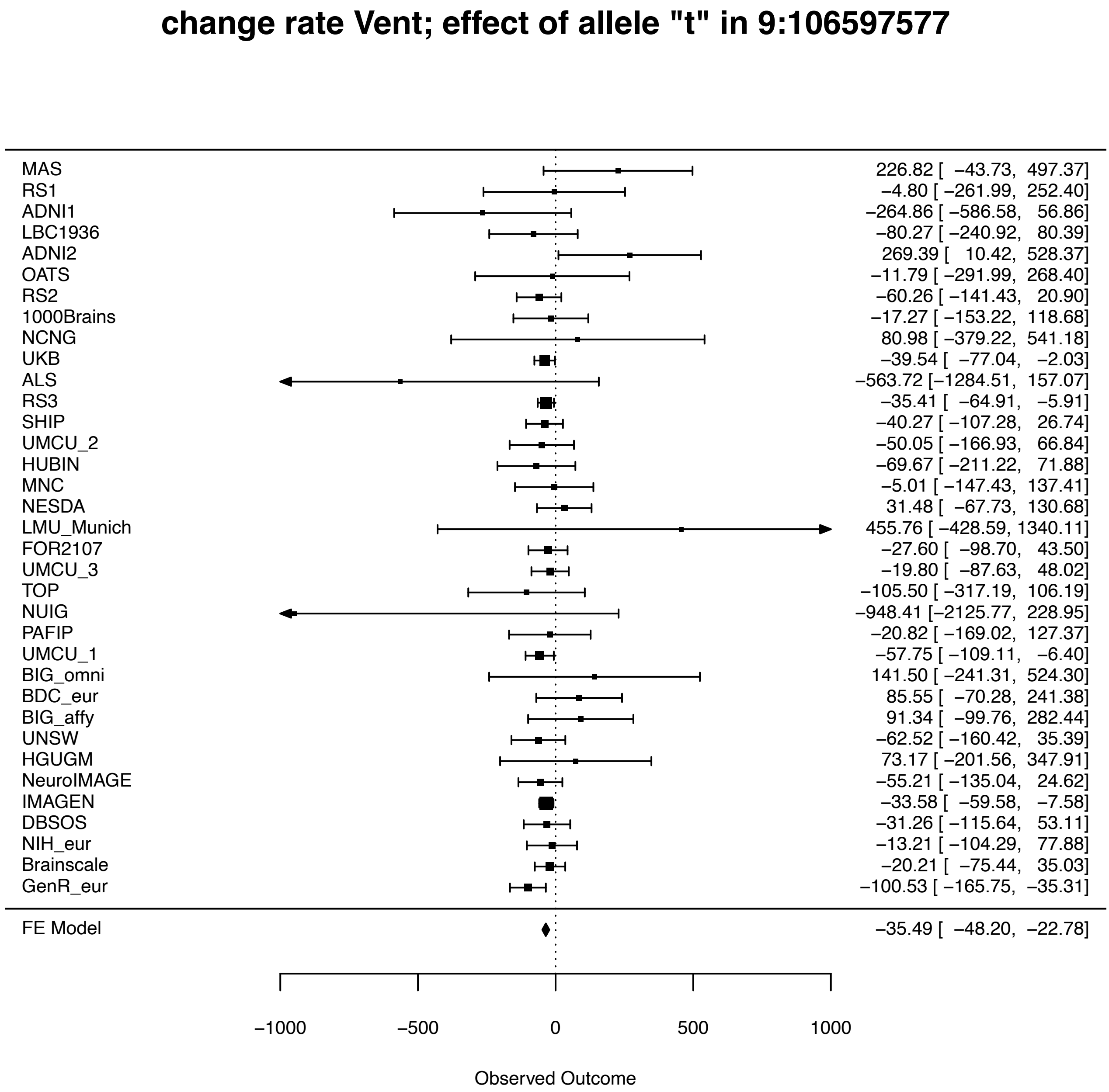
change rate Vent; effect of allele "t" in 9:106597577
MAS 226.82 [ −43.73, 497.37]
RS1 −4.80 [ −261.99, 252.40]
ADNI1 −264.86 [ −586.58, 56.86]
LBC1936 −80.27 [ −240.92, 80.39]
ADNI2 269.39 [ 10.42, 528.37]
OATS −11.79 [ −291.99, 268.40]
RS2 −60.26 [ −141.43, 20.90]
1000Brains −17.27 [ −153.22, 118.68]
NCNG 80.98 [ −379.22, 541.18]
UKB −39.54 [ −77.04, −2.03]
ALS −563.72 [−1284.51, 157.07]
RS3 −35.41 [ −64.91, −5.91]
SHIP −40.27 [ −107.28, 26.74]
UMCU_2 −50.05 [ −166.93, 66.84]
HUBIN −69.67 [ −211.22, 71.88]
MNC −5.01 [ −147.43, 137.41]
NESDA 31.48 [ −67.73, 130.68]
LMU_Munich 455.76 [ −428.59, 1340.11]
FOR2107 −27.60 [ −98.70, 43.50]
UMCU_3 −19.80 [ −87.63, 48.02]
TOP −105.50 [ −317.19, 106.19]
NUIG −948.41 [−2125.77, 228.95]
PAFIP −20.82 [ −169.02, 127.37]
UMCU_1 −57.75 [ −109.11, −6.40]
BIG_omni 141.50 [ −241.31, 524.30]
BDC_eur 85.55 [ −70.28, 241.38]
BIG_affy 91.34 [ −99.76, 282.44]
UNSW −62.52 [ −160.42, 35.39]
HGUGM 73.17 [ −201.56, 347.91]
NeuroIMAGE −55.21 [ −135.04, 24.62]
IMAGEN −33.58 [ −59.58, −7.58]
DBSOS −31.26 [ −115.64, 53.11]
NIH_eur −13.21 [ −104.29, 77.88]
Brainscale −20.21 [ −75.44, 35.03]
GenR_eur −100.53 [ −165.75, −35.31]
FE Model −35.49 [ −48.20, −22.78]
−1000
−500
0
500
1000
Observed Outcome

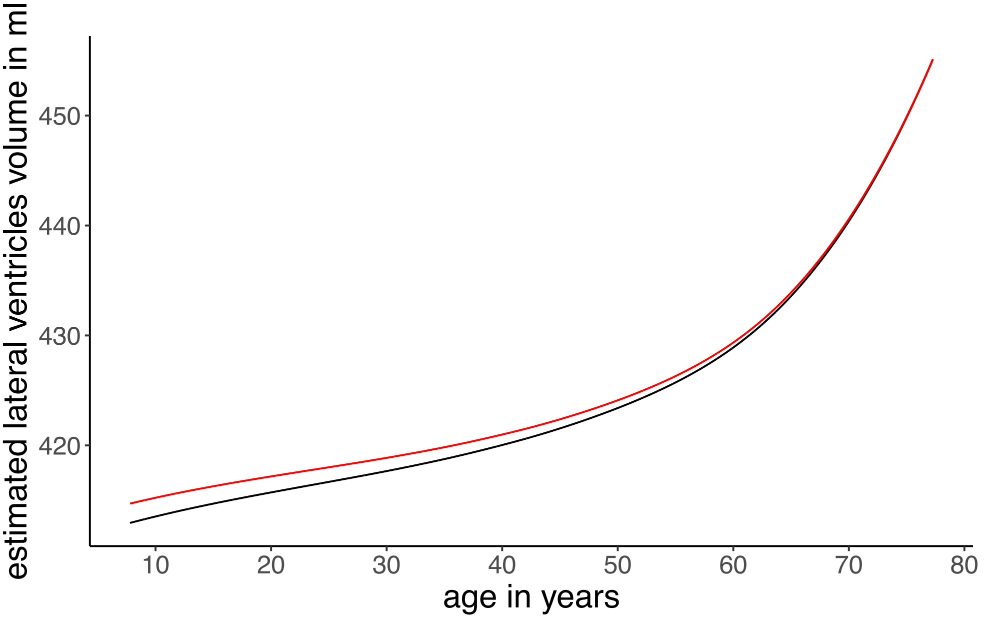
estimated lateral ventricles volume in ml
450
440
430
420
10
20
30
40
50
60
70
80
age in years

5E

**change rate cerebellum white matter**

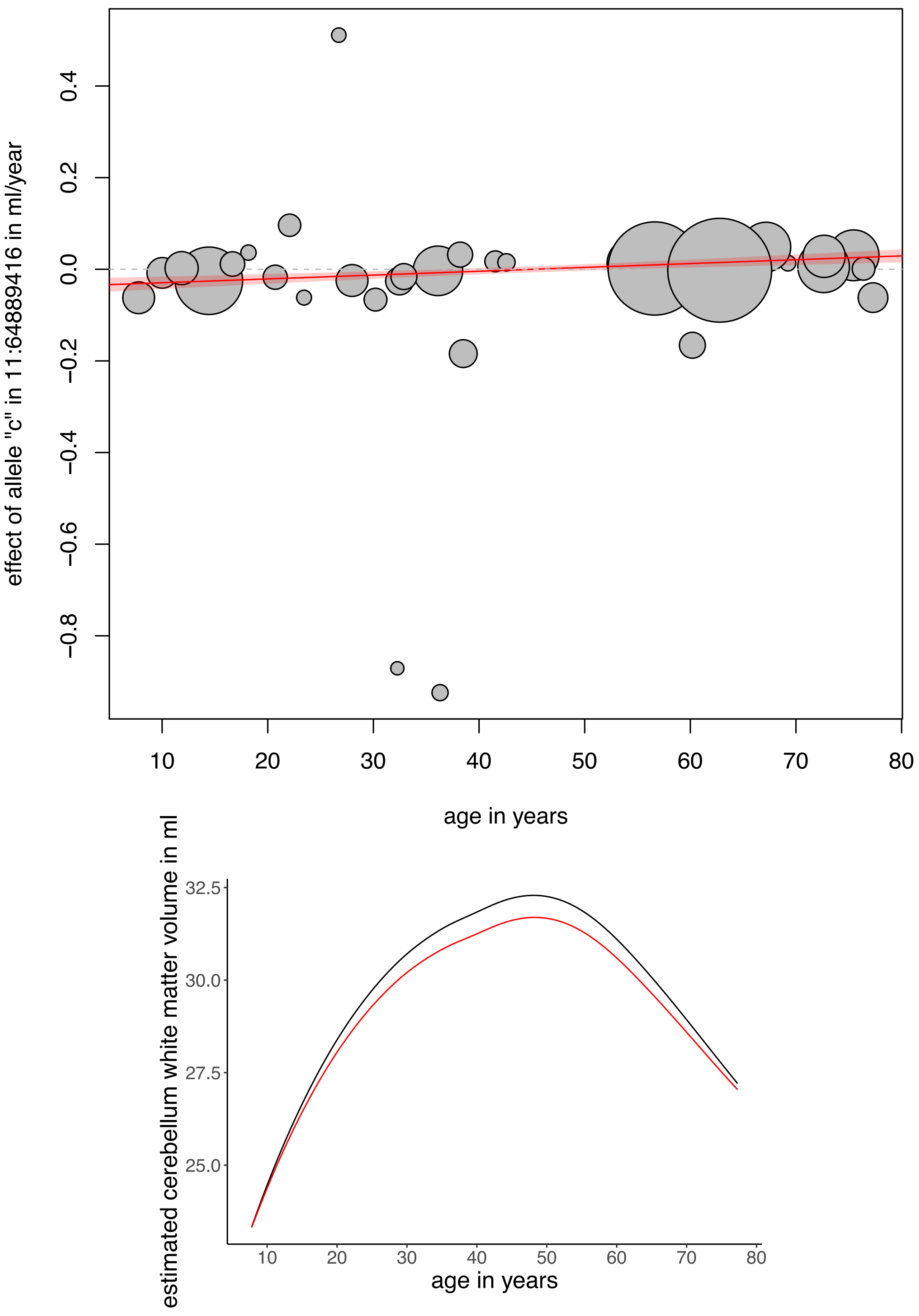

5F

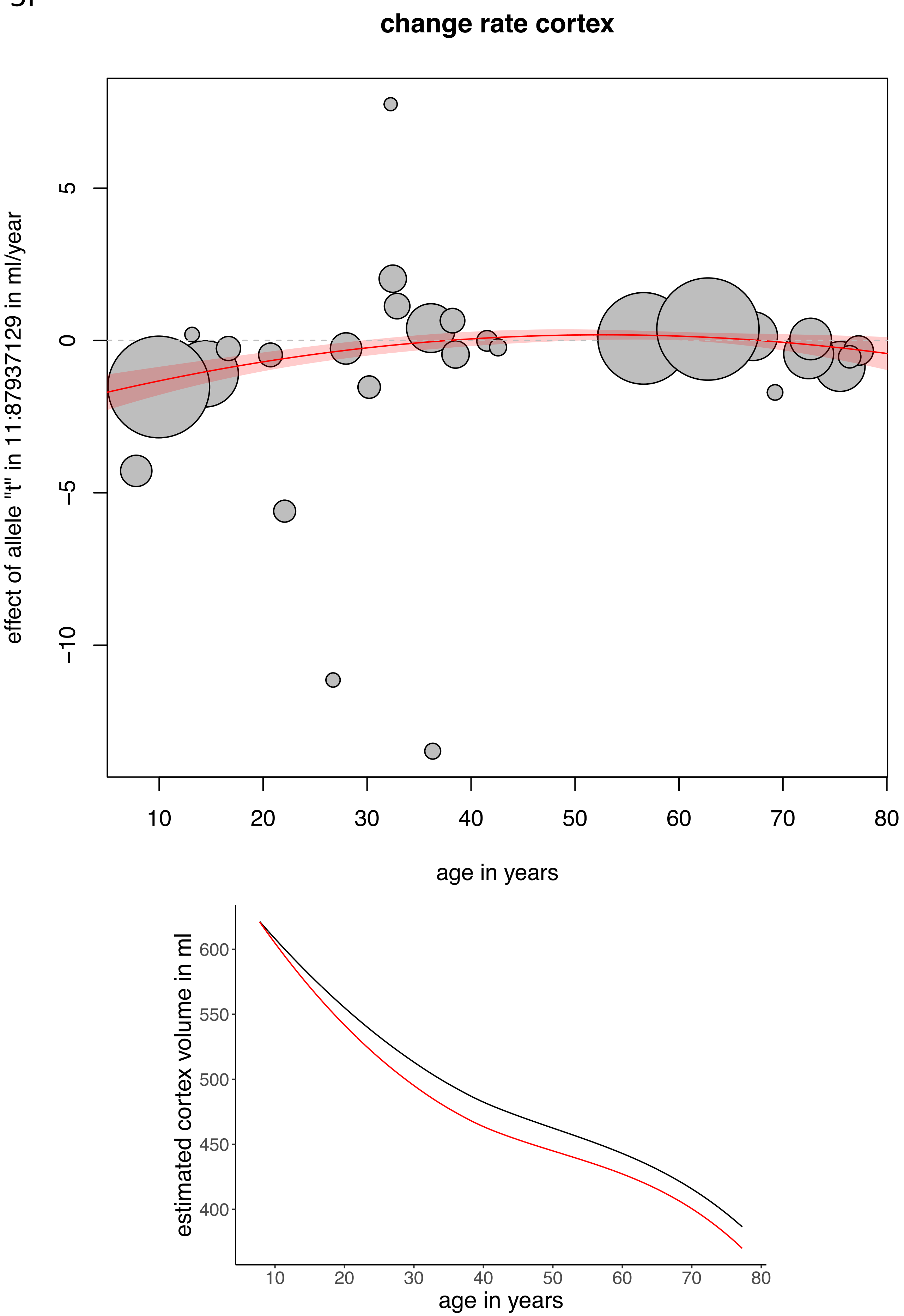

change rate cortex
effect of allele "t" in 11:87937129 in ml/year
age in years
estimated cortex volume in ml
age in years

5G

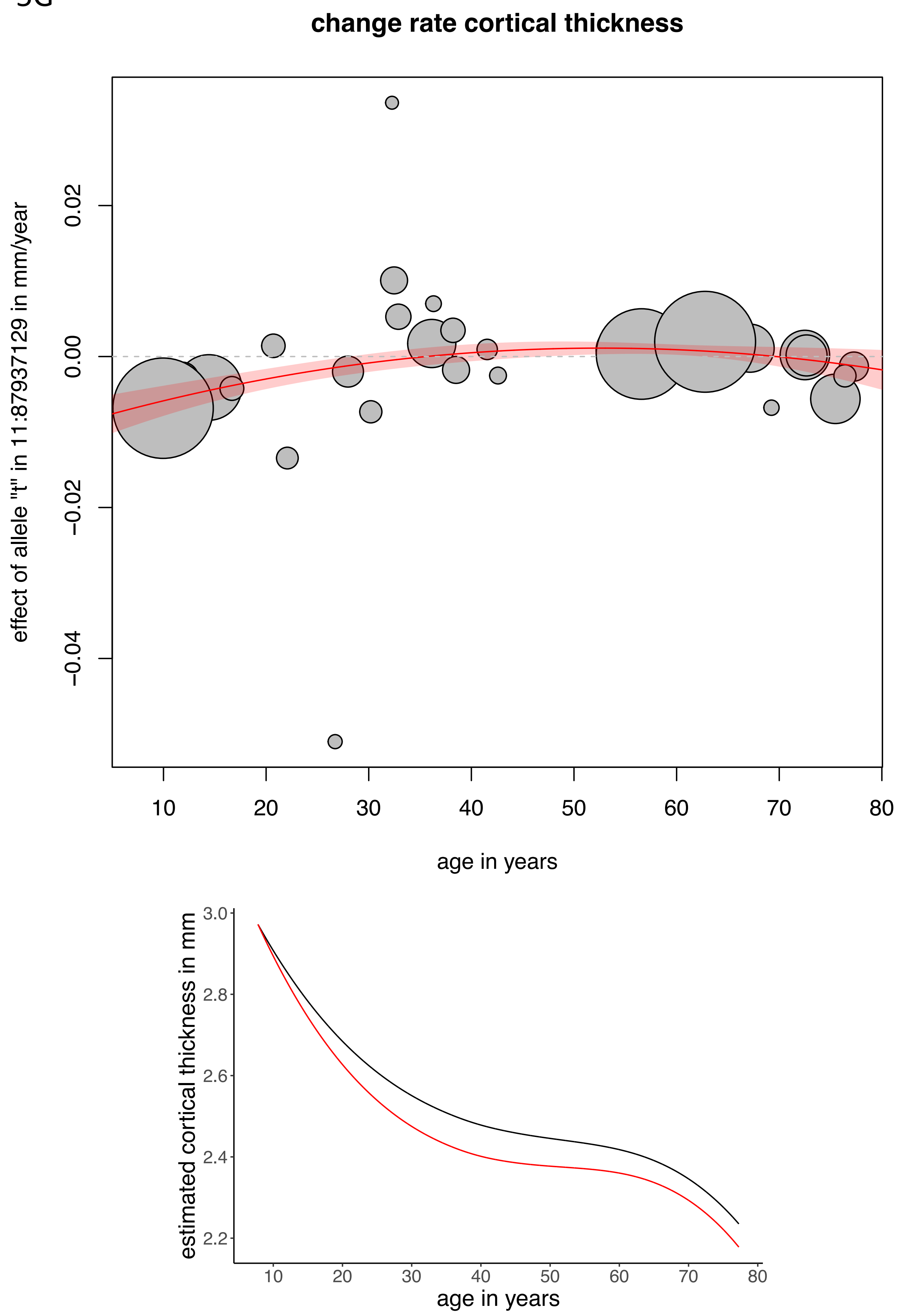

change rate cortical thickness
effect of allele "t" in 11:87937129 in mm/year
0.02
0.00
−0.02
−0.04
10
20
30
40
50
60
70
80
age in years
estimated cortical thickness in mm
3.0
2.8
2.6
2.4
2.2
10
20
30
40
50
60
70
80
age in years

5H

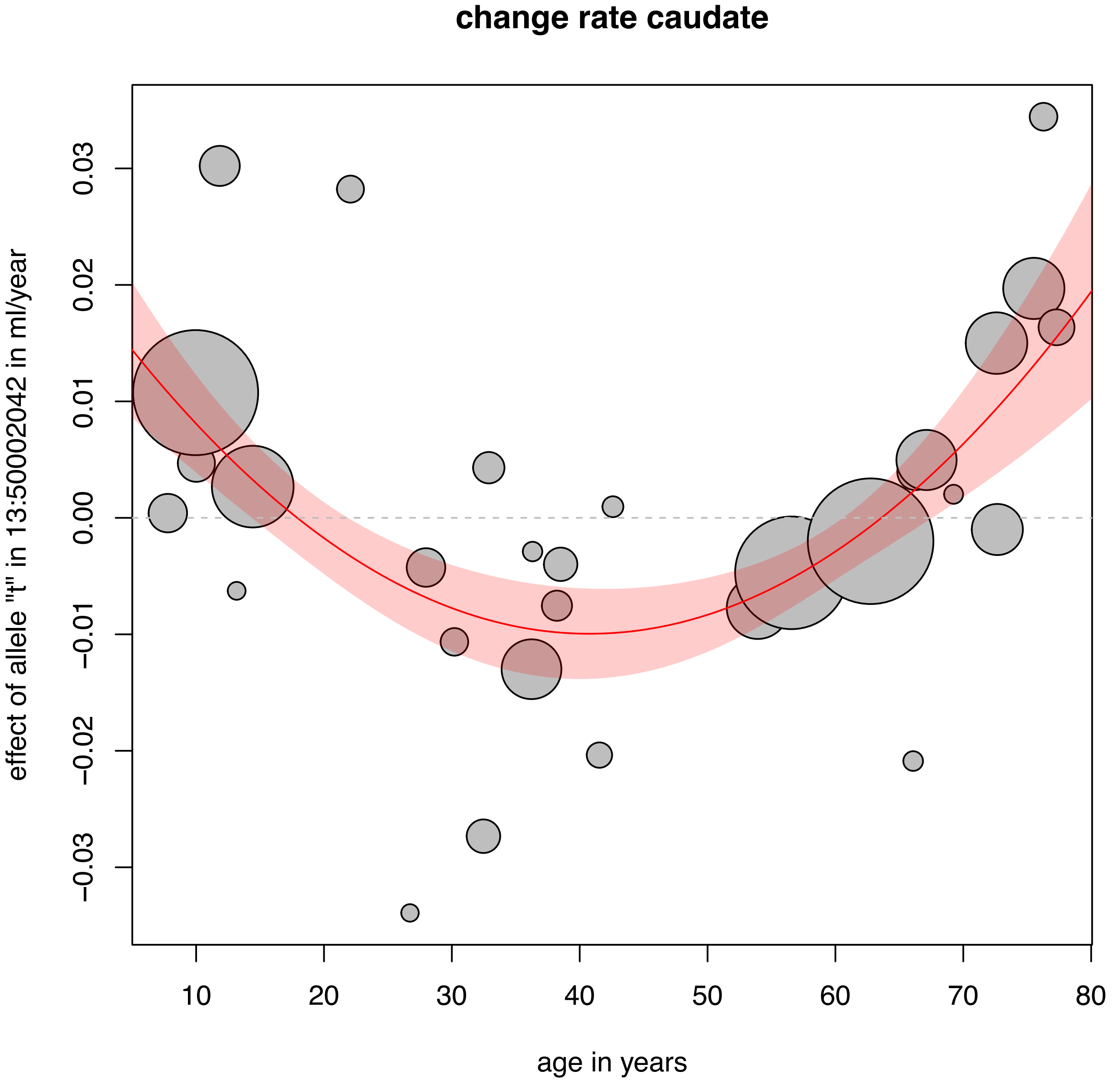

change rate caudate
effect of allele "t" in 13:50002042 in ml/year
0.03
0.02
0.01
0.00
−0.01
−0.02
−0.03
10
20
30
40
50
60
70
80
age in years


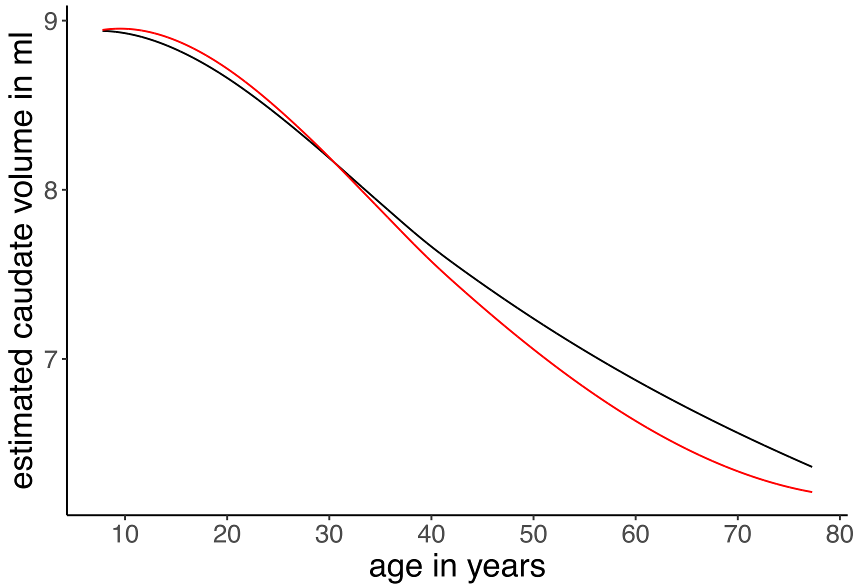

estimated caudate volume in ml
9
8
7
10
20
30
40
50
60
70
80
age in years

5l

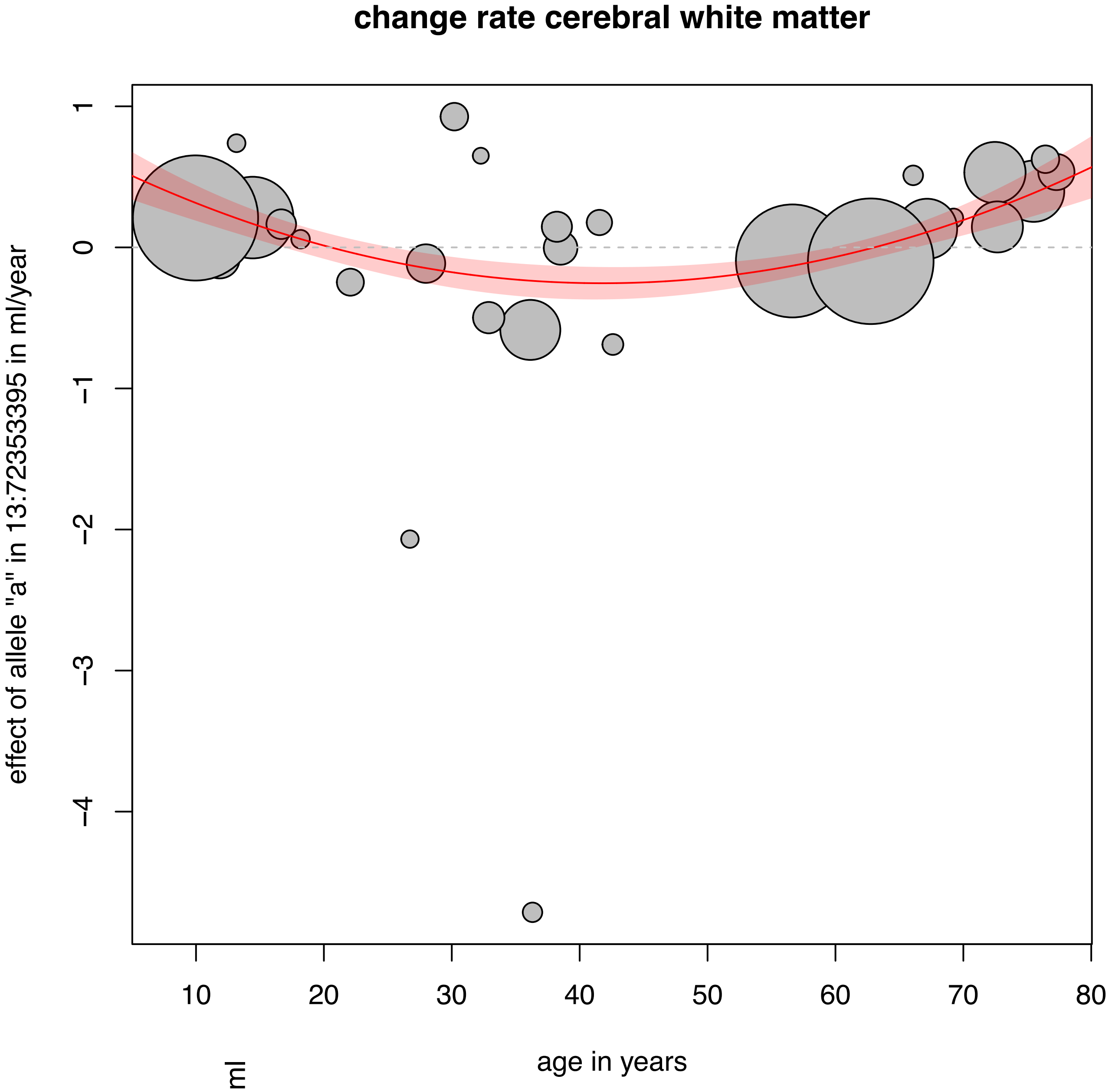
change rate cerebral white matter
effect of allele "a" in 13:72353395 in ml/year
1
0
−1
−2
−3
−4
10
20
30
40
50
60
70
80
age in years

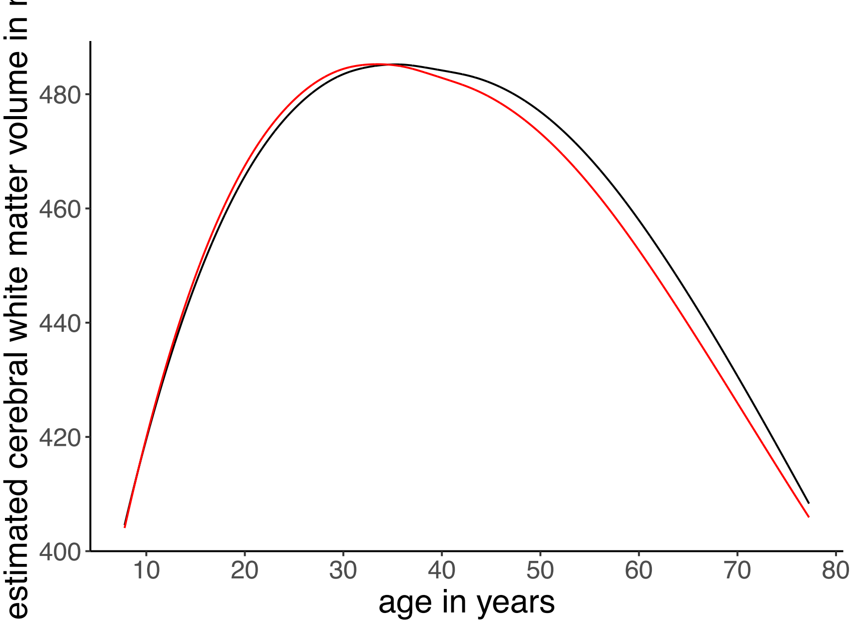
estimated cerebral white matter volume in ml
400
420
440
460
480
10
20
30
40
50
60
70
80
age in years

5J

#### change rate Vent; effect of allele "a" in 16:20065681

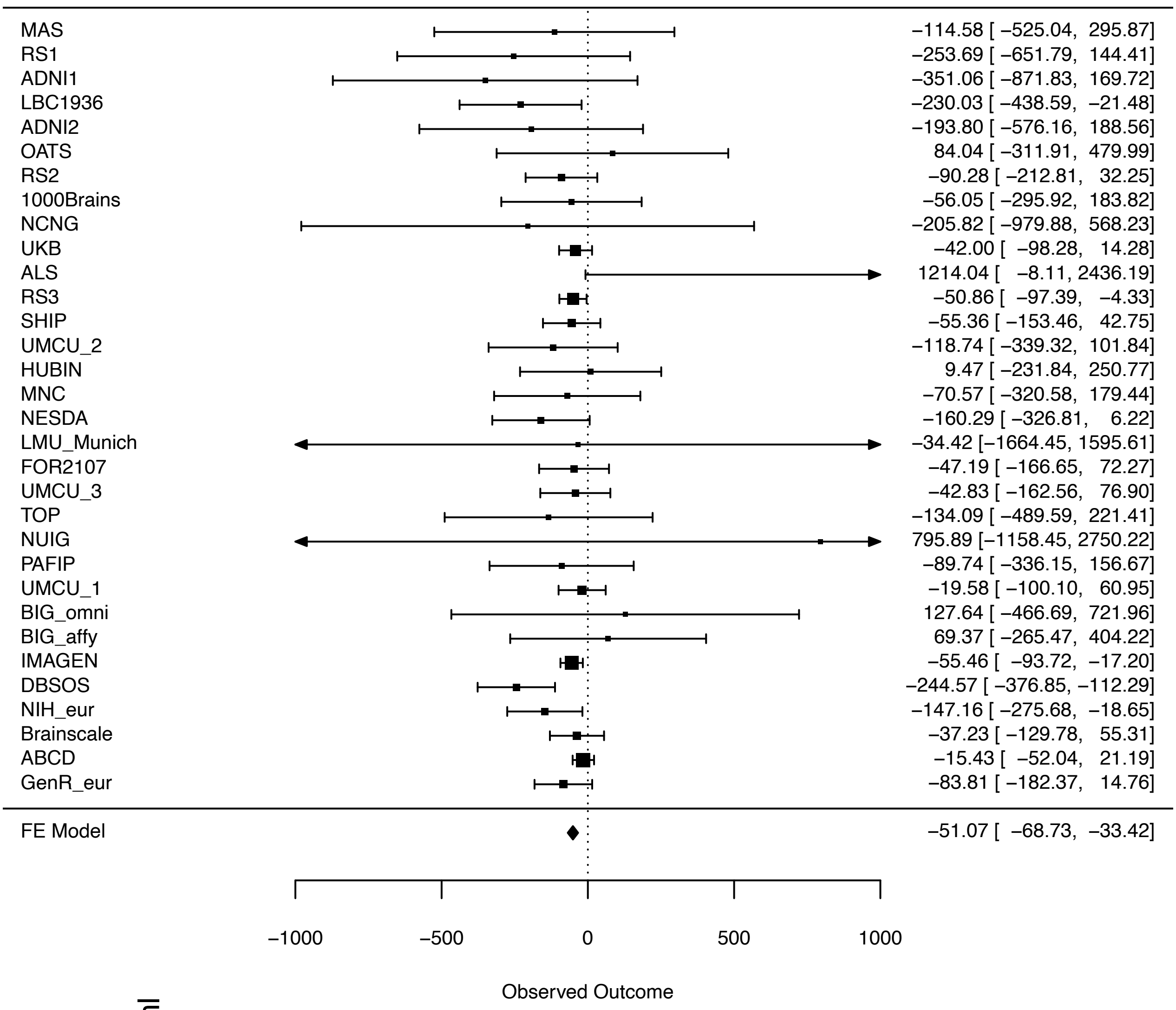


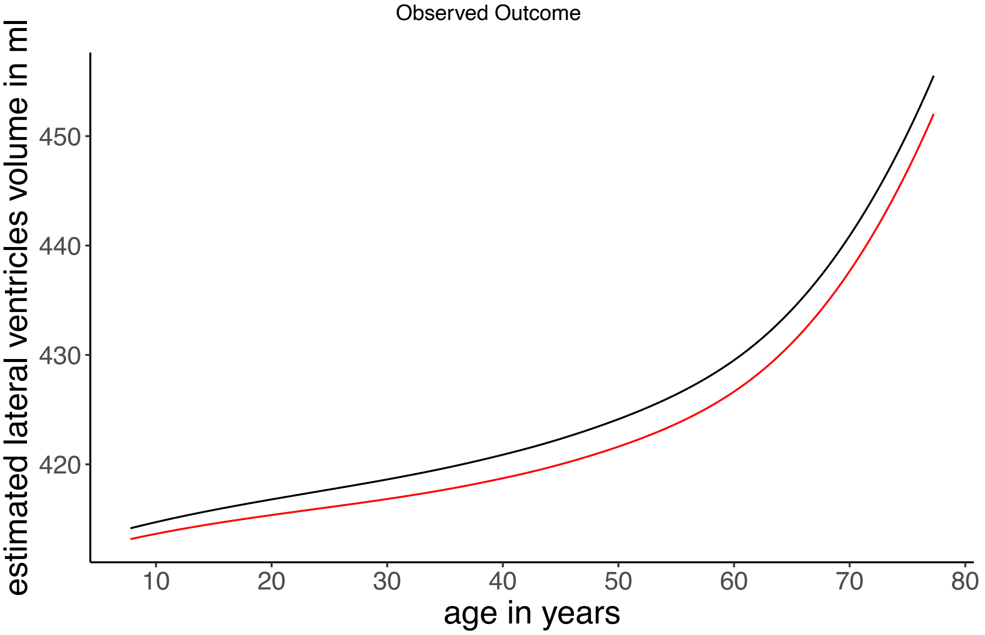

5K

#### change rate BrainSegNotVent; effect of allele "a" in 16:62059248

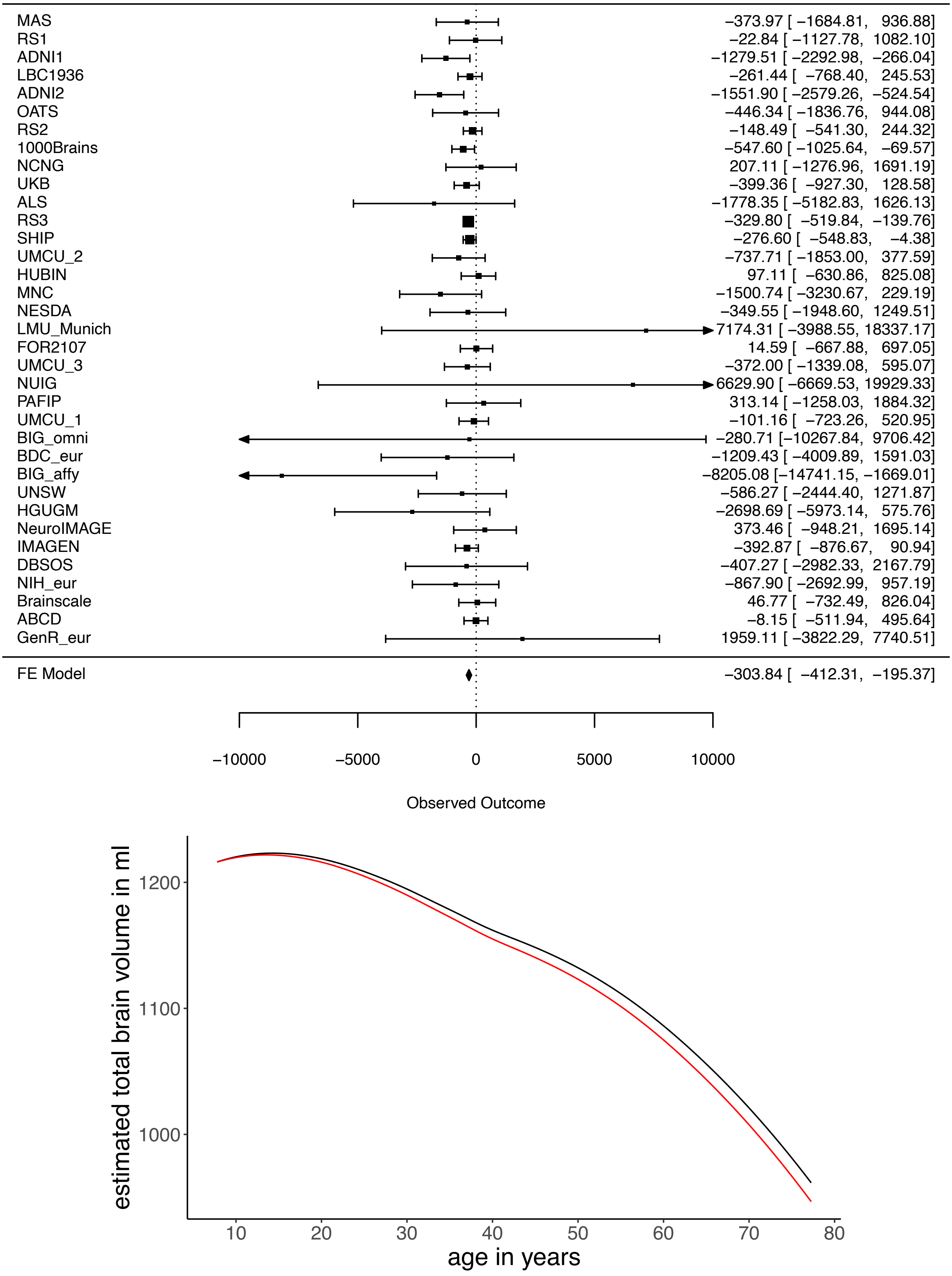

5L

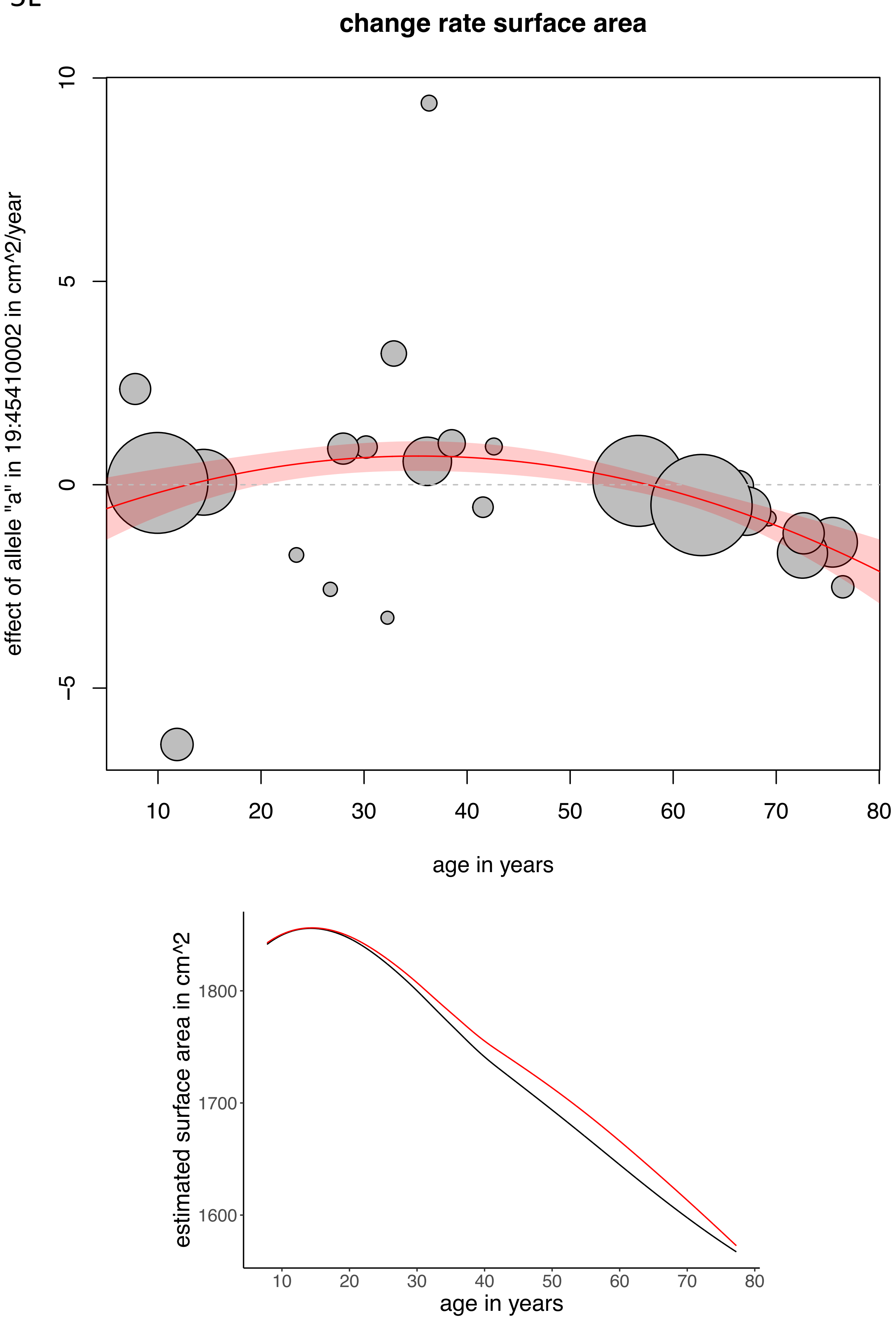

change rate surface area
effect of allele "a" in 19:45410002 in cm^2/year
10
5
0
−5
10
20
30
40
50
60
70
80
age in years
estimated surface area in cm^2
1800
1700
1600
10
20
30
40
50
60
70
80
age in years

5M

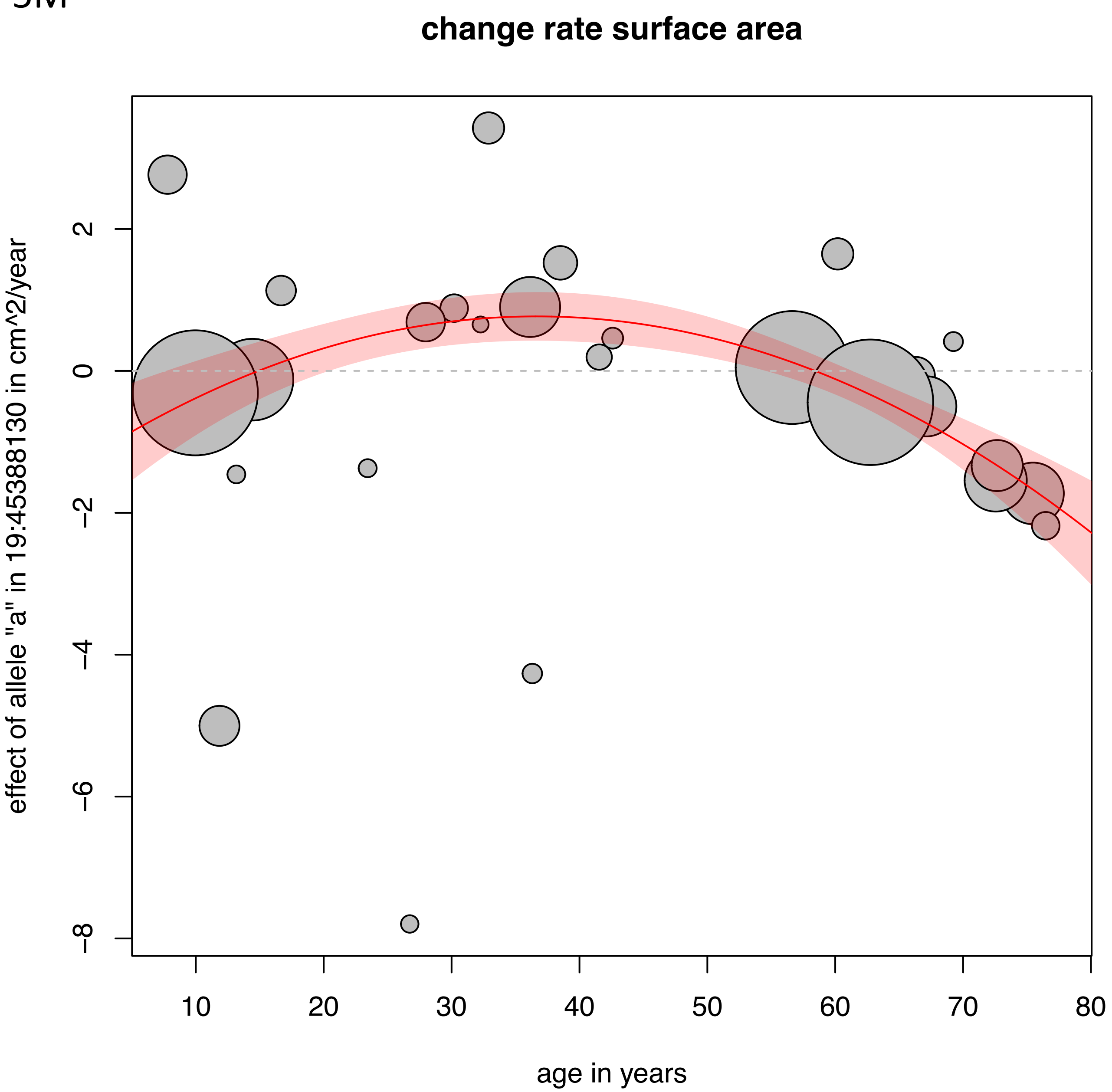

change rate surface area
effect of allele "a" in 19:45388130 in cm^2/year
age in years
2
0
−2
−4
−6
−8
10
20
30
40
50
60
70
80


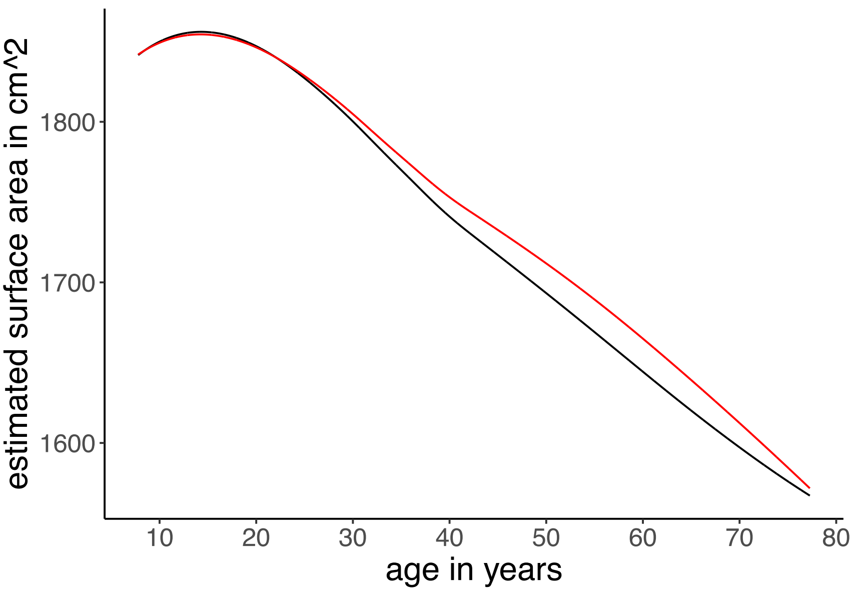

estimated surface area in cm^2
age in years
1800
1700
1600
10
20
30
40
50
60
70
80

5N

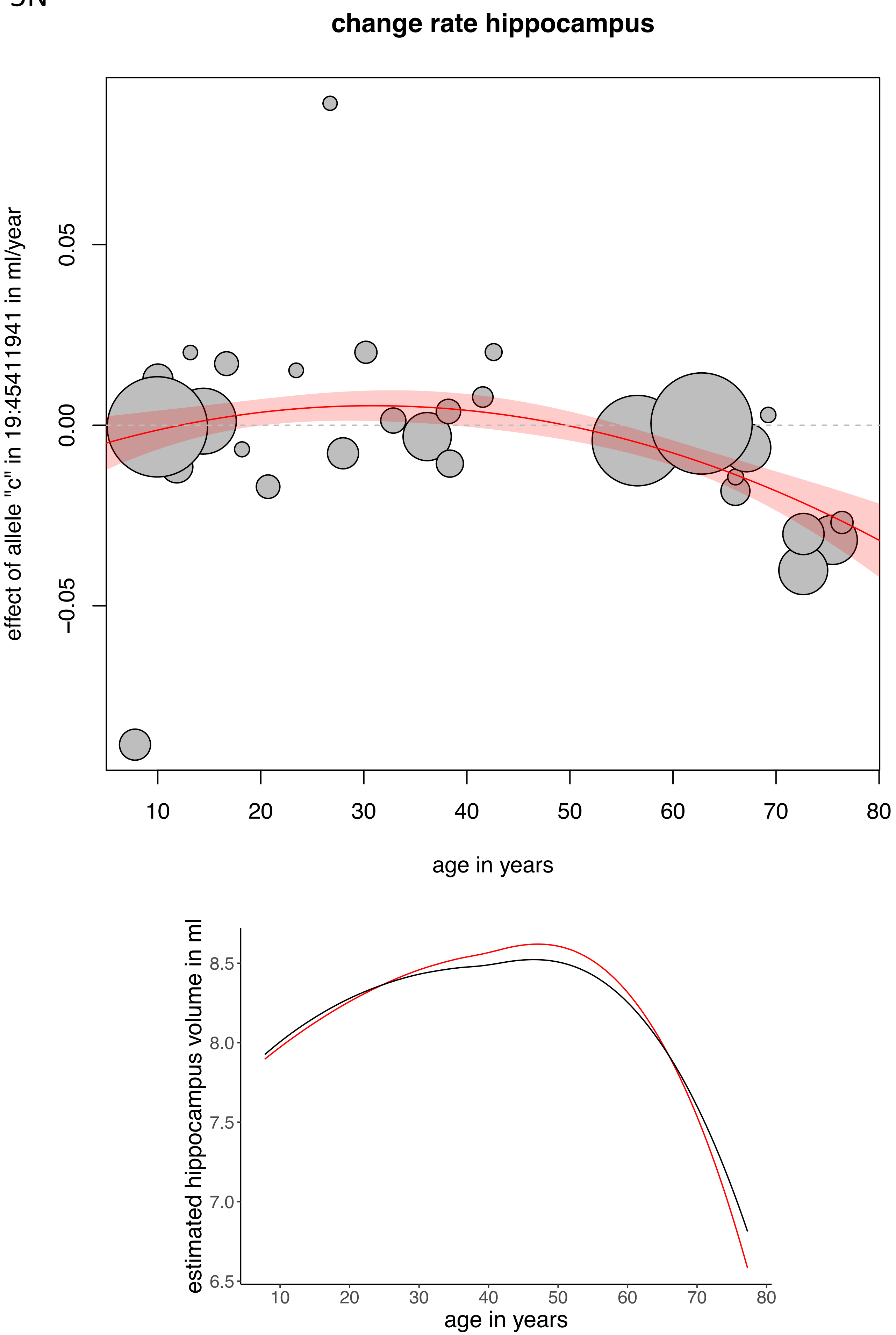

change rate hippocampus
effect of allele "c" in 19:45411941 in ml/year
0.05
0.00
−0.05
10
20
30
40
50
60
70
80
age in years
estimated hippocampus volume in ml
8.5
8.0
7.5
7.0
6.5
10
20
30
40
50
60
70
80
age in years

50

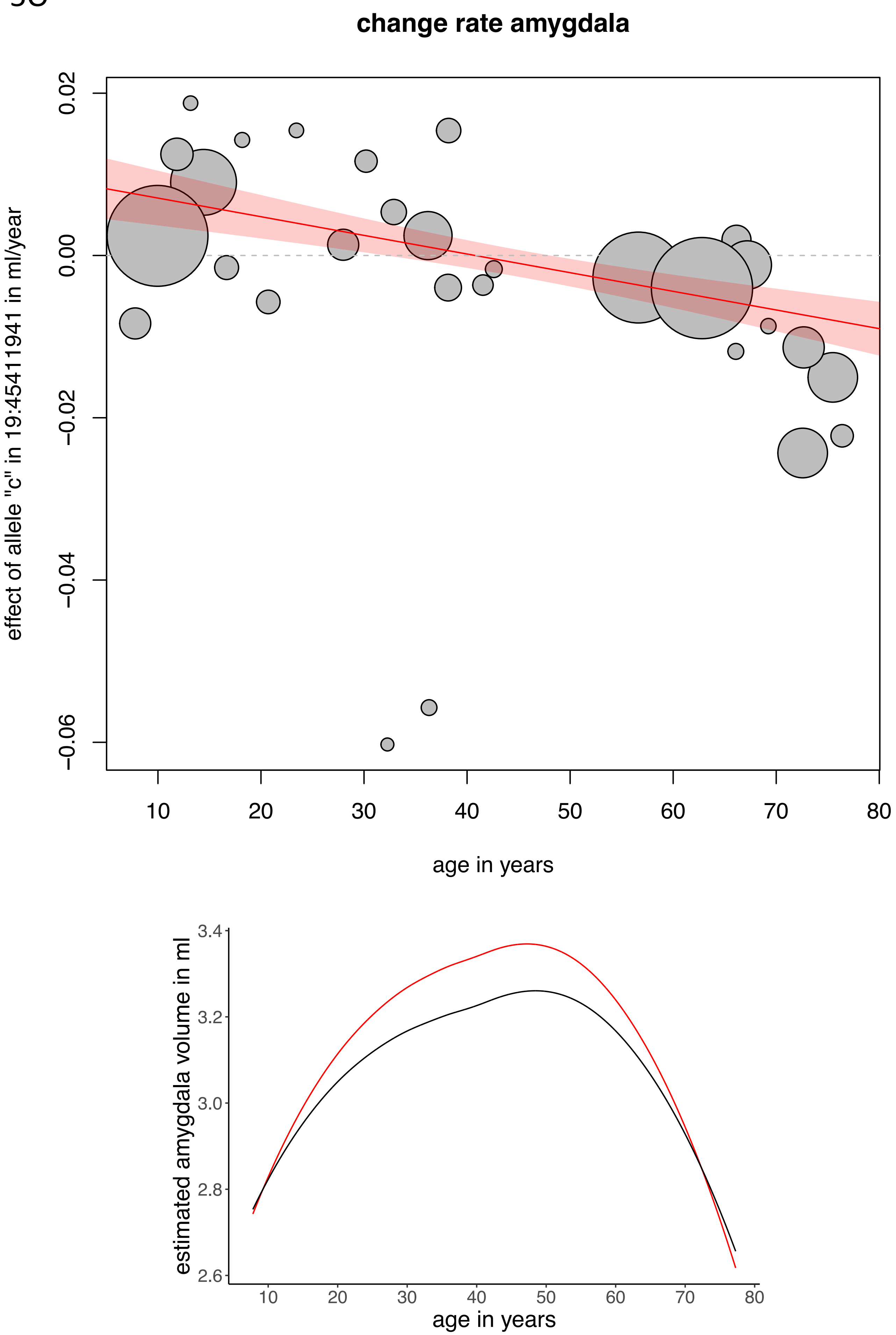

change rate amygdala
effect of allele "c" in 19:45411941 in ml/year
0.02
0.00
−0.02
−0.04
−0.06
10
20
30
40
50
60
70
80
age in years
estimated amygdala volume in ml
3.4
3.2
3.0
2.8
2.6
10
20
30
40
50
60
70
80
age in years

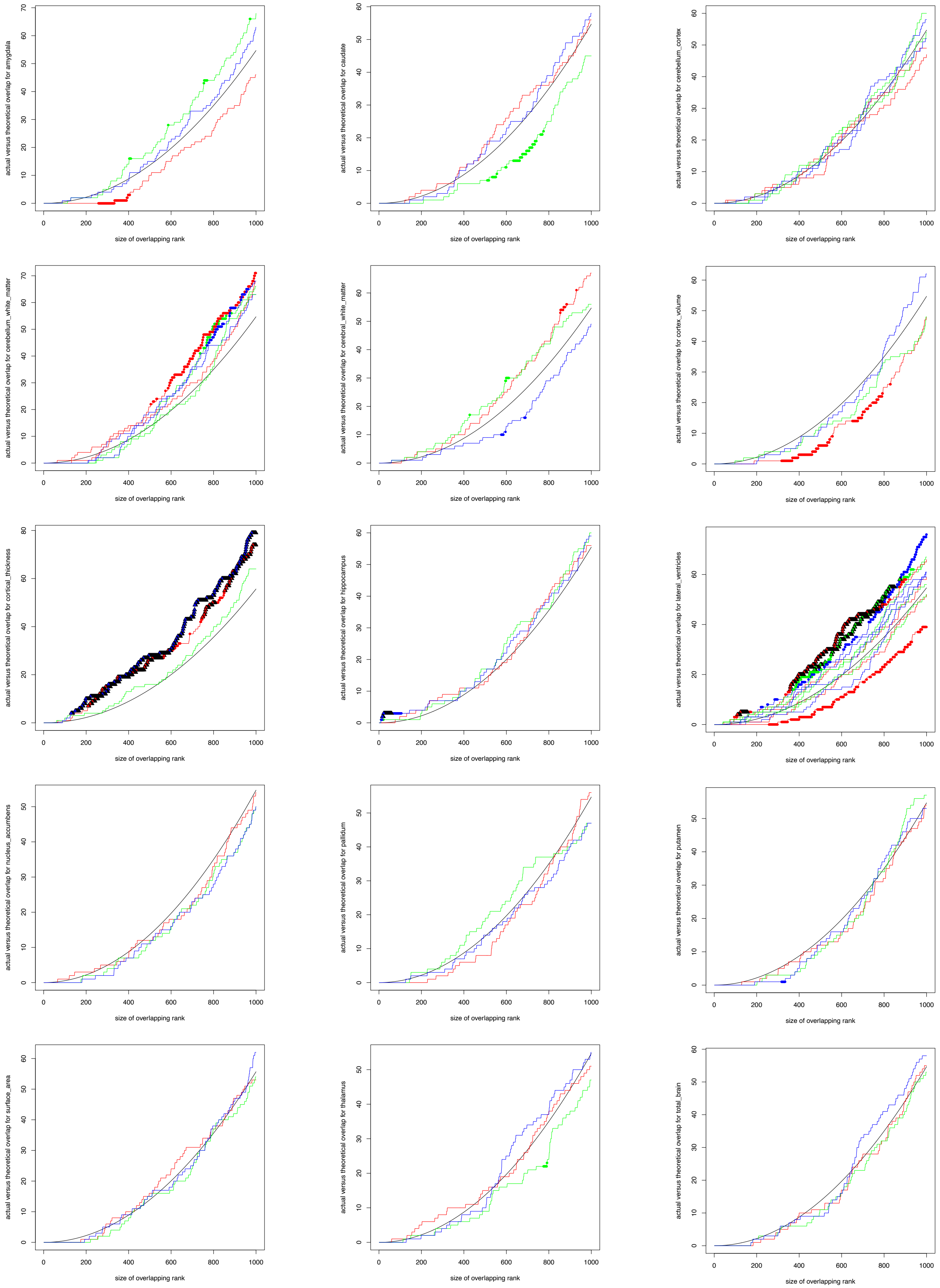
actual versus theoretical overlap for amygdala
actual versus theoretical overlap for caudate
actual versus theoretical overlap for cerebellum_cortex
actual versus theoretical overlap for cerebellum_white_matter
actual versus theoretical overlap for cerebral_white_matter
actual versus theoretical overlap for cortex_volume
actual versus theoretical overlap for cortical_thickness
actual versus theoretical overlap for hippocampus
actual versus theoretical overlap for lateral_ventricles
actual versus theoretical overlap for nucleus_accumbens
actual versus theoretical overlap for pallidum
actual versus theoretical overlap for putamen
actual versus theoretical overlap for surface_area
actual versus theoretical overlap for thalamus
actual versus theoretical overlap for total_brain
size of overlapping rank

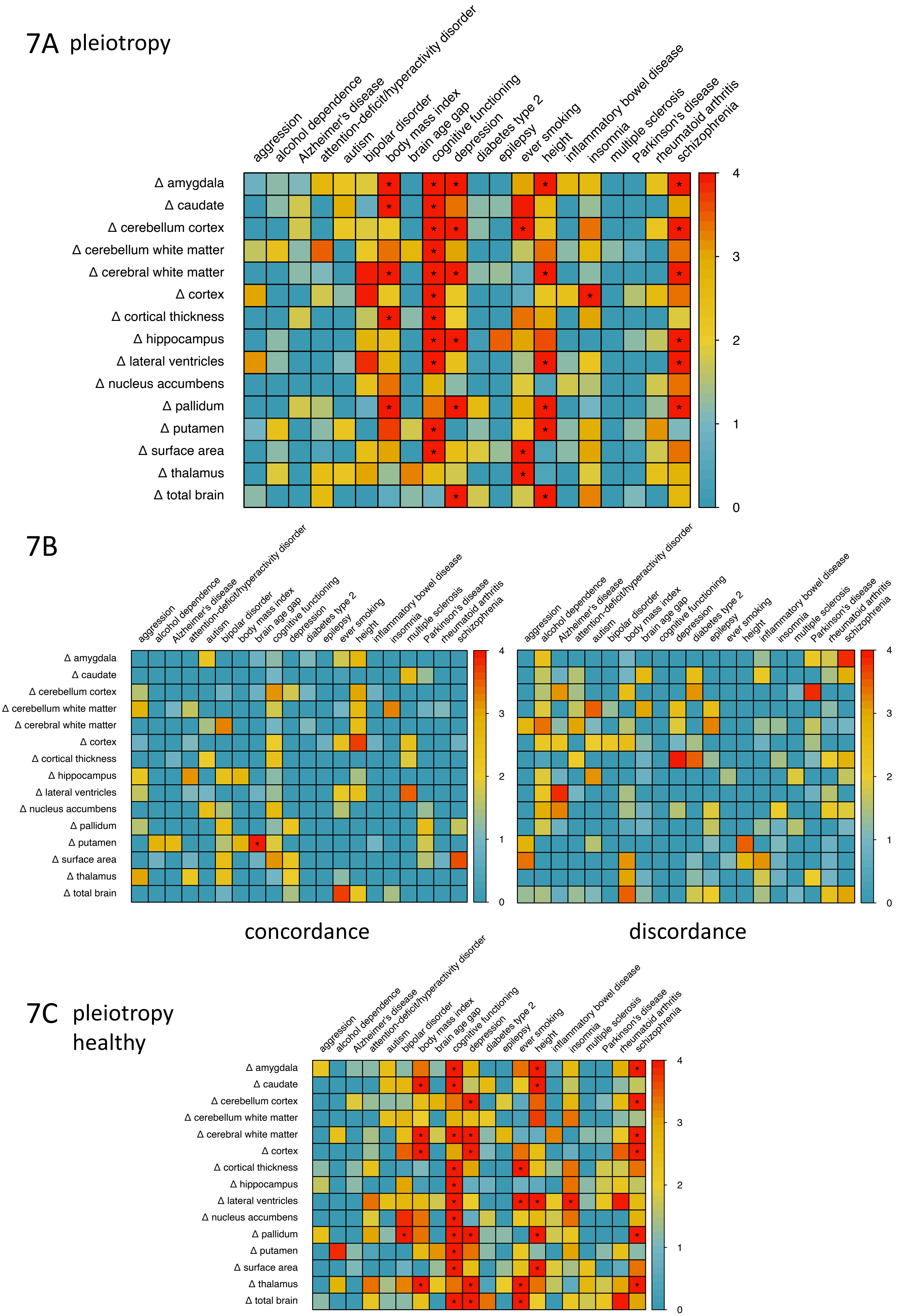
7A pleiotropy
aggression
alcohol dependence
Alzheimer's disease
attention-deficit/hyperactivity disorder
autism
bipolar disorder
body mass index
brain age gap
cognitive functioning
depression
diabetes type 2
epilepsy
ever smoking
height
inflammatory bowel disease
insomnia
multiple sclerosis
Parkinson's disease
rheumatoid arthritis
schizophrenia
Δ amygdala
Δ caudate
Δ cerebellum cortex
Δ cerebellum white matter
Δ cerebral white matter
Δ cortex
Δ cortical thickness
Δ hippocampus
Δ lateral ventricles
Δ nucleus accumbens
Δ pallidum
Δ putamen
Δ surface area
Δ thalamus
Δ total brain
4
3
2
1
0
7B
concordance
discordance
7C pleiotropy healthy

7D pleiotropy

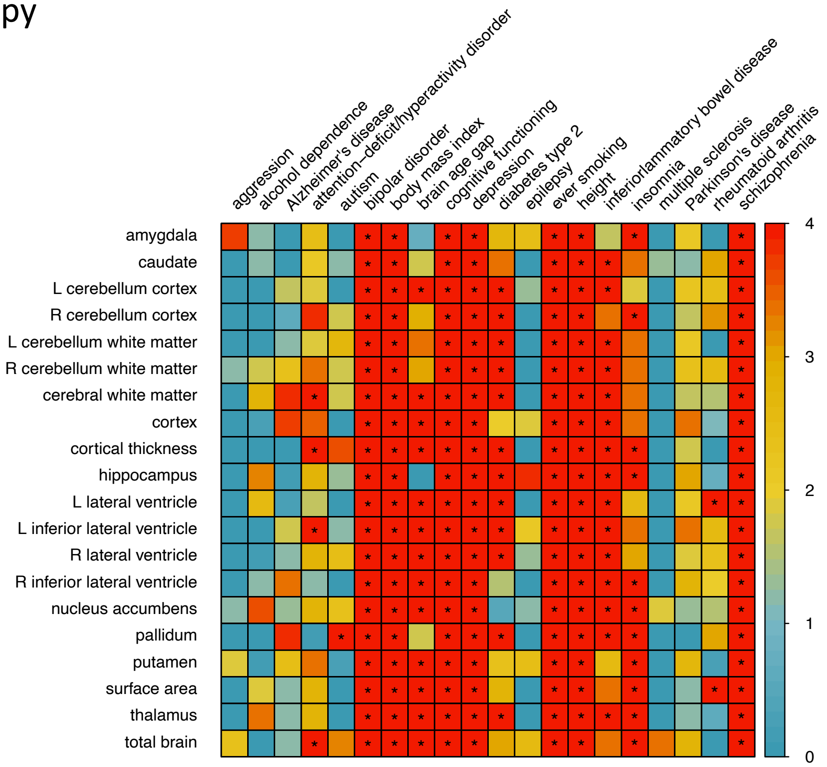


7E

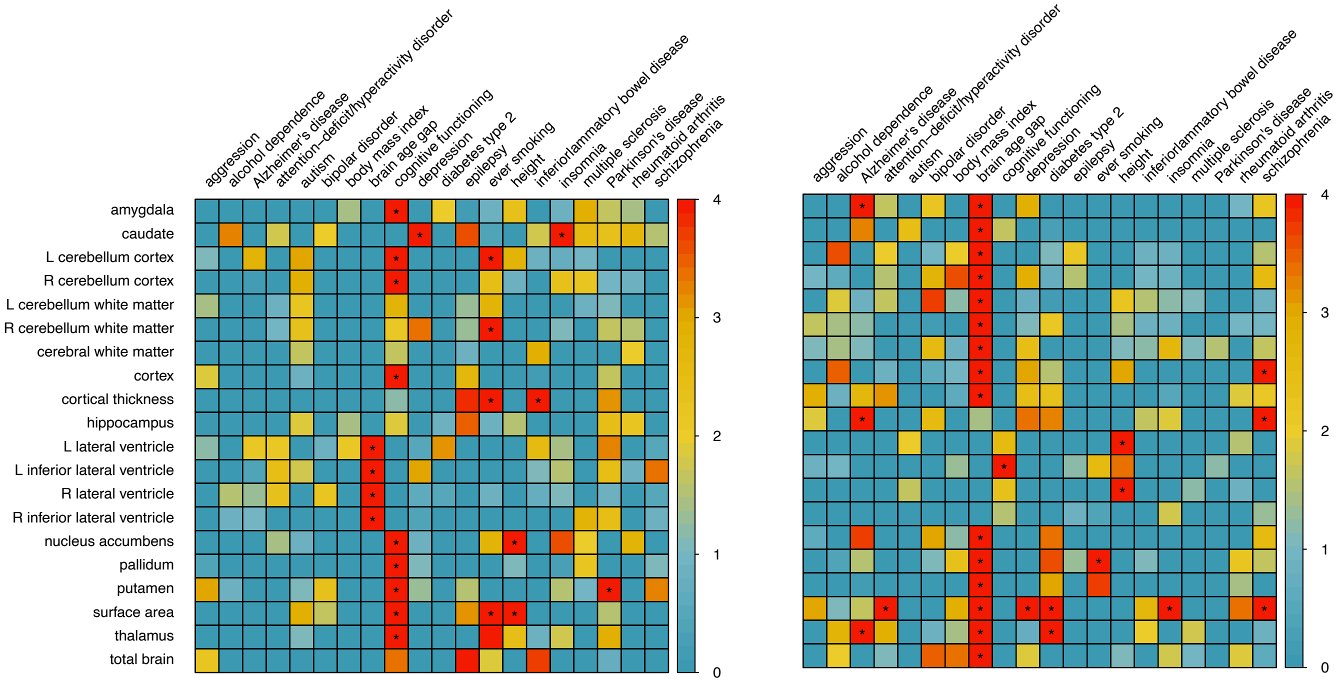


concordance discordance

8A

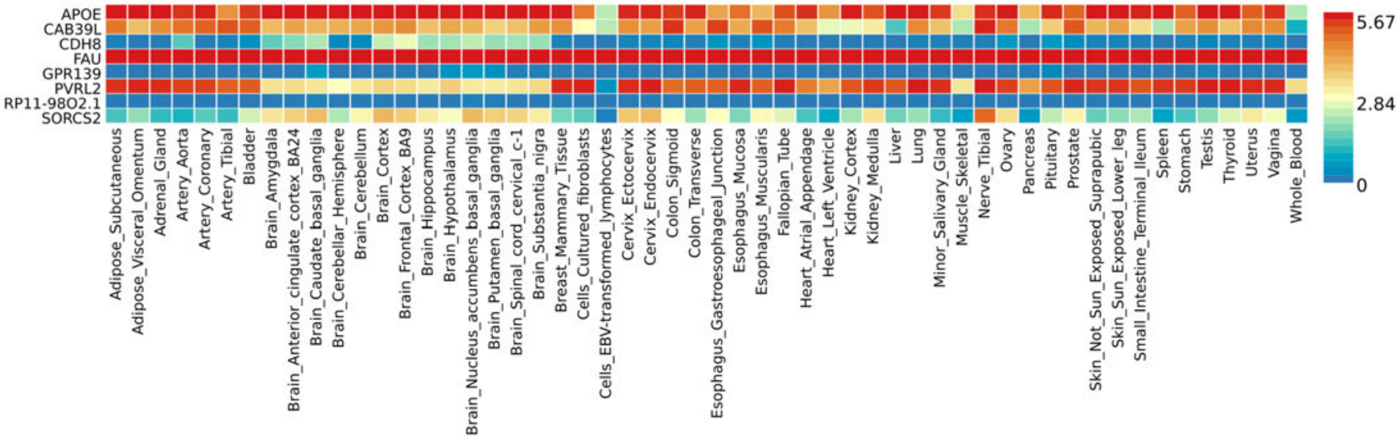
APOE
CAB39L
CDH8
FAU
GPR139
PVRL2
RP11-98O2.1
SORCS2
Adipose_Subcutaneous
Adipose_Visceral_Omentum
Adrenal_Gland
Artery_Aorta
Artery_Coronary
Artery_Tibial
Bladder
Brain_Amygdala
Brain_Anterior_cingulate_cortex_BA24
Brain_Caudate_basal_ganglia
Brain_Cerebellar_Hemisphere
Brain_Cerebellum
Brain_Cortex
Brain_Frontal_Cortex_BA9
Brain_Hippocampus
Brain_Hypothalamus
Brain_Nucleus_accumbens_basal_ganglia
Brain_Putamen_basal_ganglia
Brain_Spinal_cord_cervical_c-1
Brain_Substantia_nigra
Breast_Mammary_Tissue
Cells_Cultured_fibroblasts
Cells_EBV-transformed_lymphocytes
Cervix_Ectocervix
Cervix_Endocervix
Colon_Sigmoid
Colon_Transverse
Esophagus_Gastroesophageal_Junction
Esophagus_Mucosa
Esophagus_Muscularis
Fallopian_Tube
Heart_Atrial_Appendage
Heart_Left_Ventricle
Kidney_Cortex
Kidney_Medulla
Liver
Lung
Minor_Salivary_Gland
Muscle_Skeletal
Nerve_Tibial
Ovary
Pancreas
Pituitary
Prostate
Skin_Not_Sun_Exposed_Suprapubic
Skin_Sun_Exposed_Lower_leg
Small_Intestine_Terminal_Ileum
Spleen
Stomach
Testis
Thyroid
Uterus
Vagina
Whole_Blood
5.67
2.84
0

8B

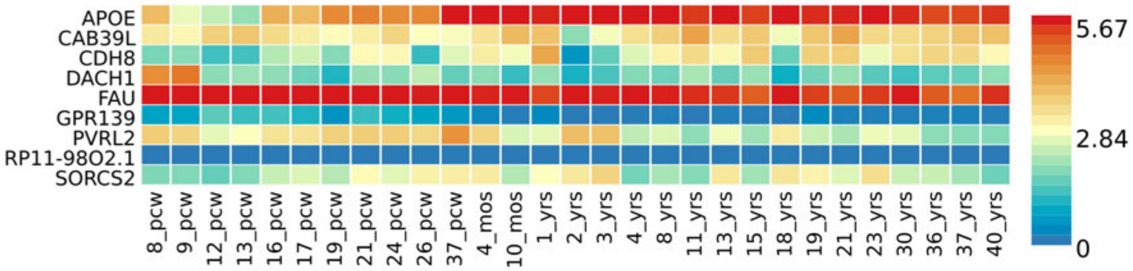
APOE
CAB39L
CDH8
DACH1
FAU
GPR139
PVRL2
RP11-98O2.1
SORCS2
8_pcw
9_pcw
12_pcw
13_pcw
16_pcw
17_pcw
19_pcw
21_pcw
24_pcw
26_pcw
37_pcw
4_mos
10_mos
1_yrs
2_yrs
3_yrs
4_yrs
8_yrs
11_yrs
13_yrs
15_yrs
18_yrs
19_yrs
21_yrs
23_yrs
30_yrs
36_yrs
37_yrs
40_yrs
5.67
2.84
0

# 9A

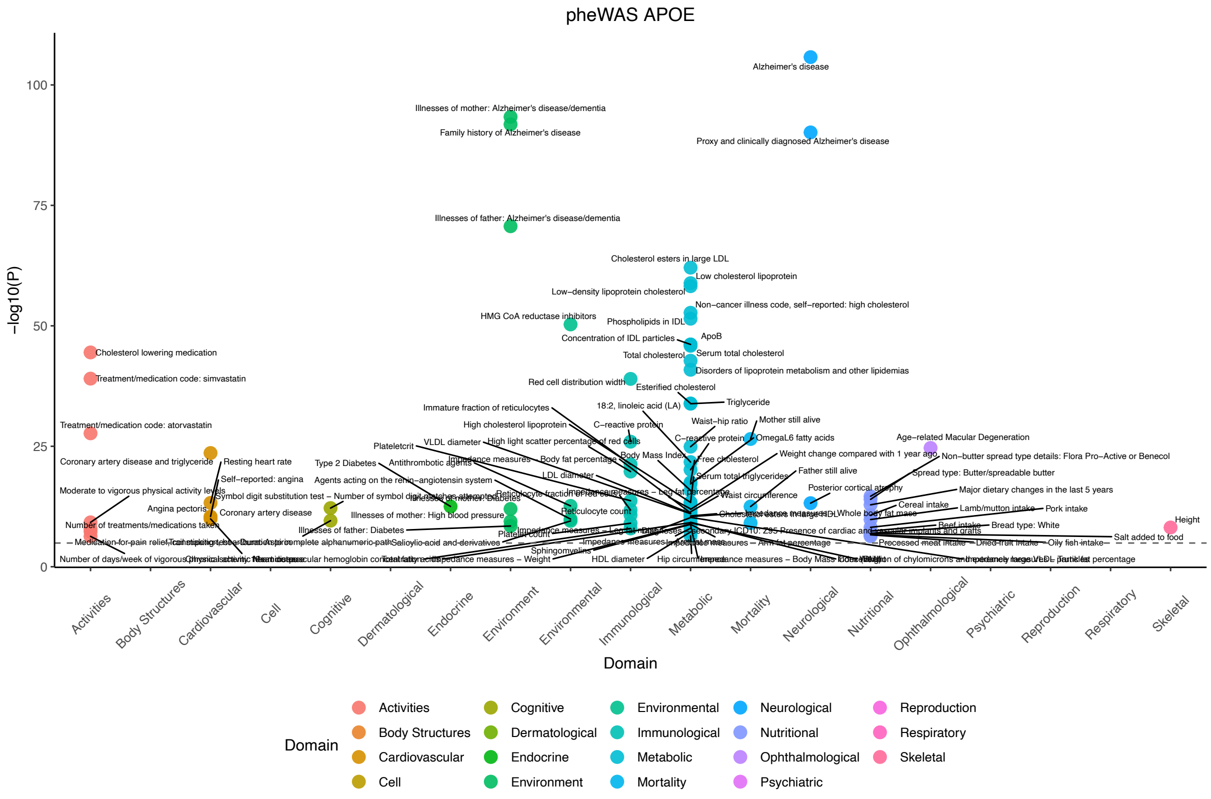
pheWAS APOE
Alzheimer's disease
Illnesses of mother: Alzheimer's disease/dementia
Family history of Alzheimer's disease
Proxy and clinically diagnosed Alzheimer's disease
Illnesses of father: Alzheimer's disease/dementia
Cholesterol esters in large LDL
Low cholesterol lipoprotein
Low−density lipoprotein cholesterol
Non−cancer illness code, self−reported: high cholesterol
HMG CoA reductase inhibitors
Phospholipids in IDL
ApoB
Concentration of IDL particles
Serum total cholesterol
Total cholesterol
Disorders of lipoprotein metabolism and other lipidemias
Cholesterol lowering medication
Treatment/medication code: simvastatin
Red cell distribution width
Esterified cholesterol
Triglyceride
Immature fraction of reticulocytes
18:2, linoleic acid (LA)
Treatment/medication code: atorvastatin
High cholesterol lipoprotein
C−reactive protein
Waist−hip ratio
Mother still alive
Plateletcrit
VLDL diameter
High light scatter percentage of red cells
OmegaL6 fatty acids
Age−related Macular Degeneration
Coronary artery disease and triglyceride
Resting heart rate
Type 2 Diabetes
Antithrombotic agents
Body Mass Index
Free cholesterol
Weight change compared with 1 year ago
Non−butter spread type details: Flora Pro−Active or Benecol
Impedance measures
Body fat percentage
Father still alive
Spread type: Butter/spreadable butter
Self−reported: angina
Agents acting on the renin−angiotensin system
LDL diameter
Serum total triglycerides
Moderate to vigorous physical activity levels
Posterior cortical atrophy
Major dietary changes in the last 5 years
Symbol digit substitution test − Number of symbol digit matches attempted
Waist circumference
Angina pectoris
Coronary artery disease
Reticulocyte count
Cereal intake
Lamb/mutton intake
Pork intake
Illnesses of mother: High blood pressure
Whole body fat mass
Height
Number of treatments/medications taken
Beef intake
Bread type: White
Illnesses of father: Diabetes
Platelet count
Salt added to food
Salicylic acid and derivatives
Processed meat intake
Dried fruit intake
Oily fish intake
Sphingomyelins
HDL diameter
Hip circumference
Number of days/week of vigorous physical activity
Domain
Activities
Body Structures
Cardiovascular
Cell
Cognitive
Dermatological
Endocrine
Environment
Environmental
Immunological
Metabolic
Mortality
Neurological
Nutritional
Ophthalmological
Psychiatric
Reproduction
Respiratory
Skeletal
−log10(P)
0
25
50
75
100

# 9B

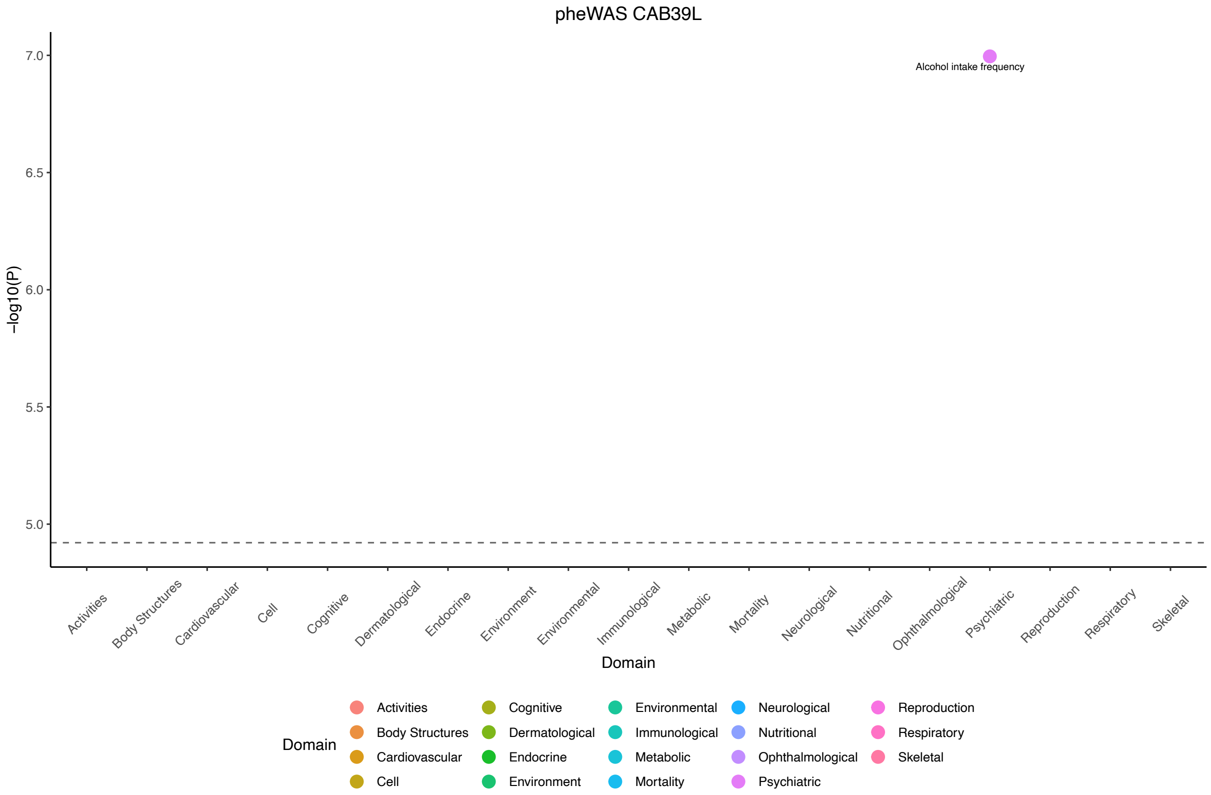
pheWAS CAB39L
Alcohol intake frequency
−log10(P)
5.0
5.5
6.0
6.5
7.0
Domain
Activities
Body Structures
Cardiovascular
Cell
Cognitive
Dermatological
Endocrine
Environment
Environmental
Immunological
Metabolic
Mortality
Neurological
Nutritional
Ophthalmological
Psychiatric
Reproduction
Respiratory
Skeletal

9C

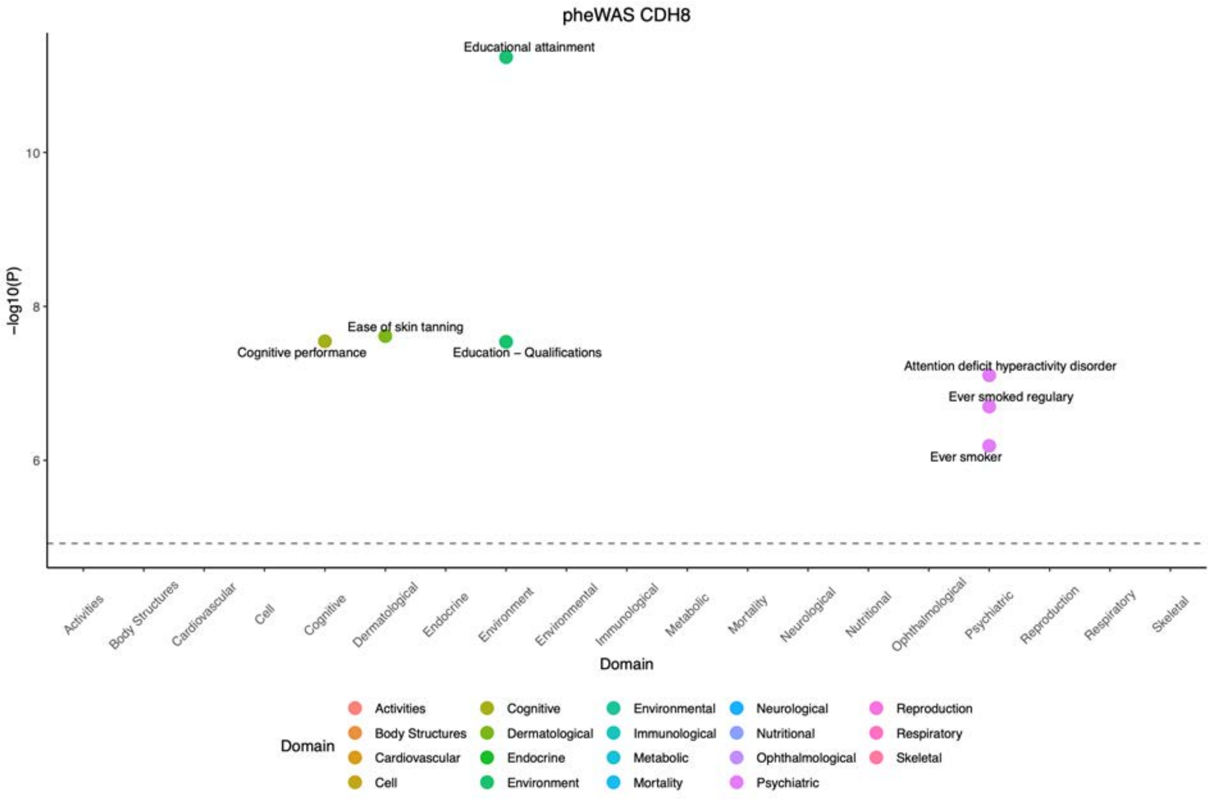

pheWAS CDH8
Educational attainment
Ease of skin tanning
Cognitive performance
Education – Qualifications
Attention deficit hyperactivity disorder
Ever smoked regulary
Ever smoker
-log10(P)
10
8
6
Activities
Body Structures
Cardiovascular
Cell
Cognitive
Dermatological
Endocrine
Environment
Environmental
Immunological
Metabolic
Mortality
Neurological
Nutritional
Ophthalmological
Psychiatric
Reproduction
Respiratory
Skeletal
Domain
Domain


9D

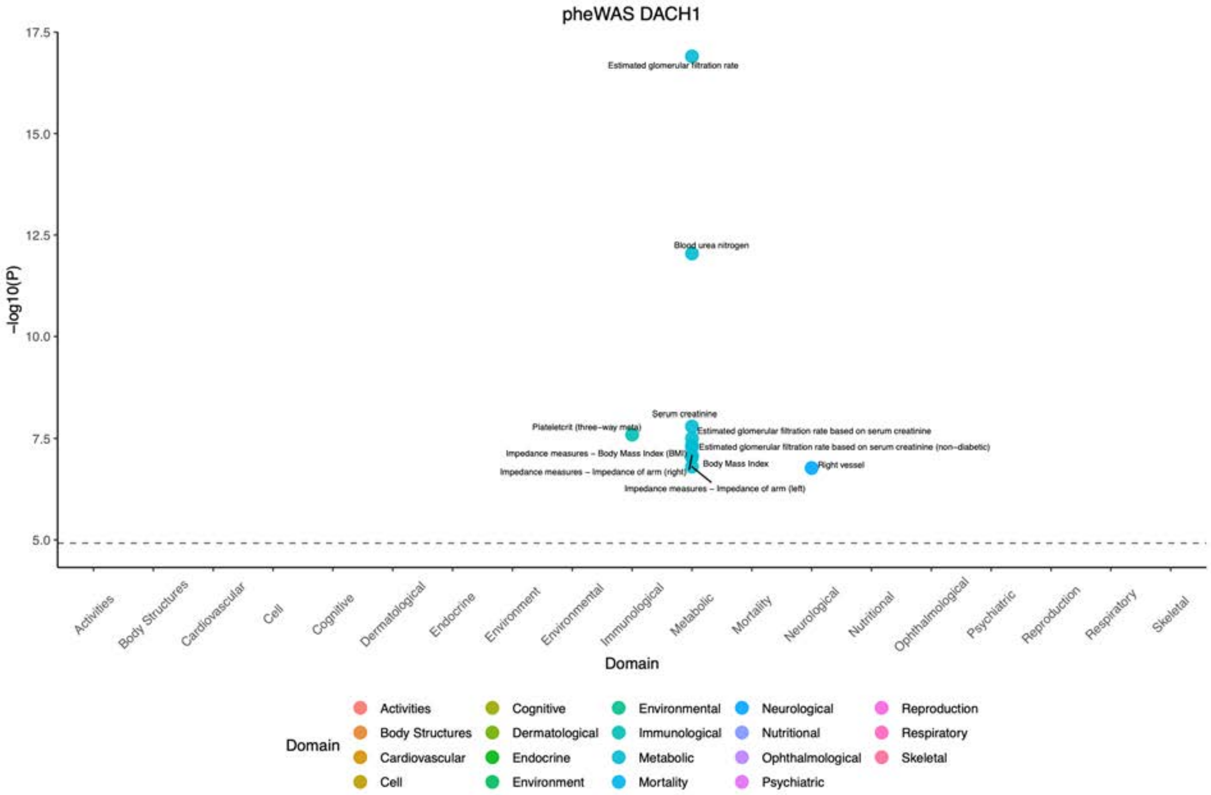

pheWAS DACH1
Estimated glomerular filtration rate
Blood urea nitrogen
Serum creatinine
Plateletcrit (three-way meta)
Estimated glomerular filtration rate based on serum creatinine
Estimated glomerular filtration rate based on serum creatinine (non-diabetic)
Impedance measures – Body Mass Index (BMI)
Body Mass Index
Right vessel
Impedance measures – Impedance of arm (right)
Impedance measures – Impedance of arm (left)
-log10(P)
17.5
15.0
12.5
10.0
7.5
5.0
Activities
Body Structures
Cardiovascular
Cell
Cognitive
Dermatological
Endocrine
Environment
Environmental
Immunological
Metabolic
Mortality
Neurological
Nutritional
Ophthalmological
Psychiatric
Reproduction
Respiratory
Skeletal
Domain
Domain

9E

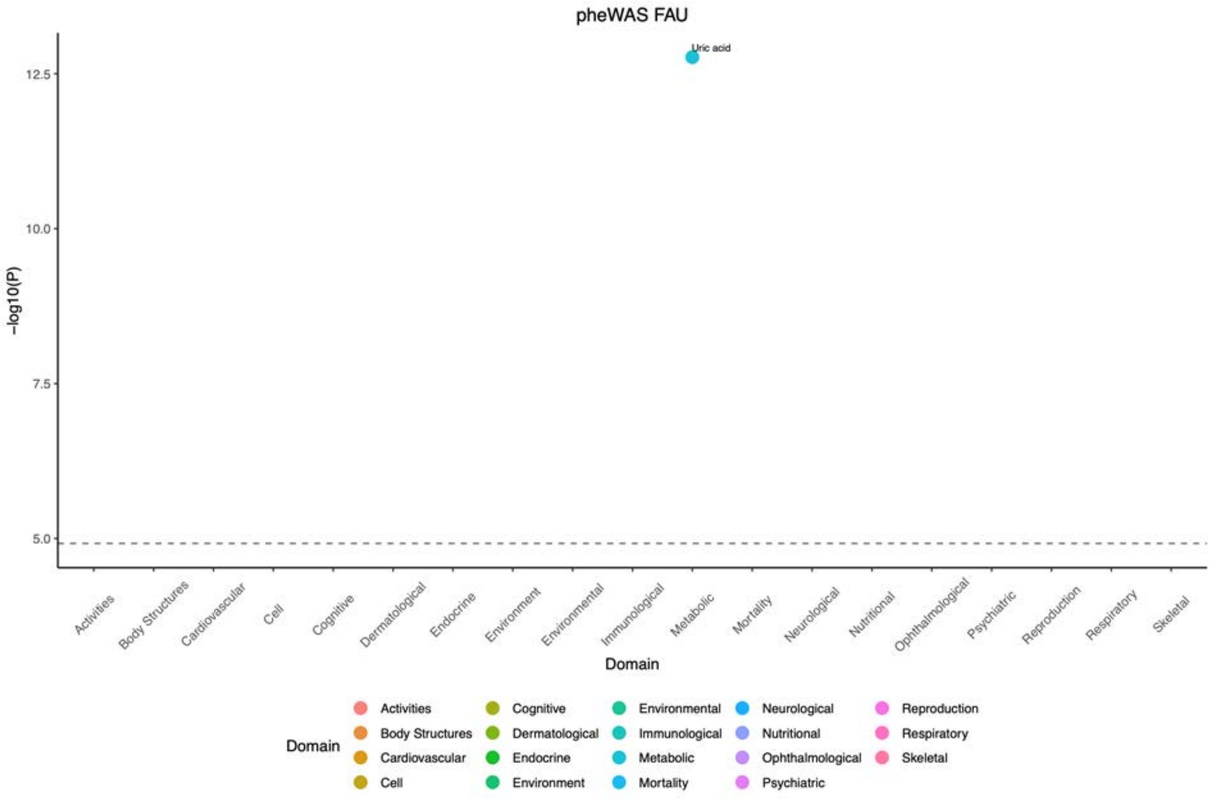
pheWAS FAU
Uric acid
12.5
10.0
7.5
5.0
-log10(P)
Activities
Body Structures
Cardiovascular
Cell
Cognitive
Dermatological
Endocrine
Environment
Environmental
Immunological
Metabolic
Mortality
Neurological
Nutritional
Ophthalmological
Psychiatric
Reproduction
Respiratory
Skeletal
Domain
Domain
Activities
Body Structures
Cardiovascular
Cell
Cognitive
Dermatological
Endocrine
Environment
Environmental
Immunological
Metabolic
Mortality
Neurological
Nutritional
Ophthalmological
Psychiatric
Reproduction
Respiratory
Skeletal

9F

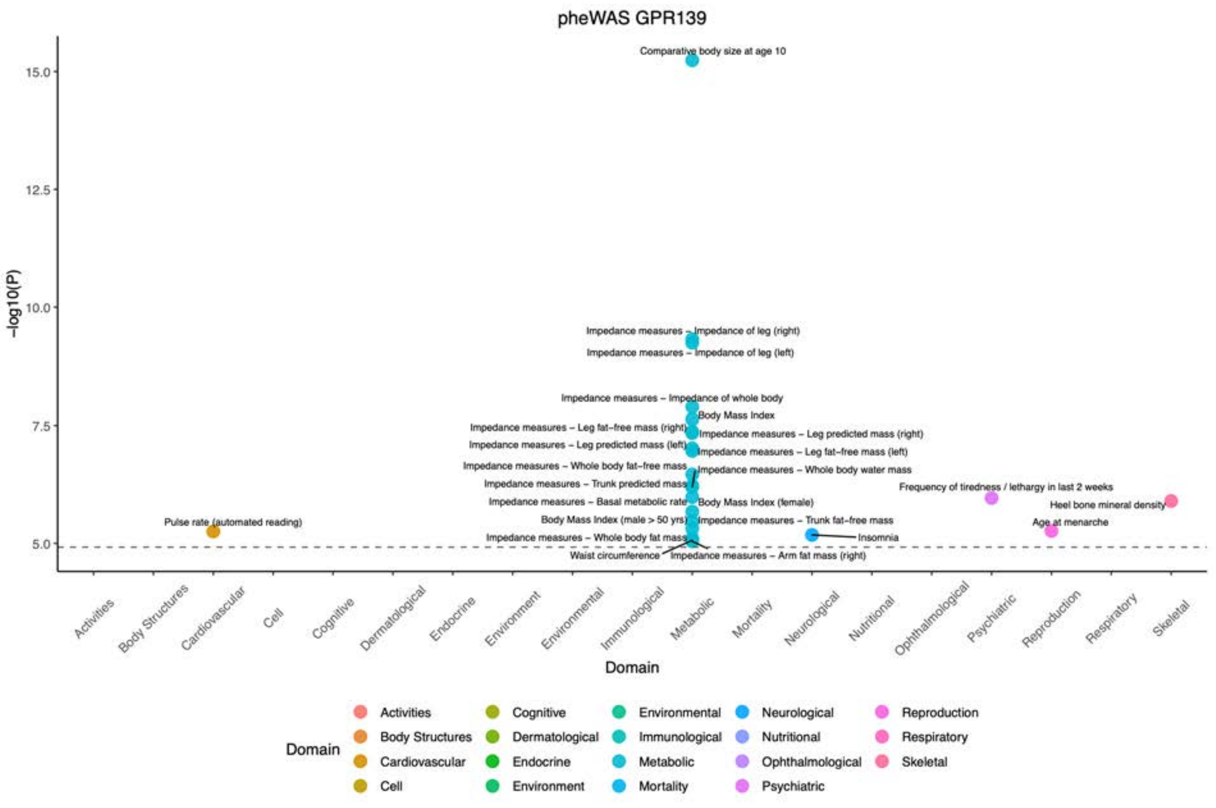
pheWAS GPR139
Comparative body size at age 10
Impedance measures – Impedance of leg (right)
Impedance measures – Impedance of leg (left)
Impedance measures – Impedance of whole body
Body Mass Index
Impedance measures – Leg fat–free mass (right)
Impedance measures – Leg predicted mass (right)
Impedance measures – Leg predicted mass (left)
Impedance measures – Leg fat–free mass (left)
Impedance measures – Whole body fat–free mass
Impedance measures – Whole body water mass
Impedance measures – Trunk predicted mass
Frequency of tiredness / lethargy in last 2 weeks
Impedance measures – Basal metabolic rate
Body Mass Index (female)
Heel bone mineral density
Pulse rate (automated reading)
Body Mass Index (male > 50 yrs)
Impedance measures – Trunk fat–free mass
Age at menarche
Impedance measures – Whole body fat mass
Insomnia
Waist circumference
Impedance measures – Arm fat mass (right)
15.0
12.5
10.0
7.5
5.0
-log10(P)
Activities
Body Structures
Cardiovascular
Cell
Cognitive
Dermatological
Endocrine
Environment
Environmental
Immunological
Metabolic
Mortality
Neurological
Nutritional
Ophthalmological
Psychiatric
Reproduction
Respiratory
Skeletal
Domain
Domain
Activities
Body Structures
Cardiovascular
Cell
Cognitive
Dermatological
Endocrine
Environment
Environmental
Immunological
Metabolic
Mortality
Neurological
Nutritional
Ophthalmological
Psychiatric
Reproduction
Respiratory
Skeletal

9G

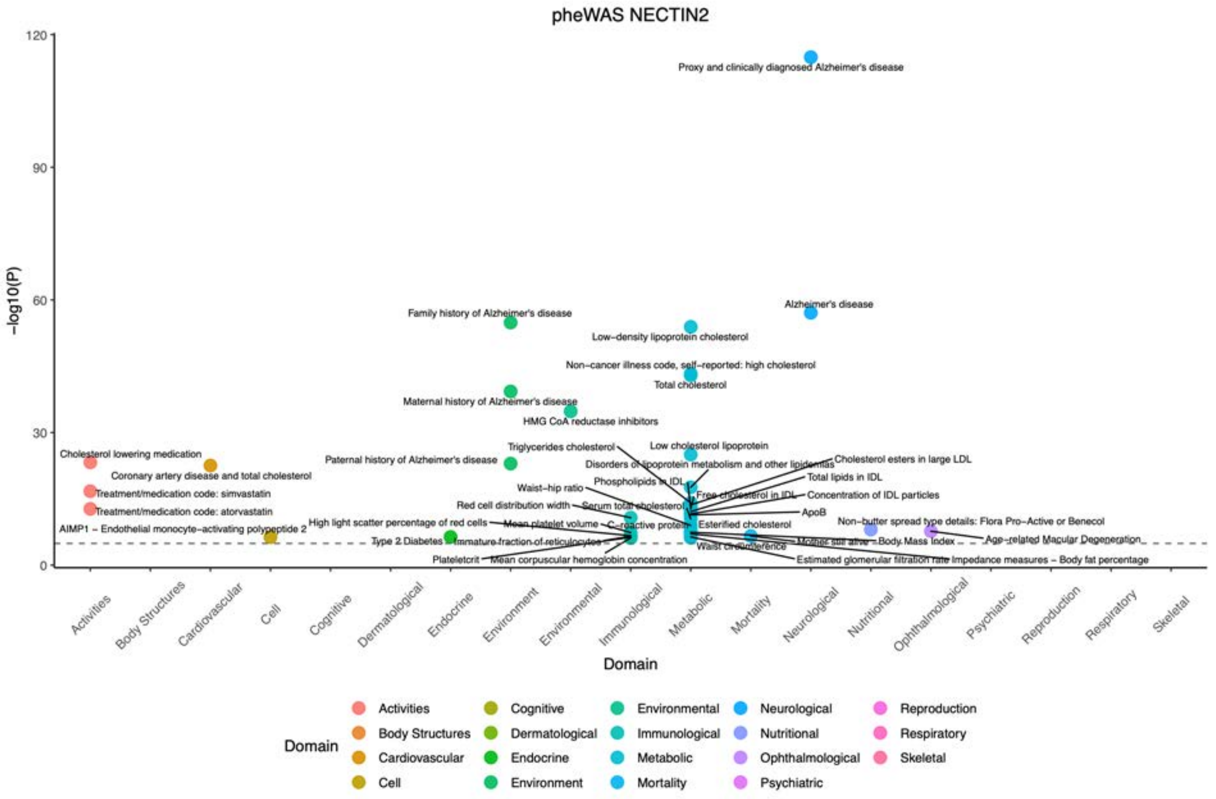

pheWAS NECTIN2
−log10(P)
Domain
Proxy and clinically diagnosed Alzheimer's disease
Alzheimer's disease
Family history of Alzheimer's disease
Low−density lipoprotein cholesterol
Non−cancer illness code, self−reported: high cholesterol
Total cholesterol
Maternal history of Alzheimer's disease
HMG CoA reductase inhibitors
Cholesterol lowering medication
Coronary artery disease and total cholesterol
Treatment/medication code: simvastatin
Treatment/medication code: atorvastatin
AIMP1 − Endothelial monocyte−activating polypeptide 2
Paternal history of Alzheimer's disease
Triglycerides cholesterol
Low cholesterol lipoprotein
Disorders of lipoprotein metabolism and other lipidemias
Cholesterol esters in large LDL
Total lipids in IDL
Phospholipids in IDL
Waist−hip ratio
Free cholesterol in IDL
Concentration of IDL particles
Red cell distribution width
Serum total cholesterol
ApoB
High light scatter percentage of red cells
Mean platelet volume
C−reactive protein
Esterified cholesterol
Non−butter spread type details: Flora Pro−Active or Benecol
Type 2 Diabetes
Immature fraction of reticulocytes
Waist circumference
Mother still alive
Body Mass Index
Age−related Macular Degeneration
Plateletcrit
Mean corpuscular hemoglobin concentration
Estimated glomerular filtration rate
Impedance measures − Body fat percentage
Activities
Body Structures
Cardiovascular
Cell
Cognitive
Dermatological
Endocrine
Environment
Environmental
Immunological
Metabolic
Mortality
Neurological
Nutritional
Ophthalmological
Psychiatric
Reproduction
Respiratory
Skeletal


9H

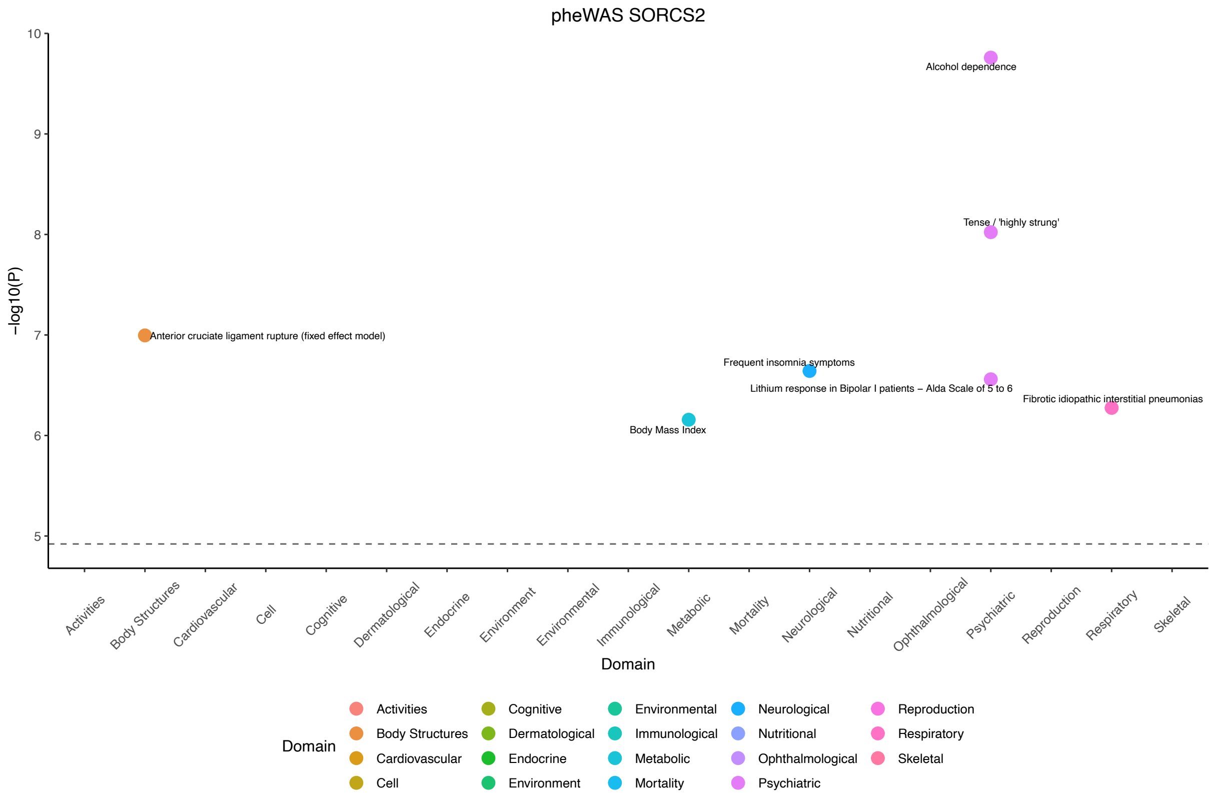

pheWAS SORCS2
−log10(P)
Domain
Alcohol dependence
Tense / 'highly strung'
Anterior cruciate ligament rupture (fixed effect model)
Frequent insomnia symptoms
Lithium response in Bipolar I patients − Alda Scale of 5 to 6
Fibrotic idiopathic interstitial pneumonias
Body Mass Index
Activities
Body Structures
Cardiovascular
Cell
Cognitive
Dermatological
Endocrine
Environment
Environmental
Immunological
Metabolic
Mortality
Neurological
Nutritional
Ophthalmological
Psychiatric
Reproduction
Respiratory
Skeletal

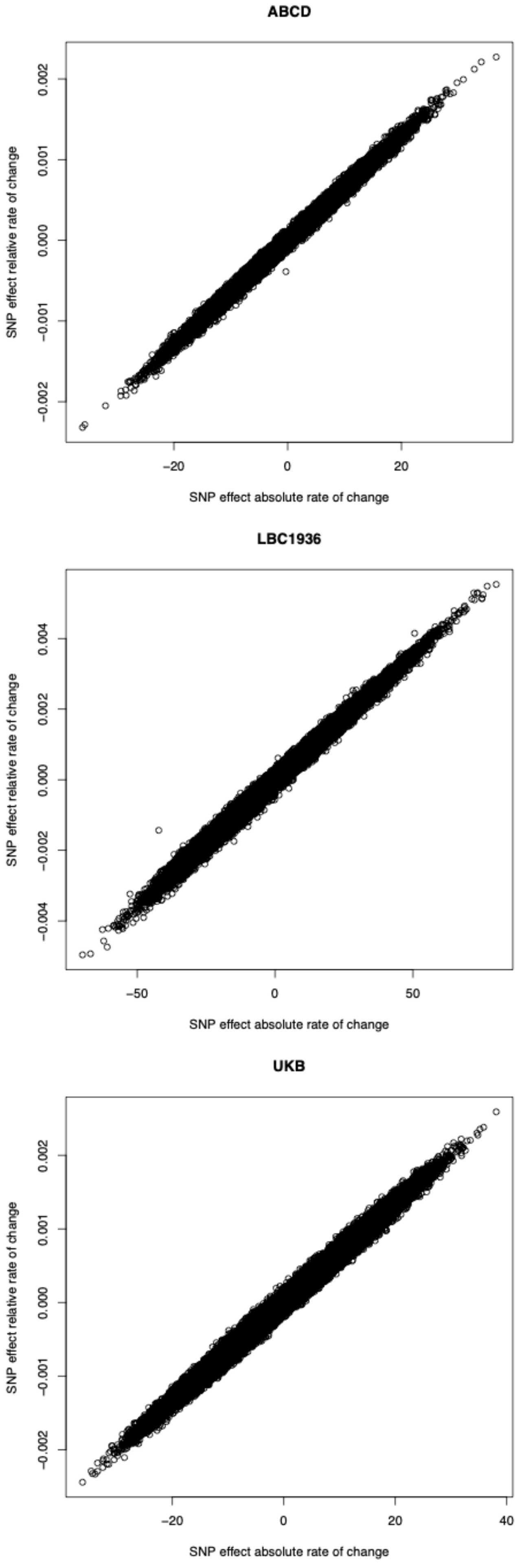
ABCD
SNP effect relative rate of change
SNP effect absolute rate of change
LBC1936
SNP effect relative rate of change
SNP effect absolute rate of change
UKB
SNP effect relative rate of change
SNP effect absolute rate of change

11

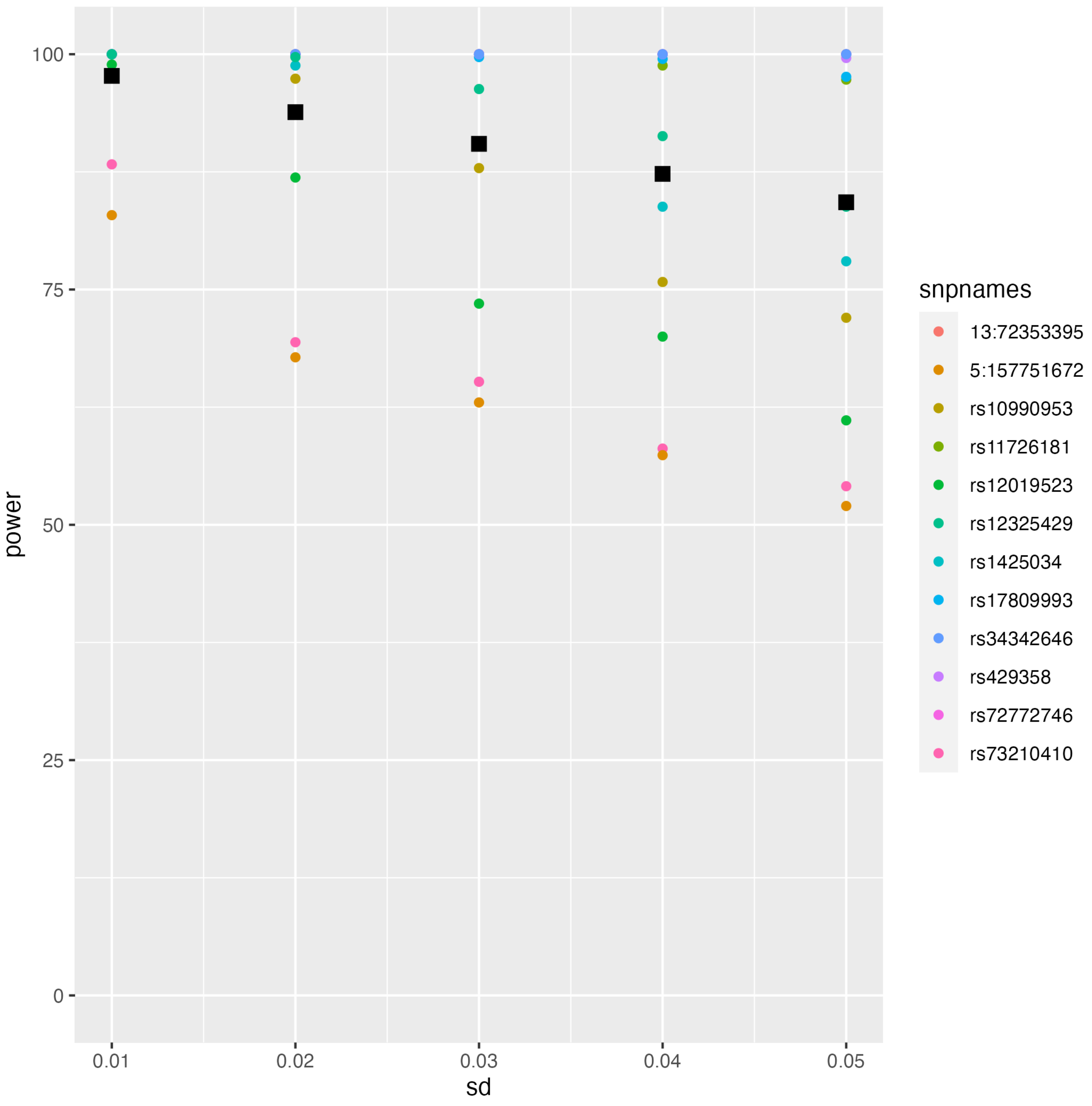

100
75
50
25
0
power
0.01
0.02
0.03
0.04
0.05
sd
snpnames
13:72353395
5:157751672
rs10990953
rs11726181
rs12019523
rs12325429
rs1425034
rs17809993
rs34342646
rs429358
rs72772746
rs73210410

### QQ plots Amygdala

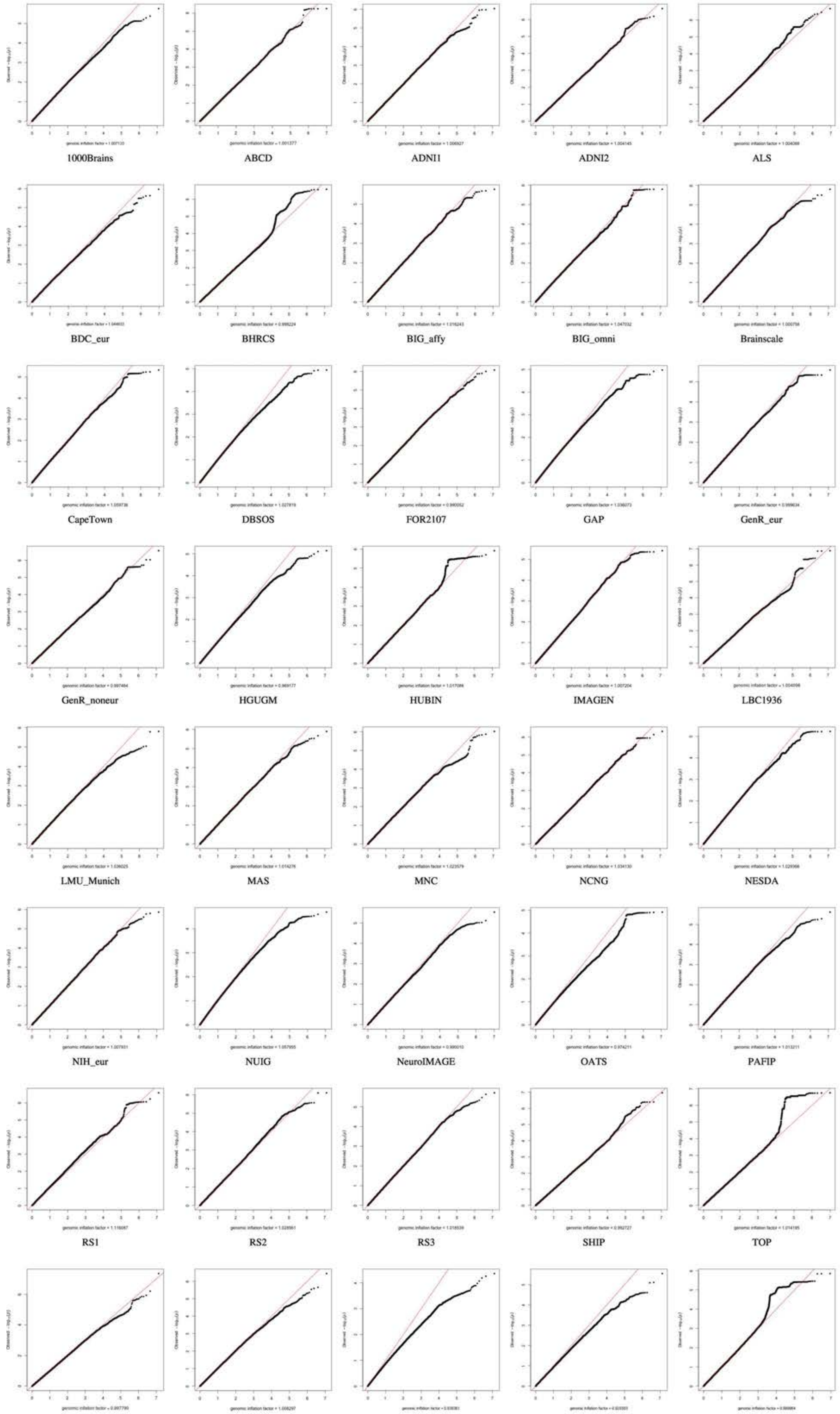

### QQ plots Caudate

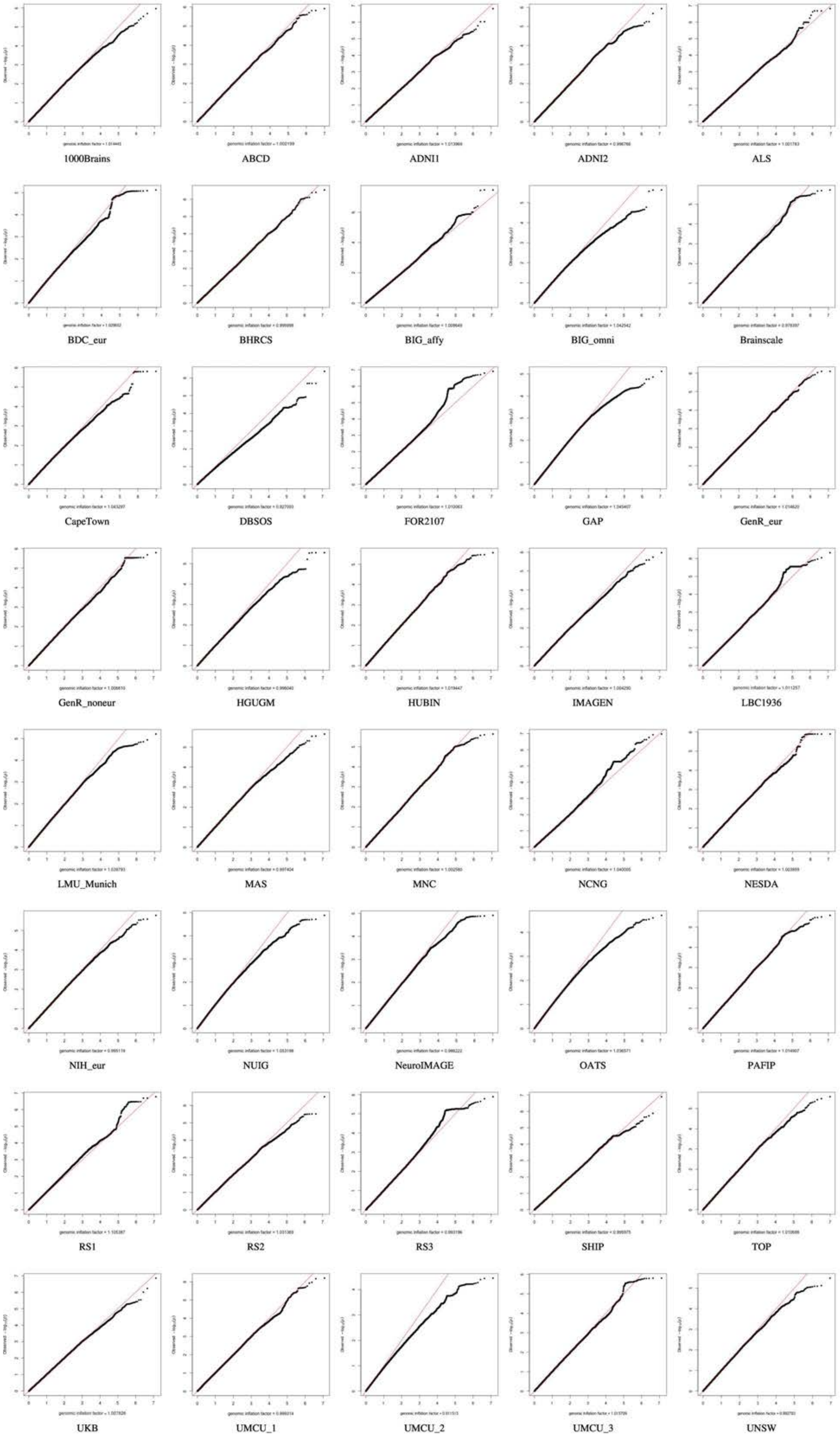

12

### QQ plots Cerebellum Cortex

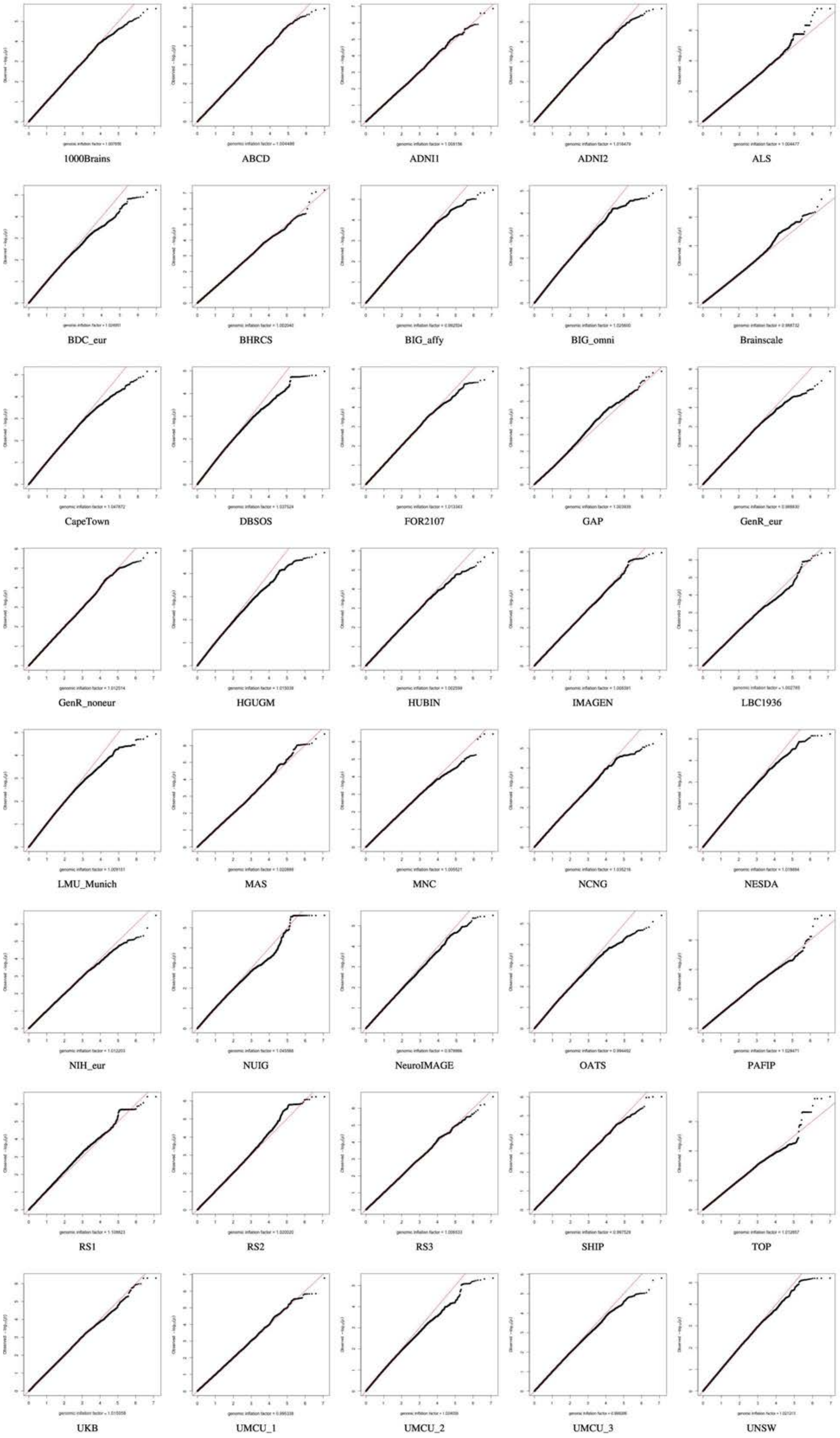

12

### QQ plots Cerebellum White Matter

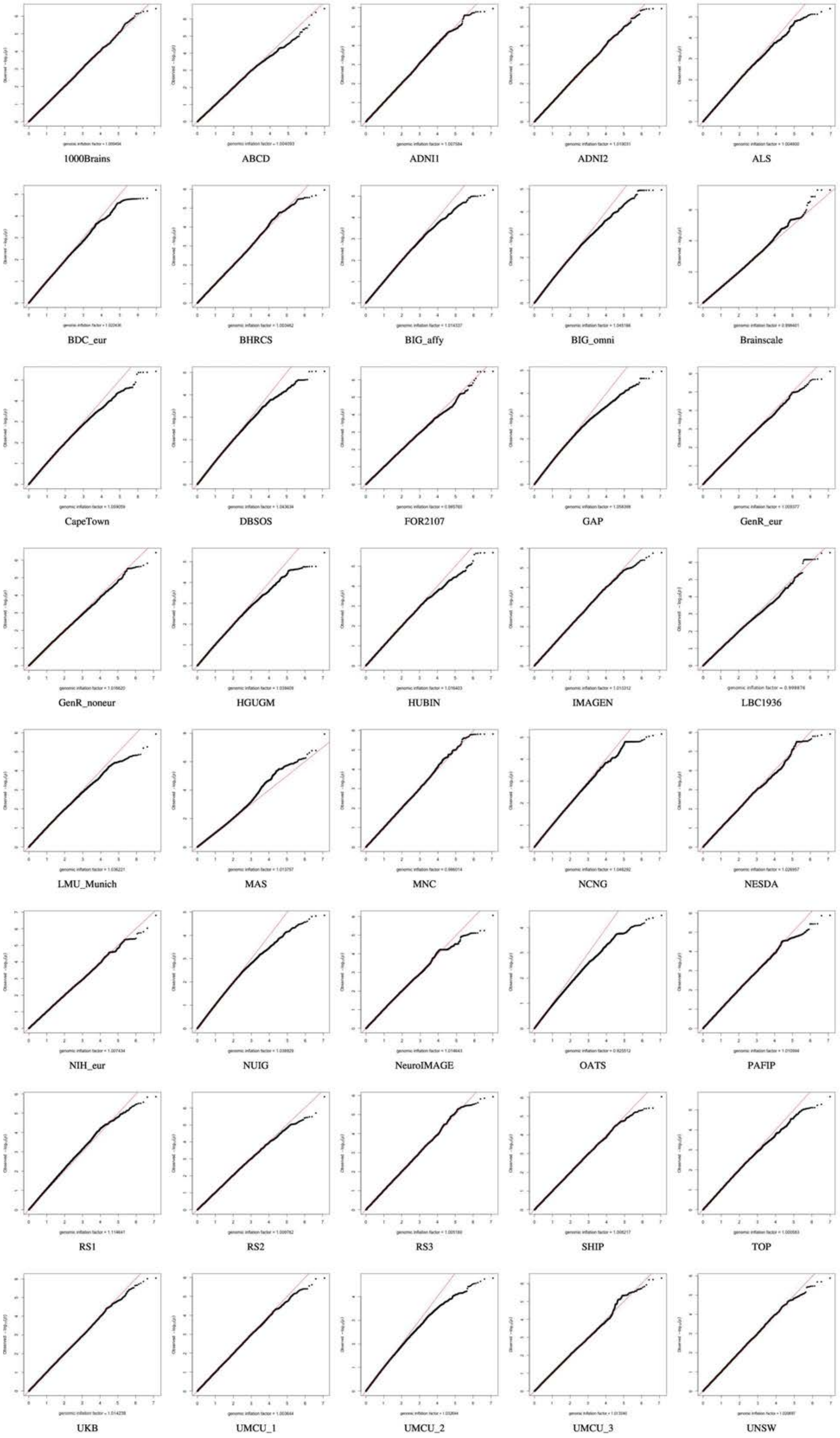

### QQ plots Cerebral White Matter

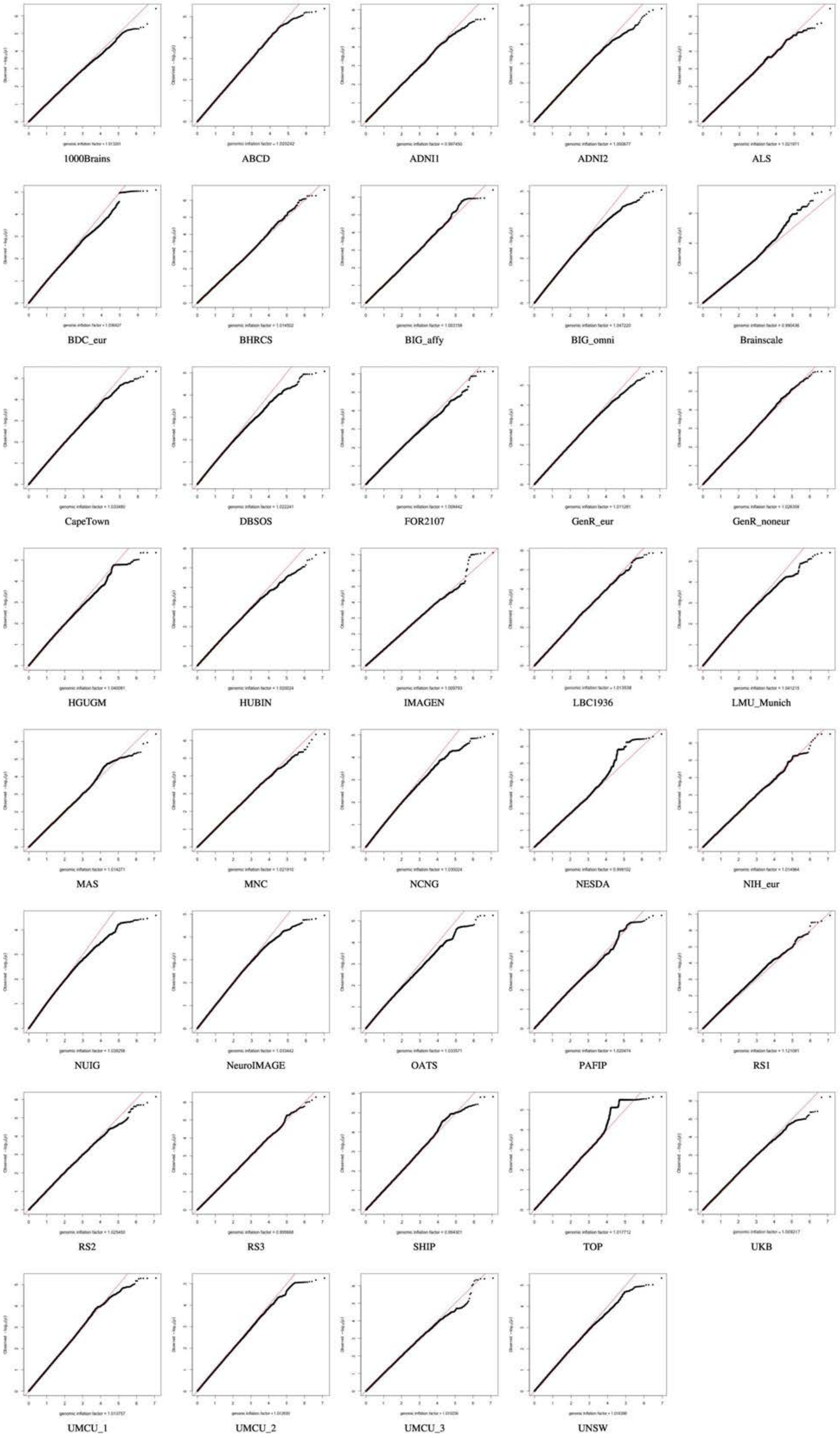

12

QQ plots Cortex

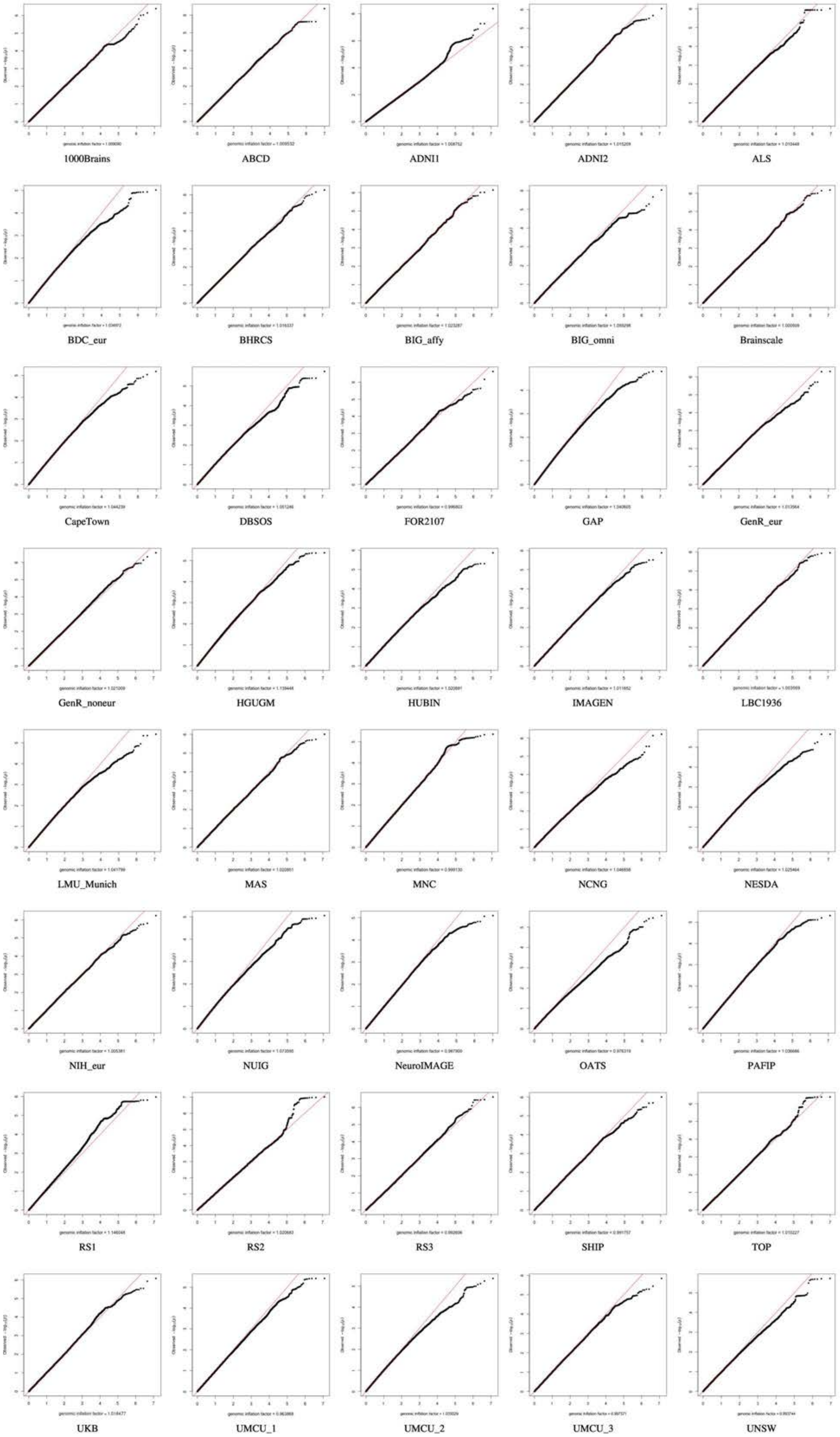

12

### QQ plots Cortical Thickness

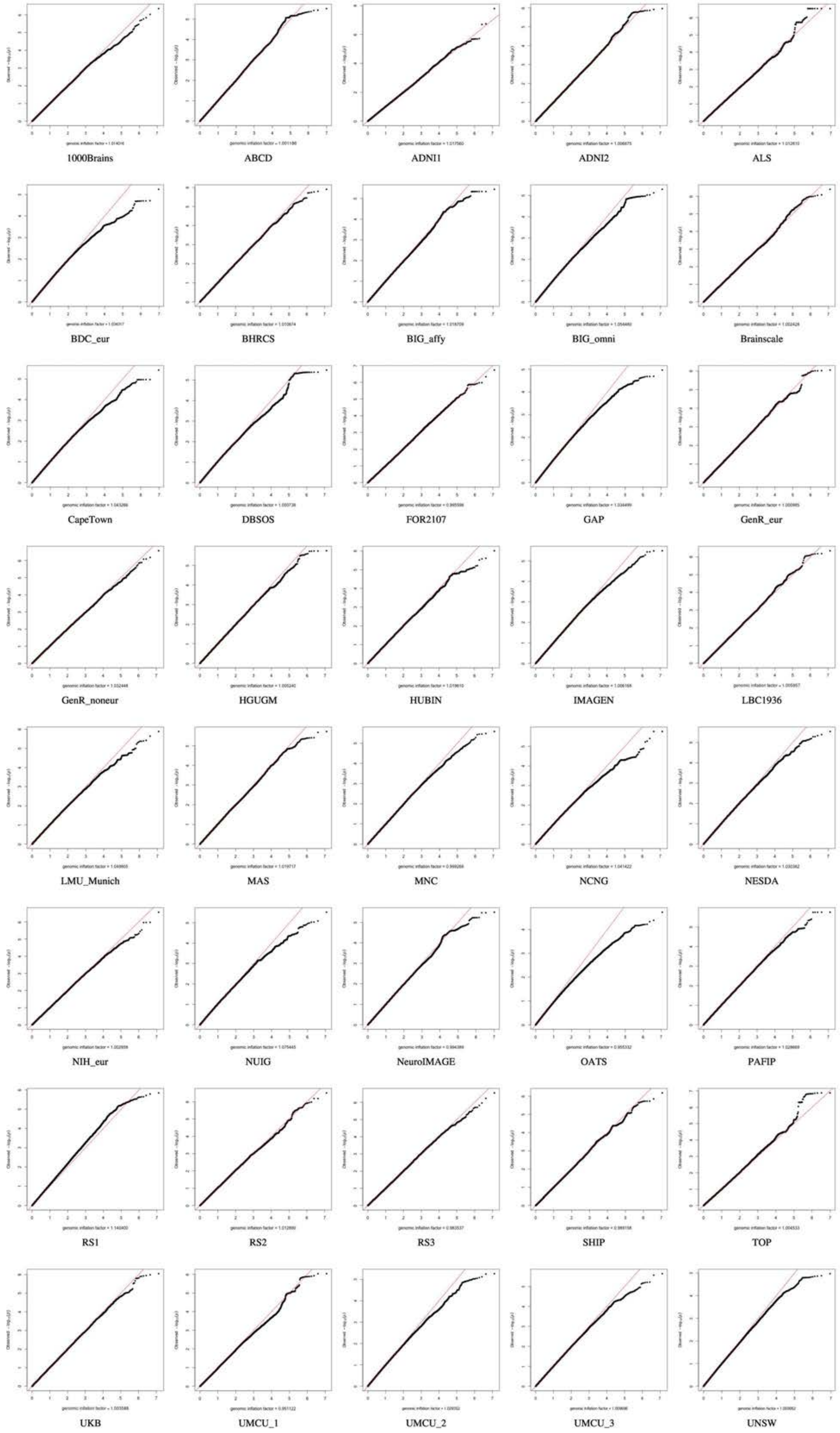

12

### QQ plots Hippocampus

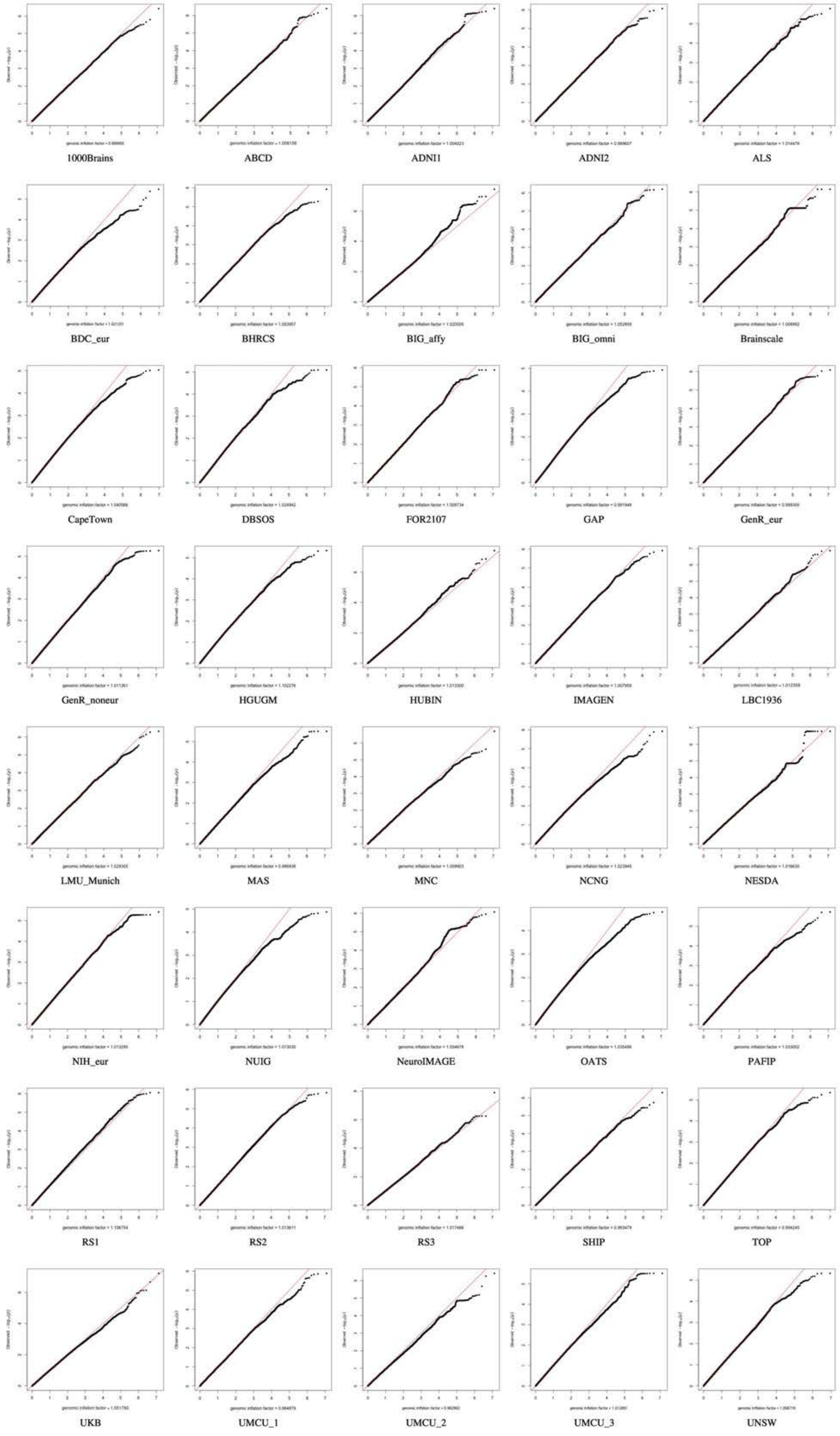

QQ plots Lateral Ventricles

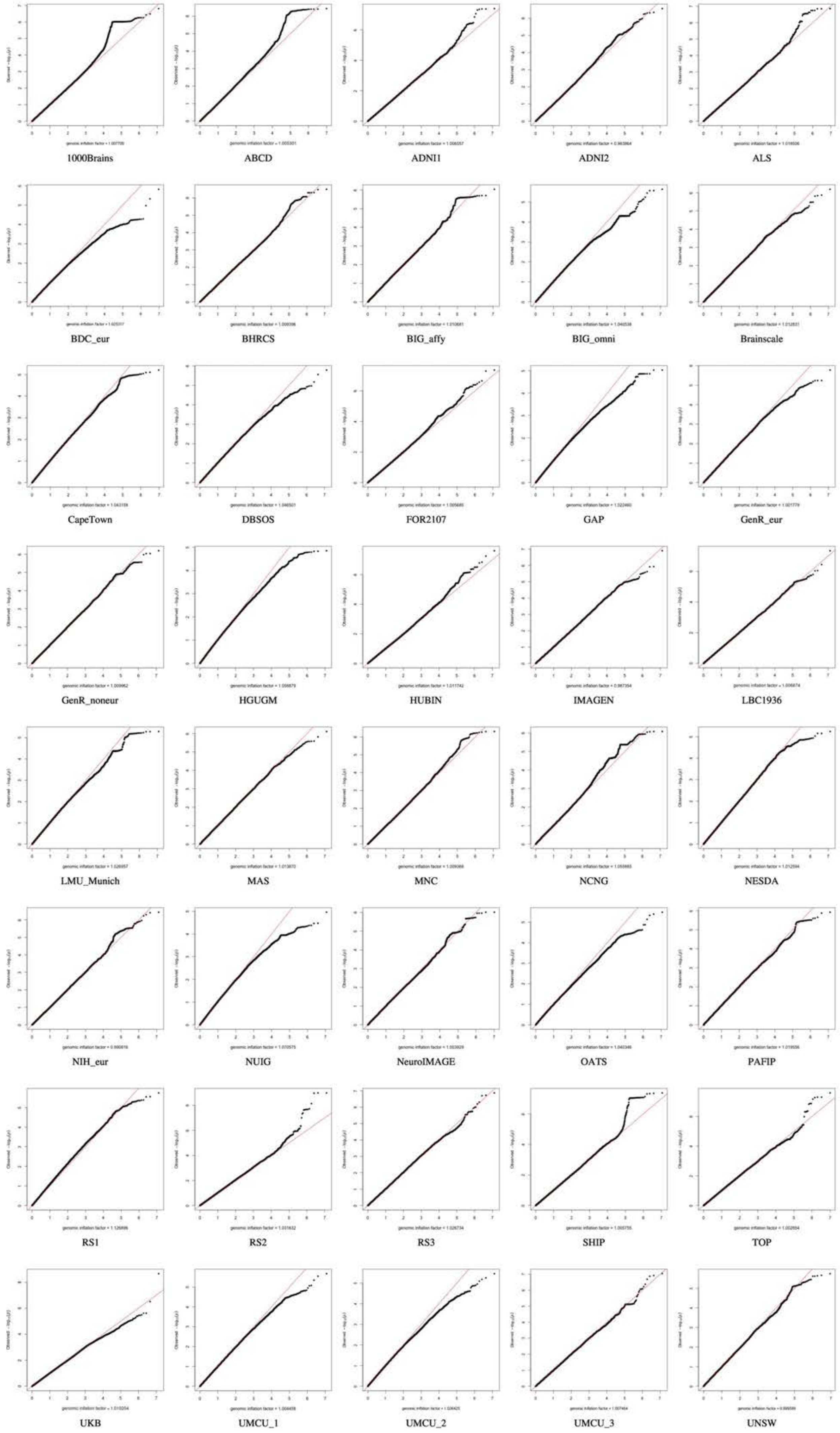

### QQ plots Nucleus Accumbens

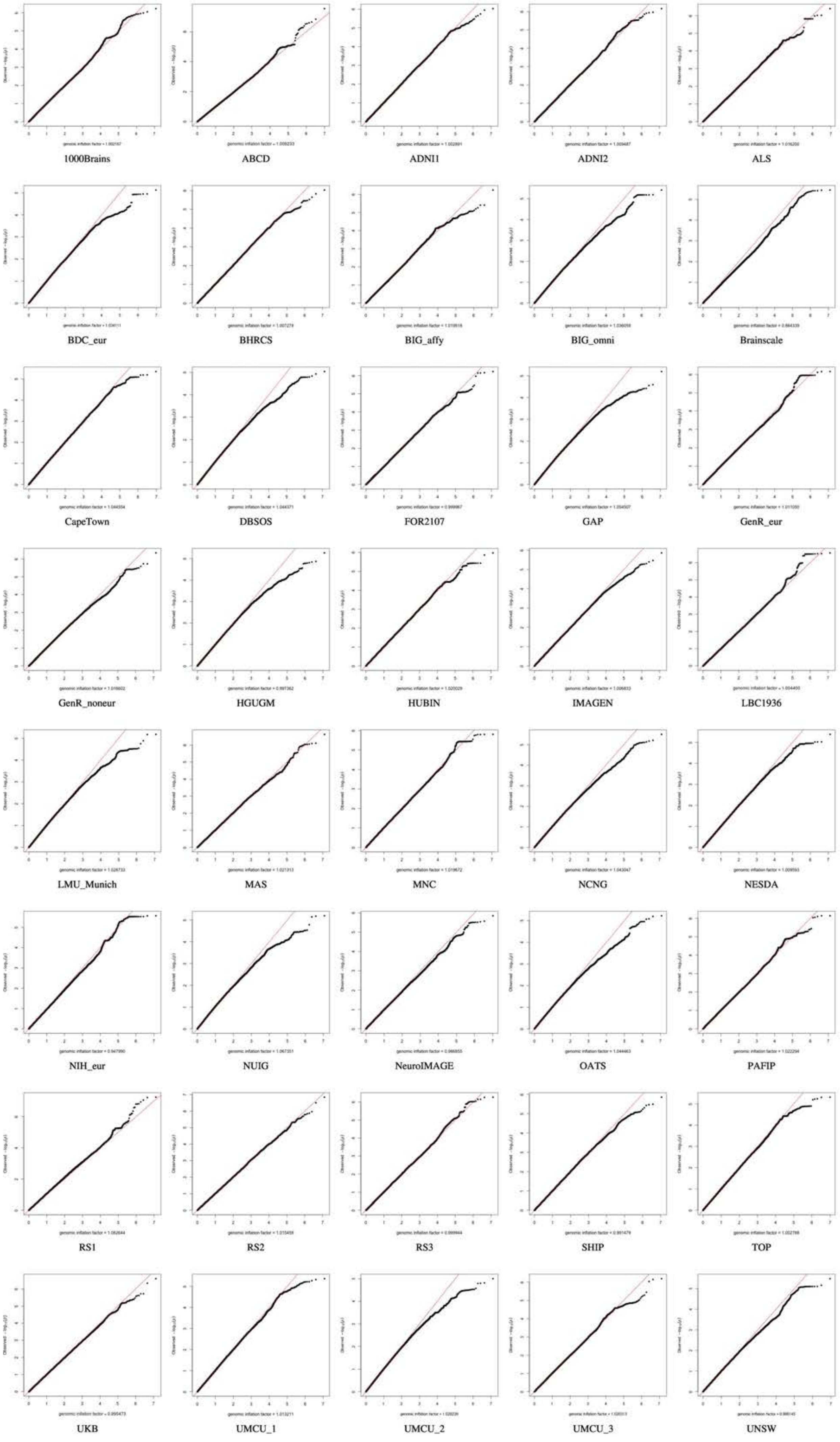

### QQ plots Pallidum

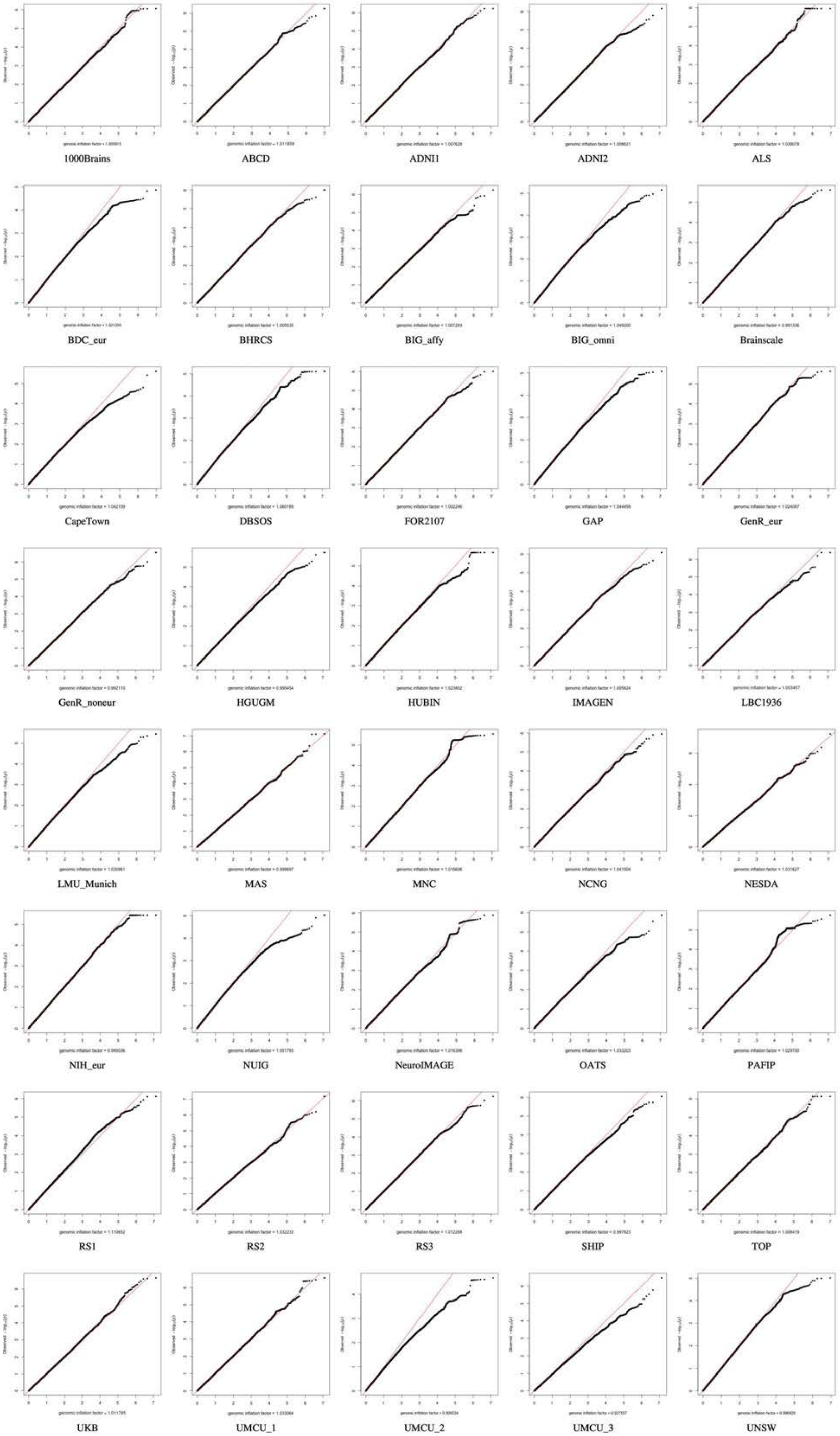

12

### QQ plots Putamen

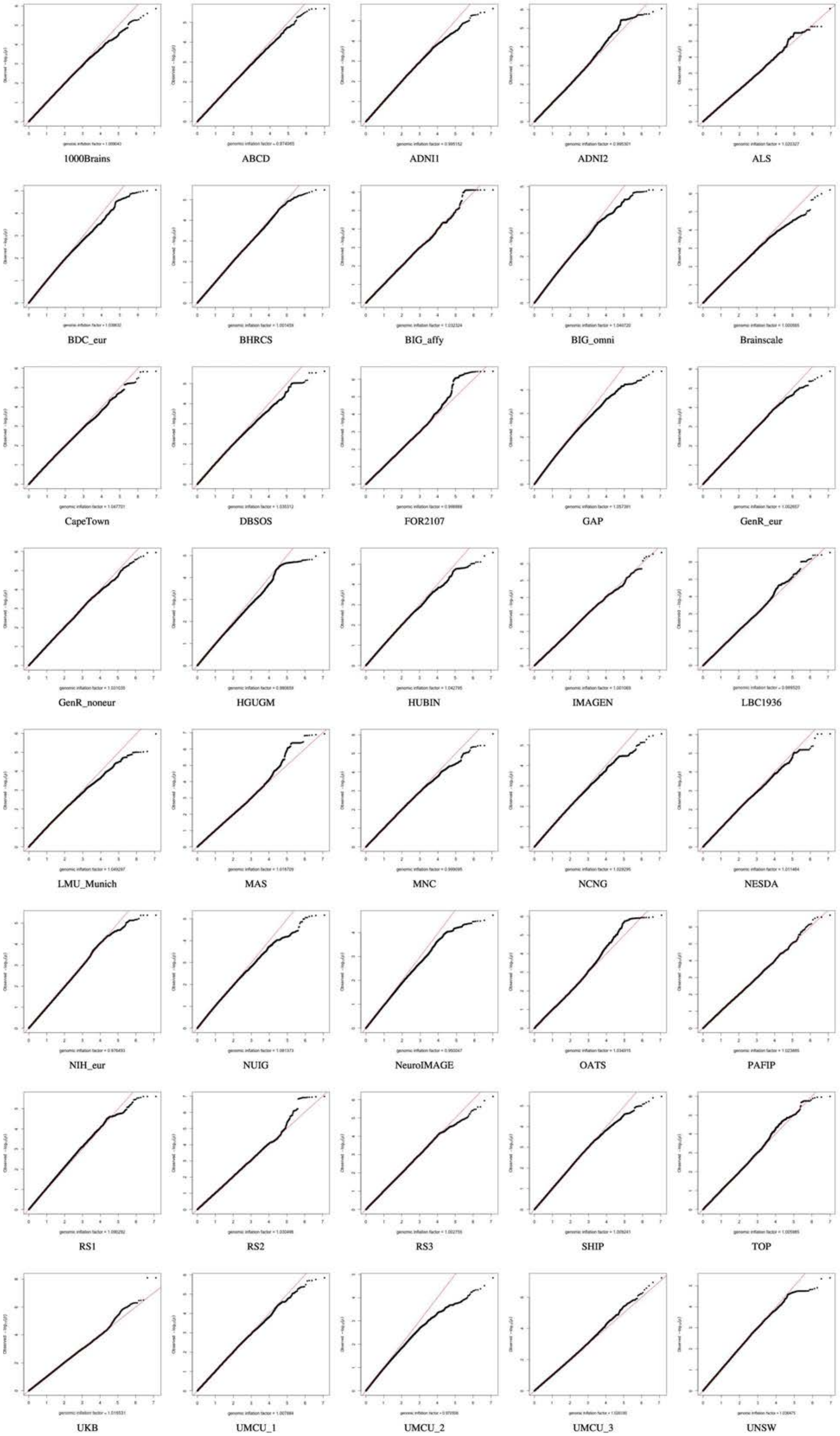

12

### QQ plots Surface Area

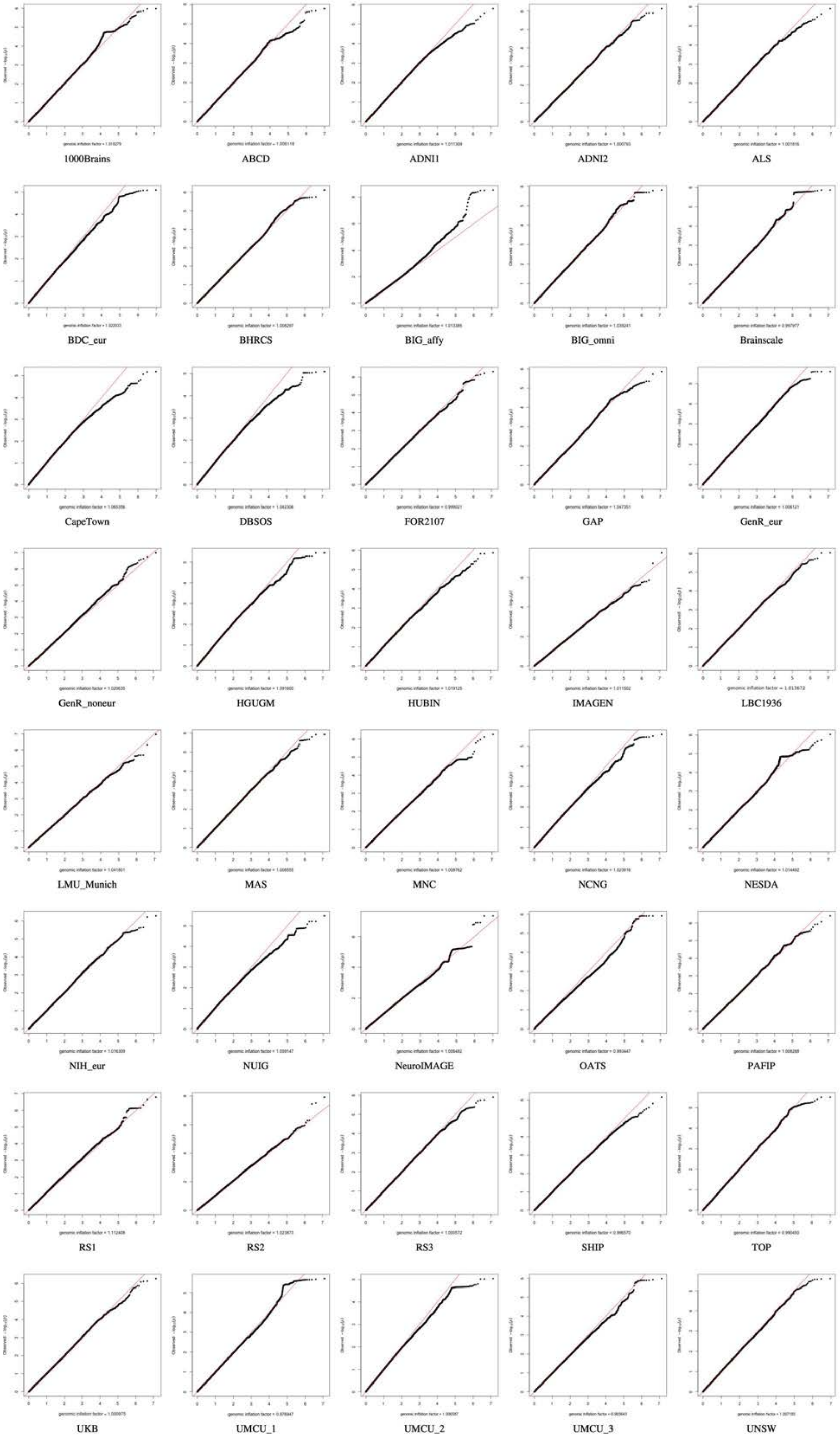

### QQ plots Thalamus

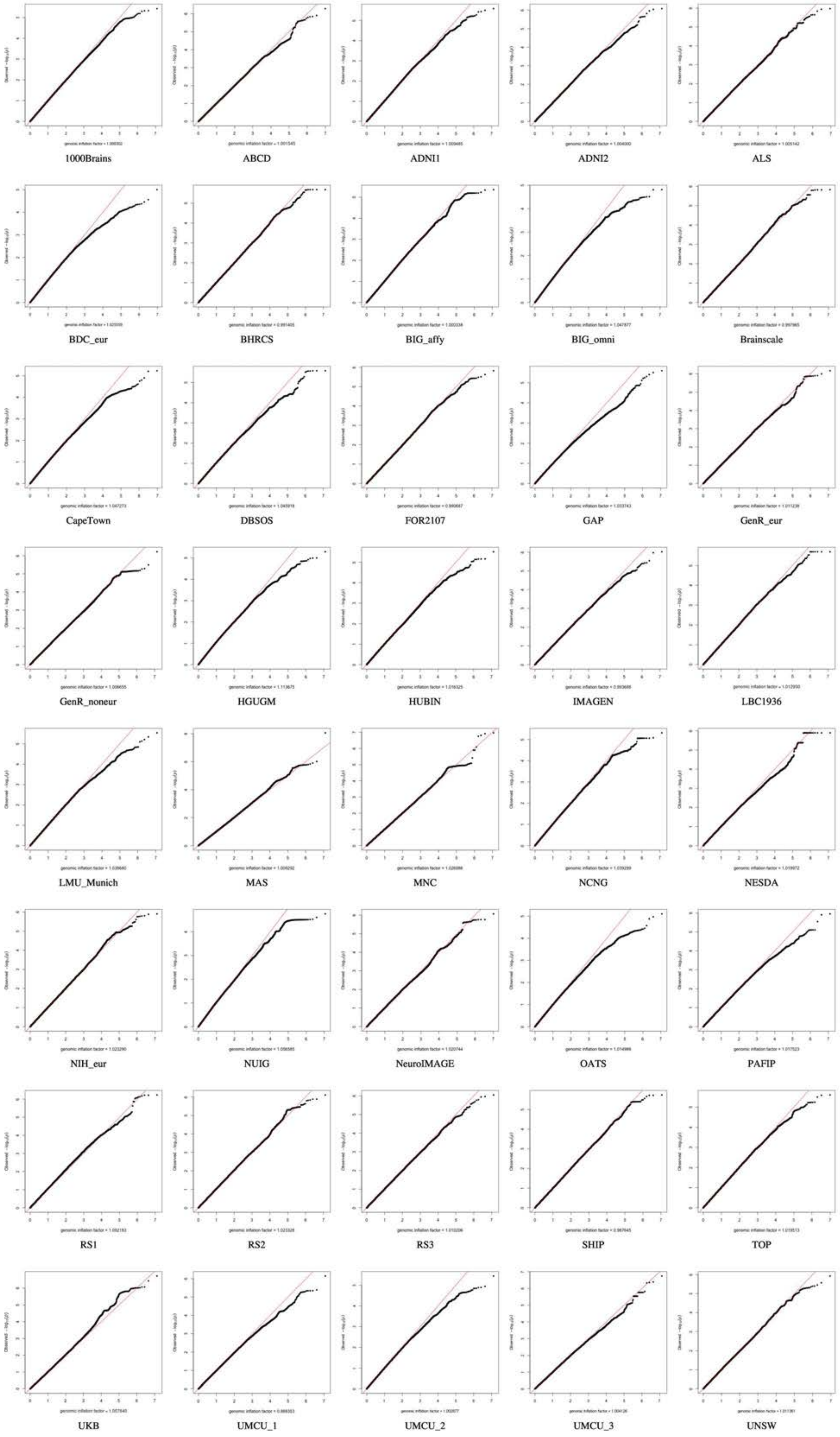

### QQ plots Total Brain

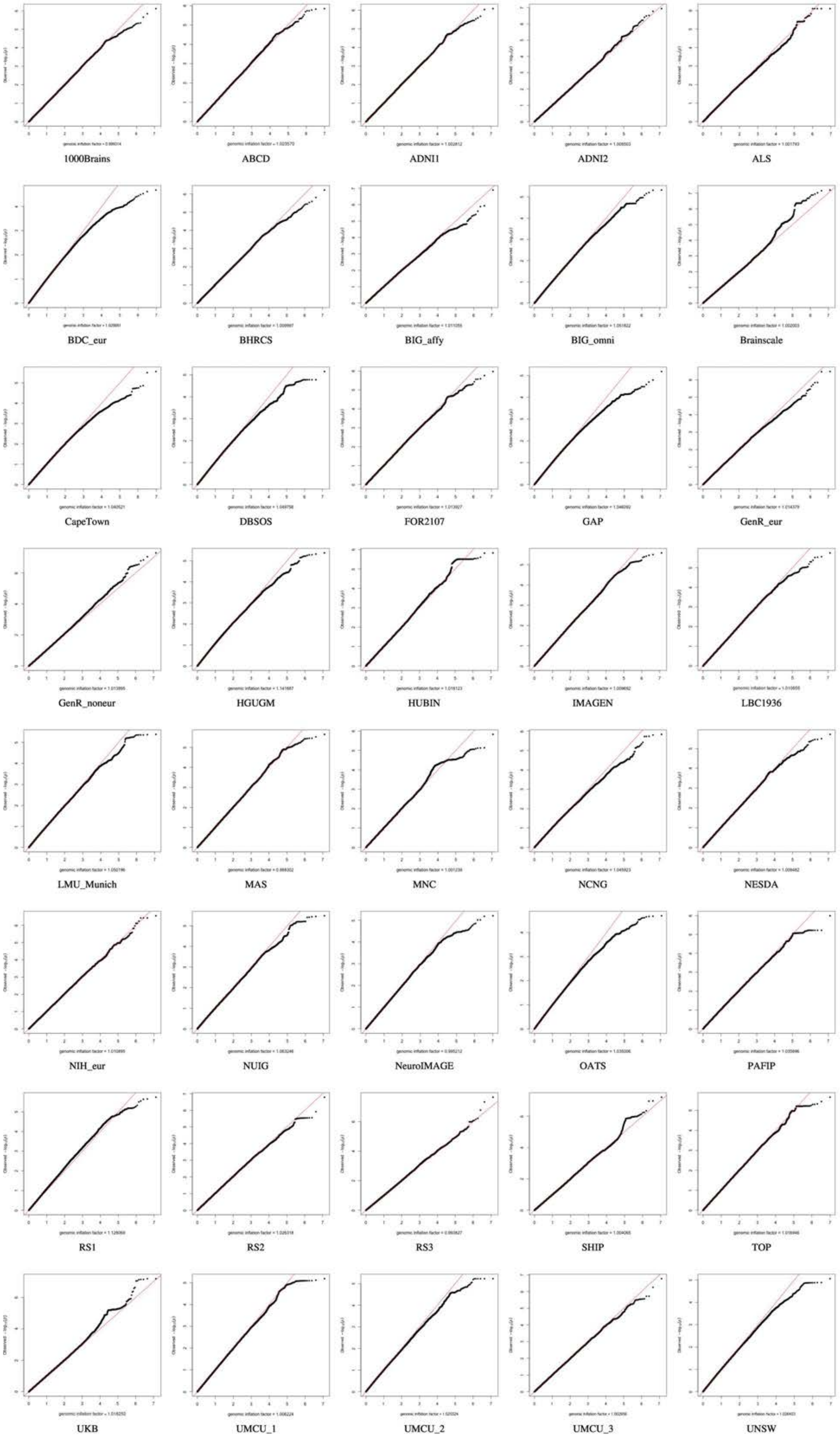
